# Comparing the effects of host species richness on different metrics of disease

**DOI:** 10.1101/2022.11.01.514730

**Authors:** Michael H. Cortez

## Abstract

Changes in host species richness can alter infection risk and disease levels in communities. I review theoretical predictions for direct and environmental transmission pathogens about the effects of host additions (or removals) on three common disease metrics: the pathogen community reproduction number (ℛ_0_) and the proportion (i.e., infection prevalence) and density of infected individuals in a focal host. To help explain why predictions differ between studies and between metrics, I analyze an SIR-type model of an environmentally transmitted pathogen and multiple host species that compete for resources. I use local sensitivity analysis to explore and compare how variation in an added host’s ability to transmit a pathogen, its density, and the pathogen transmission mechanism affect each disease metric. I find that if an added host species is a weak interspecific competitor, then each metric typically increases or decreases, respectively, when the added host has a high or low ability to transmit the pathogen. However, if the added host species is a strong interspecific competitor, then the response of each metric can be strengthened, weakened, or reversed depending on how the competitive effects of the added host propagate through the community and alter the densities of all host species in the community. The three metrics often respond in the same direction, but the metrics can respond in different directions for three reasons: (1) strong density-mediated feedbacks driven by high disease-induced mortality; (2) host additions or removals cause large changes in focal host density via competition or disease-induced mortality; and (3) the metrics use different quantities to define a host’s ability to transmit disease: the quantities for infection prevalence and infected density depend on the instantaneous production rates of infectious propagules whereas the quantities for ℛ_0_ depend on lifetime production of infectious propagules. This study provides a framework that unifies prior theoretical studies and identifies rules governing the context-dependent relationships between host species richness and the three metrics of disease.

## 1 Introduction

The gain or loss of host species from a community can alter levels of disease and disease risk for focal host populations. In empirical systems, increased host species richness can decrease or increase disease risk and levels of disease (Randolph and Dobson, 2012; Salkeld et al., 2013; Wood et al., 2014; Civitello et al., 2015; Ostfeld and Keesing, 2017; Rohr et al., 2019). Theoretical studies have used mathematical models to explore potential mechanisms driving this variation in host species-disease relationships (Dobson, 2004; Joseph et al., 2013; Mihaljevic et al., 2014; Faust et al., 2017). However, because the models make different assumptions about the host and pathogen species and use different metrics of disease, it remains unclear what biological and mathematical assumptions are responsible for the variation and differences in predictions (Buhnerkempe et al., 2015; Johnson et al., 2015; Rohr et al., 2019). Here, I present a framework that helps explain and extend these prior predictions, accounting for the effects of the characteristics of the host and pathogen species and the metric of disease.

Across empirical systems, increased host species richness often decreases levels of disease and disease risk (Civitello et al., 2015), but it can increase disease levels in many systems as well (Wood et al., 2014). The effects of altered host species richness are likely affected by which specific species are already present in the community and which are added to or removed from the community (Wood et al., 2016; Randolph and Dobson, 2012). For example, infection prevalence of the fungus *Batrachochytrium dendrobatidis* in three frog genera depends on which specific genera are present in a community: *Gastrophryne* species decrease, *Bufo* species increase, and *Hyla* species have no effect on infection prevalence in the other genera. Similarly, in laboratory experiments, the prevalence of fugal infections in *Daphnia dentifera* increases when the alternative host *D. lumholtzi* is present but decreases when the alternative host *Ceriodaphnia dubia* is present. In total, empirical studies show that relationships between host species richness and disease are likely to be context-dependent (Salkeld et al., 2013).

Despite the observed variation in how disease levels respond to changes in host species richness, we have a limited understanding of the mechanisms causing that variation and their context-dependent effects. This has led to calls for theory on host richness-disease relationships and, more generally, host biodiversity-disease relationships (Buhnerkempe et al., 2015; Johnson et al., 2015; Halsey, 2019; Rohr et al., 2019). Current theory predicts that responses to host additions and removals are affected by host competence (the ability to transmit the pathogen to susceptible individuals), interspecific host competition, and the pathogen transmission mechanism (Dobson, 2004; Rudolf and Antonovics, 2005; Joseph et al., 2013; Mihaljevic et al., 2014; O’Regan et al., 2015; Strauss et al., 2015; Faust et al., 2017; Cortez and Duffy, 2020; Cortez, 2021); specific predictions are discussed in the next section. While this body of work provides insight into how host species richness can affect levels of disease, general predictions remain unclear because the studies make different assumptions about the characteristics of the host and pathogen species.

Another limitation of current theory is that modeling studies use different metrics of disease and it is unclear how the choice of disease metric influences predictions. Three metrics of disease commonly used in empirical and theoretical studies are: (i) the pathogen’s community reproduction number 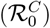, which is the average number of new infections produced by a single infected individual in an otherwise susceptible community; (ii) the proportion of infected individuals in a focal population (hereafter, focal host infection prevalence; *I/N*); and (iii) the density of infected individuals in a focal population (hereafter, focal host infected density; *I*). I note that decreased and increased disease risk in response to increased host species richness are often referred to as amplification and dilution, respectively (Keesing et al., 2006; Rohr et al., 2019), but to avoid confusion about which disease metric is being referenced, I will not use that terminology in this study. The three metrics provide different information about disease dynamics and disease risk. Specifically, 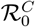 provides a measure of the risk of an outbreak and how fast an outbreak will grow. In prevalence and infected density both provide measures of disease burden for a focal population. In addition, infection prevalence measures disease risk for a focal host population whereas infected density measures the potential risk for a pathogen to spread or spillover into another population or species (Roche et al., 2012; Wood and Lafferty, 2013). Theoretical studies predict that changes in host species richness can have effects of different signs on the three metrics (Roche et al., 2012; Roberts and Heesterbeek, 2018). As an empirical example, infection prevalence and infected density in *Daphnia lumholtzi* both increase when *D. dentifera* is present. In contrast, fungal infection prevalence in *Daphnia dentifera* increases but infected density decreases when *D. lumholtzi* is present (Searle et al., 2016). Thus, the metrics can respond in the same or opposite directions to changes in host species richness, but it is currently unknown when and why the metrics respond similarly versus differently.

In this study, I review the theoretical predictions about the relationships between host species richness and the three metrics of disease, focusing on pathogens with direct transmission (transmission via host-host contact) and environmental transmission (transmission via infectious propagules that are shed into an environment). Then, I present a multihost environmental transmission model and use it to explore the factors influencing the relationships between each metric of disease and host species richness. To do this, I use sensitivity analysis to calculate how each metric responds to variation in components of host competence, host density, and the pathogen transmission mode. This study extends prior work (Roberts and Heesterbeek, 2018; Cortez and Duffy, 2021; Cortez, 2021) using the same approach by computing the relationships for all three metrics and identifying when and why the three metrics can have relationships with the same or different signs. My results provide a framework that unifies prior theoretical studies and explains the biological and mathematical assumptions that are responsible for the variation in prior theoretical predictions. That in turn helps identify the rules and biological mechanisms shaping the context-dependent relationships between host species richness and disease in empirical systems.

## 2 Current theory on how host species richness affects disease

Many studies have reviewed and debated the empirical evidence for positive and negative effects of host species richness on levels of disease (Keesing et al., 2010; Randolph and Dobson, 2012; Salkeld et al., 2013; Wood et al., 2014; Civitello et al., 2015; Ostfeld and Keesing, 2017; Rohr et al., 2019). Here, I review current theory for directly and environmentally transmitted pathogens.

The foundational work in Keesing et al. (2006) predicts that disease risk in a focal host increases when host additions or removals increase host encounter rates (*α*), increase per exposure transmission rates (Δ), increase susceptible host densities (*S*), decrease mortality rates of infected individuals (*μ*), and decrease host recovery rates (*ν*). These predictions are derived from an equation describing changes in infected host density,

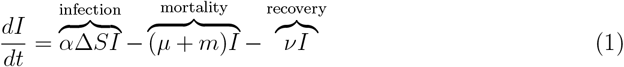

where *S* and *I* are susceptible and infected host densities, respectively, and *m* is the non-disease mortality rate. The predicted responses are based on the idea that increases and decreases in the growth rate of infected individuals (*dI/dt*) increase and decrease disease risk, respectively. While the five mechanisms are defined for direct transmission pathogens (i.e., transmission via host-host contact), analogues exist for environmental transmission pathogens (i.e., transmission via infectious propagules shed into the environment; Johnson and Thieltges 2010) and vector-borne pathogens (Ostfeld and Keesing, 2000; Keesing et al., 2006). A key limitation of these predictions is that they focus on changes in the growth rate of infected individuals and do not directly address other quantities measured in populations (e.g., infection prevalence and infected density).

Other theoretical studies have explored how the presence and absence of host species affect the pathogen’s community reproduction number 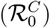 and infection prevalence (*I*^*^*/N*^*^, where *N*^*^ is total host density) and infected density (*I*^*^) of a focal host at equilibrium.For density-dependent direct transmission (DDDT) models (host-host contact rates are proportional to the density of infected individuals), host additions often increase 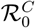 and focal host infection prevalence (Dobson, 2004; Rudolf and Antonovics, 2005; Faust et al., 2017), but both metrics can decrease if there is strong interspecific competition (Joseph et al., 2013; Mihaljevic et al., 2014; O’Regan et al., 2015; Cortez, 2021). For frequency-dependent direct transmission (FDDT) models (host-host contact rates are proportional to the frequency of infected individuals), host additions often decrease 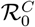 and focal host infection prevalence (Dobson, 2004; Rudolf and Antonovics, 2005; Joseph et al., 2013; Mihaljevic et al., 2014), but both metrics can increase if added host species have sufficiently high intraspecific and interspecific transmission rates (O’Regan et al., 2015; Faust et al., 2017). Combined, this suggests that host additions increase 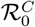 and focal host infection prevalence more often under DDDT than FDDT (Dobson, 2004; Rudolf and Antonovics, 2005; Mihaljevic et al., 2014; Faust et al., 2017).

For environmental transmission (ET) models (transmission via infectious propagules shed into the environment), responses in 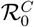 and focal host infection prevalence depend on interspecific host competition and whether host species are net producers (sources) or net removers (sinks) of infectious propagules (Cáceres et al., 2014; Cortez and Duffy, 2021; Cortez, 2021; Espira et al., 2022). In particular, added host species are more likely to increase 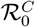 and focal host infection prevalence if they are sources (Cortez, 2021; Espira et al., 2022). Added host species are also more likely to increase focal host infection prevalence if they have stronger competitive effects on sink hosts than on source hosts (Cortez, 2021).

Taken together, current theory points to possible general rules about how host competence, host competition, and the pathogen transmission mechanism affect host species richness-disease relationships. In addition, current theory suggests responses in 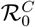 and infection prevalence are similar. This is useful to modelers because 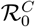 is typically easier to compute from models than infection prevalence. However, current theory is limited in two key ways. First, studies make different assumptions about the number of host species, the strength of interspecific host interactions, and the rates and mechanisms of disease transmission. For example, some FDDT studies (Dobson, 2004; Joseph et al., 2013; Mihaljevic et al., 2014) assume interspecific competition is absent and interspecific transmission rates are lower than intraspecific transmission rates whereas other studies (O’Regan et al., 2015; Faust et al., 2017) assume interspecific competition is present and intraspecific and interspecific transmission rates are comparable. Similarly, some ET studies (Begon and Bowers, 1994; Fenton et al., 2015) assume interspecific competition is absent and all host species are net producers of infectious propagules, whereas others studies (Strauss et al., 2015; Cáceres et al., 2014; Cortez and Duffy, 2021; Espira et al., 2022) assume interspecific competition is present and host species can be net producers or net removers of infectious propagules. Differences in the assumptions about competition and transmission rates or infectious propagule production rates make it difficult to identify which assumptions are responsible for differences in model behavior. Moreover, the difference in transmission mechanisms complicate efforts to find general rules that apply to all pathogens.

The second limitation of current theory is that prior studies have used different metrics of disease and the three metrics listed above can respond differently to host additions and removal (Keesing et al., 2006; Roche et al., 2012; Roberts and Heesterbeek, 2018). Indeed, Keesing et al. (2006) noted that variation in transmission rates (via *α* or Δ) could result in infection prevalence and infected density changing in different directions under some conditions (page 488). The differing responses of the three metrics means that predictions for one metric may not directly apply to a different metric. In addition, because we currently do not understand when and why the metrics respond similarly versus differently, it is unclear how to fairly compare the predictions for the different metrics.

These limitations point to a need for theory that can extend and unify the results for each particular metric as well as explain why predictions differ between metrics. The following presents the model and approach I use as a step toward addressing this need.

## 3 Models and Methods

### 3.1 Single-habitat *n*-host environmental transmission model

I focus on the model in Cortez (2021) of *n* competing host species and a shared environmentally transmitted pathogen. Infection occurs when susceptible individuals encounter infectious propagules (e.g., spores) in the environment that were shed by infected individuals. All host species shed infectious propagules into a shared habitat and are equally exposed to the infectious propagules. Examples of environmentally transmitted pathogens include the human pathogens *Vibrio cholera* and *Giardia lamblia* (Kaper et al., 1995; Wolfe, 1992), the fungal pathogen *Metschnikowia bicuspidata* of *Daphnia* species (Strauss et al., 2015; Searle et al., 2016), the chytrid fungus *Batrachochytrium dendrobatidis* of amphibians (Daszak et al., 2003; Skerratt et al., 2007), whirling disease in fish (Hedrick et al., 1998; Bartholomew and Reno, 2002), and trematode parasites of snails (Johnson et al., 2012). Without loss of generality, I refer to host 1 is the focal host and the others as alternative hosts; this choice is arbitrary, but in practice, the designation depends on the specific system being studied.

Each host species is divided into subpopulations of susceptible (*S*_*i*_), infected (*I*_*i*_), and recovered (*R*_*i*_) individuals, where the total density for host *i* is *N*_*i*_ = *S*_*i*_ + *I*_*i*_ + *R*_*i*_. The model describes the changes in each host class and infectious propagule density (*P*),

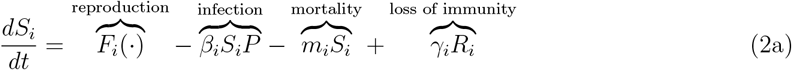

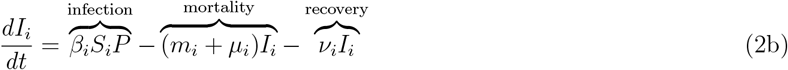

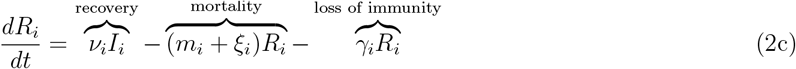

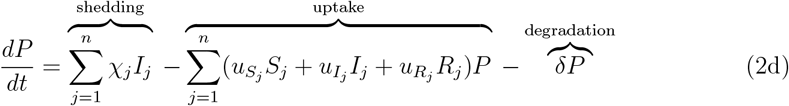

The parameter values are summarized in Table 1 and additional details are provided in Appendix S1. In equation (2a), host *i* susceptible density increases due to reproduction (*F*_*i*_), decreases due to infection (*β*_*i*_*S*_*i*_*P*) and non-disease mortality (*m*_*i*_*S*_*i*_), and increases due to waning immunity (*γ*_*i*_*R*_*i*_). In equation (2b), host *i* infected density increases due to new infections (*β*_*i*_*S*_*i*_*P*) and decreases due to non-disease mortality (*m*_*i*_*I*_*i*_), mortality due to disease (*μ*_*i*_*I*_*i*_), and recovery (*ν*_*i*_*Y*_*i*_). Because infection occurs when infectious propagules are taken up by susceptible individuals, the transmission parameter is the product of the uptake rate of susceptible individuals and the per infectious propagule probability of infection 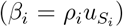. In equation (2c), host *i* recovered density increases due to recovery (*ν*_*i*_*I*_*i*_) and decreases due to mortality (*m*_*i*_*R*_*i*_ +*ξ*_*i*_*R*_*i*_) and waning immunity (*γ*_*i*_*R*_*i*_). I assume susceptible individuals and recovered individuals are identical except that recovered individuals may experience increased mortality due to prior exposure to the pathogen (*ξ*_*i*_ ≥ 0). In equation (2d), infectious propagule density increases due to shedding by infected individuals (first sum) and decreases due to uptake of infectious propagules by all individuals of all host species (second sum) and degradation (*δP*).

**Table 1:**
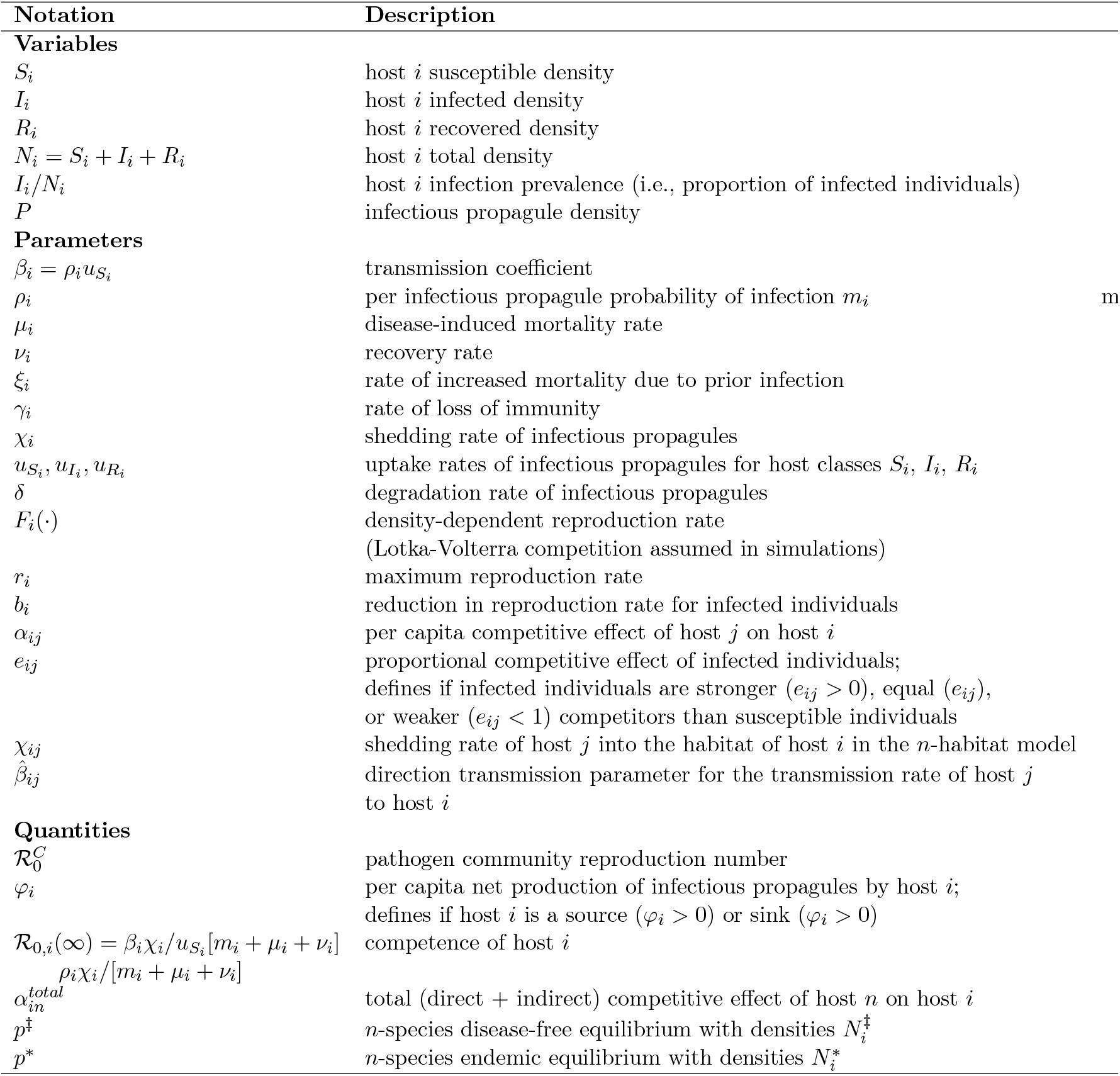
Model variables, parameters, and important quantities

The reproduction rates, *F*_*i*_, account for intraspecific and interspecific host competition. For the numerical simulations presented in figures, I use the Lokta-Volterra competition functions,

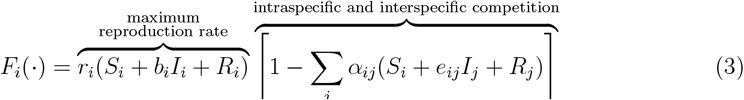

where for host *i, r*_*i*_ is the maximum reproduction rate, *b*_*i*_ is the reduction in growth rate for infected individuals, *α*_*ij*_ is the per capita competitive effect of host *j*, and *e*_*ij*_ determines if infected individuals of host *j* are weaker (*e*_*ij*_ < 1), equal (*e*_*ij*_ = 1), or stronger (*e*_*ij*_ > 1) competitors than susceptible individuals. The latter accounts for the possibility that infected individuals may be stronger competitors when infection causes increased appetite or resource acquisition rates (Ponton et al., 2011; Shikano and Cory, 2016; Bernardo and Singer, 2017).

I note a few important assumptions about model (2); additional mathematical details about these and other assumptions are provided in Section S1.2.4. First, I assume intraspecific density dependence and interspecific host competition only affect the reproduction rates of the host species (*F*_*i*_) and do not affect the host mortality rates. These assumptions simplify the mathematical analysis and align with many multi-host-pathogen models (but see, e.g., Marini et al. (2017)). Second, while I describe and interpret the interspecific host interactions in terms of competitive interactions (i.e., −, − interactions), my results also hold for species that have mutualistic (+, −) or contramensalistic (+, −) interactions, provided the interactions only affect the reproduction rates of the host species. My results may not apply to species with predator-prey or consumer-resource interactions because those interspecific interactions necessarily affect the mortality rates of a species.

Third, I assume the *n* host species stably coexist in the presence and absence of the pathogen, respectively, at a disease-free equilibrium (with densities 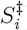) and an endemic equilibrium (with densities 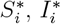, and 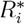). Without this assumption, my equilibrium-based analyses are biologically uninformative. In addition, to explore the effects of host additions on disease and without loss of generality, I assume host *n* is the added host species and assume that host species 1,…,*n* − 1 can stably coexist in presence and absence of the pathogen. This allows me to compare disease levels when the added host species is absent and present. However, this assumption means that my results may not apply to systems where host coexistence is pathogen-mediated or systems where the gain or loss of the added host causes other host species to go extinct.

### 3.2 Model Generality

The predictions for the environmental transmission (ET) model (2) apply to direct transmission models (Cortez, 2021). This is because the ET model (2) can reduce to a model with density-dependent direct transmission (DDDT), frequency-dependent direct transmission (FDDT), or direct transmission that is intermediate to DDDT and FDDT; see Box 1 for specific conditions. This means predictions from model (2) simultaneously apply to ET, DDDT, and FDDT.

While model (2) can reduce to a direct transmission form, it is not flexible enough to reduce down to all possible parameterizations of DDDT and FDDT models. The underlying reason is that model (2) reduces to a direct transmission form where the interspecific (*β*_*ij*_) and intraspecific (*β*_*ii*_) transmissions coefficients are correlated and not independent (mathematical details provided in Appendix S1.2). To address this issue, in Appendix S2, I extend my results to an *n*-habitat version of the model that allows for partially overlapping host habitats.

The key difference between the 1-habitat and *n*-habitat models is that each host species can have different shedding rates in their own habitat (*χ*_*jj*_) and the habitats of heterospecifics (*χ*_*ij*_). This allows for each species to have different rates of transmission to conspecifics and heterospecifics. That makes the *n*-habitat model flexible enough to reduce down to all possible parameterizations of DDDT and FDDT models (mathematical details provided in Appendix S2.2).

A full analysis of the *n*-habitat ET model is not tractable. Thus, Appendix S2 primarily focuses on systems where (i) interspecific host competition is absent and (ii) each host species has low shedding and uptake rates in the habitats of other host species. When the *n*-habitat ET model is reduced down to a direct transmission form, these conditions result in a direct transmission model where (i) interspecific host competition is absent and (ii) interspecific direct transmission rates are lower than intraspecific direct transmission rates (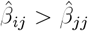, where 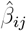 is the direct transmission rate parameter for host *j* transmitting the pathogento host *i*). While restrictive, these conditions align with the assumptions in prior studies on DDDT and FDDT models (Dobson, 2004; Joseph et al., 2013; Mihaljevic et al., 2014). Thus, the restricted results for the *n*-habitat model help explain patterns from these prior studies.

In total, analyzing the 1-habitat model (2) and the *n*-habitat model provides a way to extend and unify results for ET, DDDT, and FDDT. The main text focuses on the simpler 1-habitat model because the results from the 1-habitat and *n*-habitat models are qualitatively similar. The only difference is that the predictions for the *n*-habitat model depend the host’s rates of shedding into the habitats of heterospecifics. These differences are indicated in the text.

### 3.3 Defining host competence, source, and sink

My mathematical analysis identifies two characteristics of host species that measure how well each host species can transmit the pathogen.

First, host competence measures the potential ability of an exposed individual to transmit the pathogen over the entire length of time the individual is infected. Mathematically, I define host competence as

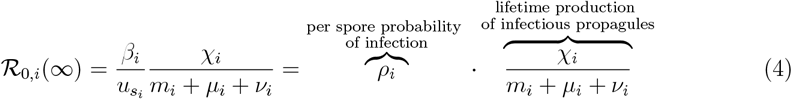

where the second equality follows from the transmission rate (*β*_*i*_) being the product of the per infectious propagule probability of infection (*ρ*_*i*_) and the uptake rate of infectious propagules 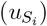. On the right hand side of the equation, the first term is the probability of infection given exposure and the fraction is the shedding rate (*χ*_*i*_) multiplied by the average duration of infection (1*/*[*μ*_*i*_+*m*_*i*_+*γ*_*i*_]), which equals the average lifetime production of infectious propagules by an infected individual of host *i*. Combined, this yields the average lifetime production of infectious propagules by individuals that are exposed to the pathogen. This definition of competence matches that proposed in prior studies (Merrill and Johnson, 2020). I note that my notation for competence comes from equation (4) being mathematically equivalent to the pathogen’s basic reproduction number for an arbitrarily large population of host *i*; see Appendix S1.3 for additional details.

Second, source and sink measure a host species’ instantaneous per capita net production rate of infectious propagules. Mathematically, source and sink are defined by,

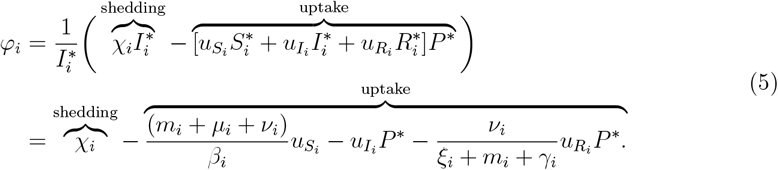

(The second line follows from solving the equilibrium conditions *dI*_*i*_*/dt* = 0 and *dR*_*i*_*/dt* = 0 for 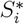 and 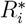.) Source hosts are net producers of infectious propagules, meaning the host’s per capita shedding rate is larger than its per capita uptake rate (*φ*_*i*_ > 0). Sink hosts are net removers of infectious propagules, meaning the host’s per capita shedding rate is smaller than its per capita uptake rate (*φ*_*i*_ < 0).

While similar, competence and sink/source are distinct because competence measures the lifetime production of infectious propagules per individual whereas sink and source measure the net instantaneous production of infectious propagules per individual. The similarities between competence and sink/source mean that high and low competence often means source and sink, respectively. However, the differences between the definitions mean that high competence hosts can be sinks and low competence hosts can be sources. First, high competence hosts are sinks when they have low mortality and recovery rates (small *μ*_*i*_ and *ν*_*i*_, making ℛ_0,*i*_(∞) large) and small shedding rates (small *χ*_*i*_, making *φ*_*i*_ negative or small and positive). Such hosts have low instantaneous production rates of infectious propagules, but because individuals remain infected for long periods of time, their lifetime production rates of infectious propagules are large. Second, low competence hosts are large sources when they have high mortality and recovery rates (large *μ*_*i*_ + *ν*_*i*_, making ℛ_0,*i*_(∞) small) and large shedding rates (large *χ*_*i*_, making *φ*_*i*_ large and positive). Such hosts have high instantaneous production rates of infectious propagules, but because individuals remain infected for short periods of time, their lifetime production rates of infectious propagules are small.

I note two additional things about the definitions of competence, sink, and source.

First, in addition to arising naturally in the calculations, an advantage of these definitions is that they provide a density-independent ranking of the host species from highest to lowest competence and from largest source to largest sink. This avoids complications that arise when densities shift as host species are added or removed from a community. However, the rankings can differ across environments because both definitions depend on the non-disease mortality rates, *m*_*i*_. Second, both definitions apply to direct transmission pathogens because of the connection between direct and environmental transmission models (Box 1). For competence, higher competence hosts have larger intraspecific transmission rates and lower mortality and recovery rates. For source and sink, all hosts are sources (*φ*_*i*_ > 0) under DDDT because DDDT corresponds to no uptake in the ET model (2). Under FDDT, hosts are sources (*φ*_*i*_ > 0) when their intraspecific and interspecific transmission rates are sufficiently large and sinks (*φ*_*i*_ < 0) otherwise; see Appendices S1.3 and S2.2 for mathematical details.

### 3.4 Computing the sensitivities of the disease metrics

I use model (2) to predict how additions and removals of alternative hosts affect three disease metrics: (i) the parasite’s basic reproduction number for the community 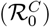), (ii) focal host infection prevalence at equilibrium 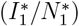, and (iii) focal host infected density at equilibrium 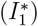. Because explicit formulas do not exist for the metrics except in special cases, I cannot directly compute how additions and removals of alternative hosts affect each metric. However, it is possible to compute explicit formulas for the slopes of the relationships between each metric of disease and alternative host competence (ℛ_0,*n*_(∞)), alternative host equilibrium density 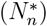, and the pathogen transmission mode. The signs of the slopes yield insight into the effects of additions/removals on each metric of disease. As in prior studies (Roberts and Heesterbeek, 2018; Cortez and Duffy, 2021; Cortez, 2021), I compute the slopes using local sensitivities (i.e., derivatives). Here, I describe the idea behind the approach. Box 2 and Appendix S2.4 explain how the 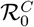 sensitivities are computed using the theory for stage-structured models. Appendix S1.5 explains how the sensitivities of the other two metrics are computed using the Jacobian-based approach in Cortez and Duffy (2021) and Cortez (2021).

The slopes of the relationships between each metric of disease and the components of alternative host competence are computed using derivatives with respect to the alternative host’s transmission (*β*_*n*_), mortality (*μ*_*n*_), recovery (*ν*_*n*_), shedding (*χ*_*n*_), and uptake 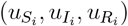 rates. For example, the sensitivities for the alternative host’s shedding rate are computed using the derivatives,

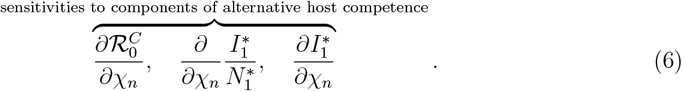

Positive and negative values imply that each metric increases and decreases, respectively, as the alternative host’s shedding rate increases. These sensitivities are useful because they identify how variation in the epidemiological traits of the alternative host species influences the disease metrics. In addition, the sensitivities can provide insight about the consequences of intervention strategies that target a specific biological process, e.g., vaccination that lowers infection rates of an alternative host (decreased *β*_*n*_) or the culling of infected individuals of an alternative host (increased *μ*_*n*_).

The slopes of the relationships between each metric of disease and alternative host density are computed using the derivatives

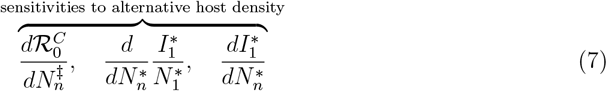

where 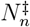 and 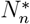 are the densities of host *n* at the disease-free and endemic equilibria, respectively. (The change in derivative notation is mathematically necessary to account for the total effect of variation in alternative host density.) Positive and negative values imply that each metric increases and decreases, respectively, as the alternative host’s density increases. This yields insight about the effects of host additions because the effect of adding an alternative host to the community is predicted by the slope of the response as alternative host density is increased from zero to its equilibrium value. In particular, positive and negative values suggest that the addition of the alternative host will increase and decrease, respectively, each metric of disease.

In total, the sensitivity formulas identify the specific conditions under which each metric has positive versus negative relationships with alternative host competence, alternative host density, and the pathogen transmission mechanism. Comparing those conditions yields insight into when the three metrics will have similar or different responses to host additions and removals.

## 4 Results

I first present the general mechanisms causing the sensitivities for the three metrics to differ in sign. I then compare how the metrics respond to changes in alternative host competence, alternative host density, and the pathogen transmission mode.

### 4.1 General mechanisms for why sensitivities can have different signs

The three general mechanisms explaining why the sensitivities for the metrics can differ in sign are (1) differences in the definitions of competence and source/sink, (2) density-mediated feedbacks driven by disease-induced mortality, and (3) large changes in focal host density.

For mechanism 1, differences in the definitions of competence and source/sink can cause the sensitivities of 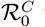 to differ in sign from the sensitivities of focal host infection prevalence 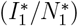 and infected density 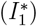 at equilibrium. The reason is that the sensitivity formulas for 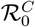 depend on host competence (ℛ_0,*i*_(∞)) whereas the sensitivity formulas for focal host infection prevalence and infected density depend on whether hosts are sources or sinks (*φ*_*i*_). Consequently, predictions for 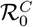 can disagree with predictions for the other two metrics when high and low competence do not imply source and sink, respectively.

For mechanism 2, density-mediated feedbacks driven by disease-induced mortality can cause the sensitivities of 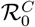 to differ in sign from the sensitivities of the other two metrics. The underlying reason is that 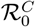 is computed using the disease-free equilibrium densities 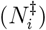, which are independent of disease-induced mortality rates, whereas focal host infection prevalence and infected density depend on the disease-induced mortality rates. For the latter, as conditions are varied (e.g., a host species is added to the community), changes in infectious propagule densities will lead to changes in infection rates, that will lead to changes in total disease-induced mortality rates for each host (*μ*_*i*_*I*_*i*_), and that will lead to changes in the total and infected densities of each host. If the feedbacks due to disease-induced mortality are strong, then the changes in total and infected densities will be large. That in turn can cause focal host infection prevalence and infected density to respond in the opposite direction of 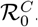.

For mechanism 3, large changes in focal host density can cause the sensitivities of focal host infection prevalence and focal host infected density to differ in sign. The mathematical reason is that the quotient rule from calculus implies that the sensitivities to alternative host density (equation (7)) are related by

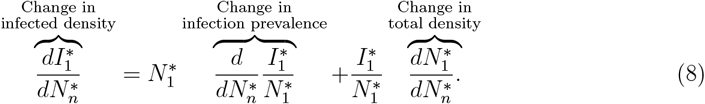

The same relationship applies to the sensitivities to parameter values (equation (6)). The sensitivities of infected density (left side of equation) and infection prevalence (first term on the right side of the equation) can only differ in sign if the changes in total focal host density (second term on the right side of the equation) are large and in the opposite direction of the changes in infection prevalence. For example, increased density of host *n* could increase focal infection prevalence 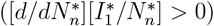 but also decrease focal host infected density 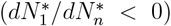because of large decreases in focal host density 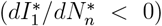. My calculations show that the sensitivities of focal host infection prevalence and infected density have opposite signs only if (i) some host species are more strongly regulated by disease and competition with alternative host species than by competition with the focal host species or (ii) the focal host is a sink or small source and a subset of the alternative hosts are large sources; see Remark S1.6 of Appendix S1.1.5 and Appendix S1.5 for mathematical details. The most likely scenario for condition (i) to be met both biologically and mathematically is when the focal host is strongly regulated by disease or interspecific competition and weakly regulated by intraspecific competition. Because of this, I will only discuss condition (i) in terms of regulation of the focal host.

### 4.2 Sensitivities of disease metrics to alternative host competence

I now compare the sensitivities of the three metrics to the components that define the competence of the alternative host, 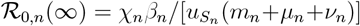. These sensitivities are useful because they help identify how variation in the epidemiological traits of the alternative host species influences disease levels and disease risk in the focal host. For instance, the sensitivities can provide insight about the consequences of intervention strategies that target a specific biological process, such as vaccination that lowers infection rates of an alternative host (decreased *β*_*n*_).

#### Community reproduction number

Any variation in epidemiological traits that increases the competence of the alternative host always increases 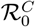; mathematical details provided in Appendix S1.4.2. In particular, increased shedding rates (*χ*_*n*_), decreased uptake rates 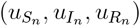, increased transmission rates (*β*_*n*_; Figure 1A), decreased mortality rates (*μ*_*n*_; Figure 1D), and decreased recovery rates (*ν*_*n*_) of an alternative host always increase 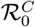. These predictions explain why 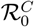 always increased with increased host competence in prior studies of direct (Dobson, 2004; Joseph et al., 2013; O’Regan et al., 2015; Huang et al., 2019) and environmental (Feng et al., 2009; Espira et al., 2022) transmission pathogens; see Appendix S3 for details about each study.

**Figure 1:**
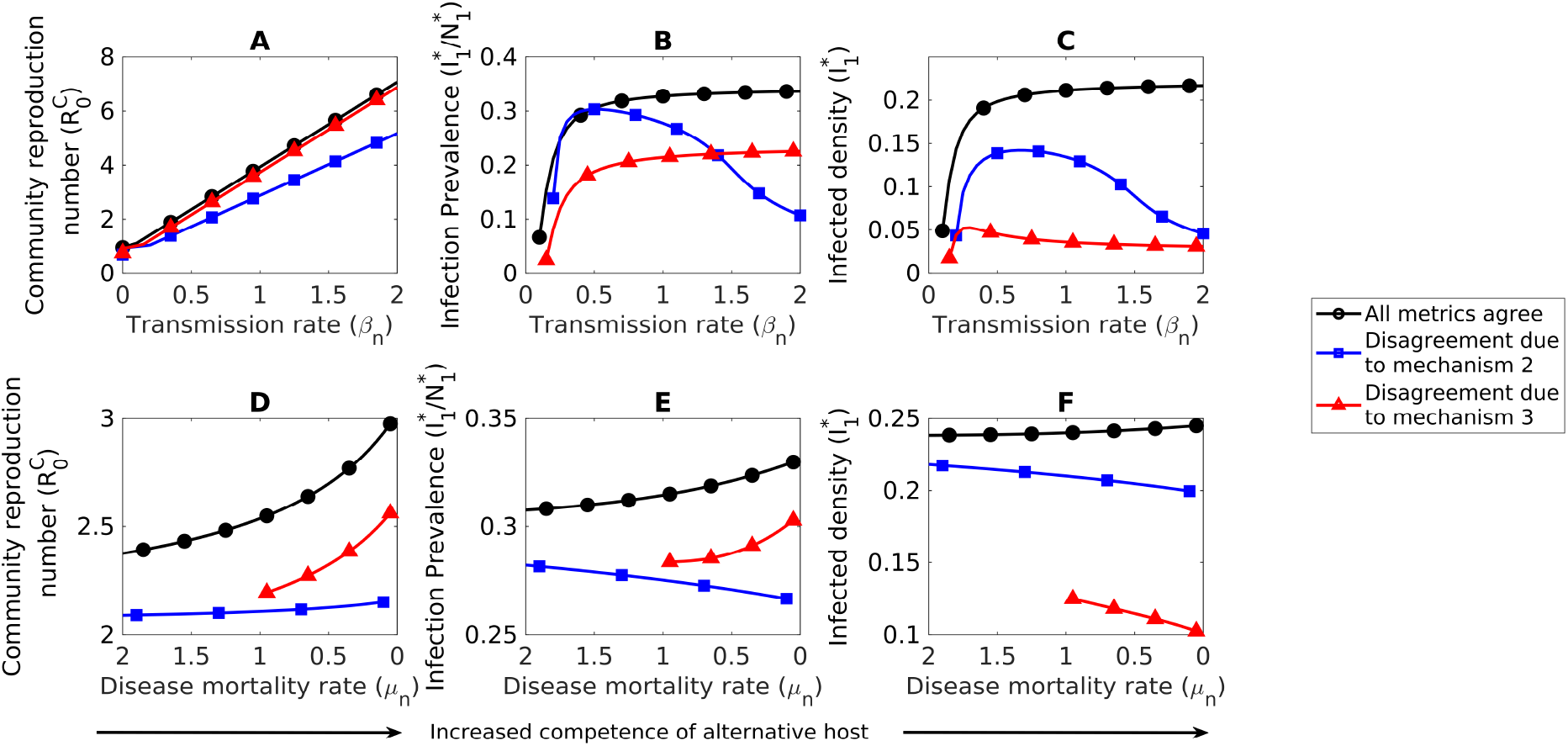
Increased competence of an alternative host increases all metrics of disease (black circles), unless there are strong density-mediated feedbacks due to disease-induced mortality (mechanism 2; blue squares) or large changes in focal host density (mechanism 3; red triangles). Note that the x-axes in the bottom row are reversed. (A,D) 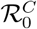 always increases with increased transmission rates and decreased disease-induced mortality rates of an alternative host. (B,E) Focal host infection prevalence 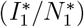 increases with increased transmission rates and decreased disease-induced mortality rates of an alternative host (black circles and red triangles in B,E), unless there are strong density-mediated feedbacks due to disease-induced mortality (mechanism 2), which occurs when interspecific competition is sufficiently large (blue squares in B) or the alternative host is a large sink (blue squares in E). (C,F) Focal host infected density 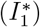responds in the same way as focal host infection prevalence (black circles and blue squares), unless there are large changes in focal host density (mechanism 3), which occurs when the focal host is more strongly regulated by the disease or interspecific competition than intraspecific competition (red triangles in C) or the focal host is a sink and other hosts are large sources (red triangles in F). In each row, curves of the same color are from the same parameterization of a four-host version of model (2); see Appendix S4 for parameter values.

#### Focal host infection prevalence

Variation in epidemiological traits that increases the competence of the alternative host often increases focal host infection prevalence, but not always; mathematical details provided in Appendix S1.5.3. Specifically, increases in the alternative host’s shedding rate (*χ*_*n*_) or decreases in its uptake rates 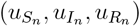 always increase focal host infection prevalence. In comparison, increases in an alternative host’s transmission parameter (*β*_*n*_) and decreases in its mortality and recovery rates (*μ*_*n*_, *ν*_*n*_) often increase focal host infection prevalence (black circles and red triangles in Figure 1BE). However, decreased focal host infection prevalence is also possible; here, I focus on the three most biologically likely scenarios and refer the reader to Cortez (2021) and Appendix S1.5.3 for additional details. First, increases in the alternative host’s transmission parameter (*β*_*n*_) can decrease focal host infection prevalence when the alternative host is a large source (*φ*_*n*_ > 0) and interspecific competition is sufficiently strong (blue squares in Figure 1B). Second, decreases in the alternative host’s mortality rate (*μ*_*n*_) can decrease focal host infection prevalence when the alternative host is a large sink (*φ*_*n*_ < 0; blue squares in Figure 1E). Third, increases in *β*_*n*_ and decreases in *μ*_*n*_ or *ν*_*n*_ can decrease focal host infection prevalence when the alternative host has stronger interspecific competitive effects on sink hosts than source hosts. For all three scenarios, increased competence of the alternative host causes shifts in its densities that reduce the production of infectious propagules, which in turn causes decreases in focal host infection prevalence.

#### Focal host infected density

The predictions for focal host infected density are often the same as focal host infection prevalence. However, the predictions can be reversed if (i) the focal host is more strongly regulated by the disease or interspecific competition than by intraspecific competition or (ii) the focal host is a sink or small source and other hosts are large sources. Mathematical details are given in Appendix S1.5.3.

#### Comparison of metrics

Intuition suggests that higher competence in the alternative host (larger ℛ_0,*n*_(∞)) will lead to increases in all metrics of disease. This is often the case, resulting in all metrics responding to variation in alternative host competence in the same direction (black circles in Figure 1). However, the intuition can be incorrect for focal host infection prevalence and infected density. This results in variation in the competence of the alternative host having effects of one sign on 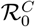 and effects of the opposite sign on the other focal host infection prevalence and infected density (blue squares in 1). This difference in responses is due to density-mediated feedbacks (mechanism 2) where changes in alternative host competence alter infectious propagule densities, which affects disease-induced mortality in the focal host.

Variation in the competence of the alternative host also can have effects of opposite sign on focal host infection prevalence and focal host infected density. This difference in responses is due to variation in the competence of the alternative host causing large changes in focal host density (mechanisms 3). This occurs when (i) the focal host is more strongly regulated by the disease or interspecific competition than intraspecific competition (red triangles in Figure 1A-C) or (ii) the focal host is a sink or small source and other hosts are large sources (red triangles in Figure 1D-F).

### 4.3 Sensitivities of metrics to alternative host density

Here, I compare how each disease metric responds to variation in the density of an alternative host. These sensitivities yield insight about the effects of host additions because positive and negative values suggest that the addition of an alternative host will increase and decrease a metric, respectively. To facilitate that connection and avoid confusion, I vary the density of host *n* and refer to host *n* as the “added host”.

The response of each metric depends on how the densities of all host species shift as the density of the added host increases. In the following formulas, 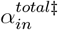 and 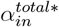 denote the total competitive effects (i.e., direct + indirect effect) of the added host on host *i* at the disease-free and endemic equilibria, respectively (see Appendix S1.1.5 for details). The total competitive effects can be positive or negative depending on the strength and asymmetry of interspecific host competition across the community. If the added host has a positive (negative) total effect on another host, then increased density of the added host causes an increase (decrease) in the density of the other host.

#### Community reproduction number

The sensitivity of 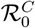 to the density of the added host is

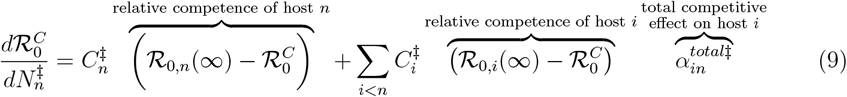

where the 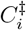 are positive constants; see Appendix S1.4.1. Equation (9) shows that the change in 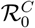 depends on whether the competence of each host species relative to 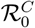 is high 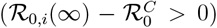 or low 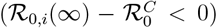 and whether the added host has a positive 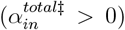 or negative 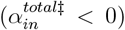 total competitive effect on each species. In particular, increased density of the added host is more likely to increase 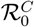 when the added host has higher competence 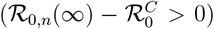, the added host has less negative total effects on high competence hosts 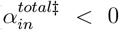small in magnitude for hosts with 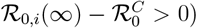, and the added host has more negative total effects on low competence hosts (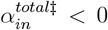 large in magnitude for hosts with 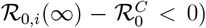). In comparison, increased density of the added host is more likely to decrease 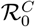 when the added host has lower competence 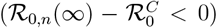, the added host has less negative total effects on low competence hosts (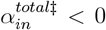 small in magnitude for hosts with 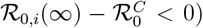), and the added host has more negative total effects on high competence hosts (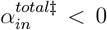 large in magnitude for hosts with 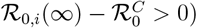).

Combining the above yields the following predictions about how 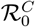 responds to increased density of the added host. First, if interspecific competition is absent, then the response is determined solely by the competence of the added host: 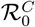 increases when the added host has sufficiently high competence (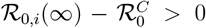; black circles in Figure 2A) and decreases otherwise (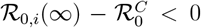; black circles in Figure 2D). Second, these predictions can be reversed if interspecific competition is sufficiently strong, depending on the competitive effects of the added host. In particular, increased density of an added host with high competence can decrease 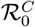 when the added host has more negative (or less positive) total competitive effects on high competence hosts than low competence hosts (blue squares in Figure 2A). Similarly, increased density of an added host with low competence can increase 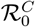 when the added host has more negative (or less positive) total competitive effects on low competence hosts than high competence hosts (blue squares in Figure 2D).

**Figure 2:**
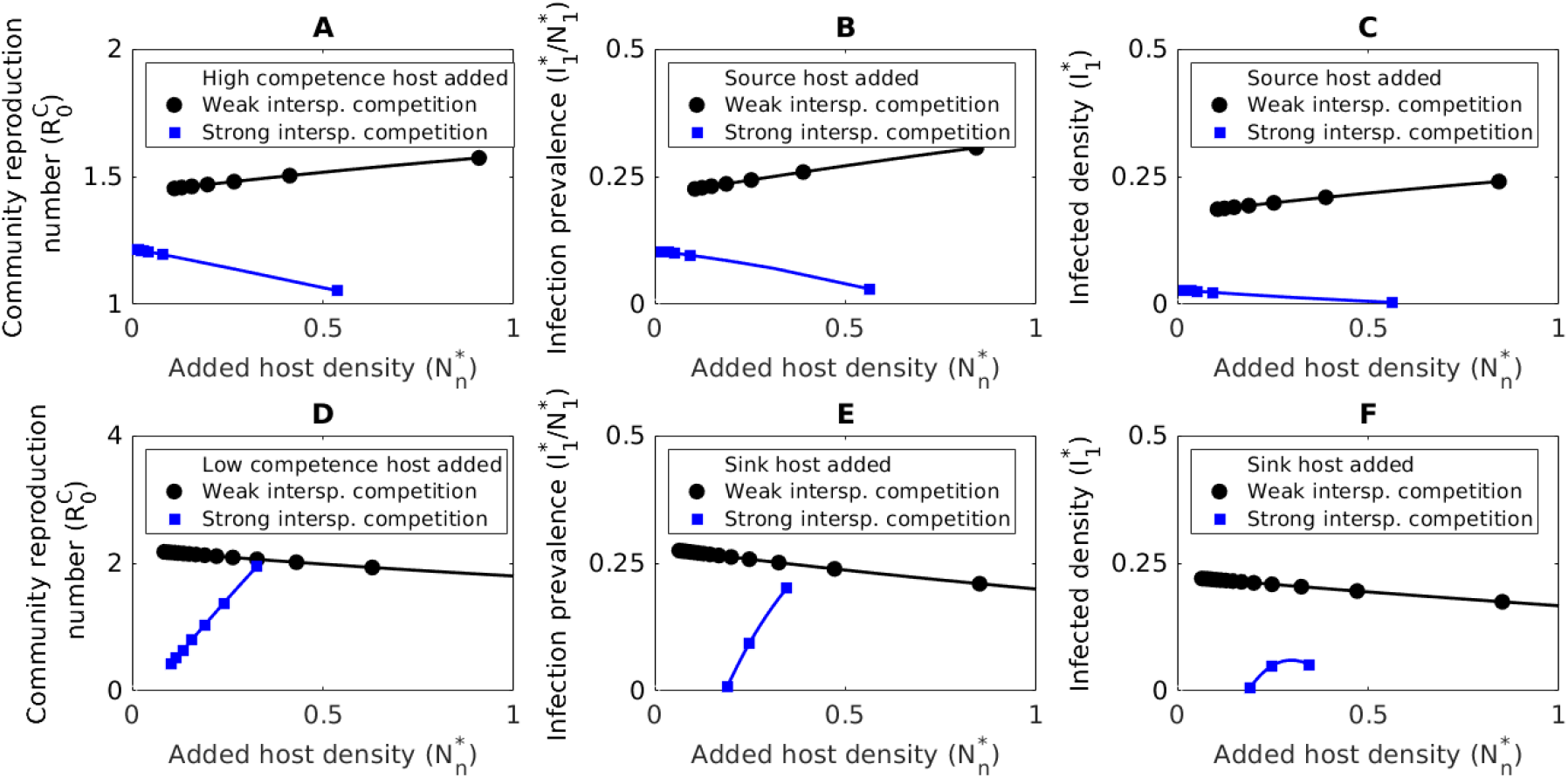
Strong interspecific competition can reverse predictions about the effects of increased aded host density on each disease metric. (A-C) Increased density of an added high competence source host increases all disease metrics in the absence of interspecific competition (black circles), but decreases all disease metrics when the added host has more negative (or less positive) total competitive effects on low competence source hosts than high competence sink hosts (blue squares). (D-F) Increased density of an added low competence sink host decreases all disease metrics in the absence of interspecific competition (black circles), but increases all disease metrics when the added host has more negative (or less positive) total competitive effects on low competence sink hosts than high competence source hosts (blue squares). In each row, curves of the same color are from the same parameterization of a four-host version of model (2); see Appendix S4 for parameter values.

The above predictions are only slightly modified when accounting for partially overlapping habitats. In particular, the conditions depend both on host competence and inter-habitat shedding rates (i.e., shedding rates into other species’ habitats); mathematical details for the analysis of the *n*-habitat model are provided in Appendix S2.4.2. Specifically, in the absence of interspecific competition, 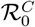 increases when the added host has sufficiently high competence and inter-habitat shedding rates and 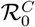 decreases when the added host has sufficiently low competence and inter-habitat shedding rates. When interspecific competition is present, these predictions can be reversed if the added host has more negative total competitive effects on hosts with high competence and high inter-habitat shedding rates or hosts with low competence and low inter-habitat shedding rates, respectively.

#### Focal host infection prevalence

The effect of increased density of the added host on focal host infection prevalence is

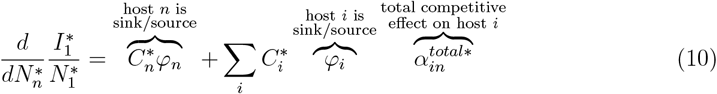

where the 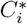 are positive constants; see Appendix S1.5.2. Due to their similar forms, the interpretation of equation (10) is the same as equation (9) except that high and low competence are replaced with source and sink, respectively. In particular, increased density of the added host is more likely to increase focal host infection prevalence when the added host is a source (*φ*_*n*_ > 0), the added host has less negative total effects on source hosts (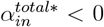 small in magnitude for hosts with *φ*_*i*_ > 0), and the added host has more negative total effects on sink hosts (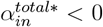 large in magnitude for hosts with *φ*_*i*_ < 0). In comparison, increased density of the added host is more likely to decrease focal host infection prevalence when the added host is a sink (*φ*_*n*_ < 0), the added host has less negative total effects on sink hosts (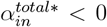 small in magnitude for hosts with *φ*_*i*_ < 0), and the added host has more negative total effects on source hosts (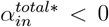 large in magnitude for hosts with *φ*_*i*_ > 0).

Combining the above yields the following predictions about how focal host infection prevalence responds to increased density of the added host. First, if interspecific competition is absent 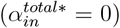, then the response is determined solely by whether the added host is a source or sink: focal host infection prevalence increases when the added host is a source (*φ*_*n*_ > 0; black circles in Figure 2BC) and decreases when the added host a sink (*φ*_*n*_ < 0; black circles in Figure 2EF). Second, these predictions can be reversed if interspecific competition is sufficient strong, depending on how competition affect other source and sink hosts. In particular, increased density of an added source host can decrease focal host infection prevalence when the added host has more negative (or less positive) total competitive effects on source hosts than sink hosts (blue squares in Figure 2BC). Similarly, increased density of an added sink host can decrease focal host infection prevalence when the added host has more negative (or less positive) total competitive effects on sink hosts than source hosts (blue squares in Figure 2EF).

I note that when accounting for partially overlapping habitats, the above predictions are extended to account for whether each host species is a source or sink in each species’ habitat; see Appendix S2.5.2 for mathematical details.

#### Focal host infected density

The predictions for focal host infected density are often the same as focal host infection prevalence (black circles and blue squares in Figure 3ABEF). However, the predictions can be reversed if (i) the focal host is more strongly regulated by the disease or interspecific competition than by intraspecific competition or (ii) the focal host is a sink or small source and other hosts are large sources. Mathematical details are given in Appendix S1.5.2.

**Figure 3:**
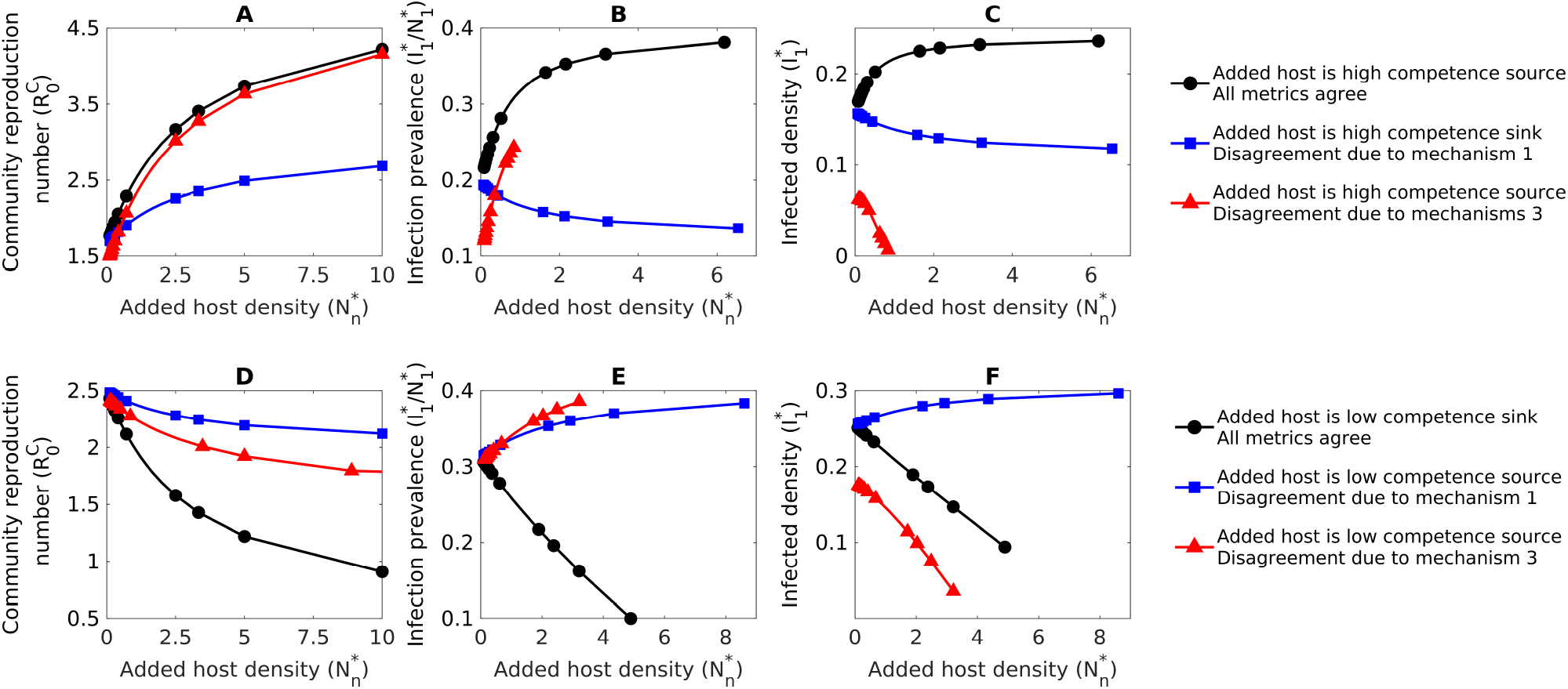
Increased density of an added host causes all metrics to change in the same direction (black circles), unless source and sink do not imply high and low competence (mechanism 1; blue squares) or there are large changes in focal host density (mechanism 3; red triangles). Increased density of the added host (A) always increases 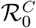 when the added host has sufficiently high competence and (D) always decreases 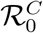 when added host has sufficiently low competence. (B,C) Increased density of the added host increases focal host infection prevalence 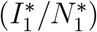 when the added host is a source (black circles and red triangles in B; blue squares and red triangles in E) and decreases focal host infection prevalence when the added host is a sink (blue squares in B; black circles in E). (C,F) Increased density of the added host often increases focal host infected density 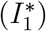 when the added host is a source (black circles in C; blue squares in F) and decreases focal host infected density when the added host is a sink (blue squares in C; black circles in F). However, focal host infected density can decrease when the added host is a source if there are large changes in focal host density (mechanism 3), which occurs when the focal host is more strongly regulated by disease and interspecific competition than by intraspecific competition (red triangles in C,F). In each row, curves of the same color are from the same parameterization of a four-host version of model (2); see Appendix S4 for parameter values.

#### Comparison of metrics

Increased alternative host density often has effects of the same sign on all metrics (black circles in Figure 3). However, responses in 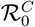 can differ in sign from the other two metrics due to differences in the definitions of competence and source/sink (mechanism 1). For example, increased density of a high competence sink host increases 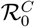 but decreases focal host infection prevalence and infected density (blue squares in Figure 3A-C). Similarly, increased density of a low competence source host decreases 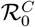 but increases focal host infection prevalence and infected density (blue squares in Figure 3D-F). In addition, while focal host infection prevalence and infected density often respond in the same direction (black circles and blue squares in Figure 3ABEF), the two metrics can response in opposite directions due to large changes in focal host density (mechanism 3) when (i) the focal host is more strongly regulated by disease or interspecific competition than intraspecific competition (red triangles in Figure 3BCEF) or (ii) the focal host is a sink or small source and other hosts are large sources (not shown).

### 4.4 Predictions for host species richness-disease relationships

The sensitivities in the previous section predict how increased density of an added host affect the three disease metrics. Here, I used those results to predict how interspecific competition and correlations between host competence and arrival order shape the relationships between host species richness and each disease metric.

#### Community reproduction number

The predicted effects of host additions on 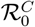 yield the following kinds of relationships between host species richness and 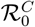. First, 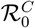 is predicted to increase when added hosts have sufficiently high competence (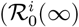 large) and inter-habitat shedding rates (large 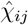), unless the added host has sufficiently strong negative competitive effects on other hosts with high competence and inter-habitat shedding rates. As an example, consider a system where higher competence hosts are only present in communities with more species, i.e., there is a positive correlation between host competence and host arrival order. In such a system, there is typically a positive relationship between host species richness and 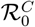 (black circle in Figure 4A). However, a negative relationship can arise if higher competence hosts are also stronger interspecific competitors because the later arriving hosts suppress host densities (blue squares in Figure 4A).

**Figure 4:**
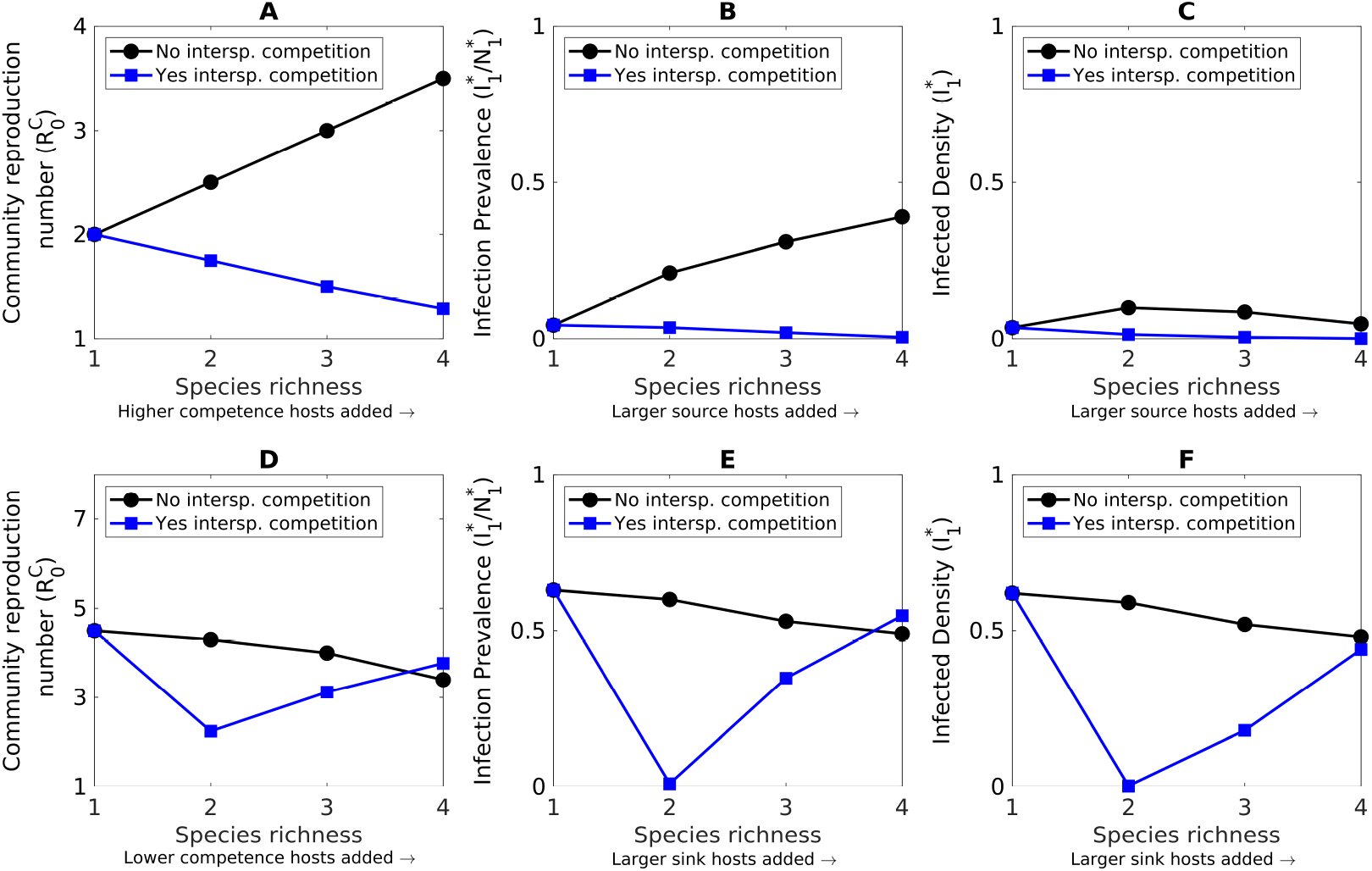
The shapes of host species richness-disease relationships depend on the characteristics of added host species and levels of interspecific competition. (A) In a system where higher competence hosts are added later, 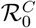 increases with host species richness (black circles), unless interspecific competition is sufficiently strong (blue squares). (B) In a system where larger source hosts are added later, focal host infection prevalence increases with host species richness (black circles), unless interspecific competition is sufficiently strong (blue squares). (D) In a system where lower competence hosts are added later, 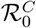 decreases with host species richness (black circles), unless added low competence hosts have strong negative competitive effects on other sink low competence hosts (blue squares). (E) In a system where larger sink hosts are added later, focal host infection prevalence decreases with host species richness (black circles), unless added sink hosts have strong negative competitive effects on other low competence hosts (blue squares). (C,F) Focal host infected density often responds to increases in host species richness in the same way that focal host infection prevalence responds (blue squares in C, all curves in F). However, differences can arise if there are large changes in focal host density (mechanism 3; black circles in C). Values are from a four-host version of model (2); see Appendix S4 for parameter values.

Second, 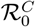 is predicted to decrease when added hosts have sufficiently low competence (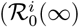 small) and inter-habitat shedding rates (small 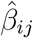), unless the added host has sufficiently strong negative competitive effects on other hosts with low competence and inter-habitat shedding rates. As an example, consider a system where lower competence hosts are only present in communities with more species, i.e., there is a negative correlation between host competence and host arrival order. In such a system there is typically a negative relationship between host species richness and 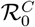 (filled black circles in Figure 4D). However, U-shaped relationships between host species richness and 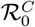 are possible if low competence hosts also have strong interspecific competitive effects on other low competence hosts (blue squares in Figure 4D). In this case, the densities of the low competence hosts are suppressed as more species are added to the community, which leads to increases in 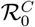.

I note that if lower competence hosts are only present in communities with more species and the competences of the lower competence hosts are not too low, then 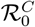 can have a positive relationship with host species richness In these counter-intuitive cases, later arriving hosts species are worse at spreading the disease than earlier arriving hosts, but all host species facilitate disease spread across the community. Mathematically, this case corresponds to equation (9) where the later arriving hosts have smaller ℛ_0,*i*_(∞) values, but all values are large enough that 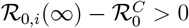.

#### Focal host infection prevalence

The predicted effects of host additions on focal host infection prevalence yield the following kinds of relationships between host species richness and focal host infection prevalence. First, added source hosts (*φ*_*n*_ > 0) are predicted to increase focal host infection prevalence, unless the added host has sufficiently strong negative competitive effects on other source hosts. As an example, consider a system where larger source hosts are only present in communities with more species, i.e., there is a positive correlation between net production of infectious propagules and host arrival order. In such a system there is typically a positive relationship between host species richness and focal host infection prevalence. However, a negative relationship can arise if larger source hosts are also stronger interspecific competitors because the later arriving hosts suppress host densities.

Second, added sink hosts (*φ*_*n*_ < 0) are predicted to decrease focal host infection prevalence, unless the added host has sufficiently strong negative competitive effects on other sink hosts. As an example, consider a system where large sink hosts are only present in communities with more species, i.e., there is a negative correlation between net production of infectious propagules and host arrival order. In such a system there is typically a negative relationship between host species richness and focal host infection prevalence. However, U-shaped relationships between host species richness and focal host infection prevalence are possible if sink hosts also have strong interspecific competitive effects on other sink hosts. In this case, the densities of the sink hosts are suppressed as more species are added to the community, which leads to increases in infectious propagule density and focal host infection prevalence.

#### Focal host infected density

Because the predicted effects of host additions on focal host infected density and infected density are often the same, the relationships of those two metrics with host species richness are often the same (compare blue squares in Figure 4B,C and all curves in Figure 4E,F). For example, if large sink hosts are only present in a community with more species (Figure 4,E,F), then focal host infection prevalence and focal host infected density both decrease with increased host richness (black circles), unless sink hosts also have strong interspecific competitive effects on other sink hosts (blue squares). However, the relationships can differ if the focal host is (i) the focal host is more strongly regulated by the disease or interspecific competition than by intraspecific competition or (ii) the focal host is a sink or small source and other hosts are large sources. Both of these conditions explain why adding large source hosts in Figure 4BC causes increased infection prevalence (black circles in Figure 4B) and decreased infected density (black circles in Figure 4C).

#### Explaining prior results about richness-disease relationships

The sensitivities help explain prior results for environmental transmission models (Table 3); see Appendix S3.2 for mathematical details. First, in prior studies (Rudge et al., 2013; Fenton et al., 2015; Espira et al., 2022) added high competence sources increased 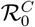 and focal host infection prevalence whereas added low competence sinks decreased both metrics because those studies assumed no interspecific competition 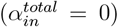. Second, other studies of two-host models (Cáceres et al., 2014; Searle et al., 2016) assumed the focal host was a source (*φ*_1_ > 0) and there was strong interspecific competition (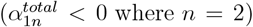where *n* = 2). Consequently, a low competence alternative sink host will necessarily decrease 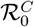 and focal host infection prevalence (Cáceres et al., 2014), but an added alternative source host can increase or decrease focal host infection prevalence (Searle et al., 2016). Third, the added alternative source host in Searle et al. (2016) increased focal host infection prevalence, but decreased focal host infected density. This difference is due to large changes in focal host density (mechanism 3) caused by the focal host being a small source and the alternative host being a very large source; I return this particular example in Section “Application to model of *Daphnia*-fungus system.”

Translating the predictions to direct transmission models explains prior results for density-dependent (DDDT) and frequency-dependent (FDDT) direct transmission models (Table 3); mathematical details are provided in Appendix S3.1. Under DDDT, all hosts have high competence 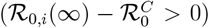 and are sources (*φ*_*i*_ > 0). Thus, equations (9) and (10) are positive in the absence of interspecific competition 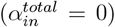 and negative only if interspecific competition is sufficiently strong 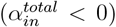. This explains why host additions always increased 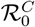 and focal host infection prevalence in the absence of interspecific competition (Begon et al., 1992; Dobson, 2004; Rudolf and Antonovics, 2005; McCormack and Allen, 2007; Faust et al., 2017; Huang et al., 2019), but host additions decreased those metrics when interspecific competition was strong (Bowers and Turner, 1997; Peixoto and Abramson, 2006; Joseph et al., 2013; Mihaljevic et al., 2014; O’Regan et al., 2015).

Under FDDT, an alternative host is a source if it has sufficiently high interspecific transmission rates and a sink otherwise (mathematical details given in Special Cases B and C of Appendix S2.4.2 and Special Case 2 of Appendix S2.5.2). Prior studies of FDDT models assume interspecific transmission rates are fractions of intraspecific transmission rates (i.e, 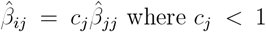 where *c*_*j*_ < 1), which means an alternative host is source only if it has sufficiently high competence. As a result, additions of lower competence alternative hosts decreased 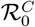 and focal host infection prevalence (Dobson, 2004; Rudolf and Antonovics, 2005; Faust et al., 2017; Mihaljevic et al., 2014), but additions of sufficiently high competence alternative hosts increased both metrics (Joseph et al., 2013; O’Regan et al., 2015).

### 4.5 Sensitivities of metrics to pathogen transmission mechanism

Prior studies (e.g., Dobson 2004; Rudolf and Antonovics 2005; Mihaljevic et al. 2014; Faust et al. 2017) suggest that host additions decrease all three disease metrics more often under frequency-dependent direct transmission (FDDT) than density-dependent direct transmission (DDDT), but not always for focal host infection prevalence (Cortez and Duffy, 2021; Cortez, 2021). I explore this further by comparing how the response of each disease metric to host additions depends on the pathogen transmission mechanism.

To explain the approach, I use 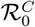 as a concrete example (see Appendix S1.4.3 for mathematical details). The goal is to determine if larger increases or larger decreases in 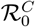 occur under FDDT or DDDT when a subcommunity of *m* resident species (*m* < *n*) has species *m* + 1, …, *n* added to it. To do this, I use a new parameter, *q*, that converts model (2) from a FDDT form (*δ* = 0) to a DDDT form 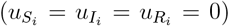. To allow for a fair comparison between the FDDT and DDDT forms of the model, the change of parameters varies the host uptake and degradation rates while holding constant the value of 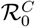 for the *m*-species subcommunity. I then compute how 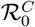 for the full *n*-species community changes using the sensitivity 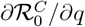. Positive values mean host additions cause larger decreases (or smaller increases) in 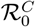 under FDDT than DDDT. Negative values mean that host additions cause larger increases (or smaller decreases) in 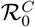 under FDDT than DDDT. The same approach is used for focal host infection prevalence and infected density, with the only difference being that the change of parameters holds constant the host and propagule densities at the endemic equilibrium for the *m*-species subcommunity; this allows for a fair comparison with those two metrics (see Appendix S1.5.4 for details).

#### Community reproduction number

For 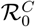, host additions cause larger decreases (or smaller increases) in 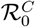 under FDDT than DDDT if (i) there is no interspecific competition (Figure 5A), (ii) added host species are sufficiently weak intraspecific and interspecific competitors, or (iii) added host species have higher uptake rates than resident host species (black circles and red triangles in Figure 5D). However, additions can cause larger increases (or smaller decreases) in 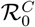 under FDDT than DDDT when added host species are stronger intraspecific and interspecific competitors and have lower uptake rates than resident host species (blue squares in Figure 5D).

**Figure 5:**
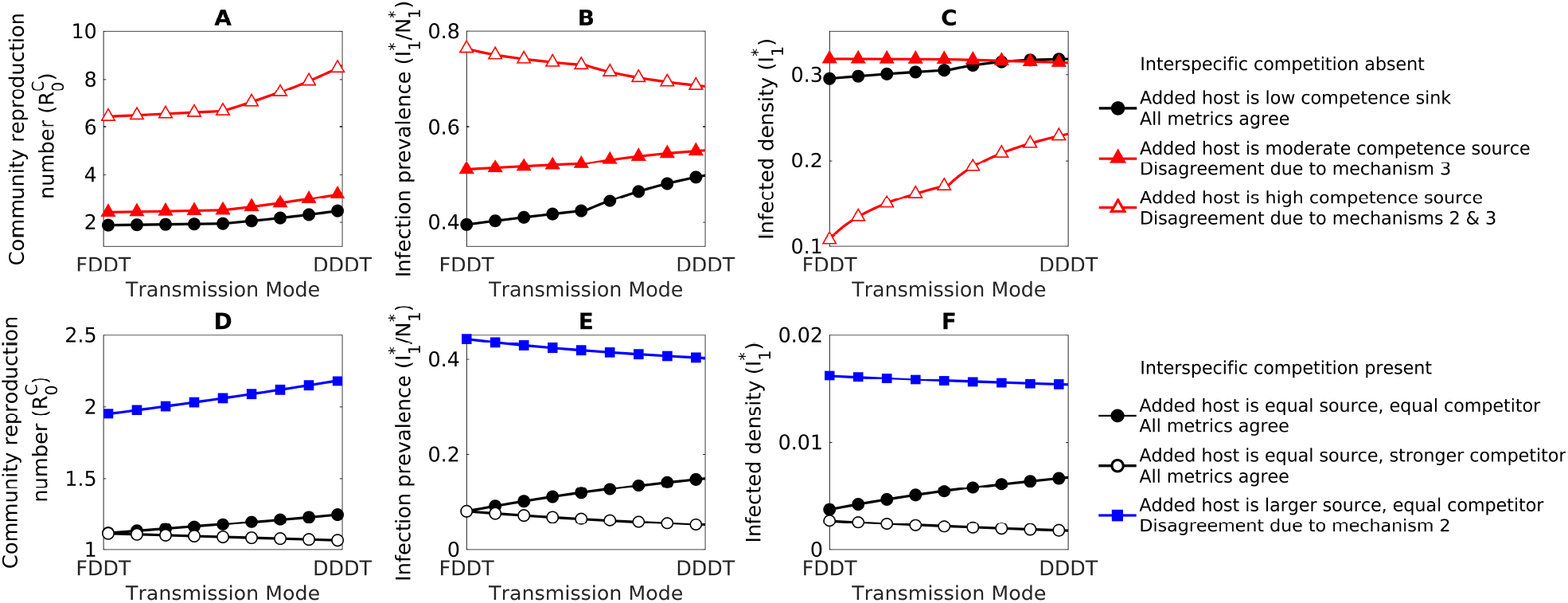
Host additions cause larger decreases (or smaller increases) in each metric under frequency-dependent direct transmission (FDDT) than density-dependent direct transmission (DDDT), unless the added host species have sufficiently high competence, the added host species are sufficiently strong interspecific competitors, or there are large changes in focal host density. In each panel, a change of parameters is used to convert the environmental transmission model from a FDDT form to a DDDT form; see Appendices S1.4.3 and S1.5.4 for details. Positive slopes mean large decreases or smaller increases in the metric under FDDT than DDDT when a host is added to a community of resident host species; negative slopes mean larger increases or smaller decreases in the metric under FDDT and DDDT. (A,D) Host additions cause larger decreases in 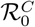 under FDDT than DDDT (all curves in A, filled black circles and blue squares in D), unless interspecific competition is sufficiently strong (open black circles in D). (B,E) Host additions cause larger decreases in focal host infection prevalence 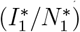 under FDDT than DDDT (black circles and filled red triangles in B, filled black circles in E), unless the added host is a sufficiently large source (open red triangles in B) or interspecific competition is sufficiently strong (open black circles and blue squares in E). (C,F) The responses in focal host infected density 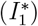 are the same as those in focal host infection prevalence (black circles in C, all curves in F), unless there are large changes in focal host density (mechanism 3), which occurs when the focal host is more strongly regulated by disease and interspecific competition than by intraspecific competition (open and filled red triangles in C). In each row, curves of the same color are from the same parameterization of a four-host version of model (2); see Appendix S4 for parameter values.

#### Focal host infection prevalence

Host additions are more likely to cause larger decreases (or smaller increases) in focal host infection prevalence under FDDT than DDDT when the added host species (i) are weak intraspecific and interspecific competitors and (ii) have lower competence than resident host species. For direct transmission models where interspecific transmission rates are fractions of intraspecific transmission rates (i.e, 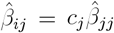where *c*_*j*_ < 1), condition (ii) implies that added host species have relatively lower rates of interspecific transmission (smaller *c*_*j*_) than resident host species; see Appendix S2.5.4 for details. Host additions are more likely to cause larger increases (or smaller decreases) in focal host infection prevalence under FDDT than DDDT when the opposite conditions hold. Consequently, in the absence of interspecific competition, an added low competence host causes smaller increases in focal host infection prevalence under FDDT than DDDT (black circles and blue squares in Figure 5B), but an added high competence host can cause the opposite (red triangles in Figure 5B). When interspecific competition is present, an added host can cause larger increases in focal host infection prevalence under FDDT than DDDT when it is a strong interspecific competitor or has high competence (blue squares and red triangles in Figure 5E, respectively).

#### Focal host infected density

The predictions for focal host infected density are often the same as focal host infection prevalence. However, the predictions can be reversed if host additions cause large changes in focal host density (mechanism 3) because the focal host is more strongly regulated by disease or interspecific competition than intraspecific competition.

#### Comparison of metrics

The pathogen transmission mechanism often has effects of the same sign on all three metrics (black circles in Figure 5). In these cases, host additions cause larger decreases (or smaller increases) in all metrics either under FDDT or under DDDT. However, the pathogen transmission mechanism can have effects of different signs on the metrics under two conditions. First, responses in 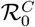 can differ in sign from the other two metrics when the added host species are large sources (red triangles in Figure 5D-F). This is due to density-mediated feedbacks (mechanism 2) where the added large sources increase infectious propagule density, which increases disease-induced mortality in the focal host. Second, as noted above, the responses in focal host infection prevalence and infected density can have opposite signs. This occurs when there are large changes in focal host density because the focal host is more strongly regulated by disease or interspecific competition than intraspecific competition (blue squares and red triangles in Figure 5BC).

#### Explaining prior results about richness-disease relationships

These predictions explain patterns in prior studies comparing DDDT and FDDT models. First, the 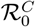 predictions explain why 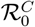 increased more often under DDDT than FDDT in the absence of interspecific competition (Dobson, 2004; Rudolf and Antonovics, 2005; Joseph et al., 2013; Mihaljevic et al., 2014), but not when interspecific competition was stronger (Joseph et al., 2013; Mihaljevic et al., 2014). Second, Rudolf and Antonovics (2005) and Faust et al. (2017) observed that host additions decreased focal host infection prevalence more often under FDDT than DDDT because those studies assumed low interspecific transmission and no interspecific competition, respectively.

### 4.6 Application to model of *Daphnia* -fungus system

To show how my theory can help explain the differing effects host additions have on the disease metrics, I apply my theory to the parameterized model in Searle et al. (2016). Here, I present a summary of the results and refer the reader to Appendix S3.2.5 for details about the analysis and to Searle et al. (2016) for details about the empirical system and model parameterization.

The model describes the dynamics of two host species, *Daphnia dentifera* and *D. lumholtzi*, and an environmentally transmitted fungal pathogen, *Metschnikowia bicuspidata*. The model assumes Lotka-Volterra competition and the structure is identical to model (2) except there are no recovered classes (*R*_*i*_) because infected individuals do not recover from infection. To avoid confusion, the host species are indexed using *i* = *D* for *D. dentifera* and *i* = *L* for *D. lumholtzi*.

The estimated parameter values from Searle et al. (2016) show that *D. dentifera* has lower competence than *D. lumholtzi*, i.e., ℛ_0,*D*_(∞) = 1.7 is less than ℛ_0,*L*_(∞) = 6, and is a smaller source, i.e., 0 < *φ*_*D*_ < *φ*_*L*_. In addition, *D. dentifera* is a weaker intraspecific and interspecific competitor than *D. lumholtzi*, i.e., *α*_*DD*_ < *α*_*LL*_ and *α*_*DL*_ < 0 < *α*_*LD*_ in equation (3). The small positive direct effect of *D. dentifera* on *D. lumholtzi* (0 < *α*_*LD*_) is likely an artifact due to fitting noisy data and it is expected that the actual value is small and negative (Searle et al., 2016). In any case, the predicted effects of host additions on each disease metric are qualitatively unchanged if a small positive or a small negative value is used for *α*_*DL*_; see Appendix S3.2.5. For the estimated parameter values, the host species can stably coexist in the absence of the pathogen (at a two-host disease-free equilibrium) and each host can stably coexist with the pathogen (at a one-host endemic equilibrium).

#### Community reproduction number

The pathogen’s reproduction number is lowest when *D. dentifera* is alone (ℛ_0,*D*_ = 1.5), highest when *D. lumholtzi* is alone 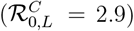, and intermediate when both hosts are present 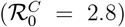. This means that 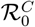 increases with additions of *D. lumholtzi* and decreases with additions of *D. dentifera*. The explanations for the two different effects on 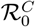 are the following.

First, the addition of *D. lumholtzi* to a community with just *D. dentifera* increases 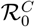. This is because the addition of *D. lumholtzi* causes an increase in the density of a high competence host (*D. lumholtzi*) and the decrease in the density of a low competence host (*D. dentifera*) due to competition. Mathematically, we see this in equation (9) where the first and second terms, respectively, are positive because (1) the competence of *D. lumholtzi* is large, making 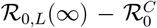 large and positive; (2) the competence of *D. dentifera* is small, making 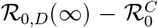 negative; and (3) *D. lumholtzi* has a negative competitive effect on *D. dentifera*, making 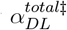 negative. Overall, equation (9) shows that increased density of *D. lumholtzi* increases 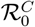, which agrees with 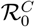 being larger when *D. lumholtzi* is added to a community with just *D. dentifera*.

Second, the addition of *D. dentifera* to a community with just *D. lumholtzi* reduces 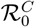. This is because the addition of *D. dentifera* causes a large increase in the density of a lower competence host (*D. dentifera*) and a small increase in the density of a higher competence host (*D. lumholtzi*) due to the positive interspecific effect. Mathematically, we see this in equation (9) where the first term is large and negative and the second term is smaller and positive, respectively, because (1) the competence of *D. dentifera* is small, making 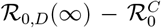 negative; (2) the competence of *D. lumholtzi* is large, making 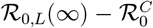 smaller and positive; and (3) *D. dentifera* has a small positive direct effect on *D. lumholtzi*, making 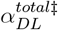 positive and small. Overall, equation (9) shows that increased density of *D. dentifera* decreases 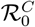, which agrees with 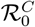 being larger when *D. dentifera* is added to a community with just *D. lumholtzi*.

#### Focal host infection prevalence

*D. dentifera* and *D. lumholtzi* both cause infection prevalence in the other species to increase. The explanations for the similar responses are the following.

First, the addition of *D. lumholtzi* causes infection prevalence in *D. dentifera* to increase. In this case, the positive effect of adding the large source *D. lumholtzi* is greater than the negative effect of reduced density of the small source *D. dentifera*. Mathematically, we see this in equation (10) where the first term is large and positive and the second term is smaller and negative, respectively, because (1) *D. lumholtzi* is a large source (*φ*_*L*_ > 0 large); (2) *D. dentifera* is a small source (*φ*_*D*_ > 0 small); and (3) *D. lumholtzi* has a negative competitive effect on *D. dentifera* 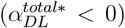. Thus, equation (10) shows that increased density of *D. lumholtzi* increases infection prevalence in *D. dentifera*, which agrees with the observed response.

Second, the addition of *D. dentifera* causes infection prevalence in *D. lumholtzi* to increase. This is caused by the positive effect of adding the small source *D. dentifera* and the positive effect of increased density of the large source *D. lumholtzi*. Mathematically, we see this in equation (10) where the first and second terms are positive, respectively, because (1) *D. dentifera* is a small source (*φ*_*D*_ > 0 large); (2) *D. lumholtzi* is a large source (*φ*_*L*_ > 0 small); and (3) *D. dentifera* has a positive effect on *D. lumholtzi* 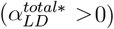. Thus, equation (10) shows that increased density of *D. dentifera* increases infection prevalence in *D. lumholtzi*, which agrees with the observed response.

#### Focal host infected density

The density of infected *D. dentifera* decreases whereas the density of infected *D. lumholtzi* increases when the other host species is added. The explanations for the two different effects on infected density are the following.

First, the addition of *D. lumholtzi* causes the density of infected *D. dentifera* to decrease. Infected density and infection prevalence in *D. dentifera* have responses of opposite signs because the addition of *D. lumholtzi* causes a large decrease in the total density of *D. dentifera*. The large decrease in total density of *D. dentifera* is due to *D. lumholtzi* being a strong interspecific competitor. This aligns with condition (ii) because the focal host *D. dentifera* is more strongly regulated by interspecific competition than by intraspecific competition.

Second, the addition of *D. dentifera* causes the density of infected *D. lumholtzi* to increase. Infected density and infection prevalence in *D. lumholtzi* have responses of the same sign because the addition of *D. dentifera* does not cause a large increase or decrease in the total density of *D. lumholtzi*. Moreover, condition (i) is not satisfied because *D. lumholtzi* is strongly regulated by intraspecific competition and weakly regulated by disease and interspecific competition. Condition (ii) is not satisfied because the focal host *D. lumholtzi* is not a small source.

#### Comparison of metrics

For both host species, the addition of a second host has different effects on the three metrics. The addition of *D. lumoltzi* causes 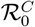 and infection prevalence in *D. dentifera* to increase, but infected density in *D. dentifera* to decrease. The differing response for infected density is due large changes in the total density of *D. dentifera* (mechanism 3), driven by *D. dentifera* being more strongly regulated by interspecific competition with *D. lumholtzi* than by intraspecific competition (condition ii). In comparison, the addition of *D. dentifera* causes 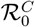 to decrease but infection prevalence and infected density in *D. lumholtzi* to increase. The differing response for 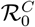 is due to differences in the definition of competence and source/sink (mechanism 1). In particular, *D. dentifera* is a lower competence host, which leads to lower 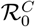, but also a small source, which leads to higher infection prevalence and infected density in *D. lumholtzi*.

## 5 Discussion

Across empirical studies, increased host species richness can decrease (dilute) or increase (amplify) disease risk (Keesing et al., 2010; Randolph and Dobson, 2012; Salkeld et al., 2013; Wood et al., 2014; Civitello et al., 2015; Ostfeld and Keesing, 2017; Rohr et al., 2019). To understand the variation in responses, theory is needed that identifies the rules driving context-dependent biodiversity-disease relationships and explains how predicted outcomes depend on the choice of disease metric (Buhnerkempe et al., 2015; Halsey, 2019; Rohr et al., 2019). Prior theoretical studies have shown that disease levels and risk can increase or decreases with increased host species richness, depending on mechanisms related to host competence, changes in host densities, and transmission (Dobson, 2004; Joseph et al., 2013; Mihaljevic et al., 2014; Faust et al., 2017; Cortez, 2021). The main goal of this study was to synthesize this prior theoretical work and clarify how differences between model predictions were driven by differing assumptions about host competence, interspecific host competition, the pathogen transmission mechanism, and the metric of disease. My results (Tables 1 and 2) help explain the variation in predictions across theoretical studies and identify the rules governing the shapes of host species richness-disease relationships.

**Table 2:** Predicted responses of three metric of disease to variation in components of alternative host competence

| Component of<br>alternative host<br>competence | Community<br>Reproduction No.<br>($\mathcal{R}_0^C$) | Refs.<sup>†</sup> | Focal host infection prevalence<br>($I_1^*/N_1^*$) | Refs.<sup>†</sup> | Focal host infection density<br>($I_1^*$) |
| --- | --- | --- | --- | --- | --- |
| Increased shedding ($\chi_i$) | Always increases | [1]-[6] | Always increases | [7],[8] | Increases unless condition (i) |
| Increased uptake ($u_{S_i}$) | Always decreases | [6] | Always decreases | [7],[8] | Decreases unless condition (i) |
| Increased transmission<br>rate ($\beta_i$) | Always increases | [4],[6] | Increases unless<br>(1) alternative host is a source and interspecific competition<br>is strong, or<br>(2) alternative host has larger negative competitive effects<br>on sink hosts than source hosts and infected individuals<br>are weaker competitors than susceptible individuals | [7],[8] | Same as infection prevalence<br>unless conditions (i) or (ii) |
| Increased mortality<br>rate ($\mu_i$) | Always decreases | | Decreases unless<br>(1) alternative host is a large sink<br>(2) alternative host has larger negative competitive effects<br>on sinks hosts than source hosts | [7],[8] | Same as infection prevalence<br>unless conditions (i) or (ii) |
| Increased recovery<br>rate ($\nu_i$) | Always decreases | [6] | Always decreases | [7],[8] | Same as infection prevalence<br>unless conditions (i) or (ii) |

Responses in focal host infected density can differ from the responses in focal host infection prevalence when (i) the focal host is more strongly regulated by disease and interspecific competition with alternative hosts than by intraspecific competition or (ii) the focal host is a sink or small source and some alternative hosts are large sources.
† Prior studies whose predictions are explained or generalized are [1] Dobson (2004), [2] (Huang et al., 2019), [3] Fenton et al.
(2015), [4] O'Regan et al. (2015), [5] Joseph et al. (2013), [6] Espira et al. (2022), [7] Cortez and Duffy (2021), and [8] Cortez
(2021). Mathematical details about each study are provided in Appendix S3.

**Table 3:**
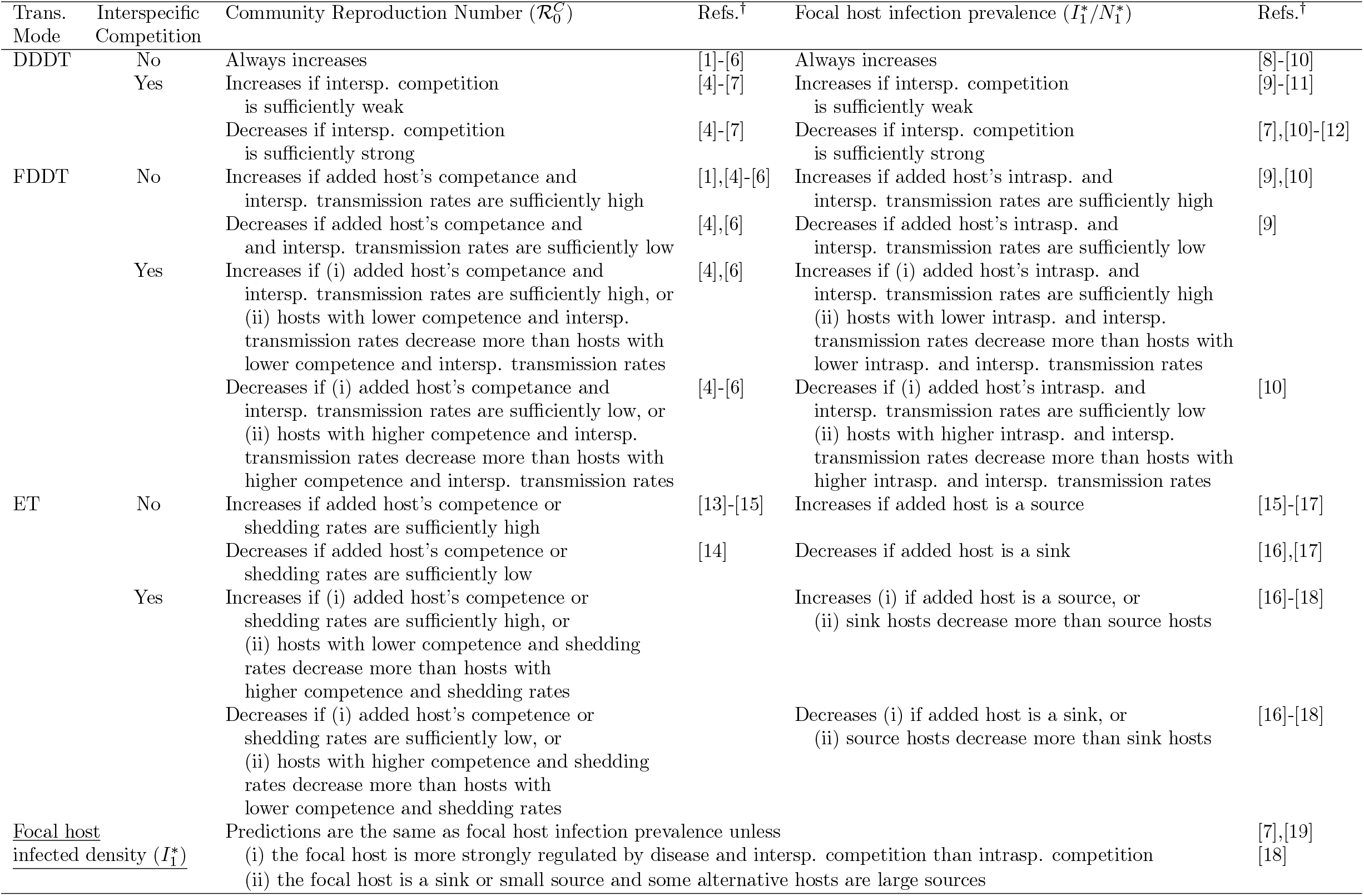
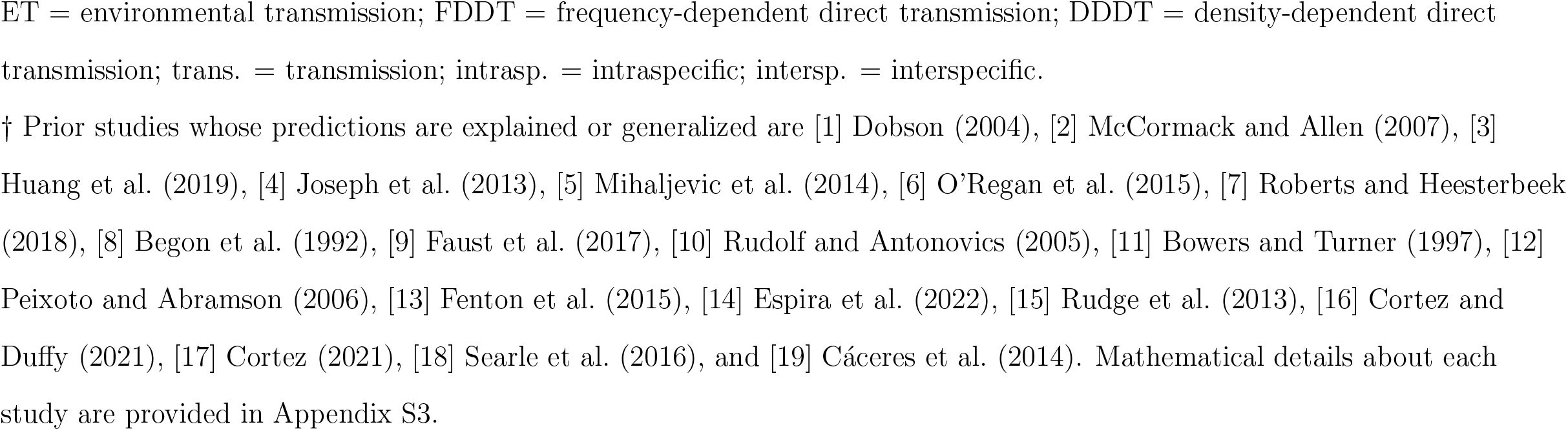
Predicted responses of three metrics of disease to the addition of a host species to a community

| Trans. Mode | Interspecific Competition | Community Reproduction Number ($\mathcal{R}_0^C$) | Refs.<sup>†</sup> | Focal host infection prevalence ($I_1^*/N_1^*$) | Refs.<sup>†</sup> |
| --- | --- | --- | --- | --- | --- |
| DDDT | No | Always increases | [1]-[6] | Always increases | [8]-[10] |
|  | Yes | Increases if intersp. competition is sufficiently weak<br>Decreases if intersp. competition is sufficiently strong | [4]-[7]<br>[4]-[7] | Increases if intersp. competition is sufficiently weak<br>Decreases if intersp. competition is sufficiently strong | [9]-[11]<br>[7],[10]-[12] |
| FDDT | No | Increases if added host's competence and intersp. transmission rates are sufficiently high | [1],[4]-[6] | Increases if added host's intrasp. and intersp. transmission rates are sufficiently high | [9],[10] |
|  |  | Decreases if added host's competence and intersp. transmission rates are sufficiently low | [4],[6] | Decreases if added host's intrasp. and intersp. transmission rates are sufficiently low | [9] |
|  | Yes | Increases if (i) added host's competence and intersp. transmission rates are sufficiently high, or (ii) hosts with lower competence and intersp. transmission rates decrease more than hosts with lower competence and intersp. transmission rates | [4],[6] | Increases if (i) added host's intrasp. and intersp. transmission rates are sufficiently high (ii) hosts with lower intrasp. and intersp. transmission rates decrease more than hosts with lower intrasp. and intersp. transmission rates |  |
|  |  | Decreases if (i) added host's competence and intersp. transmission rates are sufficiently low, or (ii) hosts with higher competence and intersp. transmission rates decrease more than hosts with higher competence and intersp. transmission rates | [4]-[6] | Decreases if (i) added host's intrasp. and intersp. transmission rates are sufficiently low (ii) hosts with higher intrasp. and intersp. transmission rates decrease more than hosts with higher intrasp. and intersp. transmission rates | [10] |
| ET | No | Increases if added host's competence or shedding rates are sufficiently high | [13]-[15] | Increases if added host is a source | [15]-[17] |
|  |  | Decreases if added host's competence or shedding rates are sufficiently low | [14] | Decreases if added host is a sink | [16],[17] |
|  | Yes | Increases if (i) added host's competence or shedding rates are sufficiently high, or (ii) hosts with lower competence and shedding rates decrease more than hosts with higher competence and shedding rates |  | Increases (i) if added host is a source, or (ii) sink hosts decrease more than source hosts | [16]-[18] |
|  |  | Decreases if (i) added host's competence or shedding rates are sufficiently low, or (ii) hosts with higher competence and shedding rates decrease more than hosts with lower competence and shedding rates |  | Decreases (i) if added host is a sink, or (ii) source hosts decrease more than sink hosts | [16]-[18] |
| <u>Focal host infected density ($I_1^*$)</u> |  | Predictions are the same as focal host infection prevalence unless<br>(i) the focal host is more strongly regulated by disease and intersp. competition than intrasp. competition<br>(ii) the focal host is a sink or small source and some alternative hosts are large sources |  |  | [7],[19]<br>[18] |

ET = environmental transmission; FDDT = frequency-dependent direct transmission; DDDT = density-dependent direct transmission; trans. = transmission; intrasp. = intraspecific; intersp. = interspecific. † Prior studies whose predictions are explained or generalized are [1] Dobson (2004), [2] McCormack and Allen (2007), [3] Huang et al. (2019), [4] Joseph et al. (2013), [5] Mihaljevic et al. (2014), [6] O'Regan et al. (2015), [7] Roberts and Heesterbeek (2018), [8] Begon et al. (1992), [9] Faust et al. (2017), [10] Rudolf and Antonovics (2005), [11] Bowers and Turner (1997), [12] Peixoto and Abramson (2006), [13] Fenton et al. (2015), [14] Espira et al. (2022), [15] Rudge et al. (2013), [16] Cortez and Duffy (2021), [17] Cortez (2021), [18] Searle et al. (2016), and [19] Cáceres et al. (2014). Mathematical details about each study are provided in Appendix S3.

Accurate predictions about host richness-disease relationships require measurements of the ability of each host species to transmit the pathogen, and critically, how that ability is measured depends on the disease metric of interest. Specifically, predictions about 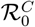require estimates of life-time production of infectious propagules given exposure (i.e, competence, ℛ_0,*i*_(∞), as defined in Merrill and Johnson (2020)). In comparison, predictions about focal host infection prevalence and infected density require estimates of instantaneous per capita net production rates of infectious propagules (*ϕ*_*i*_). In prior studies, a host’s ability to transmit the pathogen has been measured in many ways, including infection intensity given exposure (Johnson et al., 2013), quantities that include transmission rates to conspecific and heterospecifics (Gervasi et al., 2015; Downs et al., 2019), and the pathogen’s basic reproduction number for each species when alone (Strauss et al., 2015). These metrics differ from the current study’s definitions of competence and sink/source because they include additional factors (e.g., host densities) or omit factors (e.g., the average length of infection, 1*/*[*m*_*i*_ + *μ*_*i*_ + *ν*_*i*_]). As a result, the metrics used in these empirical studies could lead to inaccurate predictions about host species richness-disease relationships in nature.

Accurate predictions about host richness-disease relationships also require knowledge about density of each host species responses to altered host species richness. One component of this (only relevant to infection prevalence and infected density) is changes in host densities due to density-mediated feedbacks driven by disease-induced mortality (mechanism 2). The other component (relevant to all metrics) is changes in host densities due to interspecific competition. Prior theoretical studies (Rudolf and Antonovics, 2005; Joseph et al., 2013; Mihaljevic et al., 2014; O’Regan et al., 2015; Faust et al., 2017) have shown that host richness-disease relationships can differ depending on whether host additions and removals have additive effects on host density (corresponding to no interspecific competition) or substitutive effects on host density (corresponding to strong, symmetric interspecific competition where total host density is held constant); these effects were referred to as subtractive and compensatory effects in Joseph et al. (2013). Because both of these scenarios assume symmetric competitive effects, my results extend this prior work by explaining how asymmetric interspecific competitive effects can be important for shaping host richness-disease relationships. For example, added low competence sink hosts can increase all disease metrics if they have stronger competitive effects on low competence sink hosts than high competence source hosts (Figure 4).

In combination, my results show that accurate predictions about host richness-disease relationships require detailed information about specific measures of each host’s ability to transmit the pathogen and changes in host densities in response to altered host species richness. While all of these quantities can be estimated in studies of tractable laboratory systems (Cáceres et al., 2014; Strauss et al., 2015; Searle et al., 2016) or systems where interspecific competition is likely negligible (Fenton et al., 2015), they are likely to be difficult to estimate for many field systems. My analytical results and predictions suggest that transmission abilities (competence and sink/source) are the most important quantities to estimate. The reasoning is that if the transmission ability of an added or removed host correctly predicts the response in a metric, then this suggests the effects of altered host densities are either small in magnitude or in the same direction (i.e., a synergistic effect). For example, Venesky et al. (2014) estimated how the presence and absence of three amphibian species affected the intensity of *Batrachochytrium dendrobatidis* infections in laboratory experiments and found qualitatively similar relationships between presence/absence and infection prevalence in a field study. This agreement suggests that estimates of changes in host densities are not needed to accurately predict the qualitative effect (i.e., increases or decreases in infection prevalence) of altered host species richness. However, an important caveat to the above is that one needs to use the correct quantity when estimating the ability of each host species to transmit the disease; the Venesky et al. (2014) study is not ideal in this regard because its metric of transmission ability (infection intensity) does not account for the production of new infectious propagules by infected individuals. A second important caveat is that without predictions about changes in host densities, it is impossible to predict, *a priori*, if infection prevalence and infected density will respond in opposite directions.

My work builds on prior studies that compared the responses of different disease metrics (Roche et al., 2012; Roberts and Heesterbeek, 2018) by identifying when and why the three metrics of disease change in the same or different directions. Agreement across all metrics is an ideal situation for disease management strategies that use culling or host removal (Laurenson et al., 2004; Donnelly et al., 2006) because disease risk and disease burden can be simultaneously minimized by removing or suppressing specific alternative host species in a community. However, cases where the metrics respond differently imply the presence of trade-offs. Opposing responses in 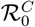 and focal host infection prevalence (mechanism 1 or 2) produce a trade-off between the risk of an outbreak and the risk of a focal host individual becoming infected. For example, additions of *D. dentifera* lower risk the risk of an outbreak for *D. lumhotzi* by reducing 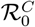, but increase the risk of infection for *D. lumhotzi* when an outbreak occurs by increasing infection prevalence in *D. lumhotzi* (see “Application to model of *Daphnia*-fungus systems”). In comparison, opposing responses in focal host infection prevalence and infected density (mechanism 2) produce a trade-off between infection risk for focal host individuals (infection prevalence) and the risk of spillover to other species (for which infected density is considered a better metric; Roche et al. 2012; Wood and Lafferty 2013). Thus, my work helps identify when disease management or conservation strategies need to account for trade-offs between different metrics of disease risk.

My results support prior claims that altered host species richness has context-dependent effects on disease risk (Randolph and Dobson, 2012; Salkeld et al., 2013; Wood et al., 2016), while also providing testable predictions about the mechanisms driving those context-dependent effects (Table 2). For example, increases in focal host infection prevalence are predicted to be driven by later added hosts being sources (black curves in Figure 4B) or having stronger competitive effects on sink hosts than source hosts (blue curves in Figure 4E). In comparison, decreases in focal host infection prevalence are predicted to be driven by later added hosts being sinks (black curves in Figure 4E) or having stronger competitive effects on source hosts than sink hosts (blue curves in Figure 4B). Pairing my work with knowledge about how host competence (or sink/source) and competitive effect are correlated with community (dis)assembly also could help explain patterns identified across empirical studies. As one example, based on life-history theory and hypotheses about pathogen adaptation, prior studies (reviewed in Ostfeld and Keesing (2012); Joseph et al. (2013); Huang et al. (2019)) have argued that more competent host species are more likely to be present in low richness communities whereas higher competence host species will only be present in high richness communities. The resulting negative relationship between arrival order and competence can lead to negative relationships between richness and disease (Joseph et al., 2013). However, my results explain why interspecific competition could alter those predictions and cause positive relationships between richness and disease (bottom row of Figure 4) that were also observed in prior theoretical work (Joseph et al., 2013). As a second example, Halliday and Rohr (2019) found that biodiversity gradients driven by mechanisms associated with biodiversity loss (e.g., habitat fragmentation or urbanization) often result in negative host species richness-disease relationships whereas biodiversity gradients driven by other mechanisms (e.g., environmental heterogeneity or elevation) can yield positive or negative host species richness-disease relationships. My theory suggests that these patterns could be driven by different relationships between extinction order and competence or different relationships between extinction order and host densities (due to competition or increased mortality, e.g., due to habitat reduction). More generally, pairing my work with system-specific knowledge about correlations between species traits and extinction/arrival order could explain the different frequencies of unimodal and monotonic relationships between richness and disease in empirical studies (Halliday and Rohr, 2019).

While this study explains prior theoretical work and has the potential to explain empirical patterns, additional theory is needed to fully explain the observed variation in empirical host species richness-disease relationships. Specific areas directly related model (2) include incorporating the effects of dose-dependent infection and mortality rates (e.g., Clay et al. 2021) and transmission via both direct and environmental pathways (e.g., Eisenberg et al. 2013); these were not addressed in this study because the supplementary material is already at a ridiculous length. More generally, advances in theory are needed in four areas. First, theory is needed for other metrics used in empirical studies, including community prevalence (Suzán et al., 2009; Searle et al., 2011), disease incidence (Haas et al., 2011), infectious propagule density (Roberts and Heesterbeek, 2018; Cortez and Duffy, 2020), and infection intensity (Hechinger and Lafferty, 2005; Becker et al., 2014; Venesky et al., 2014). This is important because the metrics can provide different and complementary information about disease burden and disease risk. Second, more theory is needed to extend our understanding of multi-host systems where host species have non-competitive interspecific interactions (Rohr et al., 2015), including systems where pathogens obligately switch between host species during their life cycle (Hechinger and Lafferty, 2005; Hopkins et al., 2016). Third, additional theory is needed to explain why increased host species richness often, but not always, decreases disease for vector-borne pathogens in empirical studies (Ezenwa et al., 2006; Keesing et al., 2009; Moore and Borer, 2012; Xavier et al., 2012) and theoretical studies (Saul, 2003; Normal et al., 1999; Ogden and Tsao, 2009; Miller and Huppert, 2013). Because vectors are analogous to infectious propagules, some of the variation may be explained by the net production of infected vectors by each host species (analogous to *φ*_*i*_). However, this analogy is limited because vectors can have complex dynamics and behavior (e.g., biting preferences), which may influence how host additions affect disease metrics (Miller and Huppert, 2013; Marini et al., 2017). Fourth, new theory is needed to explain why host species can have temporally varying effects on disease metrics. For example, empirical studies show that an additional host species can have effects of the same sign at all points in time (e.g., always decreasing focal host infection prevalence; Strauss et al. 2015) or effects of varying sign (e.g., transition from decreased to increased focal host infection prevalence; Dallas et al. 2016). My results suggest a possible explanation for the temporally varying effects: a low competence source host could decrease infection prevalence early in time because decreases in 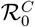 imply slower rates of spread, but also increase infection prevalence later in time because it is a net producer of infectious propagules. Recent work (Hanthanan Arachchilage et al., 2023) suggests sensitivity-based approaches could be fruitful, however new theory is needed to fully explain when and why host species have temporally variable effects on disease dynamics. Combined, advances in these four areas will help address a current need to understand how the interactions between parasites, pathogens, and their ecosystems shape disease burden and the risk of disease emergence (Cunningham et al., 2017).

This study has made a step towards addressing that need by providing a framework that unifies prior theoretical studies and identifies the rules governing the context-dependent relationships between host species richness and different metrics of disease. By providing testable predictions about underlying mechanisms, this study may help resolve some of the debate about the variation between and frequencies of different patterns of biodiversity and disease.

## 6 Acknowledgments

I thank MA Duffy, the Duffy lab, the Ecology Reading Group in FSU’s Department of Biological Science, and three anonymous reviewers for helpful comments and feedback on the manuscript. MHC was supported by the National Science Foundation under Award DEB-2015280.

## Box 1

Relationships between environmental and direct transmission models

Environmental transmission (ET) models can reduce down to models with density-dependent direct transmission (DDDT), frequency-dependent direct transmission (FDDT), or direct transmission that is intermediate to DDDT and FDDT (Eisenberg et al., 2013; Cortez and Weitz, 2013; Cortez and Duffy, 2021; Cortez, 2021; Espira et al., 2022). In general, an environmental transmission model reduces to a direct transmission model when the shedding rates and either the uptake rates or degradation rates are very large. Under these conditions infectious propagules only persist in the environment for short periods of time, and as a result a susceptible individual will encounter infectious propagules only if it is in close proximity to an infected individual. Thus, the environmental transmission pathogen effectively behaves like a direct transmission pathogen.

The ET model (2) reduces down to (i) a DDDT model when uptake is negligible compared to degradation 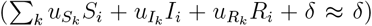, (ii) an FDDT model when degradation is negligible compared to uptake (*δ* ≈ 0) and all uptake rates are equal 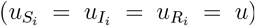, and (iii) a model with transmission intermediate to DDDT and FDDT when uptake and degradation are both non-negligible. Here, I provide a brief mathematical explanation; additional details for the 1-habitat and *n*-habitat models are provided in Appendices S1.2 and S2.2, respectively. When the shedding rates are large and the uptake or degradation rates are large, the infected density equation in the 1-habitat ET model (2) is approximately described by

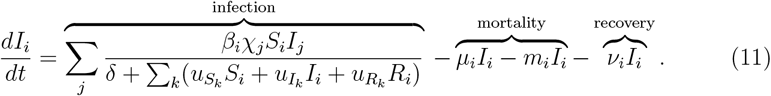

The first term in equation (11) is the sum of the direct transmission rates from each host *j* to host *i*; the second and third terms are the mortality rates due to disease and non-disease factors; and the fourth term is the recovery rate. I consider three cases. First, if uptake of infectious propagules is negligible 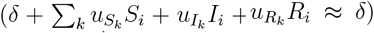, then the transmission rate in equation (11) reduces to 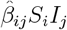 where 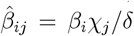. This is a DDDT rate because the rate of infection is proportional to the density of infected individuals (*I*_*j*_). Second, if degradation is negligible (*δ* ≈ 0) and all uptake rates are equal 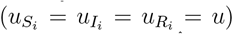, then the transmission rate in equation (11) reduces to 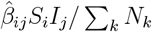 where 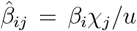. This is a FDDT rate because the rate of infection is proportional to the frequency of infected individuals in the community (*I*_*j*_*/* Σ_*k*_ *N*_*k*_). Third, if uptake and degradation rates are non-negligible, then the direct transmission rate in equation (11) is intermediate to FDDT and DDDT. The rate is more similar to DDDT when degradation is large and uptake small, and the rate is more similar to FDDT under the opposite conditions.

In total, the ET model (2) can be converted from a DDDT form to a FDDT form by varying the magnitudes of the infectious propagule degradation rate and host uptake rates. Consequently, the results for the ET model 2 apply to DDDT and FDDT models, after accounting for the restrictions on the parameter values. This allows one to simultaneously study ET, DDDT, and FDDT pathogens and identify how the transmission mechanism affects model predictions.

## Box 2

Connection between 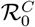 and theory for stage-structured models

The pathogen’s basic reproduction number, 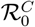, is the average number of new infections produced by a single infected individual in an otherwise susceptible community. The value of 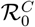 determines if a single infected individual will cause an outbreak 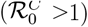 or not 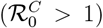. Conveniently, 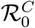 for the 1-habitat model (2) has an explicit formula,

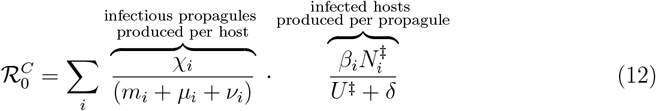

where 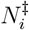 is the density of host *i* and 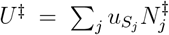 is the total uptake rate of infectious propagules at the disease-free equilibrium; see Appendix S1.4 for mathematical calculations. Each term in the sum defines the average number of new infections a single infected individual of host *i* will produce in its lifetime. In particular, the first fraction is the shedding rate (*χ*_*j*_) multiplied by the average duration of infection (1*/*[*μ*_*i*_+*m*_*i*_+*γ*_*i*_]), which equals the average lifetime production of infectious propagules by an infected individual of host *i*. The second fraction is the per infectious propagule infection rate for host 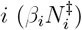 multiplied by the average lifespan of a spore (1*/*[*U*^‡^ + *δ*]), which equals the average lifetime production of new infected individuals of host *i* by an infectious propagule. The sensitivities of 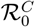 (e.g., 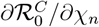 and 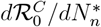) can be computed by differentiating equation (12).

For many models, explicit formulas for 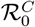 cannot be computed. This is true for the *n*-habitat models analyzed in Appendix S2 and direct transmission models with three or more host species. Consequently, one cannot compute the sensitivities of 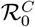 by differentiating a formula analogous to equation (12). However, the matrix theory developed for discrete-time models of stage-structured populations allows one to derive formulas for the sensitivities of 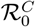, even when an explicit formula for 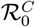 does not exist.

The intuition for the connection between 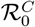 and stage-structured models is that infected individuals of each host species represent different life history “stages” for the pathogen. Matrix models for stage-structured populations describe how each stage contributes to the abundances of all stages in the next time step. Analogously, the Next Generation Matrix (Van den Driessche and Watmough, 2008; Diekmann et al., 2010) describes how the infected class in each host species contributes to infections in all host species in the next pathogen generation. As a specific example, the Next Generation Matrix for a 3-species version of model (2) is

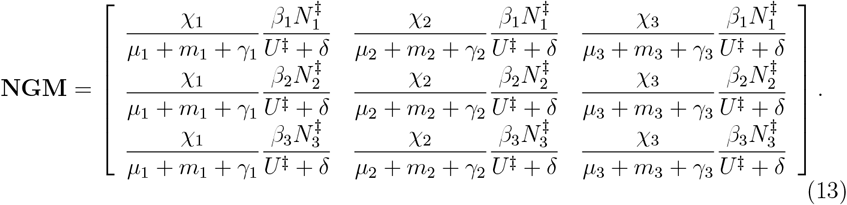

Entry *i, j* defines the average number of new infections in host *j* that a single infected individual of host *i* will produce in its lifetime. In particular, the first fraction in each entry (*χ*_*j*_*/*[*μ*_*j*_ + *ν*_*j*_ + *γ*_*j*_]) equals the average lifetime production of infectious propagules by an infected individual of host *j*. The second fraction in each entry 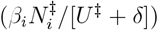 equals the average lifetime production of new infected individuals of host *i* by an infectious propagule. When the fractions are multiplied, they yield the average lifetime production of new infections in host *j* by a single infected individual of host *i*.

Importantly, the sensitivity theory for the asymptotic growth rates of stage-structured populations (Caswell, 2019) can be used to compute the sensitivities of 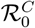. The reason is that for matrix models, the leading eigenvalue of the matrix defines the asymptotic growth rate of the species. Analogously, the leading eigenvalue of the Next Generation Matrix (13) defines the pathogen’s reproduction number 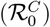, which in turn defines the asymptotic growth rate of an outbreak. In total, this means the theory for matrix models can be used to compute the sensitivities of 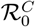 for the *n*-habitat model and direct transmission models with three or more host species, even though explicit formulas for 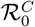 cannot be computed. Mathematical details are given in Appendix S2.4.

## Notes

### Competing Interest Statement

The authors have declared no competing interest.

### Summary of Updates

Paper was reformatted for clarify. xAdditional results sections were added to (i) illustrate host species-richness relationships when hosts are added to a community and (ii) demonstrate how the theory can be applied to a model parameterized to an empirical system.

