## Supplemental Appendices for "Comparing the effects of host species richness on different metrics of disease"

### Supplementary Information for "Comparing the effects of host species richness on different metrics of disease"

**Michael H. Cortez**

**S4 Parameter values for figures 109**

**References 114**

The matrix and vector notation used throughout is listed in Table S1. In summary, matrices are denoted using bold text (e.g., $\mathbf{M}$); the only exception is that script letters (e.g., $\mathcal{M}$) denote some matrices used to compute the basic reproduction number in Section S1.4. Vectors are denoted using overhead arrows (e.g., $\vec{v}$ or $\overrightarrow{v}$). Entries of matrices and vectors are denoted using subscripts ($\mathbf{M}_{ij}$) or brackets ($\mathbf{M}[i,j]$). Bracketed superscripts denote rows and columns that have been removed to create submatrices (e.g., $\mathbf{M}^{[\not{i},\not{j}]}$).

Table S1: Summary of matrix and vector notation

| Notation | Description |
|---|---|
| <u>Matrices</u> | |
| $\mathbf{M}$ | matrix |
| $\mathcal{M}$ | matrix, used to compute $\mathcal{R}_0$ |
| $\mathbf{M}_{ij}$, $\mathbf{M}[i,j]$ | entry $i,j$ of the matrix $\mathbf{M}$ |
| $\mathbf{M}_{\bullet i}$ | column $i$ of matrix $\mathbf{M}$ |
| $\mathbf{M}^{[\not{i},\not{j}]}$ | submatrix of $\mathbf{M}$ where row $i$ and column $j$ have been removed |
| $\mathbf{M}^{[-,\not{j}]}$ | submatrix of $\mathbf{M}$ where column $j$ has been removed |
| $\mathbf{M}^{[\not{i},-]}$ | submatrix of $\mathbf{M}$ where row $i$ has been removed |
| $\lvert\mathbf{M}\rvert$ | determinant of $\mathbf{M}$ |
| $\mathbf{D}_x$ | diagonal matrix with entries $x_i, ..., x_n$ |
| $\mathbf{A}_{I_j \to N_i}$ | matrix $\mathbf{A}$ where column $i$ is replaced with column $j$ of $\mathbf{B} + \mathbf{C}\mathbf{D}_\Gamma$; see Remark S1.5 |
| $\mathbf{0}_{a\times b}$ | matrix of zeros of size $a \times b$ |
| <u>Vectors</u> | |
| $\vec{v}$ | vector with entries $\vec{v}[1], ..., \vec{v}[n]$ |
| $\overrightarrow{\beta X}$ | vector with entries $\beta_1 X_1, ..., \beta_n X_n$ |
| $\vec{v}^{[\not{i}]}$ | vector $\vec{v}$ with entry $i$ removed |
| $\vec{0}_n$ | $n$-dimensional vector of zeros |
| $\vec{v}^{\mathrm{T}}$ | transpose of a vector |
| <u>Quantities evaluated at equilibria</u> | |
| $*$ | quantities evaluated at the $n$-host endemic equilibrium, $p^*$ |
| $'$ | quantities evaluated at the $m$-host ($m < n$) endemic equilibrium, $p'$ |
| $\ddagger$ | quantities evaluated at the $n$-host disease-free equilibrium, $p^\ddagger$ |
| $\dagger$ | quantities evaluated at the $m$-host ($m < n$) disease-free equilibrium, $p^\dagger$ |

The mnemonic for the equilibrium notation is that line segments are used for endemic equilibria ($*$, '), daggers are used for the disease-free equilibria (‡, †), and more lines are used for the equilibria with more host species ($*$, ‡).

### S1 Computing sensitivities for the 1-habitat model

#### S1.1 Additional details about 1-habitat model

Subsections S1.1.1, S1.1.2, and S1.1.3 present the different forms of the 1-habitat model that are analyzed. Model variables and parameters are summarized in Table S2. Model assumptions are discussed in Section S1.1.4. The Jacobian used in the sensitivity calculations for focal host infection prevalence and infected density is defined in Section S1.1.3 and the mathematical properties of the Jacobian are discussed in Section S1.1.5.

##### S1.1.1 SIR form of the 1-habitat model

Recall from the main text that the SIR-form of the 1-habitat model describes the dynamics of the densities of susceptible ($S_i$), infected ($I_i$), and recovered ($R_i$) individuals, respectively, in each host population ($i \leq n$) and the density of infectious propagules in the environment ($P$). For reference, the SIR-form of the 1-habitat model is

$$
\begin{aligned}
\frac{dS_i}{dt} &= \underbrace{F_i(\cdot)}_{\text{reproduction}} - \underbrace{\beta_i S_i P}_{\text{infection}} - \underbrace{m_i S_i}_{\text{mortality}} + \underbrace{\gamma_i R_i}_{\text{loss of immunity}} \\
\frac{dI_i}{dt} &= \underbrace{\beta_i S_i P}_{\text{infection}} - \underbrace{(\mu_i + m_i) I_i}_{\text{mortality}} - \underbrace{\nu_i I_i}_{\text{recovery}} \\
\frac{dR_i}{dt} &= \underbrace{\nu_i I_i}_{\text{recovery}} - \underbrace{(\xi_i + m_i) R_i}_{\text{mortality}} - \underbrace{\gamma_i R_i}_{\text{loss of immunity}} \\
\frac{dP}{dt} &= \underbrace{\sum_{j=1}^{n} \chi_j I_j}_{\text{propagule shedding}} - \underbrace{\sum_{j=1}^{n} (u_{S_j} S_j + u_{I_j} I_j + u_{R_j} R_j) P}_{\text{propagule uptake}} - \underbrace{\delta P}_{\text{degradation}}
\end{aligned}
\tag{S1}
$$

The notation is summarized in Table S2.

##### S1.1.2 NYZ form of the 1-habitat model

The sensitivity calculations for focal host infection prevalence and infected density are easier to interpret biologically if the 1-habitat model is written in terms of the total density for each host ($N_i$) and the proportions of susceptible, infected and recovered individuals in each host population ($X_i$, $Y_i$, and $Z_i$, respectively). This form of the model was studied in detail in Cortez and Duffy (2021) and Cortez (2021). The NYZ form of the 1-habitat model is

Table S2: Variables, parameters, and important quantities for the 1-habitat model (S1)

| **Variable** | **Description** |
|---|---|
| $S_i$ | susceptible density for host $i$ |
| $I_i$ | infected density for host $i$ |
| $R_i$ | recovered density for host $i$ |
| $N_i = S_i + I_i + R_i$ | total density for host $i$ |
| $X_i = S_i/N_i$ | proportion of susceptible individuals for host $i$ |
| $Y_i = I_i/N_i$ | proportion of infected individuals for host $i$ |
| $Z_i = R_i/N_i$ | proportion of recovered individuals for host $i$ |
| $P$ | density of infectious propagules in environment |
| **Parameter** | **Description** |
| $F_i$ | reproduction rate of host $i$ |
| $f_{V_i}$ | per capita reproduction rate of class $V_i$ ($V \in \{S, I, R\}$) |
| $\beta_i = \rho_i u_{S_i}$ | transmission coefficient for host $i$ |
| $\rho_i$ | per infectious propagule probability of infection |
| $m_i$ | mortality rate due to non-disease related factors |
| $\mu_i$ | disease-induced mortality rate |
| $\xi_i$ | rate of increased mortality due to prior infection |
| $\nu_i$ | recovery rate |
| $\gamma_i$ | rate of loss of immunity |
| $\chi_i$ | shedding rate of host $i$ |
| $u_{V_i}$ | uptake rate for individuals in class $V_i$ ($V \in \{S, I, R\}$) |
| $\delta$ | degradation rate of infection propagules |
| $\Gamma_i$ | $= \nu_i/(m_i + \xi_i + \gamma_i)$, ratio of $I_i$ to $R_i$ at the endemic equilibrium |
| **Quantity** | **Description** |
| $p^\ddagger$ | $n$-species disease-free community equilibrium |
| $p^*$ | $n$-species endemic community equilibrium |
| $p^\dagger$ | $m$-species ($m < n$) disease-free subcommunity equilibrium |
| $p'$ | $m$-species ($m < n$) endemic subcommunity equilibrium |
| $U$ | $= \sum_i (u_{S_i} S_i + u_{I_i} I_i + u_{R_i} R_i)$<br>total per propagule rate of uptake of infectious propagules |
| $\varphi_i$ | per capita net production of infectious propagules by host $i$, determines if host $i$ is a source ($\varphi_i > 0$) or sink ($\varphi_i > 0$) of infectious propagules |
| $\mathcal{R}_{0,i}$ | pathogen basic reproduction number for host $i$ |
| $\mathcal{R}_{0,i}(\infty)$ | $= \beta_i \chi_i / u_{S_i}(m_i + \mu_i + \gamma_i)$, competence of host $i$<br>basic reproduction number for host $i$ in the limit of infinite host density |
| $\mathcal{R}_0^C$ | pathogen community reproduction number |

Double daggers (‡) denote values evaluated at the $n$-species disease-free community equilibrium ($p^\ddagger$), asterisks ($*$) denote values evaluated at the $n$-species endemic community equilibrium ($p^*$), daggers (†) denote values evaluated at the $m$-species disease-free community equilibrium ($p^\dagger$), and primes ($'$) denote values evaluated at the $m$-species endemic community equilibrium ($p'$). The mnemonic for number of species is that more lines are needed to create the symbols denoting equilibria for communities with more species (‡, *).

$$
\begin{aligned}
\frac{dN_i}{dt} &= \overbrace{F_i(\cdot)}^{\text{reproduction}} - \overbrace{m_i N_i}^{\text{natural mortality}} - \overbrace{(\mu_i Y_i N_i + \xi_i Z_i N_i)}^{\text{disease-related mortality}} \\
\frac{dY_i}{dt} &= \overbrace{\beta_i(1 - Y_i - Z_i)P_i}^{\text{infection}} - \overbrace{(\mu_i + m_i)Y_i}^{\text{mortality}} - \overbrace{\nu_i Y_i}^{\text{recovery}} - \frac{Y_i}{N_i}\frac{dN_i}{dt} \\
\frac{dZ_i}{dt} &= \overbrace{\nu_i Y_i}^{\text{recovery}} - \overbrace{(\xi_i + m_i)Z_i}^{\text{mortality}} \overbrace{-\gamma_i Z_i}^{\text{loss of immunity}} - \frac{Z_i}{N_i}\frac{dN_i}{dt} \\
\frac{dP}{dt} &= \underbrace{\sum_{j=1}^{n} \chi_j Y_j N_j}_{\text{propagule shedding}} - \underbrace{\sum_{j=1}^{n} (u_{S_j} X_j + u_{I_j} Y_j + u_{R_j} Z_j) N_j P_i}_{\text{propagule uptake}} - \underbrace{\delta_i P_i}_{\text{degradation}}
\end{aligned}
\tag{S2}
$$

The notation is summarized in Table S2. In the $dN_i/dt$ equation, total host density of host $i$ increases due to reproduction ($F_i$) and decreases due to mortality from non-disease and disease sources ($m_i N_i$, $\mu_i Y_i N_i$, and $\xi_i Z_i N_i$). In the $dY_i/dt$ equation, host $i$ infection prevalence increases due to new infections ($\beta_i X_i P$) and decreases due to mortality ($\mu_i Y_i + m_i Y_i$) and recovery ($\nu_i Y_i$). The last term in the $dY_i/dt$ equation accounts for changes in infection prevalence due to changes in the densities of susceptible and recovered individuals; it arises from the quotient rule of calculus, $dY_i/dt = (d/dt)(I_i/N_i) = [N_i dI_i/dt - I_i dN_i/dt]N_i^2 = (1/N_i)(dI_i/dt) - (Y_i/N_i)(dN_i/dt)$. In the $dZ_i/dt$ equation, the proportion of recovered individuals of host $i$ increases due to recovery ($\nu_i Y_i$), decreases due to mortality ($m_i Z_i + \xi_i Z_i$) and waning immunity ($\gamma_i Z_i$), and changes due to changes in the densities of the susceptible and infected individuals (last term, which arises due to the quotient rule from calculus). In the $dP/dt$ equation, infectious propagule density increases due to shedding by infected individuals (first sum) and decreases due to uptake of infectious propagules by all individuals of all host species (second sum) and degradation ($\delta P$). The total per propagule uptake rate of infectious propagules is denoted $U = \sum_{j=1}^{n}(u_{S_j} X_j + u_{I_j} Y_j + u_{R_j} Z_j)N_j$.

**Jacobian**: The Jacobian of the 1-habitat NYZ model (S2) is not used in the analysis in this study, but I present it here because its interpretation is necessary when interpreting later matrices. When evaluated at an endemic equilibrium the Jacobian of the 1-habitat NYZ model (S2) has the structure,

$$
\begin{bmatrix}
\mathbf{A} & \mathbf{B} & \mathbf{C} & \vec{0}_n \\
-\mathbf{D}_{Y/N}\mathbf{A} & -\mathbf{D}_{Y/N}\mathbf{B} - \mathbf{D}_{\beta P + m + \mu + \nu} & -\mathbf{D}_{Y/N}\mathbf{C} - \mathbf{D}_{\beta P} & \overrightarrow{\beta X} \\
-\mathbf{D}_{Z/N}\mathbf{A} & -\mathbf{D}_{Z/N}\mathbf{B} + \mathbf{D}_{\nu} & -\mathbf{D}_{Z/N}\mathbf{C} - \mathbf{D}_{\xi + m + \gamma} & \vec{0}_n \\
\vec{E}_N & \vec{E}_Y & \vec{E}_Z & -U - \delta
\end{bmatrix}.
\tag{S3}
$$

where the order of the rows and columns is $N_1, ..., N_n, Y_1, ..., Y_n, Z_1, ..., Z_n, P$. The mathematical properties of the Jacobian entries are described in detail in Section S1.1.5. Here, I provide a brief biological interpretation for each term.

The entries of $\mathbf{A}$ define the direct competitive effects of each host species on the growth rates of all host species; the diagonal entries, $\mathbf{A}_{ii}$ define the intraspecific competitive effects and the off-diagonal entries, $\mathbf{A}_{ij}$ $(i \neq j)$, define the interspecific competitive effects. The entries of $\mathbf{B}$ define how the growth rate of each host species is affected by disease-induced mortality of infected individuals (diagonal entries) and competition with infected individuals (all entries). The entries of $\mathbf{C}$ define how the growth rate of each host species is affected by disease-induced mortality of recovered individuals (diagonal entries) and competition with recovered individuals (all entries). The matrices of the form $\mathbf{D}_{Y/N}\mathbf{A}$, $\mathbf{D}_{Z/N}\mathbf{A}$, $\mathbf{D}_{Y/N}\mathbf{B}$, etc arise because of the terms $(Y_i/N_i)(dN_i/dt)$ and $(Z_i/N_i)(dN_i/dt)$ in the $dY_i/dt$ and $dZ_i/dt$ equations. They define how the growth rates of infection prevalence increase and decrease when the fractions of susceptible individuals decrease and increase, respectively. The diagonal matrices $-\mathbf{D}_{\beta P+m+\mu+\nu}$ and $-\mathbf{D}_{\beta P}$ account for how infection prevalence growth rates decrease due to reduced susceptible density, mortality, and recovery. The vector $\overrightarrow{\beta X}$ defines how infection prevalence growth rates increase with increased infectious propagule density. The diagonal matrices $\mathbf{D}_{\nu}$ and $-\mathbf{D}_{\xi+m+\gamma}$ define how the growth rates of the fraction of recovered individuals increase due to recovery and decrease due to mortality and loss of immunity. In the bottom row, $\vec{E}_N$, $\vec{E}_Y$, and $\vec{E}_Z$ define how the infectious propagule growth rate changes with increases in the density, infection prevalence, and proportion of recovered individuals of each host population. The entry $-U-\delta$ defines how the infectious propagule growth rate decreases due to uptake and degradation.

##### S1.1.3 Reduced NY form of the 1-habitat model

The calculations of the sensitivities at equilibrium are simplified by analyzing a reduced version of model (S2). The reduced form is derived by setting $dZ_i/dt = 0$, solving for $Z_i^* = \nu_i Y_i^*/(\xi_i + m_i + \gamma_i) = \Gamma_i Y_i^*$, and substituting into the other equations to yield

$$\begin{aligned}
\frac{dN_i}{dt} &= \overbrace{F_i(\cdot)}^{\text{reproduction}} - \overbrace{m_i N_i}^{\text{natural mortality}} - \overbrace{(\mu_i Y_i N_i + \xi_i \Gamma_i Y_i N_i)}^{\text{disease-related mortality}} \\
\frac{dY_i}{dt} &= \overbrace{\beta_i(1 - Y_i - \Gamma_i Y_i)P}^{\text{infection}} - \overbrace{(\mu_i + m_i)Y_i}^{\text{mortality}} - \overbrace{\nu_i Y_i}^{\text{recovery}} - \frac{Y_i}{N_i}\frac{dN_i}{dt} \\
\frac{dP}{dt} &= \underbrace{\sum_j \chi_j Y_j N_j}_{\text{propagule shedding}} - \underbrace{\sum_j (u_{S_j} X_j + u_{I_j} Y_j + u_{R_j} \Gamma_j Y_j) N_j P}_{\text{propagule uptake}} - \underbrace{\delta P}_{\text{degradation}}
\end{aligned} \tag{S4}$$

where $\Gamma_i = \nu_i/(\xi_i + m_i + \gamma_i)$. Note that any equilibrium point of the 1-habitat NYZ model (S2) is also an equilibrium of the 1-habitat NY model (S4). Consequently, the sensitivities of host 1 infection prevalence $(Y_1^*)$ and infected density $(I_1^*)$ at equilibrium can be computed using either model. I compute the sensitivities using the NY form of the model because that removes algebraic steps when computing each sensitivity.

**Jacobian**: When evaluated at an endemic equilibrium the Jacobian of the 1-habitat NY model (S4) has the structure,

$$\mathbf{J} = \begin{bmatrix} \mathbf{A} & \mathbf{B} + \mathbf{C}\mathbf{D}_\Gamma & \vec{0}_n \\ -\mathbf{D}_{Y/N}\mathbf{A} & -\mathbf{D}_{\beta P/Y} - \mathbf{D}_{Y/N}(\mathbf{B} + \mathbf{C}\mathbf{D}_\Gamma) & \overrightarrow{\beta X} \\ \vec{E}_N & \vec{E}_Y + \vec{E}_Z\mathbf{D}_\Gamma & -U - \delta \end{bmatrix} \tag{S5}$$

where matrix $\mathbf{D}_{\beta P/Y}$ follows from Remark S1.1. The biological interpretation of the terms is the same as the Jacobian (S3) of the 1-habitat NYZ model (S2) in Section S1.1. The mathematical properties of the Jacobian entries are described in detail in Section S1.1.5.

##### S1.1.4 Additional details about model assumptions

Additional details about specific components of the model and assumptions are given below. Some assumptions are defined by quantities derived from the Jacobian in equation (S5); details about those assumptions are provided in Section S1.1.5.

**Pathogen transmission and relationship between transmission and uptake parameters**: I assume susceptible individuals become infected when they encounter infectious propagules in their shared environment. In addition, the models assume infectious propagules are well-mixed throughout the habitat and do not differ based on which species shed them.

Encounter rates between susceptible individuals and infectious propagules are modeled using mass action kinetics, which means the contacts rates are proportional to both susceptible densities and infectious propagule density ($\beta_i S_i P$). The transmission coefficients are composite parameters, $\beta_i = p_i u_{S_i}$, that account for the rate at which susceptible individuals of host $i$ encounter and take up infectious propagules in their habitat ($u_{S_i}$) and the per propagule probability of infection ($p_i$). When computing the sensitivities, $\beta_i$ is varied independently of $u_{S_i}$ by varying $p_i$ and $u_{S_i}$ is varied independently of $\beta_i$ by compensating for changes in $u_{S_i}$ with changes in $p_i$.

**Reproduction rates**: I assume the reproduction rate for each host population can be written as

$$F_i(\cdot) = S_i f_{S_i}(\cdot) + I_i f_{I_i}(\cdot) + R_i f_{R_i}(\cdot) \tag{S6}$$

where $(\cdot)$ is shorthand for $(S_1, ..., S_n, I_1, ..., I_n, R_1, ..., R_n)$ and $f_{S_i}$, $f_{I_i}$, and $f_{R_i}$ are the per capita reproductive outputs of susceptible, infected, and recovered individuals, respectively. I assume the functions $f_{S_i}$, $f_{I_i}$, and $f_{R_i}$ are non-increasing functions of their arguments due to intraspecific and interspecific host competition, i.e., $\frac{\partial F_{V_i}}{\partial W_j} \leq 0$ for $V, W \in \{S, I, R\}$ and all $i, j$. These assumptions result in matrix $\mathbf{A}$ of the Jacobian having negative entries; see Section S1.1.5. The assumptions are satisfied by the Lotka-Volterra competition functions used in the simulations (see equation (3) of the main text).

The form of model (S1) and the above assumptions about the reproduction rates imply that intraspecific density dependence and interspecific interactions only affect the reproduction rates of the host species. This simplifies the mathematical analysis considerably because if mortality is density dependent or affected by interspecific interactions, then many more

Jacobian entries will depend on the host densities and its determinants will yield much more complex equations. Also, while I interpret and discuss my results in terms of competitive interactions between the host species, my results also apply to mutualistic and contramensalistic interactions, provided the interactions only affect the reproduction rates of the host species. However, this means that my results do not necessarily apply to host species with predator-prey or other exploitative interactions that affect the mortality rates of species.

**Relationship between recovered and susceptible individuals**: I assume that recovered individuals are identical to susceptible individuals with the possible exception that they experience increased mortality (at rate $\xi_i$) due to prior exposure to the pathogen. In particular, I assume that susceptible and recovered individuals have identical reproduction rates ($f_{S_i} = f_{R_i}$), competitive abilities ($\partial f_{V_j}/\partial S_i = \partial f_{V_j}/\partial R_i$ for all $i, j$ and $V \in \{S, I, R\}$), and uptake rates ($u_{S_i} = u_{R_i}$).

**Coexistence equilibria, notation, and stability**: The equilibrium-based analysis in this study focuses on four different kinds of equilibria: (i) the endemic equilibrium where the pathogen and $n$ host species coexist ($p^*$), (ii) the disease-free equilibrium where the $n$ host species coexist in the absence of the pathogen ($p^\ddagger$), (iii) endemic subcommunity equilibria where the pathogen and $m$ ($m < n$) host species coexist ($p'$), and (iv) disease-free subcommunity equilibria where $m$ host species coexist in the absence of the pathogen ($p^\dagger$).

Throughout, asterisks ($*$) denote quantities evaluated at the endemic community equilibrium ($p^*$), double daggers (‡) denote quantities evaluated at the disease-free community equilibrium ($p^\ddagger$), primes ($'$) denote quantities evaluated at the endemic subcommunity equilibrium ($p'$), and daggers (†) denote quantities evaluated at the disease-free subcommunity equilibrium ($p^\dagger$). Here, the mnemonics are (i) asterisks denote endemic communities with more species because more line segments are needed to draw an asterisk than a prime and (ii) double daggers denote disease-free communities with more species because they have more lines than daggers.

Throughout, I assume all of the equilibria are stable when necessary. First, for the sensitivities of the disease metrics to the epidemiological parameters of the alternative host species (Sections S1.4.2 and S1.5.3), I assume the endemic equilibrium ($p^*$) and disease-free equilibrium ($p^\ddagger$) are stable. These assumptions assure that the predictions from the sensitivities are biologically relevant; if the equilibria are unstable, the system is unlikely to be near the equilibria, which means predictions made about those equilibria are unlikely to be biologically informative. Second, for the sensitivities of the disease metrics to the densities of the alternative host species (Sections S1.4.1 and S1.5.2), I assume the endemic equilibrium ($p^*$) and disease-free equilibrium ($p^\ddagger$) are stable and I also assume the $n-1$ host subcommunity endemic equilibrium ($p'$) and $n-1$ host disease-free equilibrium ($p^\dagger$) are stable. When these conditions are not met, the sensitivity results are unlikely to be informative about the effects of the addition or removal of host $n$. As examples, instability of an $n$-host equilibrium means the systems is unlikely to be near the equilibrium and instability of any of the equilibria can mean that the addition or removal of host $n$ causes other host species to go extinct. Third, for the sensitivities of the disease metrics to the pathogen transmission mode (Sections S1.4.3 and S1.5.4), I assume the endemic equilibrium ($p^*$) and disease-free equilibrium ($p^\ddagger$) are stable and I also assume the $m$ host subcommunity endemic equilibrium

($p'$) and $m$ host disease-free equilibrium ($p^\dagger$) are stable. When these conditions are not met, the sensitivity results are unlikely to be informative about how the transmission model influences the effects of the addition or removal of host species $m+1, ..., n$; the reasoning is the same as for the sensitivities to alternative host density. In total, this means that all of my results assume the $n$ species can stable coexist in the presence and absence of the pathogen and specific subcommunties of $m$ host species can coexist in the presence and absence of the pathogen.

**Negative density dependence at equilibrium**: I assume all host species experience negative density dependence at the disease-free and endemic equilibria, i.e., increased density of host $i$ leads to a decrease in its growth rate. Mathematically, species $i$ experiences negative density dependence at equilibrium if $(\partial/\partial N_i)(dN_i/dt) = \frac{1}{N_i}\frac{dN_i}{dt} + \frac{\partial}{\partial N_i}(\frac{1}{N_i}\frac{dN_i}{dt}) = \frac{\partial}{\partial N_i}(\frac{1}{N_i}\frac{dN_i}{dt}) = \mathbf{A}_{ii} < 0$, where all terms are evaluated at equilibrium and $\mathbf{A}$ is the submatrix of the Jacobian, $\mathbf{J}$. As shown in Section S1.1.5, $\mathbf{A}_{ii}$ is always negative under the assumptions on the host growth rates ($F_i$).

**Assumptions about Jacobian of 1-habitat NY model and biological interpretation**: Following Cortez and Duffy (2021) and Cortez (2021), I make assumptions about the model in order to simplify the analysis and focus the analysis on biological scenarios that are more likely to be observed in natural systems. Below, I list the biological assumptions and their corresponding mathematical conditions; some assumptions from above are repeated so that all of the mathematical conditions are listed in the same place. The mathematical conditions are defined by the Jacobian of the 1-habitat NY model (S4) and quantities derived from the Jacobian; see Section S1.1.5 for additional details. I note that many of the conditions are guaranteed to hold if intraspecific competition is sufficiently greater than interspecific competition for all host species.

1. Recovered and susceptible individuals have equal reproductive output, competitive abilities, and uptake rates: $f_{S_i} = f_{R_i}$, $\partial f_{V_i}/\partial S_j = \partial f_{V_i}/\partial R_j$, and $u_{S_i} = u_{R_i}$ for all $i, j$ and $V \in \{S, I, R\}$.

2. Susceptible individuals have greater or equal per capita reproductive output than infected individuals: $f_{S_i} \geq f_{I_i}$ for all $i$.

3. Infected individuals are stronger, equal, or moderately weaker (but not much weaker) competitors than susceptible individuals: $\partial f_{V_i}/\partial I_j$ is larger, equal to, or moderately smaller than $\partial f_{V_i}/\partial S_j$ for all $i, j$ and $V \in \{S, I, R\}$. I do not consider systems where infected individuals are much weaker competitors than susceptible individuals because it can lead to scenarios where the disease allows the host to reach densities higher than its carrying capacity. This is unlikely to arise in natural systems.

4. All $n$ host species coexist with the pathogen at a stable endemic equilibrium of the model ($p^*$) and all $n$ host species stably coexist in the absence of the pathogen at a disease-free equilibrium ($p^\ddagger$): $\text{sign}(|\mathbf{J}|)|_{p^*} = (-1)^{2n+1}$ and $\text{sign}(|\mathbf{A}|)|_{p^\ddagger} = (-1)^n$.

5. All host species experience negative density dependence at equilibrium:
$\mathbf{A}_{ii} = (\partial/\partial N_i)(dN_i/dt)|_{p^*} < 0$.

6. While their direct interspecific competitive effects are zero or negative ($\mathbf{A}_{ji} \leq 0$), the total interspecific effects between host species ($\alpha_{ij}^{total}$) can be negative or positive: the total competitive effect of species $i$ on species $j$ is negative ($\alpha_{ij}^{total} < 0$) when $\text{sign}(|\mathbf{A}^{[\not{i},\not{j}]}|) = (-1)^{n+i+j}$ and the total competitive effect of species $i$ on species $j$ is positive ($\alpha_{ij}^{total} > 0$) when $\text{sign}(|\mathbf{A}^{[\not{i},\not{j}]}|) = (-1)^{n+i+j+1}$. See Remark S1.3 in Section S1.1.5 for details.

7. The competitive effects between all host species, all subcommunities of $n-1$ host species, and all subcommunities of $n-2$ host species are consistent with stability: $\text{sign}(|\mathbf{A}|) = (-1)^n$, $\text{sign}(|\mathbf{A}^{[\not{i},\not{i}]}|) = (-1)^{n-1}$, and $\text{sign}(|\mathbf{A}^{[\not{i}\not{j},\not{i}\not{j}]}|) = (-1)^{n-2}$ for all $i, j$. See Remark S1.2 in Section S1.1.5 for details.

8. Increased prevalence in a host species at equilibrium has a negative total effect on its growth rate: $\text{sign}(|\mathbf{A}_{Y_j \to N_j}) = (-1)^{n-1}$ for all $i$. See Remark S1.5 in Section S1.1.5 for details.

9. Increased prevalence in host $j$ at equilibrium has a negative and positive total effect on the growth rate of host $i$ when infected individuals of host $i$ are stronger and weaker competitors, respectively, then susceptible individuals: $\text{sign}(|\mathbf{A}_{Y_j \to N_j}|)$ equals $(-1)^{n-1}$ or $(-1)^n$ when infected individuals of host $j$ are weaker or stronger competitors, respectively, than susceptible individuals. See Remark S1.5 in Section S1.1.5 for details.

##### S1.1.5 Properties and derived quantities of the Jacobian of the 1-habitat NY model

The following shows properties of the Jacobian entries and quantities derived from the Jacobian. Much of this material is taken from Cortez (2021), with minor adjustments made to account for the fact that my calculations use the 1-habitat NY model (S4) instead of the 1-habitat NYZ model (S2). Note that $\text{sign}(|\mathbf{J}|) = (-1)^{2n+1}$ when evaluated at $p^*$ because I assume the $n$-host endemic equilibrium is stable.

**Properties of A**: All entries of $\mathbf{A}$ are negative because of the assumptions about the reproduction functions $F_i$. To see this for entry $\mathbf{A}_{ii}$, differentiate to get

$$\begin{aligned}
\mathbf{A}_{ii} &= \frac{\partial}{\partial N_i}\left[\frac{dN_i}{dt}\right]\Bigg|_{p^*} = \frac{\partial}{\partial N_i}\Bigg[F_i - m_i N_i - \mu_i Y_i N_i - \xi_i Z_i N_i\Bigg]\Bigg|_{p^*} \\
&= \frac{\partial}{\partial N_i}\Bigg[N_i X_i f_{S_i} + N_i Y_i f_{I_i} + N_i Z_i f_{R_i} - m_i N_i - \mu_i Y_i N_i - \xi_i Z_i N_i\Bigg]\Bigg|_{p^*} \\
&= X_i f_{S_i} + Y_i f_{I_i} + Z_i f_{R_i} - m_i - \mu_i Y_i - \xi_i Z_i + N_i \frac{\partial}{\partial N_i}\Bigg[X_i f_{S_i} + Y_i f_{I_i} + Z_i f_{R_i}\Bigg]\Bigg|_{p^*} \\
&= N_i \frac{\partial}{\partial N_i}\Bigg[X_i f_{S_i} + Y_i f_{I_i} + Z_i f_{R_i}\Bigg]\Bigg|_{p^*} \\
&= N_i \frac{\partial}{\partial N_i}\left(\frac{F_i}{N_i}\right)\Bigg|_{p^*}.
\end{aligned} \tag{S7}$$

The third to last equality follows from the equilibrium condition $0 = dN_i/dt|_{p^*} = F_i(\cdot)|_{p^*} - m_i N_i^* - \mu_i Y_i^* N_i^* - \xi_i Z_i^* N_i^*$. To show that $\mathbf{A}_{ii}$ is negative, apply the chain rule to get

$$\begin{aligned} \mathbf{A}_{ii} =& N_i \frac{\partial}{\partial N_i}\left(\frac{F_i}{N_i}\right)\Big|_{p^*} && \text{(S8)} \\ =& N_i \frac{\partial}{\partial N_i}\left[X_i f_{S_i} + Y_i f_{I_i} + Z_i f_{R_i}\right]\Big|_{p^*} && \text{(S9)} \\ =& N_i X_i \left[X_i \frac{\partial f_{S_i}}{\partial S_i} + Y_i \frac{\partial f_{S_i}}{\partial I_i} + Z_i \frac{\partial f_{S_i}}{\partial R_i}\right]\Big|_{p^*} + N_i Y_i \left[X_i \frac{\partial f_{I_i}}{\partial S_i} + Y_i \frac{\partial f_{I_i}}{\partial I_i} + Z_i \frac{\partial f_{I_i}}{\partial R_i}\right]\Big|_{p^*} \\ & + N_i Z_i \left[X_i \frac{\partial f_{R_i}}{\partial S_i} + Y_i \frac{\partial f_{R_i}}{\partial I_i} + Z_i \frac{\partial f_{R_i}}{\partial R_i}\right]\Big|_{p^*}. && \text{(S10)} \end{aligned}$$

The last line is negative because of the assumption that $f_{V_i}$ ($V \in \{S, I, R\}$) is a decreasing function of $S_j$, $I_j$, and $R_j$ for all $i, j$.

Entry $\mathbf{A}_{ij}$ ($i \neq j$) simplifies to

$$\mathbf{A}_{ij} = \frac{\partial}{\partial N_j}\left[\frac{dN_i}{dt}\right]\Big|_{p^*} = \frac{\partial}{\partial N_j}\left[F_i - m_i N_i - \mu_i Y_i N_i - \xi_i Z_i N_i\right]\Big|_{p^*} = \frac{\partial}{\partial N_j}\left[F_i\right]\Big|_{p^*}. \quad \text{(S11)}$$

The same calculation as above shows that each entry is negative.

**Properties of C**: Applying the chain rule shows that the entries of **C** are

$$\mathbf{C}_{ii} = N_i^*[f_{R_i} - f_{S_i} - \xi_i] + X_i^* N_i^{*2}\left[\frac{\partial f_{S_i}}{\partial R_i} - \frac{\partial f_{S_i}}{\partial S_i}\right] + Y_i^* N_i^{*2}\left[\frac{\partial f_{I_i}}{\partial R_i} - \frac{\partial f_{I_i}}{\partial S_i}\right] + Z_i^* N_i^{*2}\left[\frac{\partial f_{R_i}}{\partial R_i} - \frac{\partial f_{R_i}}{\partial S_i}\right] \quad \text{(S12)}$$

$$\mathbf{C}_{ij} = X_i^* N_i^* N_j^*\left[\frac{\partial f_{S_i}}{\partial R_j} - \frac{\partial f_{S_i}}{\partial S_j}\right] + Y_i^* N_i^* N_j^*\left[\frac{\partial f_{I_i}}{\partial R_j} - \frac{\partial f_{I_i}}{\partial S_j}\right] + Z_i^* N_i^* N_j^*\left[\frac{\partial f_{R_i}}{\partial R_j} - \frac{\partial f_{R_i}}{\partial S_j}\right] \quad \text{(S13)}$$

where $i \neq j$ in the $\mathbf{C}_{ij}$ equation. Under the assumption that susceptible and recovered individuals have equal reproduction rates ($f_{S_i} = f_{R_i}$) and equal competitive abilities ($\partial f_{V_i}/\partial S_j = \partial f_{V_i}/\partial R_j$ for $V \in \{S, I, R\}$), the entries of **C** simplify to $\mathbf{C}_{ii} = -\xi_i N_i^*$ and $\mathbf{C}_{ij} = 0$ for $i \neq j$.

**Properties of B and B + CD$_\Gamma$**: The entries of **B** have the form,

$$\begin{aligned} \mathbf{B}_{ii} =& N_i^*[f_{I_i} - f_{S_i} - \mu_i] + N_i^{*2} X_i^*\left[\frac{\partial f_{S_i}}{\partial I_i} - \frac{\partial f_{S_i}}{\partial S_i}\right] \\ & + N_i^{*2} Y_i^*\left[\frac{\partial f_{I_i}}{\partial I_i} - \frac{\partial f_{I_i}}{\partial S_i}\right] + N_i^{*2} Z_i^*\left[\frac{\partial f_{R_i}}{\partial I_i} - \frac{\partial f_{R_i}}{\partial S_i}\right] && \text{(S14)} \\ \mathbf{B}_{ij} =& X_i^* N_i^* N_j^*\left[\frac{\partial f_{S_i}}{\partial I_j} - \frac{\partial f_{S_i}}{\partial S_j}\right] + Y_i^* N_i^* N_j^*\left[\frac{\partial f_{I_i}}{\partial I_j} - \frac{\partial f_{I_i}}{\partial S_j}\right] + Z_i^* N_i^* N_j^*\left[\frac{\partial f_{R_i}}{\partial I_j} - \frac{\partial f_{R_i}}{\partial S_j}\right] && \text{(S15)} \end{aligned}$$

where $i \neq j$ in the $\mathbf{B}_{ij}$ equation. The sign of $f_{I_i} - f_{S_i}$ is negative, zero and positive when infected individuals have reduced, equal, or greater reproductive output, respectively, relative to susceptible individuals. The sign of $\partial f_{V_i}/\partial I_j - \partial f_{V_i}/\partial S_j$ is positive, zero, and negative when infected individuals of host $i$ are weaker, equal, or stronger interspecific competitors than susceptible individuals, respectively.

For most empirical systems, I expect that (i) infected individuals have equal or reduced reproductive output compared to susceptible individuals ($f_{I_i} - f_{S_i} \leq 0$) and (ii) infected individuals can be weaker, equal or stronger competitors than susceptible individuals, but infected individuals are not much weaker competitors than susceptible individuals ($\partial f_{V_i}/\partial I_j$ is never much smaller in magnitude than $\partial f_{V_i}/\partial S_j$). Under these assumptions, $\mathbf{B}_{ii} + \Gamma_i \mathbf{C}_{ii}$ is negative unless infected individuals of host $i$ are sufficiently weaker intraspecific competitors than susceptible individuals. The off-diagonal entries of $\mathbf{B} + \mathbf{C}\mathbf{D}_\Gamma$ are positive, zero, and negative when infected individuals of host $j$ are weaker, equal or stronger interspecific competitors than susceptible individuals of host $j$ , respectively.

**Properties of $\vec{E}_N$, $\vec{E}_Y$, and $\vec{E}_Z$:** The three vectors have entries

$$\vec{E}_N[i] = \chi_i Y_i^* - (u_{S_i} X_i^* + u_{I_i} Y_i^* + u_{R_i} Z_i^*) P^* = Y_i^* \varphi_i \tag{S16}$$

$$\vec{E}_Y[i] = \chi_i N_i^* - (u_{I_i} - u_{S_i}) N_i^* P^* = N_i^* \varphi_i + N_i^* u_{S_i} P^*/Y_i^* \tag{S17}$$

$$\vec{E}_Z[i] = -(u_{R_i} - u_{S_i}) Z_i^* P^* = 0 \tag{S18}$$

where $\mathbf{E}_Z[i] = 0$ because I assume susceptible and recovered individuals have equal uptake rates. The quantity $\varphi_i$ defines if host $i$ is a source ($\varphi_i > 0$) or sink ($\varphi_i < 0$) of infectious propagules at the endemic equilibrium; see Section S1.3 for details.

**Remark S1.1.** *For any host species, the following holds at equilibrium*

$$\beta_i P^* + \mu_i + m_i + \nu_i + \frac{\beta_i P^* \nu_i}{\xi_i + m_i + \gamma_i} = \frac{\beta_i P^*}{Y_i^*} \tag{S19}$$

*Reasoning:* The equilibrium conditions $dN_i/dt = 0$ and $dY_i/dt = 0$ imply that $\beta_i P^* + \mu_i + m_i + \nu_i = \beta_i(1 - Z_i^*)P^*/Y_i^*$. Similarly, the equilibrium conditions $dN_i/dt = 0$ and $dR_i/dt = 0$ imply that $\nu_i/(\xi_i + m_i + \gamma_i) = Z_i^*/Y_i^*$. Substitution and a little algebra yields

$$\beta_i P^* + \mu_i + m_i + \nu_i + \frac{\beta_i P^* \nu_i}{\xi_i + m_i + \gamma_i} = \frac{\beta_i(1 - Z_i^*)P^*}{Y_i^*} + \frac{\beta_i P^* Z_i^*}{Y_i^*} = \frac{\beta_i P^*}{Y_i^*}. \tag{S20}$$

•

**Remark S1.2.** *I assume the competitive interactions between (i) all host species, (ii) all subsets of $n-1$ host species, and (iii) all subsets of $n-2$ host species are consistent with stability at equilibrium. Mathematically, I assume (i) sign($|\mathbf{A}|$) $= (-1)^n$, (ii) sign($|\mathbf{A}^{[\not{i},\not{i}]}|$) $= (-1)^{n-1}$, and (iii) sign($|\mathbf{A}^{[\not{i}\not{j},\not{i}\not{j}]}|$) $= (-1)^{n-2}$. These conditions may not hold if interspecific competition is sufficiently strong and asymmetric.*

*Reasoning:* Each submatrix determines the stability of the system when the disease levels in all species (i.e., $I_i$ and $P$) and the densities of zero, one, or two host species are frozen at their equilibrium values. I assume that the sign of the determinant of the Jacobian for each of those subsystems is consistent with stability. Because $\mathbf{A}_{ii} < 0$ for all $i$, this is guaranteed to hold if interspecific competition is sufficiently weak.

My assumptions avoid scenarios where the interspecific interactions between the host species are so strong that coexistence at the equilibrium $p^*$ is pathogen-mediated. For example, the assumption $\text{sign}(|\mathbf{A}|) = (-1)^n$ implies the $n$-host subsystem of the equilibrium is stable and the host species could stably coexist if infection levels were frozen at their equilibrium values. In comparison, if $\text{sign}(|\mathbf{A}|) \neq (-1)^n$, then the $n$-host subsystem of the equilibrium is unstable and the host species cannot stably coexist if infection levels were frozen at their equilibrium values. In this case, coexistence of the host species at the equilibrium $p^*$ requires the dynamics of the pathogen. •

**Remark S1.3.** *I interpret the quantity*

$$\alpha_{ij}^{total} = (-1)^{n+j+i} \frac{|\mathbf{A}^{[\not j,\not i]}|}{|\mathbf{J}|} \quad (i \neq j) \tag{S21}$$

*in terms of the total competitive effect of host species $j$ on host species $i$. The total competitive effect is positive when $\alpha_{ij}^{total} > 0$ and negative when $\alpha_{ij}^{total} < 0$.*

*Reasoning:* The determinant $|\mathbf{A}^{[\not j,\not i]}|$ $(i \neq j)$ can be written in terms of the sum of the direct and indirect effects (i.e., the total effect) of host $j$ on host $i$. In particular,

$$(-1)^{j+i-1}|\mathbf{A}^{[\not j,\not i]}|/|\mathbf{A}^{[\not i\,\not j,\not i\,\not j]}| = \mathbf{A}_{ij} + terms \tag{S22}$$

where $\mathbf{A}_{ij}$ is the direct competitive effect of host $j$ on host $i$ and *terms* is the indirect competitive effect. The indirect effect is normalized by $|\mathbf{A}^{[\not i\,\not j,\not i\,\not j]}|$, which represents the combined effects of all competitive interactions between all host species except species $i$ and $j$. Thus, *terms* is the indirect competitive effect of host $i$ on host $j$, mediated by the competitive interactions between all host species except species $i$ and $j$.

Under the assumption that the sign of $|\mathbf{A}^{[\not i\,\not j,\not i\,\not j]}|$ is $(-1)^{n-2}$ (see Remark S1.2), the sign of the total effect is given by

$$\text{sign}(|\mathbf{A}^{[\not j,\not i]}|) = (-1)^{j+i+n-1}\text{sign}(\mathbf{A}_{ij} + terms). \tag{S23}$$

If the total effect of species $i$ on $j$ has the same sign as the direct effect, then $\text{sign}(|\mathbf{A}^{[\not j,\not i]}|) = (-1)^{n+j+i}$ when the direct effect is negative and $\text{sign}(|\mathbf{A}^{[\not j,\not i]}|) = (-1)^{n+j+i+1}$ when the direct effect is positive. If the total effect of species $j$ on $i$ has the opposite sign as the direct effect, then $\text{sign}(|\mathbf{A}^{[\not j,\not i]}|) = (-1)^{n+j+i+1}$ when the direct effect is negative and $\text{sign}(|\mathbf{A}^{[\not j,\not i]}|) = (-1)^{n+j+i}$ when the direct effect is positive. Combined, this means that the sign of $\text{sign}(|\mathbf{A}^{[\not j,\not i]}|)$ is $(-1)^{n+j+i}$ when the total effect is negative and the sign of $\text{sign}(|\mathbf{A}^{[\not j,\not i]}|)$ is $(-1)^{n+j+i+1}$ when the total effect is positive. Using equation (S21), this means that $\alpha_{ij}^{total} < 0$ when the total effect of species $j$ on species $i$ is negative and $\alpha_{ij}^{total} > 0$ when the total effect of species $j$ on species $i$ is positive. •

**Proposition S1.4.** *For a stable system, the sign of $|\mathbf{A}|$ is consistent with stability if and only if the infectious propagules do not exhibit a hydra effect at equilibrium. That is, for a system where $(|\mathbf{J}|) < 0$, sign($|\mathbf{A}|$) $= (-1)^n$ iff $\partial P^*/\partial\delta < 0$.*

*Proof.* Elementary row operations yield

$$\frac{\partial P^*}{\partial \delta} = \left(-\frac{\partial}{\partial \delta}\frac{dP}{dt}\right)\frac{(-1)^{2n+1+2n+1}}{|\mathbf{J}|}\left|\mathbf{J}^{[\cancel{2n+1},\cancel{2n+1}]}\right| \tag{S24}$$

$$= \frac{P^*}{|\mathbf{J}|}\begin{vmatrix} \mathbf{A} & \mathbf{B}+\mathbf{D}_\Gamma\mathbf{C} \\ -\mathbf{D}_{Y/N}\mathbf{A} & -\mathbf{D}_{Y/N}\mathbf{B}-\mathbf{D}_{\beta P/Y}\end{vmatrix} = \frac{P^*}{|\mathbf{J}|}\begin{vmatrix} \mathbf{A} & \mathbf{B}+\mathbf{D}_\Gamma\mathbf{C} \\ \mathbf{0} & -\mathbf{D}_{\beta P/Y}\end{vmatrix} \tag{S25}$$

$$= \frac{P^*}{|\mathbf{J}|}|\mathbf{A}|\prod_{j\leq n}\left(-\frac{\beta_j P^*}{Y_j^*}\right) \tag{S26}$$

Because $|\mathbf{J}| < 0$ is assumed and there are $n$ host species, sign($|\mathbf{A}|$) $= (-1)^n$ if and only if $(\partial P^*/\partial\delta) < 0$. □

**Remark S1.5.** *The sign of $(|\mathbf{A}_{I_j\to N_j}|)$ is assumed to be $(-1)^n$. If there is no interspecific competition, then $|\mathbf{A}_{I_j\to N_i}| = 0$ for $i \neq j$. If interspecific competition is sufficiently weaker than intraspecific competition (i.e., $\mathbf{A}_{jj}$ larger in magnitude than $\mathbf{A}_{ji}$ for $j \neq i$), then the sign of $|\mathbf{A}_{I_j\to N_i}|$ is $(-1)^{n-1}$ when infected individuals of host $j$ are weaker interspecific competitors than susceptible individuals and the sign is $(-1)^n$ when infected individuals of host $j$ are stronger interspecific competitors than susceptible individuals. The sign of $|\mathbf{A}_{I_j\to N_i}|$ for $i \neq j$ can differ if interspecific competition is sufficiently strong and asymmetric.*

*Reasoning:* The total effect of simultaneous changes in $Y_j^*$ and $Z_j^*$ on the growth rate of host $i$ is determined by the determinant $|\mathbf{A}_{I_j\to N_i}|$. Here, $\mathbf{A}_{I_j\to N_i}$ is matrix $\mathbf{A}$ where column $i$ has been replaced by column $j$ of $\mathbf{B}+\mathbf{C}\mathbf{D}_\Gamma$. Column $j$ of $\mathbf{B}+\mathbf{C}\mathbf{D}_\Gamma$ accounts for how the competitive effects of host $j$ on all host species change with a small increase in the frequencies of infected and recovered individuals in host $j$. The determinant $|\mathbf{A}_{I_j\to N_i}|$ accounts for how those competitive effects propagate through the host community and affect the growth rate of host $i$. Another way to see this is that $|\mathbf{A}_{I_j\to N_i}|$ can be written in terms of the total competitive effect of host $i$ on host $j$. Specifically, $|\mathbf{A}_{I_j\to N_i}|/|\mathbf{A}^{[\cancel{j},\cancel{j}]}| = \mathbf{B}_{ji} + \mathbf{C}_{ji}\Gamma_j + \textit{terms}$, where $\mathbf{B}_{ji} + \mathbf{C}_{ji}\Gamma_j$ is the sum of direct competitive effects of changes in the frequencies of infected and recovered individuals of host $i$ on the growth rate of host $j$ and *terms* is the indirect competitive effect. The indirect effect is normalized by $|\mathbf{A}^{[\cancel{j},\cancel{j}]}|$, which accounts for the competitive interactions between all host species except host $j$.

The sign of $(|\mathbf{A}_{I_j\to N_j}|)$ is assumed to be $(-1)^n$. The opposite sign can occur, but it only occurred in numerical simulations when infected individuals were very weak intraspecific competitors and the disease caused the density of species $j$ to increase above its carrying capacity.

If there is no interspecific competition, then $|\mathbf{A}_{I_j\to N_i}| = 0$ for $i \neq j$. If there is interspecific competition, then the sign of $|\mathbf{A}_{I_j\to N_i}|$ $(i \neq j)$ depends on the competitive abilities of infected individuals. If interspecific competition is sufficiently weaker than intraspecific competition (i.e., $\mathbf{A}_{jj}$ larger in magnitude than $\mathbf{A}_{ji}$ for $j \neq i$), then the sign of $|\mathbf{A}_{I_j\to N_i}|$ is $(-1)^{n-1}$ when infected individuals of host $j$ are weaker interspecific competitors than susceptible individuals and the sign is $(-1)^n$ when infected individuals of host $j$ are stronger interspecific competitors than susceptible individuals. This is because the leading order term of the determinant is $\mathbf{B}_{ji}\prod_{k\neq j}\mathbf{A}_{kk}$ and $\mathbf{B}_{ji}$ is positive and negative, respectively, when infected individuals of host $j$ are weaker and stronger interspecific competitors than susceptible individuals. In general, the sign of $|\mathbf{A}_{I_j\to N_i}|$ is consistent with the above when (i) the total competitive effect of host $j$ on host $i$ has the same sign as the direct effect and (ii) the determinant of the submatrix for the $\{N_k\}_{k\neq j}$ is consistent with stability. Conditions (i) and (ii) may not hold if interspecific competition is strong and asymmetric. •

**Remark S1.6.** *For any parameter $\sigma$, the sensitivities $\partial Y_1^*/\partial\sigma$ and $\partial I_1^*/\partial\sigma$ are more likely to differ in sign when (i) regulation of a host species by competition with host species 1 is sufficiently weaker than regulation by the disease and competition with other host species or (ii) host 1 is sink or small source and some subset of hosts $j>1$ are sufficiently large sources. Condition (i) is most likely to be satisfied when the host 1 is more strongly regulated by disease or interspecific competition than by intraspecific competition. Also, condition (i) is easier to satisfy in systems where infected individuals are equal or stronger competitors than susceptible individual and condition (ii) is easier to satisfy in systems where infected individuals have uptake rates that are larger than or equal to the uptake rates of susceptible individuals.*

*Reasoning:* I define the vectors $\vec{\omega}_n$ and $\vec{\omega}_{n-1}$ as

$$\vec{\omega}_n = \begin{pmatrix} -\sum_{j=1}^{n} X_k^*Y_k^*(\mathbf{B}_{1k}+\Gamma_1\mathbf{C}_{1k}) \\ \vdots \\ -\sum_{j=1}^{n} X_n^*Y_n^*(\mathbf{B}_{n,k}+\Gamma_n\mathbf{C}_{n,k}) \end{pmatrix}, \tag{S27}$$

$$\vec{\omega}_{n-1} = \begin{pmatrix} -\sum_{j=1}^{n-1} X_k^*Y_k^*(\mathbf{B}_{1k}+\Gamma_1\mathbf{C}_{1k}) \\ \vdots \\ -\sum_{j=1}^{n-1} X_n^*Y_n^*(\mathbf{B}_{n,k}+\Gamma_n\mathbf{C}_{n,k}), \end{pmatrix} \tag{S28}$$

and I define the quantities $\theta_n$ and $\theta_{n-1}$ as

$$\theta_n = (U^*+\delta)P^* - \sum_{k\leq n} X_k^*Y_k^*\vec{E}_Y[k], \tag{S29}$$

$$\theta_{n-1} = (U^*+\delta)P^* - \sum_{k\leq n-1} X_k^*Y_k^*\vec{E}_Y[k]. \tag{S30}$$

As shown in the following sections, the matrices used to compute the sensitivities $\partial Y_1^*/\partial\sigma$ and $\partial I_1^*/\partial\sigma$ only differ in one column. In particular, matrices used to compute the sensitivities of host 1 infection prevalence and host 1 infected density, respectively, have columns of the form

$$\begin{bmatrix} S_1^*\mathbf{A}_{\bullet 1} \\ S_1^*\mathbf{E}_N[1] \end{bmatrix} \text{ versus } \begin{bmatrix} \vec{\omega}_n \\ \theta_n \end{bmatrix} \text{ or } \begin{bmatrix} \vec{\omega}_{n+1} \\ \theta_{n+1} \end{bmatrix}. \tag{S31}$$

The sensitivities for host 1 infection prevalence and host 1 infected density can have opposite signs when the signs or magnitudes of the entries of the columns differ. This occurs under two general conditions.

First, the sensitivities for host 1 infection prevalence and host 1 infected density can have opposite signs if the entries of $\vec{\omega}_n$ or $\vec{\omega}_{n-1}$ are positive and large in magnitude relative to the negative entries of $\mathbf{A}_{\bullet 1}$. Biologically, this means regulation of each host species by competition with host 1 ($\mathbf{A}_{\bullet 1}$) is weaker than regulation by disease and competition with the other host species ($\vec{\omega}_n$ or $\vec{\omega}_{n-1}$). This is more likely to occur when (a) disease-induced mortality is higher ($\mu_i$ and $\xi_i$ are large), (b) infected individuals are stronger or equal competitors than susceptible individuals ($\partial f_{V_i}/\partial I_j - \partial f_{V_i}/\partial S_j$ is negative for $V \in \{S, I, R\}$), and (c) host 1 is a relatively weak intraspecific and interspecific competitor ($\mathbf{A}_{i1}$ is small in magnitude relative to $\mu_i$ and $\xi_i$). In our numerical simulations, differences in the signs of the determinants occurred most often when regulation of host 1 by intraspecific competition was weaker than regulation by disease and competition with the other host species.

Second, the sensitivities for host 1 infection prevalence and host 1 infected density can have opposite signs if $S_1^* \vec{E}_N[1]$ is negative or small in magnitude and $\theta_n$ or $\theta_{n-1}$ is positive and large in magnitude. Biologically, this means host 1 is a sink or small source and some subset of the other host species are large sources. This is more likely to occur when (a) host 1 is either a sink or a small source ($\varphi_1$ negative or positive and small in magnitude, resulting in $S_1^* \vec{E}_N[1]$ being either negative or positive and small in magnitude), (b) some subset of the host species $j > 1$ are large sources ($\varphi_j$ positive and large for some $j \neq 1$), and (c) infected individuals have larger uptake rates than susceptible individuals ($u_{I_i} > u_{S_i}$). Note that condition (ii) can be seen by rewriting $\theta_n$ as $\theta_n = \sum_k \chi_k (1 - X_k^*) Y_k^* N_k^* + (u_{I_k} - u_{S_k}) X_k^* Y_k^* N_k^* P^*$; $\theta_{n-1}$ can be rewritten in a similar way. •

#### S1.2 Relationship between environmental and direct transmission models

As show in Cortez and Duffy (2021) and Cortez (2021), the 1-habitat environmental transmission models (S1), (S2), and (S4) reduce to direct transmission models when there is a separation of tine scales between the host dynamics and infectious propagule dynamics. In particular, when the shedding rates ($\chi_i$) and either the uptake rates ($u_{V_i}$) or degradation ($\delta$) rates are sufficiently large, then the infectious propagule dynamics will be much faster than the dynamics of the hosts. In this case, we can set $dP/dt = 0$ in the 1-habitat NY model (S4), solve for $P$, and substitute into the other equations to get a direct transmission model of the form

$$\begin{aligned}
\frac{dN_i}{dt} &= \overbrace{F_i(\cdot)}^{\text{reproduction}} - \overbrace{m_i N_i}^{\text{natural mortality}} - \overbrace{(\mu_i Y_i N_i + \xi_i \Gamma_i Y_i N_i)}^{\text{disease-related mortality}} \\
\frac{dY_i}{dt} &= \overbrace{\beta_i X_i \sum_j \frac{\chi_j Y_j N_j}{\delta + \sum_k (u_{S_k} X_k + u_{I_k} Y_k + u_{R_k} Z_k) N_k} P}^{\text{infection}} - \overbrace{(\mu_i + m_i) Y_i}^{\text{mortality}} - \overbrace{\nu_i Y_i}^{\text{recovery}} - \frac{Y_i}{N_i} \frac{dN_i}{dt}
\end{aligned} \tag{S32}$$

where the direct transmission rate from host $j$ to host $i$ is $\beta_i \chi_j S_i I_j / [\delta + \sum_k (u_{S_k} S_k + u_{I_k} I_k + u_{R_k} R_k)]$.

**Density-dependent direct transmission (DDDT):** Model (S32) reduces to a DDDT model when uptake is negligible ($u_{V_i} = 0$). Specifically, under this condition, the model reduces to a form with direct transmission rates $\hat{\beta}_{ij} S_i I_j$ where $\hat{\beta}_{ij} = \beta_i \chi_j / \delta$.

**Frequency-dependent direct transmission (FDDT):** Model (S32) reduces to a frequency-dependent direct transmission (FDDT) model when degradation is negligible ($\delta = 0$). Under this condition, the model reduces to a form with direct transmission rates $\hat{\beta}_{ij} S_i I_j / \sum_k (w_{S_k} S_k + w_{I_k} I_k + w_{R_k} R_k)$ where $\hat{\beta}_{ij} = \beta_i \chi_j$ and $w_{V_k} = u_{V_k}$ for $V \in \{S, I, R\}$; the values of $w_{V_k}$ are the weights for the frequencies of contacts between the different classes of individuals. In particular, if all host species have equal uptake rates ($u_{V_k} = 1$ for all $k$ and $V \in \{S, I, R\}$), then the direct transmission rates have the familiar form $\hat{\beta}_{ij} S_i I_j / \sum_k N_k$.

**Additional points:** First, if uptake and degradation are both non-negligible, then the transmission rate in model (S32) is intermediate to DDDT and FDDT. Thus, ET models allow one to explore how less restrictive assumptions about direct contacts between individuals influence disease risk and spread.

Second, my equilibrium-based results for the 1-habitat environmental transmission models (S1), (S2), and (S4) apply to the direct transmission models even if there is not a separation of time scales between the host and infectious propagule dynamics. That is because the equilibria of 1-habitat environmental transmission models (S1), (S2), and (S4) and the direct transmission model (S32) are identical. Thus, my results apply to DDDT models whenever uptake is negligible ($u_{V_i} = 0$) and my results apply to FDDT models whenever degradation is negligible ($\delta = 0$) and all of the uptake rates are equal to one ($u_{V_i} = 1$).

Third, the 1-habitat ET model cannot reduce down to all possible parameterizations of DDDT and FDDT models. This is because the direct transmission parameters are constrained by $\hat{\beta}_{ij} = \beta_i \chi_j / \delta$ and $\hat{\beta}_{ij} = \beta_i \chi_j$, which implies that $\hat{\beta}_{ij}$ and $\hat{\beta}_{jj}$ are not independent. This explains why the results for the $n$-habitat models in Appendix S2 are needed to fully explain the results for multi-host models with DDDT and FDDT.

#### S1.3 Definition of competence, sink and source

The following provides additional information about the definitions of competence, sink and source, and the relationship between them.

**Definition of host competence**: As in Cortez (2021), the competence of host $i$ is defined by its basic reproduction number in the limit of infinite conspecific host density,

$$\mathcal{R}_{0,i}(\infty) = \frac{\beta_i \chi_i}{u_{S_i}(m_i + \mu_i + \nu_i)} = \rho_i \frac{\chi_i}{(m_i + \mu_i + \nu_i)}; \tag{S33}$$

see Section S1.4 for the mathematical derivation of the basic reproduction number. Higher competence hosts have larger values of $\mathcal{R}_{0,i}(\infty)$ and lower competence hosts have smaller values of $\mathcal{R}_{0,i}(\infty)$.

**Definitions of source and sink**: Host $i$ is a source or sink of infectious propagules at equilibrium if increases in the density of host $i$ (while holding the frequencies of all host classes constant) cause an increase or decrease, respectively, of infectious propagules. Mathematically, sink and source are defined by the per capita net production rate of infectious propagules at equilibrium. Specifically, host $i$ is a source if $\varphi_i = (1/Y_i^*)(\partial/\partial N_i)(dP/dt)|_{p^*} > 0$ and a sink if $\varphi_i = (1/Y_i^*)(\partial/\partial N_i)(dP/dt)|_{p^*} < 0$. Using the equilibrium conditions for the 1-habitat NYZ model (S2), the quantity $\varphi_i$ can be written in terms of the host parameter values and equilibrium density of infectious propagules,

$$\varphi_i = \frac{1}{Y_i^*}\frac{\partial}{\partial N_i}\frac{dP}{dt} = \chi_i - \frac{(\mu_i + m_i + \nu_i)u_{S_i}}{\beta_i} - u_{I_i}P^* - \Gamma_i u_{R_i} P^* \tag{S34}$$

where $\Gamma_i = \nu_i/(\xi_i + m_i + \gamma_i)$.

**Notes about definitions**: The key benefit of using the above definitions of competence and source/sink is that the definitions are independent of host density. Thus, one does not need to know the specific make-up of a community in order to determine if one host species has higher versus lower competence than another or if one host is a smaller or larger source/sink.

However, the definitions of competence and sink/source depend on the host's environment, and that means the orderings of the host species depend on the hosts' environment. In particular, because both definitions depend on the non-disease mortality rates ($m_i$), the designations of high versus low competence and sink versus source may change if non-disease mortality rates are very different between two environments. In addition, the designations of sink versus source may differ between two environments if equilibrium infectious propagule densities are much higher in one environment than another. In total, the definitions allow one to order the hosts in terms of competence and net production of infectious propagules (i.e., sink/source), but those orderings may differ between different environments.

**Relationship between competence and sink/source**: As noted in the main text, the quantities defining competence and sink/source are related, but distinct, because sink/source depends on the instantaneous net production rate of infectious propagules whereas competence depends on the lifetime production of infectious propagules. As a result, both high competence and low competence hosts can be sources or sinks.

There are some restrictions on the possible combinations of values of $\mathcal{R}_{0,i}(\infty)$ and $\varphi_i$ because the quantities defining competence and sink/source are not independent. The specific restrictions are:

(i) If the competence of host $i$ is less than 1, then host $i$ is necessarily a sink. The mathematical condition is $\mathcal{R}_{0,i}(\infty) < 1$ implies $\varphi_i < 0$.

(ii) If host $i$ is a source, then the competence of host $i$ must be greater than 1. The mathematical condition is $\varphi_i > 0$ implies $\mathcal{R}_{0,i}(\infty) > 1$.

Condition (i) means that if the pathogen cannot increase from low abundance in a single-host community of just host $i$, then host $i$ must necessarily be a net remover of infectious propagules (i.e., a sink) in every community. Condition (ii) means that if a pathogen is a net producer of infectious propagules (i.e., a source) in any community, then the pathogen can increase from low abundance in a single-host community of just host $i$.

The mathematical justification is equation (S34) can be rewritten as

$$\varphi_i = \chi_i \left(1 - \frac{1}{\mathcal{R}_{0,i}(\infty)}\right) - u_{I_i} P^* - \Gamma_i u_{R_i} P^* \tag{S35}$$

$$\mathcal{R}_{0,i} = \left[1 - \frac{1}{\chi_i}(\varphi_i + u_{I_i} P^* + \Gamma_i u_{R_i} P^*)\right]^{-1}. \tag{S36}$$

When $\mathcal{R}_{0,i}$ is less than 1, the first equation must be negative. Hence, $\mathcal{R}_{0,i}(\infty) < 1$ implies $\varphi_i < 0$. When $\varphi_i > 0$, the second equation must be greater than 1. Hence, $\varphi_i > 0$ implies $\mathcal{R}_{0,i}(\infty) > 1$.

**Host competence in direct transmission models**: Competence measures the steps of infection given exposure, within-host development of infectious propagules, and pre-transmission survival of the host (Merrill and Johnson 2020). To apply this definition to DDDT models, one needs to decompose the transmission parameter as $\hat{\beta}_{ij} = \hat{E}_{ij}\hat{\chi}_j\hat{\rho}_i$ where $\hat{E}_{ij}$ is the contact rate between susceptible individuals of host $i$ and infected individuals of host $j$, $\hat{\chi}_j$ is the average infectious dose that an infected individual of host $j$ exposes a contact to, and $\hat{\rho}_i$ is the probability of infection for susceptible individuals of host $i$ given exposure. From this, host competence for direct transmission pathogens is $\mathcal{R}_{0,i}(\infty) = \hat{\rho}_i\hat{\chi}_i/[m_i+\mu_i+\nu_i]$.

**Sink and source hosts in direct transmission models**: The definition of sink and source also apply to DDDT and FDDT models. For density-dependent direct transmission (DDDT), all hosts are sources ($\varphi_i > 0$). This is because DDDT corresponds to $u_{S_i} = u_{I_i} = u_{R_i} = 0$, which means equation (S34) is always positive.

For frequency-dependent direct transmission (FDDT), the host species can be sources or sinks. FDDT corresponds to the parameter conditions $\delta = 0$ and $u_{S_i} = u_{I_i} = u_{R_i} = 1$. Under these conditions, equation (S34) is positive when $\chi_i$ is sufficiently large (meaning host $i$ is a source) and negative when $\chi_i$ is sufficiently small (meaning host $i$ is a sink). Because the frequency-dependent direct transmission rates are $\hat{\beta}_{ij} S_i I_j / \sum_k N_k = \beta_i \chi_j S_i I_j / \sum_k N_k$, host $i$ is a source ($\varphi_i > 0$) if it has sufficiently high interspecific transmission rates and host $i$ is a sink ($\varphi_i < 0$) if it has sufficiently low interspecific transmission rates.

#### S1.4 Calculations for $\mathcal{R}_0^C$ sensitivities

The community reproduction number, $\mathcal{R}_0^C$, is computed using the Next Generation Matrix method (Van den Driessche and Watmough 2008, Diekmann et al. 2010). The next generation matrix for the 1-habitat model (S1) is computed from the submatrix of the Jacobian for the infected class and infectious propagules. When evaluated at the disease-free equilibrium, $p^\ddagger$, the submatrix has the form

$$\mathcal{J} = \begin{bmatrix} -\mathbf{D}_{m+\mu+\nu} & \overrightarrow{\beta N^\ddagger} \\ \vec{\chi} & -U^\ddagger - \delta \end{bmatrix} \tag{S37}$$

where $\mathbf{D}_{m+\mu+\nu}$ is a diagonal matrix with entries $m_1+\mu_1+\nu_1, ..., m_n+\mu_n+\nu_n$, $\vec{\chi} = (\chi_1, ..., \chi_n)$, and $\overrightarrow{\beta N^\ddagger} = (\beta_1 N_1^\ddagger, ..., \beta_n N_n^\ddagger)^{\mathrm{T}}$. The submatrix is written as the sum of two matrices,

$$\mathcal{F} = \begin{bmatrix} \mathbf{0} & \overrightarrow{\beta N^\ddagger} \\ \vec{0} & 0 \end{bmatrix}, \quad \mathcal{V} = \begin{bmatrix} -\mathbf{D}_{m+\mu+\nu} & \vec{0} \\ \vec{\chi} & -U^\ddagger - \delta \end{bmatrix}, \tag{S38}$$

which are used to compute the product

$$-\mathcal{F}\mathcal{V}^{-1} = \begin{bmatrix} \mathbf{0} & \overrightarrow{\beta N^\ddagger} \\ \mathbf{0} & \mathbf{0} \end{bmatrix} \begin{bmatrix} \mathbf{D}_{1/(m+\mu+\nu)} & \vec{0} \\ \vec{\chi}\mathbf{D}_{1/(m+\mu+\nu)}/(U^\ddagger+\delta) & 1/(U^\ddagger+\delta) \end{bmatrix}. \tag{S39}$$

The $i, j$ entry of $-\mathcal{F}\mathcal{V}^{-1}$ is $\frac{\chi_j}{\mu_j+m_j+\gamma_j}\frac{\beta_i N_i^\ddagger}{U^\ddagger+\delta}$. The community reproduction number is the largest positive eigenvalue of $-\mathcal{F}\mathcal{V}^{-1}$,

$$\mathcal{R}_0^C = \frac{1}{U^\ddagger+\delta}\sum_i \frac{\beta_i\chi_i N_i^\ddagger}{(m_i+\mu_i+\nu_i)}. \tag{S40}$$

The basic reproduction number for a single host species is

$$\mathcal{R}_{0,i} = \frac{\beta_i\chi_i N_i^\ddagger}{(u_{S_i}N_i^\ddagger+\delta)(m_i+\mu_i+\nu_i)}. \tag{S41}$$

Note that the basic reproduction number in the limit of infinite host density defines the competence of host $i$, $\mathcal{R}_{0,i}(\infty) = \beta_i\chi_i/u_{S_i}(m_i+\nu_i+\nu_i)$.

##### S1.4.1 Sensitivity of $\mathcal{R}_0^C$ to host densities

**Summary of biological interpretation**: In the absence of interspecific competition, increased density of host $n$ increases or decreases $\mathcal{R}_0^C$ if its competence is sufficiently high or low relative to $\mathcal{R}_0^C$. When interspecific competition is present, increased density of host $n$ causes increased $\mathcal{R}_0^C$ if higher competence hosts increase more or decrease less in density than lower competence hosts; increased density of host $n$ causes decreased $\mathcal{R}_0^C$ if higher competence hosts decrease more or increase less in density than lower competence hosts.

The interpretation for FDDT is the same as the above. For DDDT, increased density of host $n$ always increases $\mathcal{R}_0^C$ when there is no interspecific competition. When interspecific competition is present, increased density of host $n$ increases $\mathcal{R}_0^C$ unless interspecific competition is sufficiently strong.

**Calculations for Environmental Transmission**: Let $\alpha$ be a parameter satisfying $\partial F_n/\partial\alpha < 0$ and $\partial F_i/\partial\alpha = 0$ for $i \neq n$. Assume $\partial N^\ddagger/\partial\alpha < 0$. After some algebra, the sensitivity of $\mathcal{R}_0^C$ to the density of host species $n$ is

$$\frac{d\mathcal{R}_0^C}{dN_n^\ddagger} = \frac{1}{U^\ddagger+\delta}\sum_i \frac{\beta_i\chi_i}{(m_i+\mu_i+\nu_i)}\frac{dN_i^\ddagger}{dN_n^\ddagger} - \frac{1}{(U^\ddagger+\delta)^2}\left(\sum_i \frac{\beta_i\chi_i}{(m_i+\mu_i+\nu_i)}\right)\left(\sum_i u_{S_i}\frac{dN_i^\ddagger}{dN_n^\ddagger}\right) \tag{S42}$$

$$= \frac{1}{U^{\ddagger} + \delta} \left[ \sum_i u_{S_i} \left( \mathcal{R}_{0,i}(\infty) - \mathcal{R}_0^C \right) \frac{dN_i^{\ddagger}}{dN_n^{\ddagger}} \right] \tag{S43}$$

$$= C_n^{\ddagger} \left( \mathcal{R}_{0,n}(\infty) - \mathcal{R}_0^C \right) + \sum_{i \neq n} C_i^{\ddagger} \left( \mathcal{R}_{0,i}(\infty) - \mathcal{R}_0^C \right) \alpha_{in}^{total\ddagger} \tag{S44}$$

where

$$\frac{dN_i^{\ddagger}}{dN_n^{\ddagger}} = \frac{(-1)^{i+n} |\mathbf{J}^{\ddagger[\not{n},\not{i}]}|}{|\mathbf{J}^{\ddagger}|} \Bigg/ \frac{(-1)^{n+n} |\mathbf{J}^{\ddagger[\not{n},\not{n}]}|}{|\mathbf{J}^{\ddagger}|}, \tag{S45}$$

$$C_i^{\ddagger} = \frac{-u_{S_i}}{U^{\ddagger} + \delta} \left( \frac{\partial N_n^{\ddagger}}{\partial \alpha} \right) > 0 \quad (i \neq n), \tag{S46}$$

$$C_n^{\ddagger} = \frac{u_{S_n}}{U^{\ddagger} + \delta} > 0, \tag{S47}$$

$$\alpha_{in}^{total\ddagger} = (-1)^{i+n+1} \frac{|\mathbf{J}^{\ddagger[\not{n},\not{i}]}|}{|\mathbf{J}^{\ddagger}|}. \tag{S48}$$

In the above, $\mathbf{J}^{\ddagger}$ is the submatrix of the Jacobian for the subsystem where only susceptible individuals are present, $\mathbf{J}^{\ddagger[\not{n},\not{i}]}$ is the submatrix of $\mathbf{J}^{\ddagger}$ with its $n$th row and $i$th column removed, and the total competitive effect of host $n$ on host $i$ is positive when $\alpha_{in}^{total\ddagger} > 0$ and negative when $\alpha_{in}^{total\ddagger} < 0$.

In the absence of interspecific competition, $\alpha_{in}^{total\ddagger} = 0$ and equation (S44) simplifies to its first term. In this case, equation (S44) is positive if host $n$ has sufficiently high competence relative to $\mathcal{R}_0^C$ (i.e., $\mathcal{R}_{0,n}(\infty) - \mathcal{R}_0^C > 0$) and negative if host $n$ has sufficiently low competence relative to $\mathcal{R}_0^C$ (i.e, $\mathcal{R}_{0,n}(\infty) - \mathcal{R}_0^C < 0$).

When interspecific competition is present, each term in the sum in equation (S44) is positive if host $n$ has a positive total competitive effect on host $i$ and host $i$ has high competence ($\alpha_{in}^{total\ddagger} > 0$ and $\mathcal{R}_{0,n}(\infty) - \mathcal{R}_0^C > 0$) or if host $n$ has a negative total competitive effect on host $i$ and host $i$ has low competence ($\alpha_{in}^{total\ddagger} < 0$ and $\mathcal{R}_{0,n}(\infty) - \mathcal{R}_0^C < 0$). This means that equation (S44) is negative when host $n$ has smaller negative and larger positive total competitive effects on high competence hosts and larger negative and smaller positive total competitive effects on low competence hosts.

**Density-Dependent Direct Transmission**: DDDT corresponds to $u_{V_i} = 0$ ($V \in \{S, I, R\}$), which means the sensitivity simplifies to

$$\frac{d\mathcal{R}_0^C}{dN_n^{\ddagger}} = \sum_i \frac{\beta_i \chi_i}{\delta(m_i + \mu_i + \nu_i)} \frac{dN_i^{\ddagger}}{dN_n^{\ddagger}} \tag{S49}$$

$$= \frac{-\beta_n \chi_n}{\delta(m_n + \mu_n + \nu_n)} \left( \frac{\partial N_n^{\ddagger}}{\partial \alpha} \right) + \sum_{i \neq n} \frac{-\beta_i \chi_i}{\delta(m_i + \mu_i + \nu_i)} \left( \frac{\partial N_n^{\ddagger}}{\partial \alpha} \right) \alpha_{in}^{total\ddagger}. \tag{S50}$$

where $\partial N_n^{\ddagger} / \partial \alpha < 0$ and

$$\alpha_{in}^{total\ddagger} = (-1)^{i+n+1} \frac{|\mathbf{J}^{\ddagger[\not{n},\not{i}]}|}{|\mathbf{J}^{\ddagger}|}. \tag{S51}$$

In the absence of interspecific competition, equation (S50) is positive because $\alpha_{in}^{total\ddagger} = 0$ for $i \neq n$. When interspecific competition is present, equation (S50) is positive unless the added host has sufficiently strong negative total effects on other host species ($\alpha_{in}^{total\ddagger} < 0$) such that the sum is negative and larger in magnitude than the first term in equation (S50). Biologically, this requires sufficiently strong interspecific competition.

##### S1.4.2 Sensitivity of $\mathcal{R}_0^C$ to epidemiological parameters

It is straightforward to show that the sensitivities to the epidemiological parameters satisfy

$$\frac{\partial \mathcal{R}_0^C}{\partial \beta_i} \geq 0, \quad \frac{\partial \mathcal{R}_0^C}{\partial \chi_i} \geq 0 \tag{S52}$$

$$\frac{\partial \mathcal{R}_0^C}{\partial \delta} \leq 0, \quad \frac{\partial \mathcal{R}_0^C}{\partial \mu_i} \leq 0, \quad \frac{\partial \mathcal{R}_0^C}{\partial \nu_i} \leq 0, \quad \frac{\partial \mathcal{R}_0^C}{\partial u_{S_i}} \leq 0. \tag{S53}$$

**Summary of biological interpretation**: Increases in the competence of a host species (larger $\beta_i$, $\chi_i$ or smaller $\nu_i$, $\mu_i$, $u_{S_i}$) always increase the community reproduction number. Decreases in the competence of a host species (smaller $\beta_i$, $\chi_i$ or larger $\nu_i$, $\mu_i$, $u_{S_i}$) always decrease the community reproduction number.

##### S1.4.3 Sensitivity of $\mathcal{R}_0^C$ to pathogen transmission mode

Here, I use a continuous change of parameters to convert the 1-habitat ET model (S1) from a FDDT form ($\delta = 0$ for all $i$) to a DDDT form ($u_{V_i} = 0$ for $V \in \{S, I, R\}$). The change of parameters holds all parameters constant except $\delta$ and $u_{V_i}$, which must necessarily change. The specific change of parameters defined below is chosen such that the community reproduction number for a particular assemblage of hosts species is held constant. The biological interpretation of this constraint is that the pathogen's community reproduction number is known for a particular assemblage of species, the pathogen's transmission mode is unknown, and the pathogen's community reproduction number is unknown when a set of hosts are added to or lost from the community. This method allows one to predict if greater increases (or smaller decreases) in the value of $\mathcal{R}_0^C$ would occur under FDDT or DDDT when host species are added to or lost from a community.

Note that in order to apply the following theory, one must know/assume the ratio of the uptake rates for all host species, i.e., $u_{S_i}/u_{S_j}$. That information is needed to define the change of parameters functions.

**Summary of biological interpretation**: When host species are added to a community, large reductions or smaller increases in $\mathcal{R}_0^C$ are more likely to occur under FDDT than DDDT when added host species are weak intraspecific and interspecific competitors and added host species have higher uptake rates. In comparison, larger reductions and smaller increases in $\mathcal{R}_0^C$ are more likely to occur under DDDT than FDDT when added host species are strong intraspecific and interspecific competitors and added host species have lower uptake rates. In addition, in the absence of interspecific competition, reductions in $\mathcal{R}_0^C$ are always more likely under FDDT than DDDT. The justification for this interpretation is given after the calculations.

**Additions of host species to a community**: Without loss of generality, assume host species 1,...,$m$ ($m < n$) are present in a community. Assume that species stably coexist at a disease-free equilibrium (hereafter, the 'subcommunity equilibrium'). Let $N_i^\dagger$ denote the density of host $i$ at the subcommunity equilibrium. The pathogen's reproduction number at the subcommunity equilibrium is

$$\mathcal{R}_0^\dagger = \frac{1}{U^\dagger + \delta} \sum_{i=1}^{m} \frac{\beta_i \chi_i N_i^\dagger}{(m_i + \mu_i + \nu_i)}. \tag{S54}$$

where $U^\dagger = \sum_{i=1}^{m} u_{S_i} N_i^\dagger$ denotes the per propagule rate at which infectious propagules are taken up by host species 1,...,$m$ at the subcommunity equilibrium.

The infectious propagule equation is rewritten as

$$\frac{dP}{dt} = \sum_i \chi_i I_i - g(q) \sum_{i=1}^{n} \left(u_{S_i} S_i + u_{I_i} I_i + u_{R_i} R_i\right) P - h(q) \delta P \tag{S55}$$

where the functions

$$g(q) = \begin{cases} \dfrac{-\delta}{U^\dagger} q + 1 + \dfrac{\delta}{U^\dagger}, & 0 \leq q \leq 1 \\ 2 - q, & 1 \leq q \leq 2 \end{cases} \tag{S56}$$

$$h(q) = \begin{cases} q, & 0 \leq q \leq 1 \\ \dfrac{-U^\dagger}{\delta}(2 - q) + 1 + \dfrac{U^\dagger}{\delta}, & 1 \leq q \leq 2 \end{cases} \tag{S57}$$

transform the 1-habitat ET model (S1) from a FDDT form when $q = 0$ to a DDDT form when $q = 2$. The transformation holds constant the value of the pathogen's reproduction number for the subcommunity ($\mathcal{R}_0^\dagger$).

Differentiating equation (S41) with respect to $q$ and some algebra yields the sensitivity

$$\frac{\partial \mathcal{R}_0^C}{\partial q} = \begin{cases} \frac{\delta}{U^\dagger}(U^\ddagger - U^\dagger) \frac{\mathcal{R}_0^C}{\delta + U^\ddagger}, & 0 \leq q \leq 1 \\ (U^\ddagger - U^\dagger) \frac{\mathcal{R}_0^C}{\delta + U^\ddagger}, & 1 \leq q \leq 2 \end{cases} \tag{S58}$$

where all derivatives at $q = 1$ are interpreted as right or left derivatives. The sign of the derivative depends on $U^\ddagger - U^\dagger = \sum_{i>m} u_{S_i} N_i^\ddagger + \sum_{i \leq m} u_{S_i} (N_i^\ddagger - N_i^\dagger)$.

**Removal of host species from a community**: The notation is the same as the above, where daggers denotes quantities associated with the subcommunity equilibrium for host species 1,...,$m$ ($m < n$). The infectious propagule equation is rewritten as in equation (S55) where the functions $g(q)$ and $h(q)$ are now defined as

$$g(q) = \begin{cases} \dfrac{-\delta}{U^\ddagger} q + 1 + \dfrac{\delta}{U^\ddagger}, & 0 \leq q \leq 1 \\ 2 - q, & 1 \leq q \leq 2 \end{cases} \tag{S59}$$

$$h(q) = \begin{cases} q, & 0 \leq q \leq 1 \\ \dfrac{-U^\ddagger}{\delta}(2-q) + 1 + \dfrac{U^\ddagger}{\delta}, & 1 \leq q \leq 2 \end{cases} \tag{S60}$$

Importantly, this transforms the 1-habitat ET model (S1) from a FDDT form when $q = 0$ to a DDDT form when $q = 2$ while holding constant the value of the pathogen's reproduction number for the $n$-species community ($\mathcal{R}_0^C$).

Differentiating equation (S54) with respect to $q$ and some algebra yields the sensitivity

$$\frac{\partial \mathcal{R}_0^\dagger}{\partial q} = \begin{cases} -\frac{\delta}{U^\dagger}(U^\ddagger - U^\dagger)\frac{\mathcal{R}_0^\dagger}{\delta + U^\dagger}, & 0 \leq q \leq 1 \\ -(U^\ddagger - U^\dagger)\frac{\mathcal{R}_0^\dagger}{\delta+U^\dagger}, & 1 \leq q \leq 2 \end{cases} \tag{S61}$$

where all derivatives at $q = 1$ are interpreted as right or left derivatives. The sign of the derivative depends on $U^\ddagger - U^\dagger = \sum_{i>m} u_{S_i} N_i^\ddagger + \sum_{i \leq m} u_{S_i}(N_i^\ddagger - N_i^\dagger)$.

**Justification for interpretation**: Equation (S58) is positive when equation (S61) is negative and vice versa. To avoid confusion, I will only focus on equation (S58); the interpretation is qualitatively similar for equation (S61).

The sign of equation (S58) is determined by $U^\ddagger - U^\dagger$. The quantity $U^\ddagger - U^\dagger$ is positive unless $N_i^\ddagger < N_i^\dagger$ for $i \leq m$, $u_{S_i}$ sufficiently large for $i \leq m$, and $u_{S_i} N_i^\dagger$ is small for $i > m$. This means that equation (S58) is more likely to be positive when (i) species $i > m$ have weak intraspecific interactions, (ii) species $i > m$ have weak interspecific interactions on species $i \leq m$, and (iii) species $i > m$ have higher uptake rates than species $i \leq m$. Thus, adding host species to a community is more likely to cause larger decreases (or smaller increases) in $R_0$ under FDDT than DDDT when conditions (i)-(iii) are satisfied. In comparison, equation (S58) is more likely to be negative when (iv) species $i > m$ have strong intraspecific interactions, (v) species $i > m$ have strong interspecific interactions on species $i \leq m$, and (vi) species $i > m$ have lower uptake rates than species $i \leq m$. Thus, adding host species to a community is more likely to cause larger decreases (or smaller increases) in $R_0$ under DDDT than FDDT when conditions (iv)-(vi) are satisfied.

In the absence of interspecific competition, $N_i^\ddagger = N_i^\dagger$ for $i \leq m$, which means $U^\ddagger - U^\dagger > 0$. Consequently, equation (S58) is positive. This means that if the pathogen reproduction number is held constant and one or more host species are added to a community, then (i) decreases in the value of $\mathcal{R}_0^C$ under DDDT only occur if the additions also decrease the value of $\mathcal{R}_0^C$ under FDDT and (ii) increases in the value of $\mathcal{R}_0^C$ under FDDT necessarily imply increases in the value of $\mathcal{R}_0^C$ under DDDT. Consequently, in the absence of interspecific competition, decreases in $\mathcal{R}_0^C$ are more likely under FDDT than DDDT and increases in $\mathcal{R}_0^C$ are more likely under DDDT than FDDT.

#### S1.5 Calculations for sensitivities of infection prevalence and infected density

All calculations for the sensitivities of infection prevalence and infected density use the Jacobian of the 1-habitat NY model (S4). The Jacobian is defined in equation (S5) of Section S1.1.3 and its properties are defined in Section S1.1.5. All of the interpretations in the following subsections assume the assumptions in Section S1.1.1 hold. Without loss of generality, I focus on responses in host 1 (the focal host) to changes in the density and characteristics of host $n$. I note that the results for the sensitivities of host 1 infection prevalence are mathematically equivalent to those in Cortez (2021).

##### S1.5.1 Method for computing local sensitivities

Following previous studies (Cortez 2021, Cortez and Duffy 2021), the local sensitivities of host $i$ infection prevalence at equilibrium ($Y_i^*$) and host $i$ total density ($N_i^*$) to a model parameter ($\sigma$) are computed from the Jacobian, $\mathbf{J}$, of the 1-habitat NY model (S4) via the partial derivatives,

$$\begin{aligned}\frac{\partial Y_i^*}{\partial \sigma} =&\sum_{k\leq n}\frac{(-1)^{k+i+n}}{|\mathbf{J}|}\left(-\frac{\partial}{\partial\sigma}\frac{dN_k}{dt}\right)|\mathbf{J}^{[\cancel{k},\cancel{i+n}]}|+\sum_{k\leq n}\frac{(-1)^{k+n+i+n}}{|\mathbf{J}|}\left(-\frac{\partial}{\partial\sigma}\frac{dY_k}{dt}\right)|\mathbf{J}^{[\cancel{k+n},\cancel{i+n}]}| \\ &+\frac{(-1)^{2n+1+i+n}}{|\mathbf{J}|}\left(-\frac{\partial}{\partial\sigma}\frac{dP}{dt}\right)|\mathbf{J}^{[\cancel{2n+1},\cancel{i+n}]}| \end{aligned} \tag{S62}$$

$$\begin{aligned}\frac{\partial N_i^*}{\partial \sigma} =&\sum_{k\leq n}\frac{(-1)^{k+i}}{|\mathbf{J}|}\left(-\frac{\partial}{\partial\sigma}\frac{dN_k}{dt}\right)|\mathbf{J}^{[\cancel{k},\cancel{i}]}|+\sum_{k\leq n}\frac{(-1)^{k+n+i}}{|\mathbf{J}|}\left(-\frac{\partial}{\partial\sigma}\frac{dY_k}{dt}\right)|\mathbf{J}^{[\cancel{k+n},\cancel{i}]}| \\ &+\frac{(-1)^{2n+1+i}}{|\mathbf{J}|}\left(-\frac{\partial}{\partial\sigma}\frac{dP}{dt}\right)|\mathbf{J}^{[\cancel{2n+1},\cancel{i}]}| \end{aligned} \tag{S63}$$

To compute the local sensitivities of host $i$ infected density at equilibrium ($I_i^*$) to an epidemiological parameter ($\sigma$), I use the chain rule to compute

$$\frac{\partial Y_1^*}{\partial\sigma}=\frac{\partial}{\partial\sigma}\frac{I_1^*}{N_1^*}=\frac{1}{(N_1^*)^2}\left(N_1^*\frac{\partial I_1^*}{\partial\sigma}-I_1^*\frac{\partial N_1^*}{\partial\sigma}\right) \tag{S64}$$

which can be rearranged to get

$$\frac{\partial I_1^*}{\partial\sigma}=N_1^*\frac{\partial Y_1^*}{\partial\sigma}+Y_1^*\frac{\partial N_1^*}{\partial\sigma}. \tag{S65}$$

The sensitivities of host $i$ infection prevalence ($Y_i^*$) and host $i$ infected density ($I_i^*$) to the total density of host $n$ ($N_n^*$) are computed using the partial derivatives

$$\frac{dY_i^*}{dN_n^*}=\frac{\partial Y_i^*}{\partial\sigma}\Bigg/\frac{\partial N_n^*}{\partial\sigma} \tag{S66}$$

$$\frac{dI_i^*}{dN_n^*}=\frac{\partial I_i^*}{\partial\sigma}\Bigg/\frac{\partial N_n^*}{\partial\sigma}. \tag{S67}$$

I note the following about the method. First, when computing the sensitivities to epidemiological parameters of host $n$, $\sigma$ is determined by the specific parameter of interest (e.g., $\beta_n$ or $\mu_n$).

Second, when computing the sensitivities to the density of host $n$, I chose a parameter representing the level of intraspecific competition experienced by host $n$. Throughout, I use $\alpha$ to denote such a parameter. In a Lokta-Volterra competition model, $\alpha$ would be the intraspecific competition coefficient $\alpha_{nn}$. In those calculations, I assume (i) increases in $\alpha$ decrease the growth rate of host $n$, (ii) $\alpha$ does not directly effect the growth rates of the other host species, and (iii) the equilibrium density for host $n$ decreases with increases in $\alpha$. The respective mathematical assumptions are (i) $\frac{\partial}{\partial \alpha}\frac{dN_n}{dt} < 0$, (ii) $\frac{\partial}{\partial \alpha}\frac{dN_k}{dt} = 0$ for $k \neq n$, and (iii) $\frac{\partial N_n^*}{\partial \alpha} < 0$. The first two assumptions are guaranteed to hold with the correct choice of parameter. The third assumption may not hold in all systems. When is does not hold, host $n$ experiences a generalized hydra effect (i.e., increased equilibrium density with a decrease in its per capita growth rate). I do not consider systems where host $n$ exhibits a generalized hydra effects because generalized hydra effects are less likely to be present in natural systems than absent. However, I note that all of the results in Section S1.5.2 can be applied to systems where host $n$ is exhibiting a generalized hydra effect simply by changing the sign of the predicted response.

##### S1.5.2 Sensitivities to host densities

**Summary of biological interpretation**: If interspecific host competition is sufficiently weak, then increased density of host $n$ increases host 1 infection prevalence when host $n$ is a source and decreases host 1 infection prevalence when host $n$ is a sink. If interspecific host competition is not weak, then increased density of host $n$ can increase host 1 infection prevalence when host $n$ has stronger negative (or weaker positive) competitive effects on sink hosts than source hosts and increased density of host $n$ can decrease host 1 infection prevalence when host $n$ has stronger negative (or weaker positive) competitive effects on source hosts than sink hosts.

The responses of host 1 infected density to increased density of host $n$ are often the same as the responses of host 1 infection prevalence. However, the predictions for host 1 infected density can have the opposite sign when (i) for each host species, regulation by competition with host 1 is sufficiently weaker than regulation by disease and competition with species $2, ..., n$ or (ii) host 1 is a sink or small source and a subset of the other host species are sufficiently large sources. Condition (i) is more likely to occur when host 1 is a weak intraspecific and interspecific competitor. In addition, condition (i) is easier to satisfy in systems where infected individuals are stronger competitors than susceptible individuals and condition (ii) is easier to satisfy in systems where infected individuals have larger uptake rates than susceptible individuals. See Remark S1.6 for details.

**Sensitivity of infection prevalence**: Let $\alpha$ be a parameter satisfying $\frac{\partial F_n}{\partial \alpha} < 0$ and $\frac{\partial F_j}{\partial \alpha} = 0$ for $j \neq n$. Assume $\partial N_n^*/\partial \alpha < 0$. The sensitivity of host 1 infection prevalence to $\alpha$ is computed as

$$\frac{\partial Y_1^*}{\partial \alpha} = \left(-\frac{\partial F_n}{\partial \alpha}\right)\frac{(-1)^{n+n+1}}{|\mathbf{J}|}|\mathbf{J}^{[\cancel{n},\cancel{n+1}]}| + \left(\frac{Y_n^*}{N_n^*}\frac{\partial F_n}{\partial \alpha}\right)\frac{(-1)^{2n+n+1}}{|\mathbf{J}|}|\mathbf{J}^{[\cancel{2n},\cancel{n+1}]}| \tag{S68}$$

$$= \left(-\frac{\partial F_n}{\partial \alpha}\right)\frac{-1}{|\mathbf{J}|}\begin{vmatrix} \mathbf{A}^{[\cancel{n},-]} & (\mathbf{B}+\mathbf{C}\mathbf{D}_\Gamma)^{[\cancel{n},\cancel{1}]} & \vec{0}_{n-1} \\ \mathbf{0}_{n\times n} & -\mathbf{D}_{\beta P/Y}^{[-,\cancel{1}]} & \overrightarrow{\beta X} \\ \vec{E}_N & \vec{E}_Y^{[\cancel{1}]} & -U-\delta \end{vmatrix} \tag{S69}$$

$$= \left(-\frac{\partial F_n}{\partial \alpha}\right)\frac{(-1)^{1+n+n-1}}{|\mathbf{J}|}\begin{vmatrix} \mathbf{A}^{[\not n,-]} & \vec{0}_{n-1} & (\mathbf{B}+\mathbf{CD}_\Gamma)^{[\not n,\not 1]} \\ \vec{E}_N & -U-\delta & \vec{E}_Y^{[\not 1]} \\ \mathbf{0}_{n\times n} & \overrightarrow{\beta X} & -\mathbf{D}_{\beta P/Y}^{[-,\not 1]} \end{vmatrix} \tag{S70}$$

$$= \left(-\frac{\partial F_n}{\partial \alpha}\right)\frac{1}{|\mathbf{J}|}\begin{vmatrix} \mathbf{A}^{[\not n,-]} & \vec{0}_{n-1} & (\mathbf{B}+\mathbf{CD}_\Gamma)^{[\not n,\not 1]} \\ \vec{E}_N & 0 & \vec{E}_Y^{[\not 1]} \\ \mathbf{0}_{n\times n} & \begin{bmatrix} \beta_1 X_1^* \\ \vec{0}_{n-1} \end{bmatrix} & -\mathbf{D}_{\beta P/Y}^{[-,\not 1]} \end{vmatrix} \tag{S71}$$

$$= \left(-\frac{\partial F_n}{\partial \alpha}\right)\frac{\beta_1 X_1^*}{|\mathbf{J}|}(-1)^{n-1}\prod_{j\neq 1}\left(\frac{\beta_i P^*}{Y_i^*}\right)\begin{vmatrix} \mathbf{A}^{[\not n,-]} \\ \vec{E}_N \end{vmatrix} \tag{S72}$$

$$= \left(-\frac{\partial F_n}{\partial \alpha}\right)\beta_1 X_1^*(-1)^{n-1}\prod_{j\neq 1}\left(\frac{\beta_i P^*}{Y_i^*}\right)\sum_j \frac{(-1)^{n+j}|\mathbf{A}^{[\not n,\not j]}|}{|\mathbf{J}|}\varphi_j Y_j^*. \tag{S73}$$

Under the assumption that $\partial N_n^*/\partial\alpha < 0$, the sensitivity of host 1 infection prevalence to the density of host $n$ is

$$\frac{dY_1^*}{dN_n^*} = C_n^*\varphi_n + \sum_i C_j^*\varphi_j\alpha_{in}^{total} \tag{S74}$$

where

$$C_n^* = \left(\frac{\partial N_n^*}{\partial\alpha}\right)^{-1}\left(-\frac{\partial F_n}{\partial \alpha}\right)\beta_1 X_1^*(-1)^{n-1}\prod_{j\neq 1}\left(\frac{\beta_i P^*}{Y_i^*}\right)\frac{(-1)^{n+n}|\mathbf{A}^{[\not n,\not n]}|}{|\mathbf{J}|}Y_j^* \tag{S75}$$

$$C_i^* = \left(\frac{\partial N_n^*}{\partial\alpha}\right)^{-1}\left(-\frac{\partial F_n}{\partial \alpha}\right)\beta_1 X_1^*(-1)^{n-1}\prod_{j\neq 1}\left(\frac{\beta_i P^*}{Y_i^*}\right)(-1)^n Y_j^* \tag{S76}$$

are positive under the assumptions in Remark S1.2.

The first term in equation (S74) represents the indirect effect $N_n \to P \to Y_1$ and the terms in the sum represent the indirect effects $N_n \to N_i \to P \to Y_1$ ($i \neq n$). In the absence of interspecific competition, only the first term in equation (S74) is non-zero. Thus, increased density of host $n$ increases or decreases focal host infection prevalence when host $n$ is a source ($\varphi_n > 0$) or sink ($\varphi_n < 0$), respectively. When interspecific competition is present, each term in the sum is positive when either host $n$ has a positive total competitive effect on host $i$ ($\alpha_{in}^{total} > 0$) and host $i$ is a source or host $n$ has a negative total competitive effect on host $i$ ($\alpha_{in}^{total} > 0$) and host $i$ is a sink. Thus, equation (S74) will be positive when increased density of host $n$ causes larger increases or smaller decreases in the densities of source hosts and smaller increases or larger decreases in the densities of sink hosts.

**Density-Dependent Direct Transmission**: Under DDDT, all hosts are sources ($\varphi_i > 0$). In the absence of interspecific competition, equation (S74) is positive (because the first term is positive and the sum is zero because $\alpha_{in}^{total} = 0$ for $i \neq n$). Thus, increased density of host $n$ causes increased focal host infection prevalence. When interspecific competition is present, increased density of host $n$ increases focal host infection prevalence unless interspecific competition is sufficiently strong ($\alpha_{in}^{total}$ negative and sufficiently large in magnitude) such that host $n$ greatly suppresses the densities of the other host species.

**Sensitivity of infected density**: The sensitivity of host 1 infection prevalence to $\alpha$ is computed using elementary row operations and Schur's formula to yield

$$\frac{\partial N_1^*}{\partial \alpha} = \left(-\frac{\partial F_n}{\partial \alpha}\right)\frac{(-1)^{n+1}}{|\mathbf{J}|}|\mathbf{J}^{[\not n,\not 1]}| + \left(\frac{Y_n^*}{N_n^*}\frac{\partial F_n}{\partial \alpha}\right)\frac{(-1)^{2n+1}}{|\mathbf{J}|}|\mathbf{J}^{[\not 2\not n,\not 1]}| \tag{S77}$$

$$= \left(-\frac{\partial F_n}{\partial \alpha}\right)\frac{(-1)^{n+1}}{|\mathbf{J}|}\begin{vmatrix} \mathbf{A}^{[\not n,\not 1]} & (\mathbf{B}+\mathbf{C}\mathbf{D}_\Gamma)^{[\not n,-]} & \vec{0}_{n-1} \\ \mathbf{0}_{n\times n-1} & -\mathbf{D}_{\beta P/Y} & \overrightarrow{\beta X} \\ \vec{E}_N^{[\not 1]} & \vec{E}_Y & -U-\delta \end{vmatrix} \tag{S78}$$

$$= \left(-\frac{\partial F_n}{\partial \alpha}\right)\frac{1}{|\mathbf{J}|}\begin{vmatrix} \vec{0}_{n-1} & \mathbf{A}^{[\not n,\not 1]} & (\mathbf{B}+\mathbf{C}\mathbf{D}_\Gamma)^{[\not n,-]} \\ -U-\delta & \vec{E}_N^{[\not 1]} & \vec{E}_Y \\ \overrightarrow{\beta X} & \mathbf{0}_{n\times n-1} & -\mathbf{D}_{\beta P/Y} \end{vmatrix} \tag{S79}$$

$$= \left(-\frac{\partial F_n}{\partial \alpha}\right)\frac{1}{|\mathbf{J}|}\begin{vmatrix} -\vec{\omega}_n^{[\not n]}/P^* & \mathbf{A}^{[\not n,\not 1]} & (\mathbf{B}+\mathbf{C}\mathbf{D}_\Gamma)^{[\not n,-]} \\ -\theta_n/P^* & \vec{E}_N^{[\not 1]} & \vec{E}_Y \\ \vec{0}_n & \mathbf{0}_{n\times n-1} & -\mathbf{D}_{\beta P/Y} \end{vmatrix} \tag{S80}$$

$$= \left(-\frac{\partial F_n}{\partial \alpha}\right)\frac{(-1)^{n-1}}{P^*|\mathbf{J}|}\prod_{j\geq 1}\left(\frac{\beta_i P^*}{Y_i^*}\right)\begin{vmatrix} \vec{\omega}_n^{[\not n]} & \mathbf{A}^{[\not n,\not 1]} \\ \theta_n & \vec{E}_N^{[\not 1]} \end{vmatrix} \tag{S81}$$

where $\vec{\omega}_n$ and $\theta_n$ are defined in Remark S1.6. Using equation (S65) yields the sensitivity of host 1 infected density,

$$\frac{\partial I_1^*}{\partial \alpha} = \left(-\frac{\partial F_n}{\partial \alpha}\right)\frac{(-1)^{n-1}\beta_1}{|\mathbf{J}|}\prod_{j\neq 2}\left(\frac{\beta_i P^*}{Y_i^*}\right)\left(S_1^*\begin{vmatrix} \mathbf{A}^{[\not n,-]} \\ \vec{E}_N \end{vmatrix} + \begin{vmatrix} \vec{\omega}_n^{[\not n]} & \mathbf{A}^{[\not n,\not 1]} \\ \theta_n & \vec{E}_N^{[\not 1]} \end{vmatrix}\right). \tag{S82}$$

The sensitivity $dI_1^*/dN_n^*$ is found by dividing equation (S82) by equation (S81) for $\partial N_n^*/\partial\alpha$.

The two matrices in equation (S82) only differ in the entries of their first columns. The sensitivities $dY_1^*/dN_n^*$ and $dI_1^*/dN_n^*$ can have opposite signs when the second determinant in equation (S82) has the opposite sign as the first determinant and is sufficiently large in magnitude. This is more likely to occur when (i) the entries of $\vec{\omega}_n$ are positive and large in magnitude relative to the entries of $\mathbf{A}_{\bullet 1}$ or (ii) $\theta_n$ is positive and large in magnitude and $\mathbf{E}_N[1]$ is either negative or positive and small in magnitude. Those conditions are discussed in Remark S1.6.

##### S1.5.3 Sensitivities to epidemiological parameters

**Summary of biological interpretation**: The predicted effects of varying the components defining the competence of host $n$ on host 1 infection prevalence are the following.

- Increasing the shedding of host $n$ ($\chi_n$) increases host 1 infection prevalence.

- Increasing the uptake rate of host $n$ ($u_{V_n}$) decreases host 1 infection prevalence.

- Increasing the transmission parameter of host $n$ ($\beta_n$) often increases host 1 infection prevalence. However, host 1 infection infection prevalence can decrease under the following conditions, with conditions 2 and 4 being the most likely to arise in natural systems. (1) Host $n$ is a sink ($\varphi_n < 0$) and infected individuals of host $n$ have much larger uptake rates than susceptible individuals ($u_{I_1} > u_{S_i}$). (2) Host $n$ is a source ($\varphi_i > 0$) and interspecific competition is sufficiently strong. (3) Infected individuals of host $n$ are sufficiently stronger interspecific competitors than susceptible individuals and host $n$ has stronger interspecific competitive effects on source hosts ($\varphi_j > 0$) than sink hosts ($\varphi_j < 0$). (4) Infected individuals of host $n$ are weaker or insufficiently stronger interspecific competitors than susceptible individuals and host $n$ has stronger interspecific competitive effects on sink hosts ($\varphi_j < 0$) than source hosts ($\varphi_j > 0$).

- Increasing the disease-induced mortality in host $n$ ($\mu_n$) often decreases host 1 infection prevalence. However, host 1 infection infection prevalence can increase under the following conditions, with conditions 1 and 2 being the most likely to arise in natural systems. (1) Host $n$ is a sufficiently large sink ($\varphi_n < 0$). (2) Host $n$ has stronger negative (or weaker positive) total competitive effects on sink hosts and weaker negative (or stronger positive) total competitive effects on source hosts. (3) Host $n$ is a sink ($\varphi_n < 0$) and infected individuals of host $n$ have much larger uptake rates than susceptible individuals ($u_{I_1} > u_{S_i}$). (4) Host $n$ is a source ($\varphi_n > 0$) and either (i) infected individuals of host $n$ are sufficiently stronger competitors than susceptible individuals and host $n$ has stronger competitive effects on source hosts than sink hosts or (ii) infected individuals of host $n$ are weaker or insufficiently stronger competitors than susceptible individuals and host $n$ has stronger competitive effects on sink hosts than source hosts.

- Increasing the recovery rate in host $n$ ($\nu_n$) often decreases host 1 infection prevalence. However, host 1 infection infection prevalence can increase under the following conditions, with condition (2.i) being the most likely to arise in natural systems. (1) Host $n$ is a sink ($\varphi_n < 0$) and infected individuals of host $n$ have much larger uptake rates than susceptible individuals ($u_{I_1} > u_{S_i}$). (2) Host $n$ is a source ($\varphi_n > 0$) and either (i) infected individuals of host $n$ are sufficiently stronger competitors than susceptible individuals and host $n$ has stronger competitive effects on source hosts than sink hosts or (ii) infected individuals of host $n$ are weaker or insufficiently stronger competitors than susceptible individuals and host $n$ has stronger competitive effects on sink hosts than source hosts.

The predicted effects of varying the components defining the competence of host $n$ on host 1 infected density are often the same as the predicted effects for host 1 infection prevalence. However, the predictions for host 1 infected density can have the opposite sign when (i) for each host species, regulation by competition with host 1 is sufficiently weaker than regulation by disease and competition with species $2, ..., n$ or (ii) host 1 is a sink or small source and a subset of the other host species are sufficiently large sources. Condition (ii) only applies to the predictions for $\beta_i$, $\mu_i$ and $\nu_i$. Condition (i) is easier to satisfy in systems where infected individuals are stronger competitors than susceptible individuals and condition (ii) is easier to satisfy in systems where infected individuals have larger uptake rates than susceptible individuals. See Remark S1.6 for additional details.

**Shedding rates ($\chi_i$)**: The sensitivity of host 1 infection prevalence to the shedding rate of host $i$ is

$$\frac{\partial Y_1^*}{\partial \chi_i} = \left(-\frac{\partial}{\partial \chi_i}\frac{dP}{dt}\right)\frac{(-1)^{2n+n+2}}{|\mathbf{J}|}|\mathbf{J}^{[\cancel{2n+1},\cancel{n+1}]}| \tag{S83}$$

$$= \left(-\frac{\partial}{\partial \chi_i}\frac{dP}{dt}\right)\frac{-1}{|\mathbf{J}|}\begin{vmatrix} \mathbf{A} & \vec{0}_n & (\mathbf{B}+\mathbf{C}\mathbf{D}_\Gamma)^{[-,\cancel{1}]} \\ \mathbf{0}_{n\times n} & \begin{bmatrix} \beta_1 X_1^* \\ \vec{0}_{n-1} \end{bmatrix} & -\mathbf{D}_{\beta P/Y}^{[-,\cancel{1}]} \end{vmatrix} \tag{S84}$$

$$= \overbrace{Y_i^* N_i^*}^{\chi_i \to P}\overbrace{\beta_1 X_1^*}^{P \to Y_1}(-1)^{n-1}\prod_{j>1}\left(\frac{\beta_i P}{Y_i}\right)\frac{|\mathbf{A}|}{|\mathbf{J}|}. \tag{S85}$$

Equation (S85) represents the indirect effect $\chi_i \to P \to Y_1$. Because I assume sign($|\mathbf{J}|$) = $(-1)^{2n+1}$ and sign($|\mathbf{A}|$) = $(-1)^n$ (see Remark S1.2), equation (S85) is positive. Hence, increases in shedding always lead to increased host 1 infection prevalence.

The sensitivity of the host 1 total density to the shedding rate of host $i$ is

$$\frac{\partial N_1^*}{\partial \chi_i} = \left(-\frac{\partial}{\partial \chi_i}\frac{dP}{dt}\right)\frac{(-1)^{2n+2}}{|\mathbf{J}|}|\mathbf{J}^{[\cancel{2n+1},\cancel{1}]}| \tag{S86}$$

$$= \left(-\frac{\partial}{\partial \chi_i}\frac{dP}{dt}\right)\frac{1}{|\mathbf{J}|}\begin{vmatrix} \mathbf{A}^{[-,\cancel{1}]} & \mathbf{B}+\mathbf{C}\mathbf{D}_\Gamma & \vec{0}_n \\ \mathbf{0}_{n\times n-1} & -\mathbf{D}_{\beta P/Y} & \overrightarrow{\beta X} \end{vmatrix} \tag{S87}$$

$$= \left(-\frac{\partial}{\partial \chi_i}\frac{dP}{dt}\right)\frac{(-1)^{2n-1}}{|\mathbf{J}|}\begin{vmatrix} \vec{0}_n & \mathbf{A}^{[-,\cancel{1}]} & \mathbf{B}+\mathbf{C}\mathbf{D}_\Gamma \\ \overrightarrow{\beta X} & \mathbf{0}_{n\times n-1} & -\mathbf{D}_{\beta P/Y} \end{vmatrix} \tag{S88}$$

$$= \left(-\frac{\partial}{\partial \chi_i}\frac{dP}{dt}\right)\frac{(-1)}{|\mathbf{J}|}\begin{vmatrix} -\vec{\omega}_n/P^* & \mathbf{A}^{[-,\cancel{1}]} & \mathbf{B}+\mathbf{C}\mathbf{D}_\Gamma \\ \vec{0}_n & \mathbf{0}_{n\times n-1} & -\mathbf{D}_{\beta P/Y} \end{vmatrix} \tag{S89}$$

$$= Y_i^* N_i^* \frac{(-1)^{n-1}}{P^*|\mathbf{J}|}\prod_{j\geq 1}\left(\frac{\beta_i P^*}{Y_i}\right)\left|\left[\begin{array}{cc} \vec{\omega}_n & \mathbf{A}^{[-,\cancel{1}]} \end{array}\right]\right|. \tag{S90}$$

Using equation (S65) yields

$$\frac{\partial I_1^*}{\partial \chi_i} = Y_i^* N_i^* \frac{(-1)^{n-1}\beta_1}{|\mathbf{J}|}\prod_{j>1}\left(\frac{\beta_i P^*}{Y_i}\right)\left(S_1^*|\mathbf{A}| + \left|\left[\begin{array}{cc} \vec{\omega}_n & \mathbf{A}^{[-,\cancel{1}]} \end{array}\right]\right|\right) \tag{S91}$$

Equation (S85) for $\partial Y_1^*/\partial \chi_i$ and equation (S91) for $\partial I_1^*/\partial \chi_i$ have opposite signs whenever the second determinant in equation (S91) has the opposite sign of the first determinant and is sufficiently large in magnitude. That is more likely to occur when the entries of $\vec{\omega}_n$ are positive and large in magnitude relative to the entries of $\mathbf{A}_{\bullet 1}$. Those conditions are discussed in Remark S1.6.

**Uptake rates ($u_{V_i}$)**: Nearly identical calculations yield the sensitivities to uptake rates,

$$\frac{\partial Y_1^*}{\partial u_{V_i}} = \overbrace{V_i^* P^*}^{u_{V_i}\to P}\overbrace{\beta_1 S_1^*}^{P\to Y_1}(-1)^n \prod_{j>1}\left(\frac{\beta_i P^*}{Y_i}\right)\frac{|\mathbf{A}|}{|\mathbf{J}|}. \tag{S92}$$

$$\frac{\partial I_1^*}{\partial u_{V_i}} = V_i^* P^* \frac{(-1)^n \beta_1}{|\mathbf{J}|}\prod_{j>1}\left(\frac{\beta_i P^*}{Y_i}\right)\left(S_1^*|\mathbf{A}| + \left|\begin{bmatrix} \vec{\omega}_n & \mathbf{A}^{[-,\cancel{1}]} \end{bmatrix}\right|\right). \tag{S93}$$

Equation (S92) represents the indirect effect $u_{v_i} \to P \to Y_1$. Because I assume sign($|\mathbf{J}|$) = $(-1)^{2n+1}$ and sign($|\mathbf{A}|$) = $(-1)^n$ (see Remark S1.2), equation (S85) is negative. Hence, increases in uptake always lead to decreased host 1 infection prevalence.

Equation (S92) for $\partial Y_1^*/\partial u_{V_i}$ and equation (S93) for $\partial I_1^*/\partial u_{V_i}$ have opposite signs whenever the second determinant in equation (S93) has the opposite sign of the first determinant and is sufficiently large in magnitude. That is more likely to occur when the entries of $\vec{\omega}_n$ are positive and large in magnitude relative to the entries of $\mathbf{A}_{\bullet 1}$. Those conditions are discussed in Remark S1.6.

**Transmission parameter ($\beta_n$)**: The sensitivity of host 1 infection prevalence to the transmission parameter for host $n$ is

$$\frac{\partial Y_1^*}{\partial \beta_n} = \left(-\frac{\partial}{\partial \beta_n}\frac{dY_n}{dt}\right)\frac{(-1)^{2n+n+1}}{|\mathbf{J}|}|\mathbf{J}^{[\cancel{2n},\cancel{n+1}]}| \tag{S94}$$

$$= \left(-\frac{\partial}{\partial \beta_n}\frac{dY_n}{dt}\right)\frac{1}{|\mathbf{J}|}\begin{vmatrix} \mathbf{A} & \mathbf{B}_{\bullet n} + \Gamma_n \mathbf{C}_{\bullet n} & \vec{0}_n & (\mathbf{B}+\mathbf{C}\mathbf{D}_\Gamma)^{[-,\cancel{1-n}]} \\ \vec{E}_N & \vec{E}_Y[n] & 0 & \vec{E}_Y^{[\cancel{1-n}]} \\ \mathbf{0}_{n-1\times n} & \vec{0}_{n-1} & \begin{bmatrix} \beta_1 X_1^* \\ \vec{0}_{n-2} \end{bmatrix} & -\mathbf{D}_{\beta P/Y}^{[\cancel{n},\cancel{1}]} \end{vmatrix} \tag{S95}$$

$$= \frac{(-1)^{n-1} X_n^* P^* \beta_1 X_1^*}{|\mathbf{J}|}\prod_{j=2}^{n-1}\left(\frac{\beta_j N_j^* P^*}{I_j^*}\right)\left|\begin{bmatrix} \mathbf{A} & \mathbf{B}_{\bullet n} + \Gamma_n \mathbf{C}_{\bullet n} \\ \vec{E}_N & \vec{E}_Y[n] \end{bmatrix}\right| \tag{S96}$$

$$\begin{aligned} &= \overbrace{X_n^* P^*}^{\beta_n \to Y_n}\overbrace{\frac{N_n^*}{Y_n^*}(\varphi_n Y_n^* + u_{S_n} P^*)}^{Y_n \to P}\overbrace{\beta_1 X_1^*}^{P\to Y_1}\frac{|\mathbf{A}|(-1)^{n-1}}{|\mathbf{J}|}\prod_{j=2}^{n-1}\left(\frac{\beta_j P^*}{Y_j^*}\right) \\ &\quad - \overbrace{X_n^* P^*}^{\beta_n \to Y_n}\overbrace{\sum_{j=1}^{n}\varphi_j Y_j^* |\mathbf{A}_{I_n \to N_j}|}^{Y_n \to N_j \to P}\overbrace{\beta_1 X_1^*}^{P\to Y_1}\frac{(-1)^{n-1}}{|\mathbf{J}|}\prod_{j=2}^{n-1}\left(\frac{\beta_j P^*}{Y_j^*}\right). \end{aligned} \tag{S97}$$

The first line of equation (S97) represents the indirect effect $\beta_n \to Y_n \to P \to Y_1$. Because I assume sign($|\mathbf{J}|$) = $(-1)^{2n+1}$ and sign($|\mathbf{A}|$) = $(-1)^n$ (see Remark S1.2), the sign of this indirect is determined by $\varphi_n Y_n^* + u_{S_n} P^*$. Consequently, this indirect effect is positive unless host $n$ is a sink ($\varphi_n < 0$) and infected individuals of host $n$ have much larger uptake rates than susceptible individuals ($u_{I_i}$ is much larger than $u_{S_i}$).

The second line of equation (S97) defines how changes in the number of infected individuals of host $n$ affects the densities of all host species, represented by the sum of the indirect effects $\beta_n \to Y_n \to N_j \to P \to Y_1$. The sign of the $n$th term is $-\varphi_n$ because I assume $\text{sign}(|\mathbf{J}|) = (-1)^{2n+1}$ and $\text{sign}(|\mathbf{A}_{I_n \to N_n}|) = (-1)^n$ (see Remark S1.5). Consequently, the $n$th term of the sum is positive when host $n$ is a sink ($\varphi_n < 0$) and negative when host $n$ is a source ($\varphi_n > 0$). For the other terms, the sign of each indirect effect depends on the competitive ability of infected individuals of host $n$ and whether the other hosts are sinks or sources. If infected individuals are sufficiently stronger interspecific competitors than susceptible individuals, then Remark S1.5 implies $\text{sign}(|\mathbf{A}_{I_n \to N_j}|) = (-1)^n$ and the sign of each term is $-\varphi_i$; this means each term is positive for sink hosts and negative for source hosts. Otherwise (i.e., infected individuals are weaker or insufficiently stronger interspecific competitors than susceptible individuals), Remark S1.5 implies $\text{sign}(|\mathbf{A}_{I_n \to N_j}|) = (-1)^{n-1}$ and the sign of each term is $\varphi_i$; this means each term is positive for source hosts and negative for sink hosts. Note that all terms in the sum are larger in magnitude when interspecific competition is stronger (because $\text{sign}(|\mathbf{A}_{I_n \to N_j}|) = 0$ in the absence of interspecific competition; see Remark S1.5).

Combining the above, equation (S97) is often positive. However, equation (S97) can be negative if one of the following conditions is satisfied. (1) Host $n$ is a sink ($\varphi_n < 0$) and infected individuals of host $n$ have much larger uptake rates than susceptible individuals ($u_{I_1} > u_{S_i}$); this makes the first line of equation (S97) negative. (2) Host $n$ is a source ($\varphi_i > 0$) and interspecific competition is sufficiently strong; this makes the $n$th term of the sum on the second line of equation (S97) negative. (3) Infected individuals of host $n$ are sufficiently stronger interspecific competitors than susceptible individuals and host $n$ has stronger interspecific competitive effects on source hosts ($\varphi_j > 0$) than sink hosts ($\varphi_j < 0$); this results in the second line of equation (S97) having larger negative terms for each source host and smaller positive terms for each sink host. (4) Infected individuals of host $n$ are weaker or insufficiently stronger interspecific competitors than susceptible individuals and host $n$ has stronger interspecific competitive effects on sink hosts ($\varphi_j < 0$) than source hosts ($\varphi_j > 0$); this results in the second line of equation (S97) having larger negative terms for each sink host and smaller positive terms for each source host.

The sensitivity of host 1 total density to the transmission parameter for host $n$ is

$$\frac{\partial N_1^*}{\partial \beta_n} = \left(-\frac{\partial}{\partial \beta_n}\frac{dY_n}{dt}\right)\frac{(-1)^{2n+1}}{|\mathbf{J}|}|\mathbf{J}^{[\cancel{2n},\cancel{1}]}| \tag{S98}$$

$$= \frac{X_n^* P^*}{|\mathbf{J}|}\begin{vmatrix} \mathbf{A}^{[\cancel{1},-]} & \mathbf{B}+\mathbf{C}\mathbf{D}_\Gamma & \vec{0}_n \\ \mathbf{0}_{n-1\times n-1} & -\mathbf{D}_{\beta P/Y}^{[\cancel{n},-]} & \overrightarrow{\beta X}^{[\cancel{n}]} \\ \vec{E}_N^{[\cancel{1}]} & \vec{E}_Y & -U-\delta \end{vmatrix} \tag{S99}$$

$$= \frac{X_n^* P^* (-1)^{n-1+2n-1}}{|\mathbf{J}|}\begin{vmatrix} \vec{0}_n & \mathbf{A}^{[\cancel{1},-]} & \mathbf{B}+\mathbf{C}\mathbf{D}_\Gamma \\ -U-\delta & \vec{E}_N^{[\cancel{1}]} & \vec{E}_Y \\ \overrightarrow{\beta X}^{[\cancel{n}]} & \mathbf{0}_{n-1\times n-1} & -\mathbf{D}_{\beta P/Y}^{[\cancel{n},-]} \end{vmatrix} \tag{S100}$$

$$= \frac{X_n^* P^* (-1)^{n-1+2n-1+n-1}}{|\mathbf{J}|}\begin{vmatrix} -\omega_{n-1}/P^* & \mathbf{A}^{[\cancel{1},-]} & \mathbf{B}_{\bullet n}+\Gamma_n\mathbf{C}_{\bullet n} & (\mathbf{B}+\mathbf{C}\mathbf{D}_\Gamma)^{[-,\cancel{n}]} \\ -\theta_{n-1}/P^* & \vec{E}_N^{[\cancel{1}]} & \vec{E}_Y[n] & \vec{E}_Y^{[\cancel{n}]} \\ \vec{0}_{n-1} & \mathbf{0}_{n-1\times n-1} & 0 & -\mathbf{D}_{\beta P/Y}^{[\cancel{n},\cancel{n}]} \end{vmatrix} \tag{S101}$$

$$= \frac{X_n^* P^* (-1)^{n-1}}{|\mathbf{J}|} \prod_{j<n} \left( \frac{\beta_j P^*}{Y_j^*} \right) \begin{vmatrix} \omega_{n-1}/P^* & \mathbf{A}^{[\cancel{1},-]} & \mathbf{B}_{\bullet n} + \Gamma_n \mathbf{C}_{\bullet n} \\ \theta_{n-1}/P^* & \vec{E}_N^{[\cancel{1}]} & \vec{E}_Y[n] \end{vmatrix} \tag{S102}$$

Using equation (S65) yields

$$\frac{\partial I_1^*}{\partial \beta_n} = \frac{X_n^* P^* (-1)^{n-1}}{|\mathbf{J}|} \prod_{j=2}^{n-1} \left( \frac{\beta_j P^*}{Y_j^*} \right) \left( S_1^* \begin{vmatrix} \mathbf{A} & \mathbf{B}_{\bullet n} + \Gamma_n \mathbf{C}_{\bullet n} \\ \vec{E}_N & \vec{E}_Y[n] \end{vmatrix} + \begin{vmatrix} \omega_{n-1} & \mathbf{A}^{[\cancel{1},-]} & \mathbf{B}_{\bullet n} + \Gamma_n \mathbf{C}_{\bullet n} \\ \theta_{n-1} & \vec{E}_N^{[\cancel{1}]} & \vec{E}_Y[n] \end{vmatrix} \right) \tag{S103}$$

Equation (S97) for $\partial Y_1^*/\partial \beta_n$ and equation (S103) for $\partial I_1^*/\partial \beta_n$ have opposite signs whenever the second determinant in equation (S103) has the opposite sign of the first determinant and is sufficiently large in magnitude. This is more likely to occur when (i) the entries of $\vec{\omega}_{n-1}$ are positive and large in magnitude relative to the entries of $\mathbf{A}_{\bullet 1}$ or (ii) $\theta_{n-1}$ is positive and large in magnitude and $\mathbf{E}_N[1]$ is either negative or positive and small in magnitude. Those conditions are discussed in Remark S1.6.

**Disease-induced mortality rate ($\mu_n$)**:The sensitivity of host 1 infection prevalence to the disease-induced mortality rate for host $n$ is

$$\frac{\partial Y_1^*}{\partial \mu_n} = \left( -\frac{\partial}{\partial \mu_n} \frac{dN_n}{dt} \right) \frac{(-1)^{n+n+1}}{|\mathbf{J}|} |\mathbf{J}^{[\cancel{n},\cancel{n+1}]}| + \left( -\frac{\partial}{\partial \mu_n} \frac{dY_n}{dt} \right) \frac{(-1)^{2n+n+1}}{|\mathbf{J}|} |\mathbf{J}^{[\cancel{2n},\cancel{n+1}]}| \tag{S104}$$

$$= \left( -\frac{\partial}{\partial \mu_n} \frac{dN_n}{dt} \right) \frac{1}{|\mathbf{J}|} \begin{vmatrix} \mathbf{A}^{[\cancel{n},-]} & \vec{0}_{n-1} & (\mathbf{B} + \mathbf{C}\mathbf{D}_\Gamma)^{[\cancel{n},\cancel{1}]} \\ \vec{E}_N & 0 & \vec{E}_Y^{\cancel{n}} \\ \mathbf{0}_{n\times n} & \begin{bmatrix} \beta_1 X_1^* \\ \vec{0}_{n-1} \end{bmatrix} & -\mathbf{D}_{\beta P/Y}^{[-,\cancel{1}]} \end{vmatrix} \tag{S105}$$

$$+ Y_n^* \frac{1}{|\mathbf{J}|} \begin{vmatrix} \mathbf{A} & \mathbf{B}_{\bullet n} + \Gamma_n \mathbf{C}_{\bullet n} & \vec{0}_n & (\mathbf{B} + \mathbf{C}\mathbf{D}_\Gamma)^{[-,\cancel{1\,n}]} \\ \vec{E}_N & \vec{E}_Y[n] & 0 & \vec{E}_Y^{\cancel{1\,n}} \\ \mathbf{0}_{n\times n} & \vec{0}_n & \begin{bmatrix} \beta_1 X_1^* \\ \vec{0}_{n-1} \end{bmatrix} & -\mathbf{D}_{\beta P/Y}^{[\cancel{n},\cancel{1n}]} \end{vmatrix} \tag{S106}$$

$$= Y_n^* N_n^* \frac{(-1)^{n-1}}{|\mathbf{J}|} \beta_1 X_1^* \prod_{j>1} \left( \frac{\beta_i P^*}{Y_i^*} \right) \begin{vmatrix} \mathbf{A}^{[\cancel{n},-]} \\ \vec{E}_N \end{vmatrix} \tag{S107}$$

$$+ Y_n^* \frac{(-1)^n}{|\mathbf{J}|} \beta_1 X_1^* \prod_{j=2}^{n-1} \left( \frac{\beta_i P^*}{Y_i^*} \right) \begin{vmatrix} \mathbf{A} & \mathbf{B}_{\bullet n} + \Gamma_n \mathbf{C}_{\bullet n} \\ \vec{E}_N & \vec{E}_Y[n] \end{vmatrix} \tag{S108}$$

$$= \overbrace{-Y_n^* N_n^*}^{\mu_n \to N_n} \overbrace{\sum_j (-1)^j |\mathbf{A}^{[\cancel{n},\cancel{j}]}| \varphi_j Y_j^*}^{N_n \to N_k \to P} \overbrace{\beta_1 X_1^*}^{P \to Y_1} \frac{1}{|\mathbf{J}|} \prod_{j>1} \left( \frac{\beta_i P^*}{Y_i^*} \right)$$

$$+ \overbrace{Y_n^*}^{\mu_n \to Y_n} \overbrace{\frac{N_n^*}{Y_n^*} (\varphi_n Y_n^* + u_{S_n} P^*) |\mathbf{A}|}^{Y_n \to P} \overbrace{\beta_1 X_1^*}^{P \to Y_1} \frac{(-1)^n}{|\mathbf{J}|} \prod_{j=2}^{n-1} \left( \frac{\beta_j P^*}{Y_j^*} \right) \tag{S109}$$

$$- \overbrace{Y_n^*}^{\mu_n \to Y_n} \overbrace{\sum_{j=1}^{n} \varphi_j Y_j^* |\mathbf{A}_{I_n \to N_j}|}^{Y_n \to N_j \to P} \overbrace{\beta_1 X_1^*}^{P \to Y_1} \frac{(-1)^n}{|\mathbf{J}|} \prod_{j=2}^{n-1} \left( \frac{\beta_j P^*}{Y_j^*} \right).$$

The first line of equation (S109) defines how changes in density of host $n$ affect the densities of all host species, represented as the indirect effect $\mu_n \to N_n \to P \to Y_1$ and the indirect effects $\mu_n \to N_n \to N_k \to P \to Y_1$ $(k \neq n)$. The sign of the $n$th term is $-\varphi_n$, because I assume sign$(|\mathbf{J}|) = (-1)^{2n+1}$ and sign$(|\mathbf{A}^{[\not{n},\not{n}]}|) = (-1)^{n-1}$ (see Remark S1.2). Consequently, the $n$th term of the sum is positive when host $n$ is a sink $(\varphi_n < 0)$ and negative when host $n$ is a source $(\varphi_n > 0)$. For the other terms, the sign of each indirect effect is determined by the sign of $(-1)^{j+1}|\mathbf{A}^{[\not{n},\not{j}]}|\varphi_j/|\mathbf{J}|$, which has the same sign as $\alpha_{jn}^{total}\varphi_j$ (where Remark $\alpha_{jn}^{total}$ is the total competitive effect of host $n$ on host $j$; see S1.3). This means that the sum of terms 1 through $n-1$ is negative unless host $n$ has stronger negative (or weaker positive) total competitive effects on sink hosts and weaker negative (or stronger positive) total competitive effects on source hosts.

The second line of equation (S97) defines how changes in proportion of infected individuals in host $n$ affects infectious propagule density, represented as the indirect effect $\mu_n \to Y_n \to P \to Y_1$. The sign of this indirect effect is $-(\varphi_n Y_n^* + u_{S_n} P^*)$ because I assume sign$(|\mathbf{J}|) = (-1)^{2n+1}$ and sign$(|\mathbf{A}|) = (-1)^n$ (see Remark S1.2). Consequently, this indirect effect is negative unless host $n$ is a sink $(\varphi_n < 0)$ and infected individuals of host $n$ have much larger uptake rates than susceptible individuals $(u_{I_1} > u_{S_i})$.

The third line of equation (S109) defines how changes in proportion of infected individuals in host $n$ affects the densities of all host species, represented as the indirect effects $\mu_n \to Y_n \to N_j \to P \to Y_1$. The sign of the $n$th term is $\varphi_n$ because I assume sign$(|\mathbf{J}|) = (-1)^{2n+1}$ and sign$(|\mathbf{A}_{I_n \to N_n}|) = (-1)^n$ (see Remark S1.5). Thus, the indirect effect is positive when host $n$ is a source $(\varphi_n > 0)$ and negative when host is a sink $(\varphi_n < 0)$. For the other terms, the sign of each indirect effect depends on the competitive ability of infected individuals of host $n$ and whether the other host species are sinks or sources. If infected individuals are sufficiently stronger interspecific competitors than susceptible individuals, then Remark S1.5 implies sign$(|\mathbf{A}_{I_n \to N_j}|) = (-1)^n$ and the sign of each term is $\varphi_j$; this means each term is positive for source hosts $(\varphi_j > 0)$ and negative for sink hosts $(\varphi_j < 0)$. Otherwise (i.e., infected individuals are weaker or insufficiently stronger interspecific competitors than susceptible individuals), Remark S1.5 implies sign$(|\mathbf{A}_{I_n \to N_j}|) = (-1)^{n-1}$ and the sign of each term is $-\varphi_j$; this means each term is positive for sink hosts $(\varphi_j < 0)$ and negative for source hosts $(\varphi_j > 0)$. Note that all terms in the sum are larger in magnitude when interspecific competition is stronger (because sign$(|\mathbf{A}_{I_n \to N_j}|) = 0$ in the absence of interspecific competition; see Remark S1.5).

Combining the above, equation (S109) is often negative. However, equation (S109) can be negative if one of the following conditions is satisfied. (1) Host $n$ is a sufficiently large sink $(\varphi_n < 0)$. This makes the $n$th term of the sum in the first line of equation (S109) positive. (2) Host $n$ has stronger negative (or weaker positive) total competitive effects on sink hosts and weaker negative (or stronger positive) total competitive effects on source hosts. This results in terms 1 through $n-1$ of the sum in the first line of equation (S109) have larger positive terms for sink hosts and smaller negative terms for source hosts. (3) Host $n$ is a sink $(\varphi_n < 0)$ and infected individuals of host $n$ have much larger uptake rates than susceptible individuals $(u_{I_1} > u_{S_i})$. This makes the second line of equation (S109) positive. (4) Host $n$ is a source $(\varphi_n > 0)$ and either (i) infected individuals of host $n$ are sufficiently stronger competitors than susceptible individuals and host $n$ has stronger competitive effects on source hosts than sink hosts or (ii) Infected individuals of host $n$ are weaker or insufficiently stronger competitors than susceptible individuals and host $n$ has stronger competitive effects on sink hosts than source hosts. This makes the third line of equation (S109) positive. In prior work (Cortez 2021), I was unable to find examples satisfying conditions (4.i) and (4.ii), so I expect them to be rarely satisfied in natural systems.

The sensitivity of the host 1 total density to the disease-induced mortality rate of host $n$ is

$$\frac{\partial N_1^*}{\partial \mu_n} = \left(-\frac{\partial}{\partial \mu_n}\frac{dN_n}{dt}\right)\frac{(-1)^{n+1}}{|\mathbf{J}|}|\mathbf{J}^{[\cancel{n},\cancel{1}]}| + \left(-\frac{\partial}{\partial \mu_n}\frac{dI_n}{dt}\right)\frac{(-1)^{2n+1}}{|\mathbf{J}|}|\mathbf{J}^{[\cancel{2n},\cancel{1}]}| \tag{S110}$$

$$= \left(-\frac{\partial}{\partial \mu_n}\frac{dN_n}{dt}\right)\frac{(-1)^{n+1}}{|\mathbf{J}|}\begin{vmatrix} \mathbf{A}^{[\cancel{n},1]} & (\mathbf{B}+\mathbf{C}\mathbf{D}_\Gamma)^{[\cancel{n},-]} & \vec{0}_{n-1} \\ \mathbf{0}_{n\times n-1} & -\mathbf{D}_{\beta P/Y} & \overrightarrow{\beta X} \\ \vec{E}_N^{[\cancel{1}]} & \vec{E}_Y & -U-\delta \end{vmatrix} \tag{S111}$$

$$+ Y_n^*\frac{(-1)}{|\mathbf{J}|}\begin{vmatrix} \mathbf{A}^{[-,1]} & \mathbf{B}+\mathbf{C}\mathbf{D}_\Gamma & \vec{0}_{n-1} \\ \mathbf{0}_{n-1\times n-1} & -\mathbf{D}_{\beta P/Y}^{[\cancel{n},-]} & \overrightarrow{\beta X}^{[\cancel{n}]} \\ \vec{E}_N^{[\cancel{1}]} & \vec{E}_Y & -U-\delta \end{vmatrix} \tag{S112}$$

$$\begin{aligned} =& Y_n^* N_n^* \frac{(-1)^{n-1}}{P^*|\mathbf{J}|}\prod_{j\geq 1}\left(\frac{\beta_i P^*}{Y_i^*}\right)\begin{vmatrix} \vec{\omega}_n^{[\cancel{n}]} & \mathbf{A}^{[\cancel{n},\cancel{1}]} \\ \theta_n & \vec{E}_N^{[\cancel{1}]} \end{vmatrix} \\ &+ \frac{Y_n^*(-1)^n}{|\mathbf{J}|}\prod_{j<n}\left(\frac{\beta_j P^*}{Y_j^*}\right)\begin{vmatrix} \omega_{n-1} & \mathbf{A}^{[\cancel{1},-]} & \mathbf{B}_{\bullet n}+\Gamma_n\mathbf{C}_{\bullet n} \\ \theta_{n-1} & \vec{E}_N^{[\cancel{1}]} & \vec{E}_Y[n] \end{vmatrix} \end{aligned} \tag{S113}$$

Using equation (S65) yields

$$\begin{aligned} \frac{\partial I_1^*}{\partial \mu_n} =& Y_n^* N_n^* \frac{(-1)^{n-1}}{P^*|\mathbf{J}|}\prod_{j\geq 2}\left(\frac{\beta_i P^*}{Y_i^*}\right)\left(S_1^*\begin{vmatrix} \mathbf{A}^{[\cancel{n},-]} \\ \vec{E}_N \end{vmatrix} + \begin{vmatrix} \vec{\omega}_n^{[\cancel{n}]} & \mathbf{A}^{[\cancel{n},\cancel{1}]} \\ \theta_n & \vec{E}_N^{[\cancel{1}]} \end{vmatrix}\right) \\ &+ \frac{Y_n^*(-1)^n}{|\mathbf{J}|}\prod_{j=2}^{n-1}\left(\frac{\beta_j P^*}{Y_j^*}\right)\left(S_1^*\begin{vmatrix} \mathbf{A} & \mathbf{B}_{\bullet n}+\Gamma_n\mathbf{C}_{\bullet n} \\ \vec{E}_N & \vec{E}_Y[n] \end{vmatrix} + \begin{vmatrix} \omega_{n-1} & \mathbf{A}^{[\cancel{1},-]} & \mathbf{B}_{\bullet n}+\Gamma_n\mathbf{C}_{\bullet n} \\ \theta_{n-1} & \vec{E}_N^{[\cancel{1}]} & \vec{E}_Y[n] \end{vmatrix}\right) \end{aligned} \tag{S114}$$

Equation (S109) for $\partial Y_1^*/\partial \mu_n$ and equation (S114) for $\partial I_1^*/\partial \mu_n$ have opposite signs whenever the second determinant in equation (S114) has the opposite sign of the first determinant and is sufficiently large in magnitude. This is more likely to occur when (i) the entries of $\vec{\omega}_n$ and $\vec{\omega}_{n-1}$ are positive and large in magnitude relative to the entries of $\mathbf{A}_{\bullet 1}$ or (ii) $\theta_n$ and $\theta_{n-1}$ are positive and large in magnitude and $\mathbf{E}_N[1]$ is either negative or positive and small in magnitude. Those conditions are discussed in Remark S1.6.

**Recovery rate ($\nu_n$)**: The sensitivity of host 1 infection prevalence to the recovery rate for host $n$ is

$$\frac{\partial Y_1^*}{\partial \nu_n} = \left(-\frac{\partial}{\partial \nu_n}\frac{dY_n}{dt}\right)\frac{(-1)^{2n+n+1}}{|\mathbf{J}|}|\mathbf{J}^{[\cancel{2n},\cancel{n+1}]}| \tag{S115}$$

$$
= \left(-\frac{\partial}{\partial \nu_n}\frac{dY_n}{dt}\right)\frac{(-1)}{|\mathbf{J}|}\begin{vmatrix} \mathbf{A} & \mathbf{B}_{\bullet n} + \Gamma_n \mathbf{C}_{\bullet n} & \vec{0}_n & (\mathbf{B}+\mathbf{C}\mathbf{D}_\Gamma)^{[-,\not{1}\not{n}]} \\ \vec{E}_N & \vec{E}_Y[n] & 0 & \vec{E}_Y^{\not{1}\not{n}} \\ \mathbf{0}_{n\times n} & \vec{0}_n & \begin{bmatrix} \beta_1 X_1^* \\ \vec{0}_{n-1} \end{bmatrix} & -\mathbf{D}_{\beta P/Y}^{[-,\not{1}]} \end{vmatrix} \tag{S116}
$$

$$
\begin{aligned}
= & \overbrace{Y_n^*}^{\nu_n \to Y_n} \overbrace{\frac{N_n^*}{Y_n^*}(\varphi_n Y_n^* + u_{S_n} P^*)|\mathbf{A}|}^{Y_n \to P} \overbrace{\beta_1 X_1^*}^{P \to Y_1} \frac{(-1)^n}{|\mathbf{J}|} \prod_{j=2}^{n-1}\left(\frac{\beta_j P^*}{Y_j^*}\right) \\
& - \overbrace{Y_n^*}^{\nu_n \to Y_n} \overbrace{\sum_{j=1}^{n} \varphi_j Y_j^* |\mathbf{A}_{I_n \to N_j}|}^{Y_n \to N_j \to P} \overbrace{\beta_1 X_1^*}^{P \to Y_1} \frac{(-1)^n}{|\mathbf{J}|} \prod_{j=2}^{n-1}\left(\frac{\beta_j P^*}{Y_j^*}\right).
\end{aligned} \tag{S117}
$$

The first line of equation (S117) defines how changes in proportion of infected individuals in host $n$ affects infectious propagule density, represented as the indirect effect $\nu_n \to Y_n \to P \to Y_1$. The sign of this indirect effect is $-(\varphi_n Y_n^* + u_{S_n} P^*)$ because I assume $\text{sign}(|\mathbf{J}|) = (-1)^{2n+1}$ and $\text{sign}(|\mathbf{A}|) = (-1)^n$ (see Remark S1.2). Consequently, this indirect effect is negative unless host $n$ is a sink ($\varphi_n < 0$) and infected individuals of host $n$ have much larger uptake rates than susceptible individuals ($u_{I_1} > u_{S_i}$).

The second line of equation (S117) defines how changes in proportion of infected individuals in host $n$ affects the densities of all host species, represented as the indirect effects $\nu_n \to Y_n \to N_j \to P \to Y_1$. The sign of the $n$th term is $\varphi_n$ because I assume $\text{sign}(|\mathbf{J}|) = (-1)^{2n+1}$ and $\text{sign}(|\mathbf{A}_{I_n \to N_n}|) = (-1)^n$ (see Remark S1.5). Thus, the indirect effect is positive when host $n$ is a source ($\varphi_n > 0$) and negative when host is a sink ($\varphi_n < 0$). For the other terms, the sign of each indirect effect depends on the competitive ability of infected individuals of host $n$ and whether the other host species are sinks or sources. If infected individuals are sufficiently stronger interspecific competitors than susceptible individuals, then Remark S1.5 implies $\text{sign}(|\mathbf{A}_{I_n \to N_j}|) = (-1)^n$ and the sign of each term is $\varphi_j$; this means each term is positive for source hosts ($\varphi_j > 0$) and negative for sink hosts ($\varphi_j < 0$). Otherwise (i.e., infected individuals are weaker or insufficiently stronger interspecific competitors than susceptible individuals), Remark S1.5 implies $\text{sign}(|\mathbf{A}_{I_n \to N_j}|) = (-1)^{n-1}$ and the sign of each term is $-\varphi_j$; this means each term is positive for sink hosts ($\varphi_j < 0$) and negative for source hosts ($\varphi_j > 0$). Note that all terms in the sum are larger in magnitude when interspecific competition is stronger (because $\text{sign}(|\mathbf{A}_{I_n \to N_j}|) = 0$ in the absence of interspecific competition; see Remark S1.5).

Combining the above, equation (S117) is often negative. However, equation (S117) can be negative if one of the following conditions is satisfied. (1) Host $n$ is a sink ($\varphi_n < 0$) and infected individuals of host $n$ have much larger uptake rates than susceptible individuals ($u_{I_1} > u_{S_i}$). This makes the second line of equation (S117) positive. (2) Host $n$ is a source ($\varphi_n > 0$) and either (i) infected individuals of host $n$ are sufficiently stronger competitors than susceptible individuals and host $n$ has stronger competitive effects on source hosts than sink hosts or (ii) Infected individuals of host $n$ are weaker or insufficiently stronger competitors than susceptible individuals and host $n$ has stronger competitive effects on sink hosts than source hosts. This makes the third line of equation (S117) positive. In prior work (Cortez 2021), I was unable to find examples satisfying conditions (2.i) and (2.ii), so I expect them to be rarely satisfied in natural systems.

The sensitivity of the host 1 total density to the recovery rate of host $n$ is

$$\frac{\partial N_1^*}{\partial \nu_n} = \left( -\frac{\partial}{\partial \nu_n} \frac{dI_n}{dt} \right) \frac{(-1)^{2n+1}}{|\mathbf{J}|} |\mathbf{J}^{[\not{2n},\not{1}]}| \tag{S118}$$

$$= Y_n^* \frac{(-1)}{|\mathbf{J}|} \begin{vmatrix} \mathbf{A}^{[-,1]} & \mathbf{B} + \mathbf{CD}_\Gamma & \vec{0}_{n-1} \\ \mathbf{0}_{n-1 \times n-1} & -\mathbf{D}_{\beta P/Y}^{[\not{n},-]} & \overrightarrow{\beta X}^{[\not{n}]} \\ \vec{E}_N^{[\not{1}]} & \vec{E}_Y & -U - \delta \end{vmatrix} \tag{S119}$$

$$= \frac{Y_n^* (-1)^n}{|\mathbf{J}|} \prod_{j<n} \left( \frac{\beta_j P^*}{Y_j^*} \right) \begin{vmatrix} \omega_{n-1}/P^* & \mathbf{A}^{[\not{1},-]} & \mathbf{B}_{\bullet n} + \Gamma_n \mathbf{C}_{\bullet n} \\ \theta_{n-1}/P^* & \vec{E}_N^{[\not{1}]} & \vec{E}_Y[n] \end{vmatrix} \tag{S120}$$

Using equation (S65) yields

$$\frac{\partial I_1^*}{\partial \nu_n} = \frac{Y_n^* (-1)^n}{|\mathbf{J}|} \prod_{j=2}^{n-1} \left( \frac{\beta_j P^*}{Y_j^*} \right) \left( S_1^* \begin{vmatrix} \mathbf{A} & \mathbf{B}_{\bullet n} + \Gamma_n \mathbf{C}_{\bullet n} \\ \vec{E}_N & \vec{E}_Y[n] \end{vmatrix} + \begin{vmatrix} \omega_{n-1} & \mathbf{A}^{[\not{1},-]} & \mathbf{B}_{\bullet n} + \Gamma_n \mathbf{C}_{\bullet n} \\ \theta_{n-1} & \vec{E}_N^{[\not{1}]} & \vec{E}_Y[n] \end{vmatrix} \right) \tag{S121}$$

Equation (S117) for $\partial Y_1^*/\partial \nu_n$ and equation (S121) for $\partial I_1^*/\partial \nu_n$ have opposite signs whenever the second determinant in equation (S121) has the opposite sign of the first determinant and is sufficiently large in magnitude. This is more likely to occur when (i) the entries of $\vec{\omega}_{n-1}$ are positive and large in magnitude relative to the entries of $\mathbf{A}_{\bullet 1}$ or (ii) $\theta_{n-1}$ is positive and large in magnitude and $\mathbf{E}_N[1]$ is either negative or positive and small in magnitude. Those conditions are discussed in Remark S1.6.

##### S1.5.4 Sensitivities to transmission mode

Here, I use a continuous change of parameters to convert the 1-habitat ET model (S4) from a frequency-dependent direct transmission (FDDT) form ($\delta = 0$) to a density-dependent direct transmission form (DDDT) form ($u_{V_i} = 0$ for $V \in \{S, I, R\}$). The change of parameters holds all parameters constant except $\delta$ and $u_{V_i}$, which must necessarily change. The specific change of parameters defined below holds constant all equilibrium densities and infection prevalences for a particular assemblage of host species. The biological interpretation of the constraint is that the densities and infection prevalences for a particular assemblage of host species are known, the pathogen's transmission mode is unknown, and the host densities and infection prevalences are unknown when a set of host species are added to or lost from the community. This method allows one to predict how changes in host 1 infection prevalence and host 1 infected density after the removal or addition of one or more host species depend on the pathogen transmission mode. Specifically, it identifies if greater increases (or smaller decreases) in host 1 infection prevalence and infected density occur under DDDT or FDDT.

Note that in order to apply the following theory, one must know/assume the ratio of the uptake rates for all host species and classes, i.e., know the ratios of $u_{S_i}$, $u_{I_i}$, $u_{S_j}$ and $u_{I_j}$ for all $i, j$. That information is needed to define the change of parameters functions.

**Summary of biological interpretation**: If host species are added to a community, larger reductions (or smaller increases) in host 1 infection prevalence are more likely to occur under

FDDT than DDDT when added host species are weak intraspecific and interspecific competitors and added host species have lower competence than resident hosts. In comparison, larger reductions (or smaller increases) in host 1 infection prevalence are more likely to occur under DDDT than FDDT when added host species are strong intraspecific and interspecific competitors and added host species have higher competence than resident hosts. In addition, if interspecific competition is absent, then reduced host 1 infection prevalence with added host species is more common under FDDT than DDDT, unless the added host species have sufficiently high shedding rates.

The responses of host 1 infected density to increased density of host $n$ are often the same as the responses of host 1 infection prevalence. However, the predictions for host 1 infected density can have the opposite sign when host 1 is weak intraspecific and interspecific competitor and for each host species, regulation by competition with host 1 is weaker than regulation by disease and competition with species $2, ..., n$. This condition is easier to satisfy in systems where infected individuals are stronger competitors than susceptible individuals. The justifications for the above are give after the calculations.

**Additions of host species to a community**: Without loss of generality, assume host species $1, ..., m$ ($m < n$) are present in a community. Assume the species stably coexist at an endemic equilibrium ($p'$; hereafter the 'subcommunity equilibrium'). Let $\mathbf{J}'$ denote the Jacobian of the submodel for species $1, ..., m$ evaluated at $p'$.

The infectious propagule equation is rewritten as

$$\frac{dP}{dt} = \sum_i \chi_i Y_i N_i - g(q)UP - h(q)\delta P \tag{S122}$$

where the change of parameter functions

$$g(q) = \begin{cases} \frac{-\delta}{U'}q + 1 + \frac{\delta}{U'}, & 0 \leq q \leq 1 \\ 2 - q, & 1 \leq q \leq 2 \end{cases} \tag{S123}$$

$$h(q) = \begin{cases} q, & 0 \leq q \leq 1 \\ \frac{-U'}{\delta}(2-q) + 1 + \frac{U'}{\delta}, & 1 \leq q \leq 2 \end{cases} \tag{S124}$$

transform the 1-habitat ET model (S4) from a FDDT form when $q = 0$ to a DDDT form when $q = 2$. The transformation holds constant the subcommunity equilibrium, $p'$.

The sensitivity of host 1 infection prevalence at the community equilibrium to the parameter, $q$, is

$$0 \leq q \leq 1 : \frac{\partial Y_1^*}{\partial q} = \frac{(-1)^{n+1}}{|\mathbf{J}^*|} \overbrace{\beta_1 X_1^*}^{P \to Y_1} \prod_{j=2}^{n} \left(\frac{\beta_i P^*}{Y_i^*}\right) |\mathbf{A}^*| \overbrace{\left(\frac{U^*}{U'}\delta P^* - \delta P^*\right)}^{q \to P} \tag{S125}$$

$$
\begin{aligned}
=&\frac{(-1)^{n+1}}{|\mathbf{J}^*|}\beta_1 X_1^* \prod_{j=2}^{n}\left(\frac{\beta_i P^*}{Y_i^*}\right)\frac{\delta P^*}{U'}|\mathbf{A}^*| \\
&\times\left[\sum_{i=m+1}^{n}\frac{\beta_i\chi_i}{\mu_i+m_i+\nu_i}S_i^* + \sum_{i=1}^{m}\frac{\beta_i\chi_i}{\mu_i+m_i+\nu_i}(S_i^*-S_i')\right]
\end{aligned} \tag{S126}
$$

$$
1\le q\le 2:\frac{\partial Y_1^*}{\partial q} = \frac{(-1)^{n+1}}{|\mathbf{J}^*|}\overbrace{\beta_1 X_1^*}^{P\to Y_1}\prod_{j=2}^{n}\left(\frac{\beta_i P^*}{Y_i^*}\right)|\mathbf{A}^*|\overbrace{(U^*P^*-U'P^*)}^{q\to P} \tag{S127}
$$

$$
\begin{aligned}
=&\frac{(-1)^{n+1}|\mathbf{A}^*|}{|\mathbf{J}^*|}\beta_1 X_1^* \prod_{j=2}^{n}\left(\frac{\beta_i P^*}{Y_i^*}\right)P^*|\mathbf{A}^*| \\
&\times\left[\sum_{i=m+1}^{n}\frac{\beta_i\chi_i}{\mu_i+m_i+\nu_i}S_i^* + \sum_{i=1}^{m}\frac{\beta_i\chi_i}{\mu_i+m_i+\nu_i}(S_i^*-S_i')\right].
\end{aligned} \tag{S128}
$$

where derivatives evaluated at $q = 1$ are treated as left and right derivatives. To get the last equality, solve $dI_i/dt|_{p^*} = 0$ and $dI_i/dt|_{p'} = 0$ for $I_i^*$ and $I_i'$, respectively, substitute into $dP/dt|_{p^*} = 0$ and $dP/dt|_{p'} = 0$, and solve for $U^*$ and $U'$ to get

$$
U^* = \frac{1}{f(q)}\left[-\delta + \sum_{i=1}^{n}\frac{\chi_i\beta_i S_i^*}{m_i+\mu_i+\nu_i}\right], \quad U' = \frac{1}{f(q)}\left[-\delta + \sum_{i=1}^{m}\frac{\chi_i\beta_i S_i'}{m_i+\mu_i+\nu_i}\right]. \tag{S129}
$$

Equations (S126) and (S128) represent the indirect effect $q \to P \to Y_1$. Under the assumptions in Remark S1.2, the signs of both equations are determined by $U^* - U'$.

Using equations (S65), (S318), and (S319) and following the steps in equation (S91), the sensitivity of host 1 infected density at the community equilibrium to the parameter $q$ is

$$
0\le q\le 1:\frac{\partial I_1^*}{\partial q} = \frac{(-1)^{n+1}}{|\mathbf{J}^*|}\beta_1 X_1^* \prod_{j=2}^{n}\left(\frac{\beta_i P^*}{Y_i^*}\right)\left(\frac{U^*}{U'}\delta P^* - \delta P^*\right)\left(S_1^*|\mathbf{A}^*| + \left|\begin{bmatrix}\vec{\omega}_n^* & \mathbf{A}^{*[-,\not{1}]}\end{bmatrix}\right|\right) \tag{S130}
$$

$$
\begin{aligned}
=&\frac{(-1)^{n+1}}{|\mathbf{J}^*|}\beta_1 X_1^* \prod_{j=2}^{n}\left(\frac{\beta_i P^*}{Y_i^*}\right)\frac{\delta P^*}{U'}\left(S_1^*|\mathbf{A}^*| + \left|\begin{bmatrix}\vec{\omega}_n^* & \mathbf{A}^{*[-,\not{1}]}\end{bmatrix}\right|\right) \\
&\times\left[\sum_{i=m+1}^{n}\frac{\beta_i\chi_i}{\mu_i+m_i+\nu_i}S_i^* + \sum_{i=1}^{m}\frac{\beta_i\chi_i}{\mu_i+m_i+\nu_i}(S_i^*-S_i')\right]
\end{aligned} \tag{S131}
$$

$$
1\le q\le 2:\frac{\partial I_1^*}{\partial q} = \frac{(-1)^{n+1}}{|\mathbf{J}^*|}\beta_1 X_1^* \prod_{j=2}^{n}\left(\frac{\beta_i P^*}{Y_i^*}\right)\left(U^*P^* - U'P^*\right)\left(S_1^*|\mathbf{A}^*| + \left|\begin{bmatrix}\vec{\omega}_n^* & \mathbf{A}^{*[-,\not{1}]}\end{bmatrix}\right|\right) \tag{S132}
$$

$$
\begin{aligned}
=&\frac{(-1)^{n+1}}{|\mathbf{J}^*|}\beta_1 X_1^* \prod_{j=2}^{n}\left(\frac{\beta_i P^*}{Y_i^*}\right)P^*\left(S_1^*|\mathbf{A}^*| + \left|\begin{bmatrix}\vec{\omega}_n^* & \mathbf{A}^{*[-,\not{1}]}\end{bmatrix}\right|\right) \\
&\times\left[\sum_{i=m+1}^{n}\frac{\beta_i\chi_i}{\mu_i+m_i+\nu_i}S_i^* + \sum_{i=1}^{m}\frac{\beta_i\chi_i}{\mu_i+m_i+\nu_i}(S_i^*-S_i')\right]
\end{aligned} \tag{S133}
$$

Equations (S126) and (S128) have the same sign and equations (S131) and (S133) have the same sign.

**Removal of host species from a community**: The notation and assumptions are the same as the above. The infectious propagule equation is rewritten as in equation (S122) where the change of parameter functions,

$$g(q) = \begin{cases} \dfrac{-\delta}{U^*}q + 1 + \dfrac{\delta}{U^*}, & 0 \leq q \leq 1 \\ 2 - q, & 1 \leq q \leq 2 \end{cases} \tag{S134}$$

$$h(q) = \begin{cases} q, & 0 \leq q \leq 1 \\ \dfrac{-U^*}{\delta}(2-q) + 1 + \dfrac{U^*}{\delta}, & 1 \leq q \leq 2 \end{cases} \tag{S135}$$

transform the 1-habitat ET model (S4) from a FDDT form when $q = 0$ to a DDDT form when $q = 2$. The transformation holds constant the community equilibrium, $p^*$.

The sensitivity of host 1 infection prevalence at the subcommunity equilibrium to the parameter, $q$, is

$$0 \leq q \leq 1 : \frac{\partial Y_1'}{\partial q} = \frac{(-1)^{m+1}|\mathbf{A}'|}{|\mathbf{J}'|} \overbrace{\beta_1 X_1'}^{P\to Y_1} \prod_{j=2}^{m} \left(\frac{\beta_i P'}{Y_i'}\right) \overbrace{\left(\frac{U'}{U^*}\delta P' - \delta P'\right)}^{q \to P} \tag{S136}$$

$$\begin{aligned} &= \frac{(-1)^{m}|\mathbf{A}'|}{|\mathbf{J}|'} \beta_1 X_1' \prod_{j=2}^{m} \left(\frac{\beta_i P'}{Y_i'}\right) \frac{\delta P'}{U^*} \\ &\times \left[\sum_{i=m+1}^{n} \frac{\beta_i \chi_i}{\mu_i + m_i + \nu_i} S_i^* + \sum_{i=1}^{m} \frac{\beta_i \chi_i}{\mu_i + m_i + \nu_i}(S_i^* - S_i')\right] \end{aligned} \tag{S137}$$

$$1 \leq q \leq 2 : \frac{\partial Y_1'}{\partial q} = \frac{(-1)^{m+1}|\mathbf{A}'|}{|\mathbf{J}'|} \overbrace{\beta_1 X_1'}^{P\to Y_1} \prod_{j=2}^{m} \left(\frac{\beta_i P'}{Y_i'}\right) \overbrace{(U'P' - U^*P')}^{q \to P} \tag{S138}$$

$$\begin{aligned} &= \frac{(-1)^{m}|\mathbf{A}'|}{|\mathbf{J}'|} \beta_1 X_1' \prod_{j=2}^{m} \left(\frac{\beta_i P'}{Y_i'}\right) P' \\ &\times \left[\sum_{i=m+1}^{n} \frac{\beta_i \chi_i}{\mu_i + m_i + \nu_i} S_i^* + \sum_{i=1}^{m} \frac{\beta_i \chi_i}{\mu_i + m_i + \nu_i}(S_i^* - S_i')\right]. \end{aligned} \tag{S139}$$

Equations (S137) and (S139) represent the indirect effect $q \to P \to Y_1$. Under the assumptions in Remark S1.2, the signs of both equations are determined by $-(U^* - U')$.

Using equations (S65), (S137), and (S139) and following the steps in equation (S91), the sensitivity of host 1 infected density at the subcommunity equilibrium to the parameter $q$ is

$$0 \le q \le 1: \frac{\partial I_1'}{\partial q} = \frac{(-1)^m}{|\mathbf{J}'|}\beta_1 X_1' \prod_{j=2}^{m}\left(\frac{\beta_j P'}{Y_j'}\right)\left(\frac{U'}{U^*}\delta P' - \delta P'\right)\left(S_1'|\mathbf{A}'| + \left|\left[\ \vec{\omega}_m' \quad \mathbf{A}'^{[-,1]}\ \right]\right|\right) \tag{S140}$$

$$\begin{aligned} &= \frac{(-1)^m}{|\mathbf{J}'|}\beta_1 X_1' \prod_{j=2}^{m}\left(\frac{\beta_j P'}{Y_j'}\right)\frac{\delta P'}{U^*}\left(S_1'|\mathbf{A}'| + \left|\left[\ \vec{\omega}_m' \quad \mathbf{A}'^{[-,1]}\ \right]\right|\right) \\ &\times \left[\sum_{i=m+1}^{n}\frac{\beta_i \chi_i}{\mu_i + m_i + \nu_i}S_i^* + \sum_{i=1}^{m}\frac{\beta_i \chi_i}{\mu_i + m_i + \nu_i}(S_i^* - S_i')\right] \end{aligned} \tag{S141}$$

$$1 \le q \le 2: \frac{\partial I_1'}{\partial q} = \frac{(-1)^m}{|\mathbf{J}'|}\beta_1 X_1' \prod_{j=2}^{m}\left(\frac{\beta_j P'}{Y_j'}\right)\left(U^* P' - U' P'\right)\left(S_1'|\mathbf{A}'| + \left|\left[\ \vec{\omega}_m' \quad \mathbf{A}'^{[-,1]}\ \right]\right|\right) \tag{S142}$$

$$\begin{aligned} &= \frac{(-1)^m}{|\mathbf{J}'|}\beta_1 X_1' \prod_{j=2}^{m}\left(\frac{\beta_j P'}{Y_j'}\right)P'\left(S_1'|\mathbf{A}'| + \left|\left[\ \vec{\omega}_m' \quad \mathbf{A}'^{[-,1]}\ \right]\right|\right) \\ &\times \left[\sum_{i=m+1}^{n}\frac{\beta_i \chi_i}{\mu_i + m_i + \nu_i}S_i^* + \sum_{i=1}^{m}\frac{\beta_i \chi_i}{\mu_i + m_i + \nu_i}(S_i^* - S_i')\right] \end{aligned} \tag{S143}$$

Equations (S137) and (S139) have the same sign and equations (S141) and (S143) have the same sign.

**Justification for interpretation**: Under the conditions in Section S1.1.1, equations (S126) and (S128) are positive when equations (S137) and (S137) are negative, and vice versa. Thus, to minimize confusion, I will focus on equations (S126) and (S128); the interpretation is qualitatively similar for equations (S137) and (S137).

Under the assumptions in Section S1.1.1, the signs of equations (S126) and (S128) are determined by $U^* - U'$. The quantity $U^* - U'$ is positive unless $S_i^* - S_i' < 0$ for $i \le m$, $\beta_i\chi_i/(\mu_i + m_i + \nu_i)$ is sufficiently large for $i \le m$, and $S_i^*\beta_i\chi_i/(\mu_i + m_i + \nu_i)$ is sufficiently small for $i > m$.

This means that equations (S126) and (S128) are more likely to be positive when (i) species $i > m$ have weak intraspecific interactions, (ii) species $i > m$ have weak interspecific interactions on species $i \le m$, and (iii) species $i > m$ have lower competence than species $i \le m$. Thus, adding host species to a community is more likely to cause larger decreases (or smaller increases) in host 1 infection prevalence under FDDT than DDDT when conditions (i)-(iii) are satisfied. In comparison, equations (S126) and (S128) are more likely to be negative when (iv) species $i > m$ have strong intraspecific interactions, (v) species $i > m$ have strong interspecific interactions on species $i \le m$, and (vi) species $i > m$ have higher competence than species $i \le m$. Thus, adding host species to a community is more likely to cause larger decreases (or smaller increases) in host 1 infection prevalence under DDDT than FDDT when conditions (iv)-(vi) are satisfied.

Proposition S2.1 in Appendix S2.7 of Cortez (2021) shows that in the absence of interspecific competition the addition of species $i > m$ into a community results in host 1 infection prevalence under FDDT and DDDT only if the introduced hosts have sufficiently high shedding rates ($\chi_j$ sufficiently large for $j > m$). This means that in the absence of interspecific competition, added host species dilute disease more and amplify disease less under FDDT than DDDT, unless the added host species have sufficiently high shedding rates.

Now consider the sensitivities of host 1 infected density. To minimize confusion, I will focus on equations (S131) and (S133) because under the conditions in Section S1.1.1, equations (S131) and (S133) are positive when equations (S141) and (S143) are negative, and vice versa. In most cases, the above predictions for host 1 infection prevalence also apply to host 1 infected density. This is because in most cases the signs of equations (S131) and (S133) for $\partial I_1^*/\partial q$ are the same as equations (S126) and (S128) for $\partial I_1^*/\partial q$, respectively. Disagreement only arises if the second determinant in the equations for $\partial I_1^*/\partial q$ has the opposite sign of the first determinant and is larger in magnitude. That occurs when the entries of $\vec{\omega}_n$ are positive and large in magnitude relative to the entries of $\mathbf{A}_{\bullet 1}$. Biologically, that corresponds to when host 1 is a weak intraspecific and interspecific competitor and for each host species, regulation by competition with host 1 is sufficiently weaker than regulation by disease and competition with species $2, ..., n$; see Remark S1.6 for additional details.

### S2 Computing sensitivities for the $n$-habitat model

#### S2.1 Details about the $n$-habitat model

Subsections S2.1.1, S2.1.2, and S2.1.2 provide details about the different forms of the $n$-habitat model. Model variables and parameters are summarized in Table S3. Model assumptions are discussed in Section S2.1.4. The Jacobians used in the sensitivity calculations are defined in Sections S2.1.1-S2.1.3 and the mathematical properties are discussed in Section S1.1.5.

##### S2.1.1 SIR form of the $n$-habitat model

The SIR-form of the the $n$-habitat model describes the dynamics of the densities of susceptible ($S_i$), infected ($I_i$), and recovered ($R_i$) individuals in each host population ($i \leq n$) and the densities of infectious propagules ($P_i$) in each habitat. The SIR-form of the $n$-habitat model is

$$\begin{aligned}
\frac{dN_i}{dt} &= \underbrace{F_i(\cdot)}_{\text{reproduction}} - \underbrace{\beta_i S_i P_i}_{\text{infection}} - \underbrace{m_i S_i}_{\text{mortality}} + \underbrace{\gamma_i R_i}_{\text{loss of immunity}} \\
\frac{dI_i}{dt} &= \underbrace{\beta_i S_i P_i}_{\text{infection}} - \underbrace{(\mu_i + m_i) I_i}_{\text{mortality}} - \underbrace{\nu_i I_i}_{\text{recovery}} \\
\frac{dR_i}{dt} &= \underbrace{\nu_i I_i}_{\text{recovery}} - \underbrace{(\xi_i + m_i) R_i}_{\text{mortality}} - \underbrace{\gamma_i R_i}_{\text{loss of immunity}} \\
\frac{dP_i}{dt} &= \underbrace{\sum_{j=1}^{n} \chi_{ij} I_j}_{\text{propagule shedding}} - \underbrace{\sum_{j=1}^{n} (u_{i,S_j} S_j + u_{i,I_j} I_j + u_{i,R_j} R_j) P_i}_{\text{propagule uptake}} - \underbrace{\delta_i P_i}_{\text{degradation}} .
\end{aligned} \tag{S144}$$

The parameters and variables are summarized in Table S3. The key difference between the 1-habitat SIR model (S1) and the $n$-habitat SIR model (S144) is that $P_i$ denotes the density of infectious propagules in habitat $i$ and the parameters $\chi_{ij}$ and $u_{i,V_j}$ denote the shedding and uptake rates of class $V$ of species $j$ in habitat $i$. Note that the $n$-habitat model (S144) is mathematically equivalent to the 1-habitat model (S1) when each host species has the same shedding and uptake rates in all habitats ($\chi_{ij} = \chi_j$ and $u_{i,V_j} = u_{V_j}$ for all $i, j$ and $V \in \{S, I, R\}$), the degradation rates are equal in all habitats ($\delta_i = \delta$ for all $i$), and the initial densities of the infectious propagules in all habitats are identical ($P_i(0) = P_0$ for all $i$). That is because under those conditions, the infectious propagule densities are equal in all habitats at all points in time ($P_i(t) = P_j(t)$ for all $i, j$).

##### S2.1.2 NYZ form of the $n$-habitat model

The sensitivity calculations for focal host infection prevalence and infected density are easier to interpret biologically if the SIR form of the $n$-habitat model (S144) is written in terms of

Table S3: Variables, parameters, and important quantities for the $n$-habitat model (S144)

| Variable | Description |
|---|---|
| $S_i$ | susceptible density for host $i$ |
| $I_i$ | infected density for host $i$ |
| $R_i$ | recovered density for host $i$ |
| $N_i = S_i + I_i + R_i$ | total density for host $i$ |
| $X_i = S_i/N_i$ | proportion of susceptible individuals for host $i$ |
| $Y_i = I_i/N_i$ | proportion of infected individuals for host $i$ |
| $Z_i = R_i/N_i$ | proportion of recovered individuals for host $i$ |
| $P_i$ | density of infectious propagules in habitat $i$ |
| **Parameter** | **Description** |
| $F_i$ | reproduction rate of host $i$ |
| $f_{V_i}$ | per capita reproduction rate of class $V_i$ ($V \in \{S, I, R\}$) |
| $\beta_i = \rho_i u_{S_i}$ | transmission coefficient for host $i$ |
| $\rho_i$ | per infectious propagule probability of infection |
| $m_i$ | mortality rate due to non-disease related factors |
| $\mu_i$ | disease-induced mortality rate |
| $\xi_i$ | rate of increased mortality due to prior infection |
| $\nu_i$ | recovery rate |
| $\gamma_i$ | rate of loss of immunity |
| $\chi_{ij}$ | shedding rate of host $j$ into habitat $i$ |
| $u_{i,V_j}$ | uptake rate for individuals in class $V_j$ ($V \in \{S, I, R\}$) in habitat $i$ |
| $\delta_i$ | degradation rate of infection propagules in habitat $i$ |
| $\Gamma_i$ | $= \nu_i/(m_i + \xi_i + \gamma_i)$, ratio of $I_i$ to $R_i$ at the endemic equilibrium |
| **Quantity** | **Description** |
| $p^\ddagger$ | $n$-species disease-free community equilibrium |
| $p^*$ | $n$-species endemic community equilibrium |
| $p^\dagger$ | $m$-species ($m < n$) disease-free subcommunity equilibrium |
| $p'$ | $m$-species ($m < n$) endemic subcommunity equilibrium |
| $U_i$ | $= \sum_i (u_{iS_j} S_j + u_{iI_j} I_j + u_{iR_j} R_j)$ |
| | total per propagule rate of uptake of infectious propagules in habitat $i$ |
| $\varphi_{ji}$ | per capita net production of infectious propagules by host $i$ in habitat $j$ |
| | defines if host $i$ is a source ($\varphi_{ji} > 0$) or sink ($\varphi_{ji} > 0$) for habitat $j$ |
| $\mathcal{R}_{0,i}$ | pathogen basic reproduction number for host $i$ |
| $\mathcal{R}_{0,i}(\infty)$ | $= \beta_i \chi_i / u_{S_i}(m_i + \mu_i + \gamma_i) = \rho_i \chi_i/(m_i + \mu_i + \gamma_i)$, competence of host $i$ |
| | basic reproduction number for host $i$ in the limit of infinite host density |
| $\mathcal{R}_0^C$ | pathogen community reproduction number |

Double daggers (‡) denote values evaluated at the $n$-species disease-free community equilibrium ($p^\ddagger$), asterisks ($*$) denote values evaluated at the $n$-species endemic community equilibrium ($p^*$), daggers (†) denote values evaluated at the $m$-species disease-free community equilibrium ($p^\dagger$), and primes (′) denote values evaluated at the $m$-species endemic community equilibrium ($p'$). The mnemonic for number of species is that more lines are needed to create the symbols denoting equilibria for communities with more species (‡, *).

the total density for each host ($N_i$) and the proportions of susceptible, infected and recovered individuals in each host population ($X_i$, $Y_i$, and $Z_i$). The NYZ model with $n$ habitats is

$$
\begin{aligned}
\frac{dN_i}{dt} &= \overbrace{F_i(\cdot)}^{\text{reproduction}} - \overbrace{m_i N_i}^{\text{natural mortality}} - \overbrace{(\mu_i Y_i N_i + \xi_i Z_i N_i)}^{\text{disease-related mortality}} \\
\frac{dY_i}{dt} &= \overbrace{\beta_i(1 - Y_i - Z_i)P_i}^{\text{infection}} - \overbrace{(\mu_i + m_i)Y_i}^{\text{mortality}} - \overbrace{\nu_i Y_i}^{\text{recovery}} - \frac{Y_i}{N_i}\frac{dN_i}{dt} \\
\frac{dZ_i}{dt} &= \overbrace{\nu_i Y_i}^{\text{recovery}} - \overbrace{(\xi_i + m_i)Z_i}^{\text{mortality}} \overbrace{-\gamma_i Z_i}^{\text{loss of immunity}} - \frac{Z_i}{N_i}\frac{dN_i}{dt} \\
\frac{dP_i}{dt} &= \underbrace{\sum_{j=1}^{n} \chi_{ij} Y_j N_j}_{\text{propagule shedding}} - \underbrace{\sum_{j=1}^{n} (u_{i,S_j} X_j + u_{i,I_j} Y_j + u_{i,R_j} Z_j) N_j P_i}_{\text{propagule uptake}} - \underbrace{\delta_i P_i}_{\text{degradation}}
\end{aligned}
\tag{S145}
$$

All parameters are interpreted as in the SIR form of the $n$-habitat model (S144); see Table S3. All terms in the model are interpreted as in the 1-habitat NYZ model (S2) in Section S1.1.2. The total per propagule uptake rate of infectious propagules in habitat $i$ is denoted $U_i = \sum_{j=1}^{n}(u_{i,S_j} X_j + u_{i,I_j} Y_j + u_{i,R_j} Z_j) N_j$.

**Jacobian**: While the Jacobian of the $n$-habitat NYZ model (S145) is not used in the analysis in this study, I include it here for completeness. When evaluated at an endemic equilibrium, the Jacobian of the $n$-habitat NYZ model (S145) has the structure,

$$
\begin{bmatrix}
\mathbf{A} & \mathbf{B} & \mathbf{C} & \mathbf{0}_{n\times n} \\
-\mathbf{D}_{Y/N}\mathbf{A} & -\mathbf{D}_{Y/N}\mathbf{B} - \mathbf{D}_{\beta P+\mu+m+\nu} & -\mathbf{D}_{Y/N}\mathbf{C} - \mathbf{D}_{\beta P} & \mathbf{D}_{\beta X} \\
-\mathbf{D}_{Z/N}\mathbf{A} & -\mathbf{D}_{Z/N}\mathbf{B} + \mathbf{D}_{\nu} & -\mathbf{D}_{Z/N}\mathbf{C} - \mathbf{D}_{\xi+m+\gamma} & \mathbf{0}_{n\times n} \\
\mathbf{E}_N & \mathbf{E}_Y & \mathbf{E}_Z & -\mathbf{D}_U - \mathbf{D}_\delta
\end{bmatrix}.
\tag{S146}
$$

where the order of the rows and columns is $N_1, ..., N_n, Y_1, ..., Y_n, Z_1, ..., Z_n, P_1, ..., P_n$. The interpretations of the Jacobian entries are similar to those for the Jacobian of the 1-habitat NYZ model (S3). There are three minor differences. First, the diagonal matrix $\mathbf{D}_{\beta X}$ accounts for the fact that the growth rate of infection prevalence of host $i$ only increases with changes in the density of infectious propagules in its habitat. Second, the matrices $\mathbf{E}_N$, $\mathbf{E}_Y$, and $\mathbf{E}_Z$ account for how changes in the densities and frequencies of each host species affect the infectious propagule growth rates in each habitat. Third, the diagonal form of the matrix $-\mathbf{D}_U - \mathbf{D}_\delta$ accounts for the fact that the infectious propagules in different habitats do not have direct interactions. The entries of the Jacobian are described in detail in Section S2.1.5.

##### S2.1.3 Reduced NY form of the $n$-habitat model

The calculations of the sensitivities at equilibrium are simplified by analyzing a reduced version of model (S145). The reduced form is derived by setting $dZ_i/dt = 0$, solving for $Z_i^* = \nu_i Y_i^*/(\xi_i + m_i + \gamma_i) = \Gamma_i Y_i^*$, and substituting into the other differential equations to yield,

$$
\begin{aligned}
\frac{dN_i}{dt} &= \overbrace{F_i(\cdot)}^{\text{reproduction}} - \overbrace{m_i N_i}^{\text{natural mortality}} - \overbrace{(\mu_i Y_i N_i + \xi_i \Gamma_i Y_i N_i)}^{\text{disease-related mortality}} \\
\frac{dY_i}{dt} &= \overbrace{\beta_i(1 - Y_i - \Gamma_i Y_i)}^{\text{infection}} - \overbrace{(\mu_i + m_i)Y_i}^{\text{mortality}} - \overbrace{\nu_i Y_i}^{\text{recovery}} - \frac{Y_i}{N_i}\frac{dN_i}{dt} \\
\frac{dP_i}{dt} &= \underbrace{\sum_{j=1}^{n} \chi_{ij} Y_j N_j}_{\text{propagule shedding}} - \underbrace{\sum_{j=1}^{n} (u_{i,S_j} X_j + u_{i,I_j} Y_j + u_{i,R_j} \Gamma_j Y_j) N_j P_i}_{\text{propagule uptake}} - \underbrace{\delta_i P_i}_{\text{degradation}}
\end{aligned}
\tag{S147}
$$

where $\Gamma_i = \nu_i/(\xi_i + m_i + \gamma_i)$. Note that any equilibrium point of the $n$-habitat NYZ model (S145) is also an equilibrium of the $n$-habitat NY model (S147). Consequently, the sensitivities of host 1 infection prevalence ($Y_1^*$) and infected density ($I_1^*$) at equilibrium can be computed using either model. I analyze the $n$-habitat NY model (S147) because that removes algebraic steps that must be completed when computing each sensitivity.

**Jacobian**: When evaluated at an endemic equilibrium, the Jacobian for the $n$-habitat NY model (S147) has the structure,

$$
\mathbf{J} = \begin{bmatrix} \mathbf{A} & \mathbf{B} + \mathbf{C}\mathbf{D}_\Gamma & \mathbf{0}_{n\times n} \\ -\mathbf{D}_{Y/N}\mathbf{A} & -\mathbf{D}_{\beta P/Y} - \mathbf{D}_{Y/N}(\mathbf{B} + \mathbf{C}\mathbf{D}_\Gamma) & \mathbf{D}_{\beta X} \\ \mathbf{E}_N & \mathbf{E}_Y + \mathbf{E}_Z\mathbf{D}_\Gamma & -\mathbf{D}_{U+\delta} \end{bmatrix},
\tag{S148}
$$

where the order of the rows and columns is $N_1, ..., N_n, Y_1, ..., Y_n, P_1, ..., P_n$. Details about each $n \times n$ matrix are given in Section S2.1.5. I note that the matrix $\mathbf{D}_{\beta P/Y}$ follows from Remark S2.1.

**Special case of fixed host densities**: One tractable special case is when the host densities are fixed, which results in the $n$-habitat NY model (S147) reducing to

$$
\begin{aligned}
\frac{dY_i}{dt} &= \overbrace{\beta_i(1 - Y_i - \Gamma_i Y_i)}^{\text{infection}} - \overbrace{(\mu_i + m_i)Y_i}^{\text{mortality}} - \overbrace{\nu_i Y_i}^{\text{recovery}} \\
\frac{dP_i}{dt} &= \underbrace{\sum_{j=1}^{n} \chi_{ij} Y_j N_j}_{\text{propagule shedding}} - \underbrace{\sum_{j=1}^{n} (u_{i,S_j} X_j + u_{i,I_j} Y_j + u_{i,R_j} \Gamma_j Y_j) N_j P_i}_{\text{propagule uptake}} - \underbrace{\delta_i P_i}_{\text{degradation}}
\end{aligned}
\tag{S149}
$$

where the host densities ($N_i$) are treated as fixed parameters. The Jacobian for this reduced model is

$$
\hat{\mathbf{J}} = \begin{bmatrix} -\mathbf{D}_{\beta P/Y} & \mathbf{D}_{\beta X} \\ \mathbf{E}_Y + \mathbf{E}_Z\mathbf{D}_\Gamma & -\mathbf{D}_{U+\delta} \end{bmatrix}.
\tag{S150}
$$

##### S2.1.4 Details about model assumptions

Details about the specific components of the $n$-habitat models and assumptions are given below. Some assumptions are defined by quantities derived from the Jacobian in equation (S148); details about those assumptions are provided in Section S2.1.5.

**Pathogen transmission and habitat overlap**: The $n$-habitat model assumes susceptible individuals become infected when they encounter infectious propagules in their habitat. In addition, the models assume infectious propagules are well-mixed throughout the habitat and do not differ based on which species shed them.

I assume infected individuals shed infectious propagules into their own habitat and at least one habitat of another species. In addition, I assume that the habitats are connected in such way that for any pair of host species $i$ and $j$, the pathogen can be transmitted from host species $j$ to host species $i$, possibly through intermediary species. Mathematically, this means for any $i, j$ there exists a sequence $\beta_{k_1}\chi_{k_1 i}\beta_{k_1}\chi_{k_1 k_2}...\beta_{k_l}\chi_{k_l k_{l-1}}\beta_j\chi_{j k_l} > 0$. Thus, host $j$ can transmit the pathogen to host $z$, who transmits the pathogen to host $y$, and so forth until the pathogen is transmitted from host $k$ to host $i$. One consequence of this assumption is that there are no dead end hosts in the community (host $j$ is a dead end host if it does not shed the pathogen and satisfies $\chi_{ij} = 0$ for all $i$). This assumption is made for convenience when computing the community reproduction number, $\mathcal{R}_0^C$, but all of my results are unchanged if this assumption is relaxed.

**Equilibria, stability, and notation**: I assume the $n$ species can stably coexist in the presence of the pathogen at the $n$-species community equilibrium ($p^*$) and they can stably coexist in the absence of the pathogen at the $n$-species disease-free community equilibrium ($p^\ddagger$). For the analyzes that consider communities of $m < n$ species, I assume the $m$ species can stably coexist in the presence of the pathogen at the $n$-species subcommunity equilibrium ($p'$) and they can stably coexist in the absence of the pathogen at the $m$-species disease-free subcommunity equilibrium ($p^\dagger$). Throughout, asterisks ($*$) denote quantities evaluated at the endemic community equilibrium ($p^*$), double daggers ($\ddagger$) denote quantities evaluated at the disease-free community equilibrium ($p^\ddagger$), primes ($'$) denote quantities evaluated at the endemic subcommunity equilibrium ($p'$), and daggers ($\dagger$) denote quantities evaluated at the disease-free subcommunity equilibrium ($p^\dagger$). As before, the mnemonics are (i) asterisks denote endemic communities with more species because more line segments are needed to draw an asterisk than a prime and (ii) double daggers denote disease-free communities with more species because they have more lines than daggers.

**Other model assumptions**: With the exception of the assumptions listed above, all of the assumptions in Section S1.1.4 for the 1-habitat models also apply to the $n$-habitat model. This includes the assumptions about the host reproduction rates ($F_i$), the relationship between the transmission and uptake parameters ($\beta_i = u_{i,S_i} p_i$), and negative density dependence at equilibrium ($\frac{\partial}{\partial N_i}\frac{dN_i}{dt} < 0$).

**Assumptions about Jacobian of $n$-habitat NY model and biological interpretation**: I make assumptions about the $n$-habitat models in order to simplify the analysis and focus it on biological scenarios that are more likely to be observed in natural systems. Assumptions 1-9 for the 1-habitat model in Section S1.1.1 apply to the $n$-habitat model. Only one minor difference arises due to the increased dimension of the $n$-habitat model and one additional assumption is needed to ensure that the habitats are sufficiently connected.

4. All $n$ host species coexist with the pathogen at a stable endemic equilibrium of the model: $\text{sign}(|\mathbf{J}|) = (-1)^{3n}$

10. All habitats are connected such that no species or subset of species is isolated from the others and all species shed infections propagules such that the pathogen can be transmitted from species $j$ to species $i$, possibly through intermediary species: For any $i, j$ there exists a sequence $\beta_{k_1}\chi_{k_1 i}\beta_{k_1}\chi_{k_1 k_2}...\beta_{k_l}\chi_{k_l k_{l-1}}\beta_j\chi_{j k_l} > 0$.

**Special Cases**: The formulas for the sensitivities are too complex to analyze in general. Thus, much of the following focuses on two special cases that are analytically tractable:

- Special Case 1 assumes (i) the host densities are fixed ($N_i$ fixed), (ii) susceptible, infected, and recovered individuals have equal uptake rates ($u_{i,S_j} = u_{i,I_j} = u_{i,S_j}$ for all $i, j$), and (iii) the shedding rate of host $i$ into the habitat of host $j$ is sufficiently smaller than the shedding rate of host $i$ into its own habitat ($\chi_{ii} \gg \chi_{ij}$ for $i \neq j$). For this special case, the model reduces to a simpler form given by model (S149).

- Special Case 2 assumes (i) there is no interspecific host competition ($\mathbf{A}_{ij} = 0$ and $\mathbf{B}_{ij} = 0$ for $i \neq j$), (ii) individuals from all host species have equal uptake rates ($u_{i,V_j} = u$ for all $i, j$ and $V, W \in \{S, I, R\}$), (iii) the shedding rate of host $i$ into the habitat of host $j$ is sufficiently smaller than the shedding rate of host $i$ into its own habitat ($\chi_{ii} \gg \chi_{ij}$ for $i \neq j$), and (iv) susceptible, infected, and recovered individuals of each species have equal competitive abilities ($\partial f_{V_i}/\partial S_j = \partial f_{V_i}/\partial I_j$ for all $i, j$).

Note that Special Case 2 aligns with the modeling assumptions of prior studies (Dobson 2004, Joseph et al. 2013, Mihaljevic et al. 2014) on pathogens with density-dependent direct transmission (where $u_{i,V_j} = 0$ for all $i, j$ and $V \in \{S, I, R\}$) and frequency-dependent direct transmission (where $\delta_i = 0$ for all $i$).

###### S2.1.5 Properties and derived quantities of the Jacobian of the $n$-habitat NY model

Because I assume the $n$-host endemic equilibrium is stable, $\text{sign}(|\mathbf{J}|) = (-1)^{3n}$ when evaluated at $p^*$.

**Properties of A**: All entries of $\mathbf{A}$ are negative because of the assumptions about the reproduction functions, $F_i$. The calculations are identical to those for the 1-habitat NY model (S4) in Section S1.1.5.

**Properties of C**: Under the assumption that susceptible and recovered individuals have equal reproduction rates ($f_{S_i} = f_{R_i}$) and equal competitive abilities ($\partial f_{V_i}/\partial S_j = \partial f_{V_i}/\partial R_j$

for $V \in \{S, I, R\}$), the entries of $\mathbf{C}$ simplify to $\mathbf{C}_{ii} = -\xi_i N_i^*$ and $\mathbf{C}_{ij} = 0$ for $i \neq j$.

**Properties of B and $\mathbf{B} + \mathbf{C}\mathbf{D}_\Gamma$**: These values are defined as in Section S1.1.5. For most empirical systems, I expect that (i) infected individuals have reduced reproductive output compared to susceptible individuals ($f_{I_i} - f_{S_i} < 0$) and (ii) infected individuals can be weaker, equal or stronger competitors than susceptible individuals, but infected individuals are not much weaker competitors than susceptible individuals ($\partial f_{V_i}/\partial I_j$ is never much smaller in magnitude than $\partial f_{V_i}/\partial S_j$). Under these assumptions, $\mathbf{B}_{ii} + \Gamma_i \mathbf{C}_{ii}$ is negative unless infected individuals of host $i$ are sufficiently weaker intraspecific competitors than susceptible individuals. The off-diagonal entries of $\mathbf{B} + \mathbf{C}\mathbf{D}_\Gamma$ are positive, zero, and negative when infected individuals of host $j$ are weaker, equal or stronger interspecific competitors than susceptible individuals of host $j$, respectively.

**Properties of $\mathbf{E}_N$, $\mathbf{E}_Y$, and $\mathbf{E}_Z$ and definition of sink/source**: The three matrices have entries

$$\begin{aligned}\mathbf{E}_N[i,j] &= \chi_{ij} Y_j^* - (u_{iS_j} X_j^* + u_{iI_j} Y_j^* + u_{iR_j} Z_j^*) P_i^* \\ &= Y_j^* \varphi_{ij}\end{aligned} \tag{S151}$$

$$\begin{aligned}\mathbf{E}_Y[i,j] &= \chi_{ij} N_j^* - (u_{iI_j} - u_{iS_j}) N_j^* P_i^* \\ &= N_j^* \varphi_{ij} + N_j^* u_{is_j} P_i^* / Y_j^*\end{aligned} \tag{S152}$$

$$\mathbf{E}_Z[i,j] = -(u_{iR_j} - u_{iS_j}) Z_j^* P_i^* = 0 \tag{S153}$$

where $\mathbf{E}_Z[i,j] = 0$ because I assume susceptible and recovered individuals have equal uptake rates. The quantity $\varphi_{ij}$ defines if host $j$ is a source ($\varphi_{ij} > 0$) or sink ($\varphi_{ij} < 0$) of infectious propagule for habitat $i$ at equilibrium; see Section S2.3 for details.

**Remark S2.1.** *For any host species, the following holds at equilibrium*

$$\beta_i P_i^* + \mu_i + m_i + \nu_i + \frac{\beta_i P_i^* \nu_i}{\xi_i + m_i + \gamma_i} = \frac{\beta_i P_i^*}{Y_i^*} \tag{S154}$$

*Reasoning:* The equilibrium conditions $dN_i/dt = 0$ and $dY_i/dt = 0$ imply that $\beta_i P_i^* + \mu_i + m_i + \nu_i = \beta_i (1 - Z_i^*) P_i^* / Y_i^*$. Similarly, the equilibrium conditions $dN_i/dt = 0$ and $dR_i/dt = 0$ imply that $\nu_i/(\xi_i + m_i + \gamma_i) = Z_i^*/Y_i^*$. Substitution and a little algebra yields

$$\beta_i P_i^* + \mu_i + m_i + \nu_i + \frac{\beta_i P_i^* \nu_i}{\xi_i + m_i + \gamma_i} = \frac{\beta_i (1 - Z_i^*) P_i^*}{Y_i^*} + \frac{\beta_i P_i^* Z_i^*}{Y_i^*} = \frac{\beta_i P_i^*}{Y_i^*}. \tag{S155}$$

•

**Proposition S2.2.** *The diagonal entries of* $\mathbf{E}_Y - \mathbf{D}_{(U+\delta)P/XY}$ *are negative.*

*Proof.* The diagonal entries can be algebraically manipulated to yield

$$\mathbf{E}_Y[i,i] - \mathbf{D}_{(U+\delta)P/XY}[i,i]$$

$$=\chi_{ii}N_i^* - u_{iI_i}N_i^*P_i^* + u_{iS_i}N_i^*P_i^* - \frac{(U_i^*+\delta_i)P_i^*}{X_i^*Y_i^*} \tag{S156}$$

$$=\chi_{ii}N_i^* - u_{iI_i}N_i^*P_i^* - \frac{(U_i^*+\delta_i)P_i^*}{Y_i^*} + u_{iS_i}N_i^*P_i^* - \frac{(U_i^*+\delta_i)P_i^*(Y_i^*+Z_i^*)}{Y_i^*} \tag{S157}$$

$$\begin{aligned}
=&\frac{1}{Y_i^*}\left[\chi_{ii}N_i^*Y_i^* - (U_i^*+\delta_i)P_i^*\right] - u_{iI_i}N_i^*P_i^* \\
&+ u_{iS_i}N_i^*P_i^* - (u_{iS_i}X_i^* + u_{iI_i}Y_i^* + u_{iR_i}Z_i^*)N_i^*P_i^*\frac{(Y_i^*+Z_i^*)}{X_i^*Y_i^*} \\
&- \sum_{j\neq i}(u_{iS_j}X_j^* + u_{iI_j}Y_j^* + u_{iR_j}Z_j^*)N_j^*P_i^*\frac{(Y_i^*+Z_i^*)}{X_i^*Y_i^*} - \delta_iP_i^*\frac{(Y_i^*+Z_i^*)}{X_i^*Y_i^*}
\end{aligned} \tag{S158}$$

$$\begin{aligned}
=&\frac{1}{Y_i^*}\left[\chi_{ii}N_i^*Y_i^* - (U_i^*+\delta_i)P_i^*\right] - u_{iI_i}N_i^*P_i^* \\
&+ \left[u_{iS_i}N_i^*P_i^* - u_{iS_i}X_iN_i^*P_i^*\frac{(Y_i^*+Z_i^*)}{X_i^*Y_i^*}\right] - (u_{iI_i}Y_i^* + u_{iR_i}Z_i^*)N_i^*P_i^*\frac{(Y_i^*+Z_i^*)}{X_i^*Y_i^*} \\
&- \sum_{j\neq i}(u_{iS_j}X_j^* + u_{iI_j}Y_j^* + u_{iR_j}Z_j^*)N_j^*P_i^*\frac{(Y_i^*+Z_i^*)}{X_i^*Y_i^*} - \delta_iP_i\frac{(Y_i^*+Z_i^*)}{X_i^*Y_i^*}
\end{aligned} \tag{S159}$$

The first line of equation (S159) is negative because $dP_i/d|_{p^*} = \sum_j \chi_{ij}Y_j^*N_j^* - (U_i^*+\delta_i)P_i^* = 0$. All other lines of equation (S159) are negative. □

**Remark S2.3.** *For any parameter $\sigma$, the sensitivities $\partial Y_1^*/\partial\sigma$ and $\partial I_1^*/\partial\sigma$ are more likely to differ in sign when (i) regulation of a host species by competition with host species 1 is sufficiently weaker than regulation by the disease and competition with other host species or (ii) host 1 is sink or small source and some subset of hosts $j > 1$ are sufficiently large sources. Condition (i) is most likely to be satisfied when the host 1 is more strongly regulated by disease or interspecific competition than by intraspecific competition. Also, condition (i) is easier to satisfy in systems where infected individuals are equal or stronger competitors than susceptible individual.*

*Reasoning:* As shown in the following sections, the matrices used to compute the sensitivities $\partial Y_1^*/\partial\sigma$ and $\partial I_1^*/\partial\sigma$ only differ in one column. In particular, the matrices used to compute the sensitivities for host 1 infection prevalence and host 1 infected density, respectively, have columns of the form

$$\begin{bmatrix} N_1^*\mathbf{A}_{\bullet 1} \\ N_1^*\mathbf{E}_N[\bullet,1] \end{bmatrix} \text{versus} \begin{bmatrix} -Y_1^*(\mathbf{B}_{\bullet 1} + \mathbf{C}_{\bullet 1}\Gamma_1) \\ -Y_1^*\mathbf{E}_Y[\bullet,1] + \vec{e}_1\frac{(U_1^*+\delta_1)P_1^*}{X_1^*} \end{bmatrix}, \tag{S160}$$

where $\vec{e}_1$ is the unit vector with a 1 in its first entry. The sensitivities for host 1 infection prevalence and host 1 infected density can have opposite signs when the signs or magnitudes of the entries of the columns differ. This occurs under two general conditions.

First, the sensitivities for host 1 infection prevalence and host 1 infected density can have opposite signs if the entries of $-Y_1^*(\mathbf{B}_{\bullet 1} + \mathbf{C}_{\bullet 1}\Gamma_1)$ are positive and large in magnitude relative to the negative entries of $N_1^*\mathbf{A}_{\bullet 1}$. Biologically, this means that some host species are weakly regulated by competition with host species 1 and strongly regulated by disease or competition host species 2,...,$n$. This is more likely to occur when (a) disease-induced mortality is higher ($\mu_i$ and $\xi_i$ are large), (b) infected individuals are stronger or equal competitors than susceptible individuals ($\partial f_{V_i}/\partial I_j - \partial f_{V_i}/\partial S_j$ is negative for $V \in \{S, I, R\}$; this makes $N_1^*\mathbf{A}_{j1}$ negative and $-Y_1^*\mathbf{B}_{j1}$ positive), and (c) host 1 is a relatively weak intraspecific and interspecific competitor ($\mathbf{A}_{i1}$ is small in magnitude relative to $\mu_i$ and $\xi_i$). In our numerical simulations, differences in the signs of the determinants occurred most often when regulation of host 1 by intraspecific competition was weaker than regulation by disease and competition with the other host species.

Second, the sensitivities for host 1 infection prevalence and host 1 infected density can have opposite signs if $N_1^*\vec{E}_N[\bullet, 1]$ is negative or small in magnitude and $-Y_1^*\mathbf{E}_Y[\bullet, 1] + \vec{e}_1 \frac{(U_1^*+\delta_1)P_1^*}{X_1^*}$ is positive and large in magnitude. Biologically, this means host 1 is a sink or small source and some subset of the other host species are large sources. This is more likely to occur when (a) host 1 is either a sink or a small source ($\varphi_1$ negative or positive and small in magnitude, resulting in $N_1^*\vec{E}_N[1]$ being either negative or positive and small in magnitude) and (b) some subset of the host species $j > 1$ are large sources ($\chi_{1j}Y_j^*N_j^*$ is large for some $j \neq 1$). Note that condition (b) can be seen via $(U_1^* + \delta_1)P_1^* = \sum_j \chi_{1j}Y_j^*N_j^*$. •

#### S2.2 Relationship to density-dependent and frequency-dependent direct transmission models

The $n$-habitat environmental transmission model (S144) reduces to a model with direct transmission when the dynamics of the infection propagules ($dP_i/dt$ equation) are much faster than the host dynamics ($dS_i/dt$, $dI_i/dt$, $dR_i/dt$ equations). In particular, the $n$-habitat environmental transmission (ET) model reduces to a model with density-dependent direct transmission (DDDT) if all uptake rates are zero ($u_{j,S_i} = u_{j,I_i} = u_{j,R_i} = 0$ for all $i, j$) and it reduces to a model with frequency-dependent direct transmission (FDDT) if all degradation rates are zero ($\delta_j = 0$). Moreover, there is an onto mapping of parameterizations of the $n$-habitat ET model (S144) to parameterizations of the FDDT and DDDT models with any possible set of interspecific and intraspecific transmission parameter $\hat{\beta}_{ij}$. The two important implications of this are (i) the $n$-habitat ET model (S144) can produce dynamics identical to any parameterization of an $n$-species FDDT or DDDT model and (ii) all of the results about $n$-habitat ET model (S144) apply to the DDDT and FDDT models, after accounting for the restrictions the uptake and degradation rates.

The environmental transmission model (S144) reduces to a model with direct transmission when the dynamics of all infection propagules ($dP_i/dt$ equation) are much faster than the host dynamics ($dS_i/dt$, $dI_i/dt$, $dR_i/dt$ equations). Mathematically, this occurs when the shedding rates ($\chi_{ij}$) and some combination of the uptake ($u_{i,V_j}$) and degradation ($\delta_i$) rates in each infectious propagule equation are sufficiently large such that there is a separation of time scales between the dynamics of the infectious propagules and the dynamics of the host classes. Under the assumed separation of time scales, the infectious propagule density reaches a quasi-steady state equilibrium defined by $P_i = \left(\sum_j \chi_{ij}I_j\right) \Big/ \left[\delta_i + \sum_j (u_{i,S_j}S_j + u_{i,I_j}I_j + u_{i,R_j}R_j)\right]$. Substitution into the equations for the susceptible and infected classes yields

$$
\begin{aligned}
\frac{dS_i}{dt} &= \overbrace{F_i(\cdot)}^{\text{reproduction}} - \overbrace{m_i S_i}^{\text{mortality}} - \overbrace{\frac{\sum_j \chi_{ij}\beta_i S_i I_j}{\delta_i + \sum_j (u_{i,S_j} S_j + u_{i,I_j} I_j + u_{i,R_j} R_j)}}^{\text{infection}} + \overbrace{\gamma_i R_i}^{\text{loss of immunity}} \\
\frac{dI_i}{dt} &= \overbrace{\frac{\sum_j \chi_{ij}\beta_i S_i I_j}{\delta_i + \sum_j (u_{i,S_j} S_j + u_{i,I_j} I_j + u_{i,R_j} R_j)}}^{\text{infection}} - \overbrace{(\mu_i + m_i) I_i}^{\text{mortality}} + \overbrace{\nu_i I_i}^{\text{recovery}} \\
\frac{dR_i}{dt} &= \underbrace{\nu_i I_i}_{\text{recovery}} - \underbrace{(\xi_i + m_i) I_i}_{\text{mortality}} - \underbrace{\gamma_i I_i}_{\text{loss of immunity}} .
\end{aligned}
\tag{S161}
$$

Model (S161) is a direct transmission model where the nonlinear transmission rate from host $j$ to host $i$ is

$$
\frac{\chi_{ij}\beta_i S_i I_j}{\delta_i + \sum_j (u_{i,S_j} S_j + u_{i,I_j} I_j + u_{i,R_j} R_j)}.
$$

This nonlinear transmission rate is a saturating function of the susceptible density of host $i$ ($S_i$) and the infected density of host $j$ ($I_j$) and a decreasing function of all other host densities.

**Density-dependent direct transmission (DDDT):** Model (S161) reduces to a DDDT model when uptake is negligible ($u_{j,V_i} = 0$ for all $i, j$ and $V \in \{S, I, R\}$. Specifically, under this condition, the model reduces to a form with direct transmission rates $\hat{\beta}_{ij} S_i I_j$ where $\hat{\beta}_{ij} = \chi_{ij}\beta_i/\delta_i$.

To show there is an onto mapping of parameterizations of the $n$-habitat ET model (S144) to parameterizations of DDDT models, I will show that one can convert any DDDT model into a well-defined ET model. I start with a DDDT model of the form

$$
\begin{aligned}
\frac{dS_i}{dt} &= F_i(\cdot) - m_i S_i - S_i \sum_{j=1}^{n} \hat{\beta}_{ij} I_j + \gamma_i R_i \\
\frac{dI_i}{dt} &= S_i \sum_{j=1}^{n} \hat{\beta}_{ij} I_j - (\mu_i + m_i) I_i + \nu_i I_i \\
\frac{dR_i}{dt} &= \nu_i I_i - (\xi_i + m_i) I_i - \gamma_i I_i.
\end{aligned}
\tag{S162}
$$

The simple way to show the onto mapping is to choose the parameters of the $n$-habitat ET model as $u_{i,V_j} = 0$ for all $i, j, W$, $\chi_j = \hat{\chi}_j/\epsilon$, $E_{ij} = \hat{E}_{ij}$, $\rho_i = r\hat{ho}_j$, $E_i = 1$, and $\delta = 1/\epsilon$ for $0 < \epsilon \ll 1$, and for any parameters values that are shared between the ET model (S144) and the DDDT model (S162), set the values in the ET model equal to those in the DDDT model (e.g., set $\mu_i$ in the ET model equal to the value of $\mu_i$ in the DDDT model). Under these conditions, the ET model (S144) reduces to a DDDT model with parameters identical those in the DDDT model (S162).

When defining host competence, it is useful to define the parameters of the $n$-habitat ET model in the following way. First, set $u_{j,V_i} = 0$ for all $i, j, V$. Second, for any parameters values that are shared between the ET model (S144) and the DDDT model (S162), set the values in the ET model equal to those in the DDDT model (e.g., set $\mu_i$ in the ET model equal to the value of $\mu_i$ in the DDDT model). Third, decompose the direct transmission parameter as $\hat{\beta}_{ij} = \hat{E}_{ij}\hat{\chi}_j\hat{\rho}_i$ where $\hat{E}_{ij}$ is the encounter rate between infected individuals of host $j$ and susceptible individuals of host $i$, $\hat{\chi}_j$ is the infectious dose that an infected individual of host $j$ exposes a contact to, and $\hat{\rho}_j$ is the probability of infection for host $i$ given exposure. Similarly, decompose the environmental transmission rate as $\beta_i = E_i\rho_i$ where $E_i$ is the encounter rate between host $i$ and infectious propagules in its habitat, and $\rho_i$ is the per infectious propagule probability of infection. Also, decompose the shedding rate as $\chi_{ij} = E_{ij}\chi_j$ where $E_{ij}$ is the contact rate infected individuals of host $j$ and the habitat of host $i$, and $\chi_j$ is the shedding rate of host $j$. Fourth, set $\chi_j = \hat{\chi}_j/\epsilon$, $E_{ij} = \hat{E}_{ij}$, $\rho_i = \hat{\rho}_j$, $E_i = 1$, and $\delta = 1/\epsilon$. The parameter $\epsilon$ is a free parameter that should be set to a very large value in order to ensure that there is a separation of time scales between the dynamics of the infectious propagules and the host classes. Under these conditions, the ET model (S144) reduces to a DDDT model with parameters identical those in the DDDT model (S162).

**Frequency-dependent direct transmission (FDDT):** Model (S161) reduces to a DDDT model when there is no degradation of infectious propagules degradation ($\delta_i = 0$). Specifically, under this condition, the model reduces to a form with direct transmission rates $\hat{\beta}_{ij}S_iI_j/\sum_k(w_{S_k}S_k + w_{I_k}I_k + w_{R_k}R_k)$ where $\hat{\beta}_{ij} = \beta_i\chi_j$ and $w_{V_k} = u_{V_k}$ for $V \in \{S, I, R\}$; the values of $w_{V_k}$ are the weights for the frequencies of contacts between the different classes of individuals. In particular, if all host species have equal uptake rates ($u_{V_k} = 1$ for all $k$ and $V \in \{S, I, R\}$), then the direct transmission rates have the familiar form $\hat{\beta}_{ij}S_iI_j/\sum_k N_k$.

To show there is an onto mapping of parameterizations of the $n$-habitat ET model (S144) to parameterizations of FDDT models, I will show that one can convert any FDDT model into a well-defined ET model. I start with a FDDT model of the form

$$\begin{aligned}
\frac{dS_i}{dt} &= F_i(\cdot) - m_iS_i - S_i\sum_{j=1}^{n}\frac{\hat{\beta}_{ij}I_j}{\sum_j(w_{i,S_j}S_j + w_{i,I_j}I_j + w_{i,R_j}R_j)} + \gamma_iR_i \\
\frac{dI_i}{dt} &= S_i\sum_{j=1}^{n}\frac{\hat{\beta}_{ij}I_j}{\sum_j(w_{i,S_j}S_j + w_{i,I_j}I_j + w_{i,R_j}R_j)} - (\mu_i + m_i)I_i + \nu_iI_i \\
\frac{dR_i}{dt} &= \nu_iI_i - (\xi_i + m_i)I_i - \gamma_iI_i.
\end{aligned} \tag{S163}$$

The simple way to show the onto mapping is to choose the parameters of the $n$-habitat ET model as $\delta_i = 0$ for all $i$, $\beta_i = 1$, $u_{i,V_j} = w_{i,V_j}/\epsilon$, and $\chi_{ij} = \hat{\beta}_{ij}/\epsilon$ where $0 < \epsilon \ll 1$, and for any parameters values that are shared between the ET model (S144) and the FDDT model (S163), set the values in the ET model equal to those in the FDDT model (e.g., set $\mu_i$ in the ET model equal to the value of $\mu_i$ in the FDDT model). Under these conditions, the ET model (S144) reduces to a FDDT model with parameters identical those in the FDDT model (S163).

When defining host competence for FDDT models with $w_{i,V_j} = 1$, it is useful to define the parameters of the $n$-habitat ET model in the following way. First, set $\delta_i = 0$ for all $i$. Second, for any parameters values that are shared between the ET model (S144) and the DDDT model (S162), set the values in the ET model equal to those in the DDDT model (e.g., set $\mu_i$ in the ET model equal to the value of $\mu_i$ in the DDDT model). Third, decompose the direct transmission parameter as $\hat{\beta}_{ij} = \hat{E}_{ij}\hat{\chi}_j\hat{\rho}_i$ where $\hat{E}_{ij}$ is the encounter rate between susceptible individuals of host $i$ and infected individuals of host $j$, $\hat{\chi}_j$ is the infectious dose that an infected individual of host $j$ exposes a contact to, and $\hat{\rho}_j$ is the probability of infection for host $i$ given exposure. Similarly, decompose the environmental transmission rate as $\beta_i = u_{i,S_i}\rho_i$ where $u_{S_i}$ is the encounter rate between susceptible individuals of host $i$ and infectious propagules in its habitat, and $\rho_i$ is the per infectious propagule probability of infection. Also, decompose the shedding rate as $\chi_{ij} = E_{ij}\chi_j$ where $E_{ij}$ is the contact rate between infected individuals of host $j$ and the habitat of host $i$, and $\chi_j$ is the shedding rate of host $j$. Fourth, set $\chi_j = \hat{\chi}_j/\epsilon$, $E_{ij} = \hat{E}_{ij}$, $\rho_i = \hat{\rho}_j$, $E_i = 1$, and $u_{i,V_j} = 1/\epsilon$ for all $i, j$ and $V \in \{S, I, R\}$. The parameter $\epsilon$ is a free parameter that should be set to a very large value in order to ensure that there is a separation of time scales between the dynamics of the infectious propagules and the host classes. Under these conditions, the ET model (S144) reduces to a FDDT model with parameters identical those in the FDDT model (S163).

#### S2.3 Definitions of host competence, sink, and source

**Definition host competence**: As with the 1-habitat model, I define the competence of host $i$ as its basic reproduction number in the limit of infinite host density, $\mathcal{R}_{0,i}(\infty) = \beta_i\chi_{ii}/u_{S_i}(m_i + \mu_i + \nu_i) = \rho_i\chi_{ii}/(m_i + \mu_i + \nu_i)$; see Section S1.4.

**Definitions of sink and source**: Host $j$ is a source or sink of infectious propagules in habitat $i$ at equilibrium if increases in the density of host $j$ (while holding the frequencies of all host classes constant) cause an increase or decrease, respectively, of infectious propagules in habitat $i$. Mathematically, sink and source are defined by the per capita net production rate of infectious propagules of host $j$ in habitat $i$. Specifically, host $j$ is a source for habitat $i$ if $\varphi_{ij} = (1/Y_j^*)(\partial/\partial N_j)(dP_i dt)|_{p^*} > 0$ and host $j$ is a sink for habitat $i$ if $\varphi_{ij} = (1/Y_j^*)(\partial/\partial N_j)(dP_i/dt)|_{p^*} < 0$. Using the equilibrium conditions for the $n$-habitat NYZ model (S145), the quantity $\varphi_{ij}$ can be written as

$$\varphi_{ij} = \frac{1}{Y_j^*}\frac{\partial}{\partial N_j}\frac{dP_i}{dt}\bigg|_{p^*} = \frac{1}{Y_j^*}\left[\chi_{ij}Y_j^* - (u_{i,S_j}X_i^* + u_{i,I_j}Y_j^* + u_{i,R_j}Z_j^*)P_i^*\right] \tag{S164}$$

$$= \chi_{ij} - \frac{u_{i,S_j}(\mu_j + m_j + \nu_j)}{\beta_j}\frac{P_i^*}{P_j^*} - u_{i,I_j}P_i^* - u_{i,R_j}\Gamma_j P_i^* \tag{S165}$$

where $\Gamma_j = \nu_j/(\xi_j + m_j + \gamma_j)$.

Notes about definitions: One advantage of using the above definitions is that they arise naturally in the sensitivity calculations. A second advantage is that both definitions are independent of the host densities. Thus, one does not need to know the specific make-up of a community in order to determine one host species has higher versus lower competence than another or if one host is smaller or larger source/sink in a particular habitat.

However, both definitions depend on the host's environment because $\mathcal{R}_{0,i}(\infty)$ and $\varphi_{ij}$ depend on the host's natural mortality rate ($m_j$). In addition, the definition of $\varphi_{ij}$ depends on the equilibrium densities of infectious propagules ($P_i^*$ and $P_j^*$). This means that the designations of high versus low competence and sink versus source can differ between habitats. Thus, the definitions allow one to order the hosts in terms of competence and net production of infectious propagules (i.e, sink/source), but those orderings may differ between different environments

**Relationship between competence and sink/source**: The relationship between the definitions of sink/source and host competence can be seen by rewriting the quantity $\varphi_{ij}$ as

$$\varphi_{ij} = \chi_{ij}\left[1 - \frac{1}{\mathcal{R}_{0,j}(\infty)}\frac{u_{i,S_j}P_i^*}{u_{j,S_j}P_j^*}\right] - u_{i,I_j}P_i^* - u_{i,R_j}\Gamma_i P_i^*. \tag{S166}$$

where $\Gamma_i = \nu_i/(\xi_i + m_i + \gamma_i)$.

There are two restrictions on the possible combinations of $\mathcal{R}_{0,i}(\infty)$ and $\varphi_{ii}$ because the quantities defining competence and sink/source are not independent. The restrictions are analogous to those listed in Section S1.3 for the 1-habitat model:

- If the competence of host $i$ is less than 1, then host $i$ is necessarily a sink in its own habitat. The mathematical condition is $\mathcal{R}_{0,i}(\infty) < 1$ implies $\varphi_{ii} < 0$.
- If host $i$ is a source in its own habitat, then the competence of host $i$ must be greater than 1. The mathematical condition is $\varphi_{ii} > 0$ implies $\mathcal{R}_{0,i}(\infty) < 1$.

In contrast, there are no restrictions on the possible combinations of $\mathcal{R}_{0,i}(\infty)$ and $\varphi_{ji}$ because $\mathcal{R}_{0,i}(\infty)$ is independent of $\chi_{ji}$ and $u_{i,V_j}$ ($V \in \{S, I, R\}$). This means that host $i$ can be a source in some habitats and a sink in others.

**Host competence in direct transmission models:** To apply the definition of competence to direct transmission models, one needs to decompose the direct transmission parameter as $\hat{\beta}_{ij} = \hat{E}_{ij}\hat{\chi}_j\hat{\rho}_i$ where $\hat{E}_{ij}$ is the encounter rate between susceptible individuals of host $i$ and infected individuals of host $j$, $\hat{\chi}_j$ is the infectious dose that an infected individual of host $j$ exposes a contact to, and $\hat{\rho}_j$ is the probability of infection for host $i$ given exposure. From this, host competence for direct transmission pathogens is $\mathcal{R}_{0,i}(\infty) = \hat{\rho}_i\hat{\chi}_i/[m_i+\mu_i+\nu_i]$.

**Sink and source in direct transmission models**: The definitions of sink and source also apply to direct transmission models because environmental transmission models. For density-dependent direct transmission (DDDT), all hosts are sources. This is because DDDT corresponds to $u_{i,V_j} = 0$ for all $i, j$ and $V \in \{S, I, R\}$, which means $\varphi_{ij}$ is always positive.

For frequency-dependent direct transmission (FDDT), the hosts species can be sources or sinks in each habitat. As a specific example, consider traditional FDDT rates where $u_{i,V_j} = 1$ and $\delta_i = 0$ for all $i, j$ and $V \in \{S, I, R\}$. Under these conditions, equation (S165) is positive when $\chi_{ij}$ is sufficiently large and negative when $\chi_{ij}$ is sufficiently small. Because the frequency-dependent direct transmission parameters are $\hat{\beta}_{ij} = \chi_{ij}\beta_i$ (see Subsection S2.2), host $j$ is a source when its interspecific transmission rate is sufficiently high and a sink when its interspecific transmission rate is sufficiently low. In general, host $j$ can be a source for some habitats and a sink for others. In many prior studies on FDDT pathogens, low competence hosts species are often sinks for all habitats and high competence hosts species are often source for all habitats. This is because those prior studies assume interspecific transmission rates are related to intraspecific transmission rates (e.g., Dobson (2004) assumes $\hat{\beta}_{ij} = \epsilon(\beta_{ii} + \beta_{jj})/2$), which results in host competence and sink/source being positively correlated.

#### S2.4 Calculations for $\mathcal{R}_0^C$ sensitivities

The Next Generation Matrix (Van den Driessche and Watmough 2008, Diekmann et al. 2010) for the $n$-habitat model (S144) is computed from the submatrix of the Jacobian for the infected class and infectious propagules. When evaluated at the disease-free equilibrium, $p^\ddagger$, the submatrix has the form

$$\mathcal{J} = \begin{bmatrix} -\mathbf{D}_{m+\mu+\nu} & \mathbf{D}_{\beta N^\ddagger} \\ \boldsymbol{\chi} & -\mathbf{D}_{U^\ddagger+\delta} \end{bmatrix} \tag{S167}$$

where $\boldsymbol{\chi}$ is a matrix with entries $\boldsymbol{\chi}_{ij} = \chi_{ij}$, $\mathbf{D}_{m+\mu+\nu}$ is a diagonal matrix with entries $m_i + \mu_i + \nu_i$, $\mathbf{D}_{\beta N^\ddagger}$ is a diagonal matrix with entries $\beta_i N_i^\ddagger$, and $-\mathbf{D}_{U^\ddagger+\delta}$ is diagonal matrix with entries $U_i^\ddagger + \delta_i$ where $U_i^\ddagger = \sum_i u_{i,S_j} N_j^\ddagger$ is the total per infectious propagule uptake rate of infectious propagules in habitat $i$ at the disease-free equilibrium. The submatrix is written as the sum of two matrices,

$$\mathcal{F} = \begin{bmatrix} \mathbf{0} & \mathbf{D}_{\beta N^\ddagger} \\ \mathbf{0} & \mathbf{0} \end{bmatrix}, \quad \mathcal{V} = \begin{bmatrix} -\mathbf{D}_{m+\mu+\nu} & \mathbf{0} \\ \boldsymbol{\chi} & -\mathbf{D}_{U^\ddagger+\delta} \end{bmatrix} \tag{S168}$$

which yields the product

$$\begin{aligned} -\mathcal{F}\mathcal{V}^{-1} &= \begin{bmatrix} \mathbf{0} & \mathbf{D}_{\beta N^\ddagger} \\ \mathbf{0} & \mathbf{0} \end{bmatrix} \begin{bmatrix} \mathbf{D}_{1/(m+\mu+\nu)} & \mathbf{0} \\ \mathbf{D}_{1/(U^\ddagger+\delta)}\boldsymbol{\chi}\mathbf{D}_{1/(m+\mu+\nu)} & \mathbf{D}_{1/(U^\ddagger+\delta)} \end{bmatrix} \\ &= \begin{bmatrix} \mathbf{D}_{\beta N^\ddagger}\mathbf{D}_{1/(U^\ddagger+\delta)}\boldsymbol{\chi}\mathbf{D}_{1/(m+\mu+\nu)} & \mathbf{D}_{\beta N^\ddagger}\mathbf{D}_{1/(U^\ddagger+\delta)} \\ \mathbf{0} & \mathbf{0} \end{bmatrix} \end{aligned} \tag{S169}$$

The community reproduction number is the largest positive eigenvalue of $-\mathcal{F}\mathcal{V}^{-1}$. Because of its block diagonal form, the community reproduction number is computed as

$$\mathcal{R}_0^C = \rho(\mathbf{M}) = \rho(\mathbf{D}_{\beta N^\ddagger}\mathbf{D}_{1/(U^\ddagger+\delta)}\boldsymbol{\chi}\mathbf{D}_{1/(m+\mu+\nu)}) \tag{S170}$$

where $\rho(\mathbf{M})$ is the spectral radius of $\mathbf{M}$.

Recall that I assume there exists a sequence $\beta_{k_1}\chi_{k_1 i}\beta_{k_1}\chi_{k_1 k_2}...\beta_{k_l}\chi_{k_l k_{l-1}}\beta_j\chi_{j k_l} > 0$ for all $i, j$. This means that there are no dead end hosts, and as a result, $\mathcal{R}_0^C$ is always computed using the full matrix $\mathbf{M}$. However, in systems with a dead end host, the community reproductive number is defined by $\mathcal{R}_0^C = \rho(\mathbf{M}^{[\not{j},\not{j}]})$, where $\mathbf{M}^{[\not{j},\not{j}]}$ is the submatrix of $\mathbf{M}$ where column $j$ and row $j$ have been removed.

##### S2.4.1 Method for computing the sensitivities of $\mathcal{R}_0^C$

In general, it is not possible to analytically compute $\mathcal{R}_0^C$. However, the sensitivity of $\mathcal{R}_0^C$ can be computed using the matrix theory developed for discrete-time models of stage-structured populations (Caswell 2019). To see this, note that the leading eigenvalue of matrix $\mathbf{M}$ in equation (S170) is $\mathcal{R}_0^C$, which defines the asymptotic geometric growth rate of the pathogen; the associated right eigenvector, $\vec{x} = (x_1, ..., x_n)$, is the asymptotic distribution of infected individuals across all host species; and the left eigenvector, $\vec{y} = (y_1, ..., y_n)$, is the reproduction value of each species.

To be able to apply the theory from stage-structured models, I need to show that matrix $\mathbf{M}$ is non-negative and irreducible. Matrix $\mathbf{M}$ is non-negative because each entry $\mathbf{M}_{ij} = \chi_{ij}\beta_i N_i^\ddagger/[(m_j+\mu_j+\nu_j)(U_i^\ddagger+\delta_i)]$ is zero when $\chi_{ij} = 0$ or $\beta_i = 0$ and positive otherwise. Matrix $\mathbf{M}$ is irreducible because I assume there exists a sequence $\beta_{k_1}\chi_{k_1 i}\beta_{k_1}\chi_{k_1 k_2}...\beta_{k_l}\chi_{k_l k_{l-1}}\beta_j\chi_{jk_l} \neq 0$ for all $i, j$. This implies there exists a sequence of entries satisfying $\mathbf{M}_{k_1 i}\mathbf{M}_{k_1 k_2}...\mathbf{M}_{k_l k_{l-1}}\mathbf{M}_{jk_l} \neq 0$ for all $i, j$.

As a consequence of $\mathbf{M}$ being non-negative and irreducible, $\mathcal{R}_0^C$ and its eigenvectors satisfy two important properties. First, the eigenvectors $\vec{x}$ and $\vec{y}$ have non-negative entries. In addition, the vectors can be normalized such that $\vec{y}^{\mathrm{T}}\vec{x} = 1$. Unless otherwise stated, throughout this section I assume $\vec{y}^{\mathrm{T}}\vec{x} = 1$. Second, for any entry of $\mathbf{M}$, the derivative of $\mathcal{R}_0^C$ with respect to that entry is

$$\frac{\partial \mathcal{R}_0^C}{\partial \mathbf{M}_{ij}} = \frac{\partial \rho(\mathbf{M})}{\partial \mathbf{M}_{ij}} = y_i x_j > 0. \tag{S171}$$

Moreover, if $\sigma$ is a parameter of the model, then the sensitivity of $\mathcal{R}_0^C$ to the parameter is

$$\frac{\partial \mathcal{R}_0^C}{\partial \sigma} = \sum_{k,l} \frac{\partial \mathcal{R}_0^C}{\partial \mathbf{M}_{kl}} \frac{\partial \mathbf{M}_{kl}}{\partial \sigma} = \sum_{k,l} y_k x_l \frac{\partial \mathbf{M}_{kl}}{\partial \sigma} \tag{S172}$$

where the sum is taken over all entries of matrix $\mathbf{M}$. These properties follow from the Perron-Frobenius Theorem and results about the eigenvalues of irreducible non-negative matrices; mathematical details are proved and discussed in Lemma 3.1 of Deutsch and Neumann (1984) and Section 3.3 of Caswell (2019).

##### S2.4.2 Sensitivity of $\mathcal{R}_0^C$ to host densities

Let $\alpha$ denote any parameter that satisfies $\partial F_n/\partial\alpha < 0$ and $\partial F_j/\partial\alpha = 0$ for $j \neq n$. Examples include parameters defining the level of intraspecific or interspecific competition experienced by host $n$ such as $\alpha_{nn}$ and $\alpha_{nj}$, respectively, in a Lotka-Volterra competition model.

The sensitivity of $\mathcal{R}_0^C$ to the equilibrium density of host $n$ ($N_n^\ddagger$) is computed using the ratio

$$\frac{d\mathcal{R}_0^C}{dN_n^\ddagger} = \frac{\partial \mathcal{R}_0^C}{\partial \alpha} \Big/ \frac{\partial N_n^\ddagger}{\partial \alpha} \tag{S173}$$

which yields

$$\frac{d\mathcal{R}_0^C}{dN_n^\ddagger} = \mathcal{R}_0^C \sum_k \frac{1}{N_k^\ddagger}\frac{dN_k^\ddagger}{dN_n^\ddagger}\left[y_k x_k - \sum_l x_l y_l \frac{u_{l,S_k} N_k^\ddagger}{\delta_l + U_l^\ddagger}\right] \tag{S174}$$

Details about the general calculations and special cases are provided below. The following interpretation is based on the sensitivities of $\mathcal{R}_0^C$ (see below) and its eigenvectors (Section S2.4.3).

**Summary of biological interpretation**: Increased density of host $n$ will lead to increases in $\mathcal{R}_0^C$ if the host community composition changes such that hosts with higher competence and higher shedding rates increase more or decrease less than hosts with lower competence and lower shedding rates. Alternatively, increased density of host $n$ will lead to decreases in $\mathcal{R}_0^C$ if the host community composition changes such that hosts with higher competence and higher shedding rates decrease more or increase less than hosts with lower competence and lower shedding rates.

If there is density-dependent direct transmission and no interspecific competition, then increased density of host $n$ always increases $\mathcal{R}_0^C$. If there is frequency-dependent direct transmission where all host species have equal uptake rates ($u_{V_i} = u$ for all $i$ and $V \in \{S, I, R\}$) and no interspecific competition, then increased density of host $n$ decreases $\mathcal{R}_0^C$ unless the host $n$ has a sufficiently large competence or sufficiently high interspecific transmission rate. If there is frequency-dependent direct transmission where all host species have equal uptake rates ($u_{V_i} = u$ for all $i$ and $V \in \{S, I, R\}$), interspecific transmission is much less than intraspecific transmission, then increases in $n$ are predicted to decrease $\mathcal{R}_0^C$ unless host $n$ has a sufficiently large competence or sufficiently large interspecific transmission rates.

**Calculations for Environmental Transmission**: Before computing the sensitivity of $\mathcal{R}_0^C$ to $\alpha$, it is useful to know

$$\sum_l x_l \frac{\chi_{kl}\beta_k N_k^\ddagger}{(\mu_l + m_l + \nu_l)(\delta_k + U_k^\ddagger)} = \mathcal{R}_0^C x_k \tag{S175}$$

$$\sum_k y_k \frac{\chi_{kl}\beta_k N_k^\ddagger}{(\mu_l + m_l + \nu_l)(\delta_k + U_k^\ddagger)} = \mathcal{R}_0^C y_l \tag{S176}$$

which follow from $\vec{x}$ and $\vec{y}$ being eigenvectors of $\mathbf{M}$. The sensitivity of $\mathcal{R}_0^C$ to $\alpha$ is

$$\frac{\partial \mathcal{R}_0^C}{\partial \alpha} = \sum_{k,l} \frac{\partial \mathcal{R}_0^C}{\partial \mathbf{M}_{kl}} \frac{\partial \mathbf{M}_{kl}}{\partial \alpha} \tag{S177}$$

$$= \sum_{k,l} \frac{\partial \mathcal{R}_0^C}{\partial \mathbf{M}_{kl}} \frac{\partial}{\partial \alpha}\left[\frac{\chi_{kl}\beta_k N_k^\ddagger}{(\mu_l + m_l + \nu_l)(\delta_k + U_k^\ddagger)}\right] \tag{S178}$$

$$= \sum_{k,l} y_k x_l \left[\sum_m \frac{\chi_{kl}\beta_k N_k^\ddagger}{(\mu_l + m_l + \nu_l)(\delta_k + U_k^\ddagger)} \frac{u_{k,S_m}}{(\delta_k + U_k^\ddagger)} \frac{\partial N_m^\ddagger}{\partial \alpha}\right] + \sum_k \frac{\partial N_k^\ddagger}{\partial \alpha} y_k \left[\sum_l x_l \frac{\chi_{kl}\beta_k}{(\mu_l + m_l + \nu_l)(\delta_k + U_k^\ddagger)}\right] \tag{S179}$$

$$= \sum_{k,m} y_k \frac{u_{k,S_m}}{(\delta_k + U_k^\ddagger)} \frac{\partial N_m^\ddagger}{\partial \alpha}\left[\sum_l x_l \frac{\chi_{kl}\beta_k N_k^\ddagger}{(\mu_l + m_l + \nu_l)(\delta_k + U_k^\ddagger)}\right] + \sum_k \frac{\partial N_k^\ddagger}{\partial \alpha} y_k x_k \frac{\mathcal{R}_0^C}{N_k^\ddagger} \tag{S180}$$

$$= -\sum_{k,m} \frac{u_{k,S_m}}{(\delta_k + U_k^\ddagger)} \frac{\partial N_m^\ddagger}{\partial \alpha} \mathcal{R}_0^C x_k y_k + \sum_k \frac{\partial N_k^\ddagger}{\partial \alpha} y_k x_k \frac{\mathcal{R}_0^C}{N_k^\ddagger} \tag{S181}$$

$$= -\sum_{k,l} \frac{u_{l,S_k}}{(\delta_l + U_l^\ddagger)} \frac{\partial N_k^\ddagger}{\partial \alpha} \mathcal{R}_0^C x_l y_l + \sum_k \frac{\partial N_k^\ddagger}{\partial \alpha} y_k x_k \frac{\mathcal{R}_0^C}{N_k^\ddagger} \tag{S182}$$

$$= \mathcal{R}_0^C \sum_k \frac{1}{N_k^\ddagger} \frac{\partial N_k^\ddagger}{\partial \alpha} \left[ y_k x_k - \sum_l x_l y_l \frac{u_{l,S_k} N_k^\ddagger}{\delta_l + U_l^\ddagger} \right] \tag{S183}$$

where the second to last line uses the changes of indices $k \to l$ and $m \to k$. Dividing by $\partial N_n^\ddagger/\partial\alpha$ and simplifying using $dN_j^\ddagger/dN_n^\ddagger = (\partial N_j^\ddagger/\partial\alpha)/(\partial N_n^\ddagger/\partial\alpha)$ yields equation (S174).

The sensitivities of the eigenvectors (Section S2.4.3) suggest that $x_k y_k$ is larger if host $k$ has higher competence and higher rates of shedding into the habitats of other host species. This means each term $y_k x_k - \sum_l x_l y_l u_{l,S_k} N_k^\ddagger/(U_l^\ddagger + \delta_l)$ is more likely to be positive when host $k$ has higher competence, a higher shedding rates, and a lower uptake rate in habitat $l$ relative to the other host species. Conversely, $y_k x_k - \sum_l x_l y_l u_{l,S_k} N_k^\ddagger/(U_l^\ddagger + \delta_l)$ is more likely to be negative when host $k$ has lower competence, a lower shedding rate, and a higher uptake rate in habitat $l$ relative to the other host species. As a result, equation (S174) is positive if increases in host $n$ cause host species with higher competence and higher shedding rates to increase more or decrease less in density than host species with lower competence and lower shedding rates.

**Special Case A: density-dependent direct transmission.** Assume there is density-dependent direct transmission (DDDT), which means $u_{j,V_i} = 0$ for all $i, j$ and $V \in \{S, I, R\}$. Then equation (S174) reduces to

$$\frac{d\mathcal{R}_0^C}{dN_n^\ddagger} = \mathcal{R}_0^C \sum_k \frac{1}{N_k^\ddagger} \frac{dN_k^\ddagger}{dN_n^\ddagger} y_k x_k \tag{S184}$$

If interspecific competition is absent, then $dN_k^\ddagger/dN_n^\ddagger = 0$ for $k \neq n$ and $d\mathcal{R}_0^C/dN_n^\ddagger > 0$. This means increased host richness always increases $\mathcal{R}_0^C$. When interspecific competition is present, $d\mathcal{R}_0^C/dN_n^\ddagger$ is positive unless host $n$ has strong interspecific competitive effects that cause larger decreases in the densities of host species with high competence and large interspecific shedding rates. This means that increased host species richness leads to increases in $\mathcal{R}_0^C$ under DDDT, unless interspecific competition is sufficiently strong that it causes host species with high competence and larger interspecific shedding rates to decrease.

**Special Case B: Frequency-Dependent Direct Transmission with Equal Weights.** Some studies have used models of frequency dependent direct transmission (FDDT) where the transmission rates have the form $\hat{\beta}_{ij} S_i I_j / \sum_j N_j$ (Dobson 2004, Joseph et al. 2013, Mihaljevic et al. 2014, O'Regan et al. 2015, Roberts and Heesterbeek 2018). This corresponds to setting $\delta_i = 0$ and $u_{j,S_i} = 1$ for all $i, j$ in model (S144). In this case, equation (S174) simplifies to

$$\frac{d\mathcal{R}_0^C}{dN_n^\ddagger} = \mathcal{R}_0^C \sum_k \frac{1}{N_k^\ddagger} \frac{dN_k^\ddagger}{dN_n^\ddagger} \left[ x_k y_k - \frac{N_k^\ddagger}{\sum_l N_l^\ddagger} \right]. \tag{S185}$$

If there is no interspecific competition, then $dN_k^\ddagger/dN_n^\ddagger = 0$ for $k \neq i$ and equation (S185) reduces to

$$\frac{d\mathcal{R}_0^C}{dN_n^\ddagger} = \mathcal{R}_0^C \frac{1}{N_n^\ddagger}\left[x_n y_n - \frac{N_n^\ddagger}{\sum_k N_k^\ddagger}\right]. \tag{S186}$$

Equation (S186) implies that if interspecific competition is absent, then increased density of host $n$ decreases $\mathcal{R}_0^C$ unless host $n$ has sufficiently high competence or sufficiently large interspecific transmission rates. Equation (S185) implies that if interspecific competition is present, then increased density of host $n$ decreases $\mathcal{R}_0^C$ unless the densities of hosts with high competence and high interspecific transmission rates increase more (or decrease less) than hosts with low competence and low interspecific transmission rates. This means that increased host species richness leads to decreases in $\mathcal{R}_0^C$ under FDDT unless the added host species have high competence and large interspecific shedding rates or there are shifts in the densities of species in the community such that hosts with high competence and high interspecific transmission rates increase more (or decrease less) than hosts with low competence and low interspecific transmission rates.

**Special Case C: Frequency-Dependent Direct Transmission with Equal Weights and Very Low Interspecific Transmission.** A subset of the studies cited in Special Case B (Dobson 2004, Mihaljevic et al. 2014) also assume interspecific transmission rates are much lower than intraspecific transmission rates; the interspecific transmission coefficients are of the form $\hat{\beta}_{ij} = \epsilon(\hat{\beta}_{ii} + \hat{\beta}_{jj})/2$ where $\epsilon_{ij} \ll 1$. Analytically analyzing this case in general is not possible. However, insight can be gained by analyzing the special case where all hosts species have equivalent traits, $n-1$ host species have the same density, and the density of host $n$ is being varied. For this special case, the sensitivity equations predict that $\mathcal{R}_0^C$ decreases with increased density of host $n$. From this (limited) analysis, I predict that $\mathcal{R}_0^C$ will typically decrease when there is FDDT and interspecific transmission rates are much lower than interspecific transmission rates.

To analyze this special case, I assume $N_k^\ddagger = 1$ for $k \neq n$, all host traits are equal, the interspecific transmission rates are proportional to $\epsilon$, and the total interspecific competitive effects of host $n$ on host $k \neq 1$ are negative (i.e., $dN_k^\ddagger/dN_n^\ddagger$). Under these assumption, all off-diagonal entries of $\mathbf{M}$ are proportional to $\epsilon$ and the last row of terms is proportional to $N_n^\ddagger$. The entries of the non-normalized eigenvectors have orders $(1, ..., 1, N_n^\ddagger \epsilon)$. This means that $x_n y_n \approx (N_n^\ddagger\epsilon)^2/[(N_n^\ddagger\epsilon)^2 + n - 1]$, $x_k y_k \approx 1/[(N_1^\ddagger\epsilon)^2 + n - 1]$ for $k \neq n$, and $N_k^\ddagger/\sum_l N_l^\ddagger = 1/[N_n^\ddagger + n - 1]$ for all $k \neq n$. Rewriting equation (S185) yields

$$\begin{aligned}\frac{d\mathcal{R}_0^C}{dN_n^\ddagger} &= \mathcal{R}_0^C \frac{1}{N_n^\ddagger}\left[x_n y_n - \frac{N_n^\ddagger}{\sum_l N_l^\ddagger}\right] + \mathcal{R}_0^C \sum_{k<n} \frac{1}{N_k^\ddagger}\frac{dN_k^\ddagger}{dN_n^\ddagger}\left[x_k y_k - \frac{N_k^\ddagger}{\sum_l N_l^\ddagger}\right] \\ &\approx \mathcal{R}_0^C \frac{1}{N_n^\ddagger} \underbrace{\left[\frac{(N_n^\ddagger\epsilon)^2}{(N_n^\ddagger\epsilon)^2 + n - 1} - \frac{N_n^\ddagger}{N_n^\ddagger + n - 1}\right]}_{\text{negative}} + \mathcal{R}_0^C \sum_{k\neq 1} \frac{1}{N_k^\ddagger} \underbrace{\frac{dN_k^\ddagger}{dN_n^\ddagger}}_{\text{negative}} \underbrace{\left[\frac{1}{(N_n^\ddagger\epsilon)^2 + n - 1} - \frac{1}{N_n^\ddagger + n - 1}\right]}_{\text{positive}} \\ &< 0\end{aligned} \tag{S187}$$

While equation (S187) was derived from a symmetric case, it suggests that $d\mathcal{R}/dN_n^\ddagger$ will be negative unless there are large asymmetries in the host densities or epidemiological traits (e.g., host $n$ has very high competence). This in turn suggests that host species richness will cause $\mathcal{R}_0^C$ to decrease in systems where there is FDDT and very low rates of interspecific transmission, except possibly where there are large asymmetries in the host densities or epidemiological traits.

Note that the signs of the terms in equation (S187) can switch if $\epsilon$ is sufficiently large. This means that if interspecific transmission is sufficiently higher than intraspecific transmission, then $\mathcal{R}_0^C$ can increase with increased density of host $n$. This suggests that $\mathcal{R}_0^C$ will increase with the addition of host species when there is FDDT and interspecific transmission is greater than intraspecific transmission.

**Special case D: Frequency-Dependent Direct Transmission with Fixed Interspecific Contact Rates.** Consider the case where $u_{i,S_j} = 0$ for all $i \neq j$ and $\delta_i = 0$ for all $i$. This represents the scenario where all infectious propagules in habitat $i$ are taken up by host $i$ (possibly because the overlap in habitats is not large and degradation is low). It also corresponds to an extreme case of FDDT where each host species has a fixed rate of contact with each other host species that does not depend on the densities of the other host species; such a model was analyzed in McCormack and Allen (2007). The sensitivity equation (S174) reduces to

$$\frac{d\mathcal{R}_0^C}{dN_n^\ddagger} = \mathcal{R}_0^C \sum_k \frac{1}{N_k^\ddagger} \frac{dN_k^\ddagger}{dN_n^\ddagger} \left[ y_k x_k - x_k y_k \frac{u_{k,S_k} N_k^\ddagger}{U_k^\ddagger} \right] = \mathcal{R}_0^C \sum_k \frac{1}{N_k^\ddagger} \frac{dN_k^\ddagger}{dN_n^\ddagger} \left[ y_k x_k - x_k y_k \frac{u_{k,S_k} N_k^\ddagger}{u_{k,S_k} N_k^\ddagger} \right] = 0 \tag{S188}$$

Thus, the value of $\mathcal{R}_0^C$ is independent of host density. The sensitivity is valid for any nonzero density of $N_n^\ddagger$; it is not valid at $N_n^\ddagger = 0$ because $\mathcal{R}_0^C$ is discontinuous at that value.

##### S2.4.3 Sensitivity of eigenvectors to epidemiological parameters

The sensitivities of $\mathcal{R}_0^C$ to host parameters and densities depend on the magnitudes of the entries of the right and left eigenvectors for $\mathcal{R}_0^C$. While the eigenvectors cannot be computed analytically, the sensitivities of the eigenvectors to model parameters can be computed. Those sensitivities give some insight into the conditions under which the entries of the eigenvectors are larger and smaller.

The following derives explicit formulas for the sensitivities of the eigenvectors. Those formulas are too complex to interpret in general. Thus, I focus on interpreting the formulas in the special case where all host species have equivalent parameter values and densities. This special case provides insight (albeit limited) in how the magnitudes of the entries of the eigenvalues depend on the host characteristics.

The calculations primarily focus on the sensitivities for the right eigenvector ($\vec{x}$); the sensitivities for the left eigenvector ($\vec{y}$) for $\mathcal{R}_0^C$ are computed in a similar way using the transpose of $\mathbf{M}$.

**Summary of biological interpretation**: For host $k$, $x_k y_k$ is predicted to be larger in magnitude when host $k$ has a larger transmission parameter ($\beta_k$), larger shedding rates ($\chi_{jk}$), lower mortality and recover rates ($\mu_k$, $\nu_k$), and lower uptake rates in its own habitat ($u_{k,S_k}$). This corresponds to ost $k$ having higher competence and higher intraspecific and interspecific transmission rates. Note that these predictions are based on the special case where all host species are equivalent.

**Deriving the formulas for the sensitivities**: Let $\lambda_i$ be the eigenvalues of $\mathbf{M}$ and $\vec{v}_i$ be the associated eigenvalues normalized such that $||\vec{v}_i||_1 = \vec{1}^{\mathrm{T}}\vec{v}_i = 1$, where $\vec{1}$ is a vector of ones and superscript T denotes the transpose. Without loss of generality, assume $\lambda_1 = \mathcal{R}_0^C$ and $\vec{v}_1 = \vec{x} = [x_1, x_2, ..., x_n]^{\mathrm{T}}$. Let $\partial\mathbf{M}/\partial\mathbf{M}_{ij} = \vec{e}_i\vec{e}_j^{\mathrm{T}}$ be the matrix defining the derivative of each entry of $\mathbf{M}$ with respect entry $i, j$. Here, $\vec{e}_i$ is the unit vector with a 1 in the $i$th entry and zeros elsewhere. Let $\partial\vec{x}/\partial\mathbf{M}_{ij}$ denote the vector of derivatives defining how each entry of $\vec{x}$ changes with small changes in entry $\mathbf{M}_{ij}$. Using equation (10.71) of Caswell (2019), the relations $\mathbf{M}\vec{x} = \mathcal{R}_0^C\vec{x}$ and $\vec{1}^{\mathrm{T}}\mathbf{M}\vec{x} = \mathcal{R}_0^C$, and the chain rule, the sensitivity of $\vec{v}_1$ to matrix entry $\mathbf{M}_{ij}$ satisfies

$$\frac{\partial\vec{x}}{\partial\mathbf{M}_{ij}} = \frac{1}{\mathcal{R}_0^{C2}}\left[\mathcal{R}_0^C\frac{\partial\mathbf{M}}{\partial\mathbf{M}_{ij}}\vec{x} + \mathcal{R}_0^C\mathbf{M}\frac{\partial\vec{x}}{\partial\mathbf{M}_{ij}} - \mathcal{R}_0^C\vec{x}\vec{1}^{\mathrm{T}}\frac{\partial\mathbf{M}}{\partial\mathbf{M}_{ij}}\vec{x} - \mathcal{R}_0^C\vec{x}\vec{1}^{\mathrm{T}}\mathbf{M}\frac{\partial\vec{x}}{\partial\mathbf{M}_{ij}}\right]. \tag{S189}$$

Rearranging the equation yields

$$\left[\mathcal{R}_0^C\mathbf{I} - \mathbf{M} + \vec{x}\vec{1}^{\mathrm{T}}\mathbf{M}\right]\frac{\partial\vec{x}}{\partial\mathbf{M}_{ij}} = \left[\mathbf{I} - \vec{x}\vec{1}^{\mathrm{T}}\right]\vec{e}_i\vec{e}_j^{\mathrm{T}}\vec{x} \tag{S190}$$

where $\mathbf{I}$ is the $n \times n$ identity matrix. The previous equation can be solved by simplifying the right hand side to $(\vec{e}_i - \vec{x})\vec{x}_j$ and multiplying by the inverse of $\mathbf{K} = \left[\mathcal{R}_0^C\mathbf{I} - \mathbf{M} + \vec{x}\vec{1}^{\mathrm{T}}\mathbf{M}\right]$. The chain rule yields the sensitivity of the right eigenvector to a parameter $\sigma$,

$$\frac{\partial\vec{x}}{\partial\sigma} = \sum_{ij}\left[\mathcal{R}_0^C\mathbf{I} - \mathbf{M} + \vec{x}\vec{1}^{\mathrm{T}}\mathbf{M}\right]^{-1}(\vec{e}_i - \vec{x})x_j\frac{\partial\mathbf{M}_{ij}}{\partial\sigma}. \tag{S191}$$

The previous equation is equivalent to equation (10.73) in Caswell (2019). To gain a little more biological insight, I now derive an explicit formula for the inverse of $\mathbf{K}$.

My approach is to compute the diagonalization $-\mathbf{M}+\vec{x}\vec{1}^{\mathrm{T}}\mathbf{M} = \mathbf{Q}\boldsymbol{\Lambda}\mathbf{Q}^{-1}$ and then compute the inverse of $\mathbf{K}$ as $\mathbf{K}^{-1} = (-\mathbf{M} + \vec{x}\vec{1}^{\mathrm{T}}\mathbf{M} + \mathcal{R}_0^C\mathbf{I})^{-1} = \mathbf{Q}(\Lambda + \mathcal{R}_0^C\mathbf{I})^{-1}\mathbf{Q}^{-1}$. To compute the diagonalization, note that if $\mathbf{M}\vec{v}_i = \lambda_i\vec{v}_i$, then $(-\mathbf{M} + \vec{x}\vec{1}^{\mathrm{T}}\mathbf{M})\vec{v}_i = -\lambda_i\vec{v}_i + \lambda_i\vec{x}$. This means that the eigenvalues and eigenvectors of $-\mathbf{M}+\vec{x}\vec{1}^{\mathrm{T}}\mathbf{M}$ are $0, -\lambda_2, ..., -\lambda_n$ and $\vec{x}, \vec{v}_2-\vec{x}, ..., \vec{v}_n-\vec{x}$, respectively. Thus, $\mathbf{Q} = [\vec{x}, \vec{v}_2 - \vec{x}, ..., \vec{v}_n - \vec{x}]$ and $(\Lambda + \mathcal{R}_0^C\mathbf{I})^{-1}$ is a diagonal matrix with diagonal entries $(\mathcal{R}_0^C)^{-1}, (\mathcal{R}_0^C - \lambda_2)^{-1}, ..., (\mathcal{R}_0^C - \lambda_n)^{-1}$.

I now apply the above to equation (S191). First, I compute $\mathbf{K}^{-1}\vec{x} = \vec{x}/\mathcal{R}_0$. Second, I write each unit vector as a linear combination of the eigenvectors of $\mathbf{M}$, $\vec{e}_i = \sum_k c_k^i\vec{v}_k$. Then each unit vector can be written as linear combination of the eigenvectors of $\mathbf{K}$, $\vec{e}_i = \sum_{k=1}^n c_k^i\vec{x} + \sum_{k=2}^n c_k^i(\vec{v}_k - \vec{x})$. Computing $\mathbf{K}^{-1}\vec{e}_i$ yields

$$\mathbf{K}^{-1}\vec{e}_i = \frac{1}{\mathcal{R}_0^C}\sum_{k=1}^n c_k^i\vec{x} + \sum_{k=2}^n\frac{c_k^i}{\mathcal{R}_0^C - \lambda_k}(\vec{v}_k - \vec{x}) \tag{S192}$$

$$= \frac{1}{\mathcal{R}_0^C}\left(c_1^i - \sum_{k=2}^{n} \frac{c_k^i \lambda_k}{\mathcal{R}_0^C - \lambda_k}\right)\vec{x} + \sum_{k=2}^{n} \frac{c_k^i}{\mathcal{R}_0^C - \lambda_k}\vec{v}_k. \tag{S193}$$

Finally, combining the above, substituting into equation (S191), and collecting like terms yields

$$\frac{\partial \vec{x}}{\partial \sigma} = \sum_{ij}\left[\frac{1}{\mathcal{R}_0^C}\left(c_1^i - 1 - \sum_{k=2}^{n} \frac{c_k^i \lambda_k}{\mathcal{R}_0^C - \lambda_k}\right)\vec{x} + \sum_{k=2}^{n} \frac{c_k^i}{\mathcal{R}_0^C - \lambda_k}\vec{v}_k\right] x_j \frac{\mathbf{M}_{ij}}{\partial \sigma}. \tag{S194}$$

**Special case of all host species are equivalent**: Consider the special case where all host species are equivalent in terms of their densities and ecological and epidemiological parameters. The sensitivities for this case determine how small asymmetries in the species epidemiological parameters affect the magnitudes of the entries of $\vec{x}$ and $\vec{y}$. Mathematically, I assume all host species have identical equilibrium densities ($N_1^\ddagger = ... = N_n^\ddagger$) and epidemiological parameters ($\beta_{ij} = \beta$, $\mu_i = \mu$, $\chi_i = \chi$, etc). Consequently, $\mathbf{M}$ is a matrix with identical entries ($\mathbf{M}_{ij} = M$), which means that $\mathcal{R}_0^C = nM$, $\vec{x} = \vec{y} = [1/n, ..., 1/n]^{\mathrm{T}}$, and $\lambda_i = 0$ for $i \geq 2$. This simplifies equation (S194) and the corresponding equation for $\vec{y}$ to

$$\frac{\partial \vec{x}}{\partial \sigma} = \sum_{ij}\left[\frac{1}{\mathcal{R}_0^C}\left(c_1^i - 1\right)\vec{x} + \sum_{k=2}^{n} \frac{c_k^i}{\mathcal{R}_0^C}\vec{v}_k\right] x_j \frac{\mathbf{M}_{ij}}{\partial \sigma} = \sum_{ij}\left[\frac{-1}{\mathcal{R}_0^C}\vec{x} + \sum_{k=1}^{n} \frac{c_k^i}{\mathcal{R}_0^C}\vec{v}_k\right] x_j \frac{\mathbf{M}_{ij}}{\partial \sigma} \tag{S195}$$

$$= \sum_{ij} \frac{1}{\mathcal{R}_0^C}\left(\vec{e}_i - \vec{x}\right) x_j \frac{\mathbf{M}_{ij}}{\partial \sigma} \tag{S196}$$

$$\frac{\partial \vec{y}}{\partial \sigma} = \sum_{ij} \frac{1}{\mathcal{R}_0^C}\left(\vec{e}_j - \vec{y}\right) y_i \frac{\mathbf{M}_{ij}}{\partial \sigma} \tag{S197}$$

Recall that $\mathbf{M}_{ij} = \chi_{ij}\beta_i N_i^\ddagger / [(m_j + \mu_j + \nu_j)(U_i^\ddagger + \delta_i)]$. Computing the sensitivity of the right eigenvector with respect to each parameter yields

$$\frac{\partial \vec{x}}{\partial \beta_i} = \sum_{j} \frac{1}{\mathcal{R}_0^C}\left(\vec{e}_i - \vec{x}\right) x_j \frac{\chi_{ij} N_i^\ddagger}{(m_j + \mu_j + \nu_j)(U_i^\ddagger + \delta_i)} = (\vec{e}_i - \vec{x})\frac{x_i}{\beta_i} \tag{S198}$$

$$\frac{\partial \vec{x}}{\partial \chi_{ji}} = \frac{1}{\mathcal{R}_0^C}\left(\vec{e}_j - \vec{x}\right) x_i \frac{\beta_j N_j^\ddagger}{(m_i + \mu_i + \nu_i)(U_j^\ddagger + \delta_j)} \tag{S199}$$

$$\frac{\partial \vec{x}}{\partial \mu_i} = \sum_{k} \frac{1}{\mathcal{R}_0^C}\left(\vec{e}_k - \vec{x}\right) x_i \frac{-\chi_{ki}\beta_k N_k^\ddagger}{(m_i + \mu_i + \nu_i)^2(U_k^\ddagger + \delta_k)} = 0 \tag{S200}$$

$$\frac{\partial \vec{x}}{\partial \nu_i} = \sum_{k} \frac{1}{\mathcal{R}_0^C}\left(\vec{e}_k - \vec{x}\right) x_i \frac{-\chi_{ki}\beta_k N_k^\ddagger}{(m_i + \mu_i + \nu_i)^2(U_k^\ddagger + \delta_k)} = 0 \tag{S201}$$

$$\frac{\partial \vec{x}}{\partial u_{i,S_k}} = \sum_{j} \frac{1}{\mathcal{R}_0^C}\left(\vec{e}_i - \vec{x}\right) x_j \frac{-\chi_{ij}\beta_i N_i^\ddagger N_k^\ddagger}{(m_j + \mu_j + \nu_j)(U_i^\ddagger + \delta_i)^2} = \left(\vec{e}_i - \vec{x}\right)\frac{-N_k^\ddagger}{U_i^\ddagger + \delta_i} \tag{S202}$$

The sums simplify because $\vec{y} = \vec{x} = (1/n, ..., 1/n)$, $\sum_k \vec{e}_k - \vec{x} = 0$, $\mathcal{R}_0^C = nM$, and all densities and parameters are assumed to be equal. Computing the sensitivity of the left eigenvector with respect to each parameter yields

$$\frac{\partial \vec{y}}{\partial \beta_i} = \sum_j \frac{1}{\mathcal{R}_0^C} (\vec{e}_i - \vec{y})\, x_j \frac{\chi_{ij} N_i^{\ddagger}}{(m_j + \mu_j + \nu_j)(U_i^{\ddagger} + \delta_i)} = (\vec{e}_i - \vec{y})\, x_k \tag{S203}$$

$$\frac{\partial \vec{y}}{\partial \chi_{ji}} = \frac{1}{\mathcal{R}_0^C} (\vec{e}_i - \vec{y})\, y_j \frac{\beta_j N_j^{\ddagger}}{(m_i + \mu_i + \nu_i)(U_j^{\ddagger} + \delta_j)} \tag{S204}$$

$$\frac{\partial \vec{y}}{\partial \mu_i} = \sum_j \frac{1}{\mathcal{R}_0^C} (\vec{e}_i - \vec{y})\, y_j \frac{-\chi_{ji}\beta_j N_j^{\ddagger}}{(m_i + \mu_i + \nu_i)^2(U_j^{\ddagger} + \delta_j)} = (\vec{e}_i - \vec{y}) \frac{-y_i}{(m_i + \mu_i + \nu_i)} \tag{S205}$$

$$\frac{\partial \vec{y}}{\partial \nu_i} = \sum_j \frac{1}{\mathcal{R}_0^C} (\vec{e}_i - \vec{y})\, y_j \frac{-\chi_{ji}\beta_j N_j^{\ddagger}}{(m_i + \mu_i + \nu_i)^2(U_j^{\ddagger} + \delta_j)} = (\vec{e}_i - \vec{y}) \frac{-y_i}{(m_i + \mu_i + \nu_i)} \tag{S206}$$

$$\frac{\partial \vec{y}}{\partial u_{i,S_k}} = \sum_k \frac{1}{\mathcal{R}_0^C} (\vec{e}_j - \vec{y})\, y_i \frac{-\beta_i \chi_{ij} N_i^{\ddagger} N_k^{\ddagger}}{(m_j + \mu_j + \nu_j)(U_i^{\ddagger} + \delta_i)^2} = 0 \tag{S207}$$

1594 These equations were simplified in the same way.

Using the above equations, one can show that the derivatives of $x_i y_i$ satisfy

$$\begin{aligned} &\frac{\partial x_i y_i}{\partial \beta_i} \geq 0, \quad \frac{\partial x_i y_i}{\partial \chi_{ij}} \geq 0 \\ &\frac{\partial x_i y_i}{\partial \mu_i} \leq 0, \quad \frac{\partial x_i y_i}{\partial \nu_i} \leq 0, \quad \frac{\partial x_i y_i}{\partial u_{i,S_j}} \leq 0 \end{aligned} \tag{S208}$$

The proof for the derivative with respect to $\chi_{ii}$ is

$$\begin{aligned} \frac{\partial (x_i y_i)}{\partial \chi_{ii}} &= y_i \frac{\partial x_i}{\partial \chi_{ii}} + x_i \frac{\partial y_i}{\partial \chi_{ii}} \\ &= \frac{y_i}{\mathcal{R}_0^C} (1 - x_i)\, x_i \frac{\beta_i N_i^{\ddagger}}{(m_i + \mu_i + \nu_i)(U_i^{\ddagger} + \delta_i)} + \frac{x_i}{\mathcal{R}_0^C} (1 - y_i)\, y_i \frac{\beta_i N_i^{\ddagger}}{(m_i + \mu_i + \nu_i)(U_i^{\ddagger} + \delta_i)} \geq 0 \end{aligned}$$

where $x_i = 1/n = y_i$ is used. The calculation for $\partial x_i y_i / \partial \beta_i$ is nearly identical. The proof for the derivative with respect to $\chi_{ji}$ $(i \neq j)$ is

$$\begin{aligned} \frac{\partial (x_i y_i)}{\partial \chi_{ji}} &= y_i \frac{\partial x_i}{\partial \chi_{ji}} + x_i \frac{\partial y_i}{\partial \chi_{ji}} \quad (i \neq j, n \geq 2) \\ &= \frac{y_i}{\mathcal{R}_0^C} (0 - x_i)\, x_i \frac{N_j^{\ddagger}\beta_i}{(m_i + \mu_i + \nu_i)(U_j^{\ddagger} + \delta_j)} + \frac{x_i}{\mathcal{R}_0^C} (1 - y_i)\, y_i \frac{N_j^{\ddagger}\beta_j}{(m_i + \mu_i + \nu_i)(U_j^{\ddagger} + \delta_j)} \\ &= \frac{1}{\mathcal{R}_0^C} \frac{N_j^{\ddagger}\beta_j}{(m_i + \mu_i + \nu_i)(U_j^{\ddagger} + \delta_j)} (-x_i x_i y_i + x_i(1 - y_i) y_i) \\ &= \frac{1}{\mathcal{R}_0^C} \frac{N_j^{\ddagger}\beta_j}{(m_i + \mu_i + \nu_i)(U_j^{\ddagger} + \delta_j)} \left( \frac{1}{n^2} - \frac{2}{n^3} \right) \geq 0 \end{aligned}$$

where $x_i = 1/n = y_i$ is used. The calculation for $\partial x_i y_i / \partial \beta_j$ $(j \neq i)$ is nearly identical. The proofs for the derivatives with respect to $\mu_i$, $\nu_i$, and $u_{i,S_j}$ are immediately clear after applying the product rule.

Based on the above, I predict that the $x_i y_i$ will be larger in magnitude when host $i$ has higher competence and higher shedding rates into the habitats of heterospecifics. Specifically, $x_i y_i$ is larger in magnitude when host $i$ has a higher transmission rate ($\beta_i$), higher shedding rates ($\chi_{ji}$), lower mortality ($\mu_i$) and recovery ($nu_i$) rates, and lower uptake rates in its own habitat ($u_{i,S_i}$).

##### S2.4.4 Sensitivity of $\mathcal{R}_0^C$ to epidemiological parameters

It is straightforward to show for all $i, j, k, l$ that $\partial \mathbf{M}_{kl} / \partial \sigma$ is non-negative for $\sigma \in \{\beta_{ij}, \chi_i\}$ and non-positive for $\sigma \in \{\delta_i, u^j_{S_i}, \mu_i, \nu_i\}$. Because $\vec{x}$ and $\vec{y}$ have non-negative entries, the sensitivities satisfy

$$\begin{aligned} &\frac{\partial \mathcal{R}_0^C}{\partial \beta_i} \geq 0, \quad \frac{\partial \mathcal{R}_0^C}{\partial \chi_{ij}} \geq 0 \\ &\frac{\partial \mathcal{R}_0^C}{\partial \delta_i} \leq 0, \quad \frac{\partial \mathcal{R}_0^C}{\partial \mu_i} \leq 0, \quad \frac{\partial \mathcal{R}_0^C}{\partial \nu_i} \leq 0, \quad \frac{\partial \mathcal{R}_0^C}{\partial u_{i,S_j}} \leq 0 \end{aligned} \tag{S209}$$

for all $i, j$.

**Summary of biological interpretation**: Increases in the competence of a host species or its shedding rates into the habitats of heterospecifics always increase the community reproduction number. Decreases in the competence of a host species or its shedding rates into the habitats of heterospecifics always decrease the community reproduction number. Mathematically, the community reproduction number decreases with larger $\beta_i$ and $\chi_{ji}$ and smaller $\mu_i$, $\nu_i$, and $u_{j,S_i}$.

##### S2.4.5 Sensitivity of $\mathcal{R}_0^C$ to transmission mode

Here, I use a continuous change of parameters to convert the $n$-habitat ET model (S144) from a FDDT form ($\delta_i = 0$ for all $i$) to a DDDT form ($u_{i,V_j} = 0$ for $V \in \{S, I, R\}$). The change of parameters holds all parameters constant except $\delta_i$ and $u_{i,V_j}$, which must necessarily change. The specific change of parameters defined below is chosen such that the community reproduction number for a particular assemblage of hosts species is held constant. The biological interpretation of this constraint is that the pathogen's reproduction number is known for a particular assemblage of species, the pathogen's transmission mode is unknown, and the pathogen's reproduction number is unknown when a set of hosts are added to or lost from the community. This method allows one to predict how the value of $\mathcal{R}_0^C$ in the new assemblage of host species depends on the pathogen transmission mode. Specifically, it identifies if greater increases (or smaller decreases) in $\mathcal{R}_0^C$ occur under DDDT or FDDT.

Note that in order to apply the following theory, one must know/assume the ratio of the uptake rates for all host species for each habitat, i.e., $u_{i,S_j}/u_{i,S_k}$. That information is needed to define the change of parameters functions. While FDDT studies typically use models corresponding to $u_{i,v_j} = 1$ and $\delta_i = 0$ (i.e., transmission rates of the form $\hat{\beta}_{ij} S_i I_j / \Sigma_k N_k$), my analysis allows for unequal uptake rates, which corresponds to FDDT models with weighted frequencies, e.g., $\hat{\beta}_{ij} S_i I_j / \Sigma_k \omega_k N_k$.

**Summary of biological interpretation**: When host species are added to a community, reductions in $\mathcal{R}_0^C$ are more likely to occur under FDDT than DDDT when added host species are weak interspecific and intraspecific competitors and added host species have higher uptake rates. In comparison, reductions in $\mathcal{R}_0^C$ are more likely to occur under DDDT than FDDT when added host species are strong intraspecific and interspecific competitors and added host species have lower uptake rates. In addition, in the absence of interspecific competition, reductions in $\mathcal{R}_0^C$ are always more likely under FDDT than DDDT. The justification for this interpretation is given after the calculations.

**Additions of host species to a community**: Without loss of generality, assume host species $1, ..., m$ $(m < n)$ are present in a community. Assume the species stably coexist at a disease-free equilibrium (hereafter the 'subcommunity equilibrium'). Let $N_i^\dagger$ denote the density of host $i$ at the subcommunity equilibrium, $\mathbf{M}_{jk}^\dagger$ $(j, k \leq m)$ denote the entries of the next generation matrix evaluated at the subcommunity equilibrium, $\mathcal{R}_0^\dagger$ denote the pathogen's reproduction number at the subcommunity equilibrium, and $U_i^\dagger = \sum_{j=1}^m u_{i,S_j} N_j^\dagger$ denote the per capita rate at which infectious propagules shed by host $i$ are taken up by host species $1, ..., m$ at the subcommunity equilibrium.

The infectious propagule equations are rewritten as

$$\frac{dP_i}{dt} = \sum_j \chi_{ij} I_j - g_i(q) \sum_{j=1}^n \left(u_{i,S_j} S_i + u_{i,I_j} I_i + u_{i,R_j} R_i\right) P_i - h_i(q) \delta_i P_i \tag{S210}$$

where the change of parameters functions are defined by

$$g_i(q) = \begin{cases} \dfrac{-\delta_i}{U_i^\dagger} q + 1 + \dfrac{\delta_i}{U_i^\dagger}, & 0 \leq q \leq 1 \\ 2 - q, & 1 \leq q \leq 2 \end{cases} \tag{S211}$$

$$h_i(q) = \begin{cases} q, & 0 \leq q \leq 1 \\ \dfrac{-U_i^\dagger}{\delta_i}(2 - q) + 1 + \dfrac{U_i^\dagger}{\delta_i}, & 1 \leq q \leq 2 \end{cases} \tag{S212}$$

These functions transform the ET model from a FDDT form when $q = 0$ to a DDDT form when $q = 2$. The transformation holds constant the value of the pathogen's community reproduction number for the $m$-host subcommunity ($\mathcal{R}_0^\dagger$) because the entries of the next generation matrix for the $m$-host subcommunity ($\mathbf{M}_{jk}^\dagger$, $1 \leq j, k \leq m$) are independent of $q$,

$$\mathbf{M}_{jk}^\dagger = \frac{N_k^\dagger \beta_{kl} \chi_l}{(m_l + \mu_l + \nu_l)(g_l(q) U_l^\dagger + h_l(q) \delta_l)} = \frac{N_k^\dagger \beta_{kl} \chi_l}{(m_l + \mu_l + \nu_l)(U_l^\dagger + \delta_l)}. \tag{S213}$$

The sensitivity of $\mathcal{R}_0^C$ to $q$ must be computed in two pieces because of the non-differentiable point at $q = 1$. While not explicitly denoted in the following, all derivatives evaluated at

$q = 1$ are treated as left or right derivatives. The sensitivities of $\mathcal{R}_0^C$ to $q$ are

$$0 \le q \le 1: \quad \frac{\partial \mathcal{R}_0^C}{\partial q} = \sum_{1\le k,l\le n} y_k x_l \frac{\partial \mathbf{M}_{kl}}{\partial q} \tag{S214}$$

$$= \sum_{1\le k,l\le n} y_k x_l \frac{-\chi_{kl}\beta_k N_k^\ddagger}{(\mu_l + m_l + \nu_l)(g_k U_k^\ddagger + h_k\delta_k)} \frac{\delta_k - \delta_k U_k^\ddagger / U_k^\dagger}{(g_k U_k^\ddagger + h_k\delta_k)} \tag{S215}$$

$$= \sum_{1\le k\le n} x_k y_k \mathcal{R}_0^C \frac{\delta_k}{U_k^\dagger} \frac{U_k^\ddagger - U_k^\dagger}{(g_k U_k^\ddagger + h_k\delta_k)} \tag{S216}$$

$$1 \le q \le 2: \quad \frac{\partial \mathcal{R}_0^C}{\partial q} = \sum_{1\le k,l\le n} y_k x_l \frac{\partial \mathbf{M}_{kl}}{\partial q} \tag{S217}$$

$$= \sum_{1\le k,l\le n} y_k x_l \frac{-\chi_{kl}\beta_k N_k^\ddagger}{(\mu_l + m_l + \nu_l)(g_k U_k^\ddagger + h_k\delta_k)} \frac{-U_k^\ddagger + U_k^\dagger}{(g_k U_k^\ddagger + h_k\delta_k)} \tag{S218}$$

$$= \sum_{1\le k\le n} x_k y_k \mathcal{R}_0^C \frac{U_k^\ddagger - U_k^\dagger}{(g_k U_k^\ddagger + h_k\delta_k)} \tag{S219}$$

where equations (S216) and (S219) were simplified using equation (S175).

**Losses of host species in a community**: The notation is the same as the above, where daggers denote quantities associated with the subcommunity equilibrium for host species $i = 1, ..., m$. The infectious propagule equation is rewritten as in equation (S210) where the functions $g_i(q)$ and $h_i(q)$ are now defined as

$$g_i(q) = \begin{cases} \dfrac{-\delta_i}{U_i^\ddagger} q + 1 + \dfrac{\delta_i}{U_i^\ddagger}, & 0 \le q \le 1 \\ 2 - q, & 1 \le q \le 2 \end{cases} \tag{S220}$$

$$h_i(q) = \begin{cases} q, & 0 \le q \le 1 \\ \dfrac{-U_i^\ddagger}{\delta_i}(2 - q) + 1 + \dfrac{U_i^\ddagger}{\delta_i}, & 1 \le q \le 2 \end{cases} \tag{S221}$$

Importantly, these functions transform the ET model from a FDDT form when $q = 0$ to a DDDT form when $q = 2$ while holding constant the value of the pathogen's reproduction number for the $n$-host community ($\mathcal{R}_0^C$). This is because the entries of the next generation matrix for the $n$-host community ($\mathbf{M}_{ij}$, $1 \le i, j \le n$) are independent of $q$,

$$\mathbf{M}_{ij} = \frac{\chi_{ij}\beta_i N_i^\ddagger}{(m_j + \mu_j + \nu_j)(g_i(q) U_i^\ddagger + h_i(q)\delta_i)} = \frac{\chi_{ij}\beta_i N_i^\ddagger}{(m_j + \mu_j + \nu_j)(U_i^\ddagger + \delta_i)}. \tag{S222}$$

As before, the sensitivity of $\mathcal{R}_0^\dagger$ to $q$ must be computed in two pieces because of the non-differentiable point at $q = 1$. While not explicitly denoted in the following, all derivatives evaluated at $q = 1$ are treated as left or right derivatives. The sensitivities of $\mathcal{R}_0^\dagger$ to $q$ are

$$0 \le q \le 1: \quad \frac{\partial \mathcal{R}_0^\dagger}{\partial q} = \sum_{1\le k,l\le n} y_k^\dagger x_l^\dagger \frac{\partial \mathbf{M}_{kl}^\dagger}{\partial q} \tag{S223}$$

$$= \sum_{1 \le k,l \le m} y_k^\dagger x_l^\dagger \frac{-\chi_{kl}\beta_k N_k^\dagger}{(\mu_l + m_l + \nu_l)(g_k U_k^\dagger + h_k \delta_k)} \frac{\delta_k - \delta_k U_k^\dagger / U_k^\ddagger}{(g_k U_k^\dagger + h_k \delta_k)} \tag{S224}$$

$$= \sum_{1 \le k \le m} x_k^\dagger y_k^\dagger \mathcal{R}_0^\dagger \frac{\delta_k}{U_k^\ddagger} \frac{-(U_k^\ddagger - U_k^\dagger)}{(g_k U_k^\dagger + h_k \delta_k)} \tag{S225}$$

$$1 \le q \le 2: \quad \frac{\partial \mathcal{R}_0^\dagger}{\partial q} = \sum_{1 \le k,l \le n} y_k^\dagger x_l^\dagger \frac{\partial \mathbf{M}_{kl}^\dagger}{\partial q} \tag{S226}$$

$$= \sum_{1 \le k,l \le m} y_k^\dagger x_l^\dagger \frac{-\chi_{kl}\beta_k N_k^\dagger}{(\mu_l + m_l + \nu_l)(g_k U_k^\dagger + h_k \delta_k)} \frac{-U_k^\dagger + U_k^\ddagger}{(g_k U_k^\dagger + h_k \delta_k)} \tag{S227}$$

$$= \sum_{1 \le k \le m} x_k^\dagger y_k^\dagger \mathcal{R}_0^\dagger \frac{-(U_k^\ddagger - U_k^\dagger)}{(g_k U_k^\ddagger + h_k \delta_k)} \tag{S228}$$

where the equations are simplified using $\mathbf{M}^\dagger \vec{x}^\dagger = \mathcal{R}_0^\dagger \vec{x}^\dagger$ and $(\vec{y}^\dagger)^{\mathrm{T}} \mathbf{M}^\dagger = \mathcal{R}_0^\dagger (\vec{y}^\dagger)^{\mathrm{T}}$. Importantly, the signs of each term in both derivatives are determined by $-(U_k^\ddagger - U_k^\dagger)$.

**Justification for interpretation**: To avoid confusion, I will only focus on equations (S216) and (S219); the interpretation is qualitatively similar for equations (S225) and (S228).

The signs of equations (S216) and (S219) are determined by $U_k^\ddagger - U_k^\dagger$, which can be written as

$$U_k^\ddagger - U_k^\dagger = \sum_{l > m} u_{k,S_l} N_l^\ddagger + \sum_{l \le m} u_{k,S_l} (N_l^\ddagger - N_l^\dagger). \tag{S229}$$

The quantity $U_k^\ddagger - U_k^\dagger$ is more likely to be positive when (i) added host densities are high, meaning added host are weak intraspecific competitors, (ii) resident host densities are higher in the presence of the added host species ($N_l^\ddagger - N_l^\dagger$ positive or negative and small in magnitude for $l \le m$), meaning added hosts are weak interspecific competitors, and (iii) resident host species have lower uptake rates than added host species in the habitats of resident host species ($u_{k,S_l}$ is smaller for $l \le m$ than $l > m$). The quantity $U_i^* - U_i'$ can be negative if (iv) added host densities are low, meaning added hosts are strong intraspecific competitors, (v) resident host densities are higher in the absence of the added host species ($N_l^\ddagger - N_l^\dagger < 0$, $l \le m$), meaning added hosts are strong interspecific competitors, and (vi) resident host species have higher uptake rates than added host species in the habitats of resident host species ($u_{k,S_l}$ is larger for $l \le m$ than $l > m$).

When equations (S216) and (S219) are positive, additions of host species to a community cause larger decreases (or smaller increases) in $\mathcal{R}_0^C$ under FDDT than DDDT. Thus, additions of host species to a community are more likely to cause larger decreases (or smaller increases) in $\mathcal{R}_0^C$ under FDDT than DDDT when the added host species are (i) weak intraspecific interactions, (ii) weak interspecific competitors, and (iii) have lower uptake rates than resident host species in the resident host habitats. In comparison, when equations (S216) and (S219) are negative, additions of host species to a community cause larger increases (or smaller decreases) in $\mathcal{R}_0^C$ under FDDT than DDDT. Thus, additions of host species to a community are more likely to cause larger increases (or smaller decreases) in $\mathcal{R}_0^C$ under FDDT than DDDT when the added host species are (i) strong intraspecific interactions, (ii) strong interspecific competitors, and (iii) have higher uptake rates than resident host species in the resident host habitats.

**Proposition S2.4.** *In the absence of interspecific competition, decreases in $\mathcal{R}_0^C$ are more likely under FDDT than DDDT and increases in $\mathcal{R}_0^C$ are more likely under DDDT than FDDT.*

*Proof.* In the absence of interspecific competition, the densities of species $i \leq m$ at the $m$-host subcommunity equilibrium and the $n$-host equilibrium are equal. This implies $U_k^\ddagger - U_k^\dagger = \sum_{l>m} u_{k,S_l} N_l^\ddagger > 0$, which implies equations (S216) and (S219) are positive. As a result, (i) host additions decrease the value of $\mathcal{R}_0^C$ under DDDT only if additions also decrease the value of $\mathcal{R}_0^C$ under FDDT and (ii) if host additions increase the value of $\mathcal{R}_0^C$ under FDDT then host additions also increases the value of $\mathcal{R}_0^C$ under DDDT. Consequently, in the absence of interspecific competition, decreases in $\mathcal{R}_0^C$ are more likely under FDDT than DDDT and increases in $\mathcal{R}_0^C$ are more likely under DDDT than FDDT. □

#### S2.5 Calculations for sensitivities of infection prevalence and infected density

The calculations for the sensitivities of host 1 infection prevalence and infected density use the Jacobian of the $n$-habitat NY model (S147). The Jacobian is defined in equation (S148) of Section S2.1.3 and its properties are defined in Section S2.1.5. The only exception is that matrix $\hat{\mathbf{J}}$ in equation (S150) is used for Special Case 1.

Each section presents the general formulas for the sensitivities and uses the formulas to determine when the sensitivities for host 1 infection prevalence and host 1 infected density have the same versus different signs. The general formulas are too complex to determine the conditions under which the sensitivities are positive or negative. Instead, I determine the signs of the sensitivities in Special Cases 1 and 2 defined in Section S2.1.4.

Note that unlike the formulas for the 1-habitat model, it is not practical to list the indirect effects associated with each formula. The underlying reason is that each host species can potentially interact with the infectious propagules in every habitat. This results in a combinatorial number of indirect pathways involving subsets of the host densities ($N_i$), subsets of the infection prevalences ($Y_i$), and subsets of the infectious propagules ($P_i$). The total number of indirect pathways is on the order of $n^3$.

##### S2.5.1 Method for computing local sensitivities

The sensitivities are computed using the method described in Section S1.5.1, with the only alteration being that sign($|\mathbf{J}|$) = $(-1)^{3n}$ and equations (S62) and (S63) are replaced with

$$\begin{aligned}\frac{\partial Y_1^*}{\partial \sigma} =& \sum_{k\leq n} \frac{(-1)^{k+1+n}}{|\mathbf{J}|}\left(-\frac{\partial}{\partial\sigma}\frac{dN_k}{dt}\right)|\mathbf{J}^{[\cancel{k},\cancel{1+n}]}| + \sum_{k\leq n} \frac{(-1)^{k+n+1+n}}{|\mathbf{J}|}\left(-\frac{\partial}{\partial\sigma}\frac{dY_k}{dt}\right)|\mathbf{J}^{[\cancel{k+n},\cancel{i+n}]}| \\ &+ \sum_{k\leq n} \frac{(-1)^{2n+k+1+n}}{|\mathbf{J}|}\left(-\frac{\partial}{\partial\sigma}\frac{dP_k}{dt}\right)|\mathbf{J}^{[\cancel{2n+k},\cancel{1+n}]}|\end{aligned} \tag{S230}$$

$$
\begin{aligned}
\frac{\partial N_n^*}{\partial \sigma} =& \sum_{k\leq n} \frac{(-1)^{k+n}}{|\mathbf{J}|}\left(-\frac{\partial}{\partial \sigma}\frac{dN_k}{dt}\right)|\mathbf{J}^{[\cancel{k},\cancel{n}]}| + \sum_{k\leq n} \frac{(-1)^{k+n+n}}{|\mathbf{J}|}\left(-\frac{\partial}{\partial \sigma}\frac{dY_k}{dt}\right)|\mathbf{J}^{[\cancel{k+n},\cancel{n}]}| \\
&+ \sum_{k\leq n} \frac{(-1)^{2n+k+n}}{|\mathbf{J}|}\left(-\frac{\partial}{\partial \sigma}\frac{dP_k}{dt}\right)|\mathbf{J}^{[\cancel{2n+k},\cancel{n}]}|
\end{aligned}
. \tag{S231}
$$

The sensitivity of host 1 infected density is computed using the previous two equations,

$$
\frac{\partial I_1^*}{\partial \sigma} = N_1^* \frac{\partial Y_1^*}{\partial \sigma} + Y_1^* \frac{\partial N_1^*}{\partial \sigma} \tag{S232}
$$

and the sensitivities to the equilibrium density of host $n$ are

$$
\frac{dY_i^*}{dN_j^*} = \frac{\partial Y_i^*}{\partial \sigma} \bigg/ \frac{\partial N_j^*}{\partial \sigma} \tag{S233}
$$

$$
\frac{dI_i^*}{dN_j^*} = \frac{\partial I_i^*}{\partial \sigma} \bigg/ \frac{\partial N_j^*}{\partial \sigma}. \tag{S234}
$$

As before, when computing the sensitivities to epidemiological parameters of host $n$, $\sigma$ is determined by the specific parameter of interest (e.g., $\beta_n$ or $\mu_n$). When computing the sensitivity to the density of host $n$, I chose a parameter, $\alpha$, representing the level of intraspecific competition experienced by host $n$. In a Lokta-Volterra competition model, $\alpha$ would be the intraspecific competition coefficient $\alpha_{nn}$. In those calculations, I assume (i) increases in $\alpha$ decrease the growth rate of host $n$, (ii) $\alpha$ does not directly effect the growth rates of the other host species, and (iii) the equilibrium density for host $n$ decreases with increases in $\alpha$. The respective mathematical assumptions are (i) $\frac{\partial}{\partial \alpha}\frac{dN_n}{dt} < 0$, (ii) $\frac{\partial}{\partial \alpha}\frac{dN_k}{dt} = 0$ for $k \neq n$, and (iii) $\frac{\partial N_n^*}{\partial \alpha} < 0$. The first two assumptions are guaranteed to hold with the correct choice of parameter. The third assumption may not hold in all systems. When is does not hold, host $n$ experiences a generalized hydra effect (i.e., increased equilibrium density with a decrease in its per capita growth rate). I do not consider systems where host $n$ exhibits a generalized hydra effects because generalized hydra effects are less likely to be present in natural systems than absent. However, I note that all of my results in Section S2.5.2 results can be applied to systems where host $n$ is exhibiting a generalized hydra effect simply by changing the sign of the predicted response.

##### S2.5.2 Sensitivities of infection prevalence to host densities

**Summary of biological interpretation**: The general formulas for the sensitivities are too complex to determine general conditions under which they are positive versus negative. Special Cases 1 and 2 (defined in Section S2.1.1) show that when host densities are fixed or there is no interspecific competition (and the other conditions are satisfied), increased density of host $n$ decreases host 1 infection prevalence unless the uptake rates of host $n$ in habitat $i$ are very low. Translating these results to direct transmission pathogens yields the following. If interspecific transmission is sufficiently low and there is frequency-dependent direct transmission (FDDT), increased density of host $n$ always decreases host 1 infection prevalence. If interspecific transmission is sufficiently low and there is density-dependent direct transmission (DDDT), increased density of host $n$ always increases host 1 infection prevalence.

The signs of the sensitivities of host 1 infection prevalence and host 1 infected density often agree. The signs of the sensitivities of host 1 infection prevalence and host 1 infected density can differ when (i) host 1 is more strongly regulated by the disease than by intraspecific competition or (ii) host 1 is a sink or small source for habitat 1 and some subset of hosts $j > 1$ are large sources for habitat 1.

**General calculation for environmental transmission**: Let $\alpha$ be a parameter satisfying $\partial F_n/\partial < 0$ and $\partial F_j/\partial\alpha = 0$ for $j \neq n$. Assume $\partial N_n^*/\partial\alpha < 0$. The sensitivity of host 1 prevalence to $\alpha$ is computed as

$$\frac{\partial Y_1^*}{\partial \alpha} = \left(-\frac{\partial}{\partial\alpha}\frac{dN_n}{dt}\right)\frac{(-1)^{n+n+1}}{|\mathbf{J}|}|\mathbf{J}^{[\cancel{n},\cancel{n+1}]}| + \left(-\frac{\partial}{\partial\alpha}\frac{dY_n}{dt}\right)\frac{(-1)^{2n+n+1}}{|\mathbf{J}|}|\mathbf{J}^{[\cancel{2n},\cancel{n+1}]}| \tag{S235}$$

$$= \left(-\frac{\partial}{\partial\alpha}\frac{dN_n}{dt}\right)\frac{-1}{|\mathbf{J}|}\begin{vmatrix} \mathbf{A}^{[\cancel{n},-]} & (\mathbf{B}+\mathbf{C}\mathbf{D}_\Gamma)^{[\cancel{n},\cancel{1}]} & \mathbf{0}_{n-1\times n} \\ \mathbf{0}_{n\times n} & -\mathbf{D}_{\beta P/Y}^{[-,\cancel{1}]} & \mathbf{D}_{\beta X} \\ \mathbf{E}_N & (\mathbf{E}_Y+\mathbf{E}_Z\mathbf{D}_\Gamma)^{[-,\cancel{1}]} & -\mathbf{D}_{U+\delta} \end{vmatrix} \tag{S236}$$

$$= \left(-\frac{\partial}{\partial\alpha}\frac{dN_n}{dt}\right)\frac{-1}{|\mathbf{J}|}\begin{vmatrix} \mathbf{A}^{[\cancel{n},-]} & (\mathbf{B}+\mathbf{C}\mathbf{D}_\Gamma)^{[\cancel{n},\cancel{1}]} & \mathbf{0}_{n-1\times n} \\ \mathbf{0}_{n\times n} & \mathbf{0}_{n\times n-1} & \mathbf{D}_{\beta X} \\ \mathbf{E}_N & (\mathbf{E}_Y-\mathbf{D}_{(U+\delta)P/XY})^{[-,\cancel{1}]} & -\mathbf{D}_{U+\delta} \end{vmatrix} \tag{S237}$$

$$= \left(-\frac{\partial}{\partial\alpha}\frac{dN_n}{dt}\right)\frac{(-1)^{1+n^2}}{|\mathbf{J}|}\begin{vmatrix} \mathbf{A}^{[\cancel{n},-]} & (\mathbf{B}+\mathbf{C}\mathbf{D}_\Gamma)^{[\cancel{n},\cancel{1}]} \\ \mathbf{E}_N & (\mathbf{E}_Y-\mathbf{D}_{(U+\delta)P/XY})^{[-,\cancel{1}]} \end{vmatrix}\prod_{j\leq n}\beta_j X_j^* \tag{S238}$$

The sensitivity of host 1 total density to $\alpha$ is computed as

$$\frac{\partial N_1^*}{\partial \alpha} = \left(-\frac{\partial}{\partial\alpha}\frac{dN_n}{dt}\right)\frac{(-1)^{n+1}}{|\mathbf{J}|}|\mathbf{J}^{[\cancel{n},\cancel{1}]}| + \left(-\frac{\partial}{\partial\alpha}\frac{dY_n}{dt}\right)\frac{(-1)^{2n+1}}{|\mathbf{J}|}|\mathbf{J}^{[\cancel{2n},\cancel{1}]}| \tag{S239}$$

$$= \left(-\frac{\partial}{\partial\alpha}\frac{dN_n}{dt}\right)\frac{(-1)^{n+1}}{|\mathbf{J}|}\begin{vmatrix} \mathbf{A}^{[\cancel{n},\cancel{1}]} & (\mathbf{B}+\mathbf{C}\mathbf{D}_\Gamma)^{[\cancel{n},-]} & \mathbf{0}_{n-1\times n} \\ \mathbf{0}_{n\times n-1} & -\mathbf{D}_{\beta P/Y} & \mathbf{D}_{\beta X} \\ \mathbf{E}_N^{[-,\cancel{1}]} & \mathbf{E}_Y+\mathbf{E}_Z\mathbf{D}_\Gamma & -\mathbf{D}_{U+\delta} \end{vmatrix} \tag{S240}$$

$$= \left(-\frac{\partial}{\partial\alpha}\frac{dN_n}{dt}\right)\frac{(-1)^{n+1}}{|\mathbf{J}|}\begin{vmatrix} \mathbf{A}^{[\cancel{n},\cancel{1}]} & (\mathbf{B}+\mathbf{C}\mathbf{D}_\Gamma)^{[\cancel{n},-]} & \mathbf{0}_{n-1\times n} \\ \mathbf{0}_{n\times n-1} & \mathbf{0}_{n\times n} & \mathbf{D}_{\beta X} \\ \mathbf{E}_N^{[-,\cancel{1}]} & \mathbf{E}_Y-\mathbf{D}_{(U+\delta)P/XY} & -\mathbf{D}_{U+\delta} \end{vmatrix} \tag{S241}$$

$$= \left(-\frac{\partial}{\partial\alpha}\frac{dN_n}{dt}\right)\frac{(-1)^{1+n+n^2}}{|\mathbf{J}|}\begin{vmatrix} \mathbf{A}^{[\cancel{n},\cancel{1}]} & \mathbf{B}+\mathbf{C}\mathbf{D}_\Gamma^{[\cancel{n},-]} \\ \mathbf{E}_N^{[-,\cancel{1}]} & \mathbf{E}_Y-\mathbf{D}_{(U+\delta)P/XY} \end{vmatrix}\prod_{j\leq n}\beta_j X_j^* \tag{S242}$$

$$= \left(-\frac{\partial}{\partial\alpha}\frac{dN_n}{dt}\right)\frac{(-1)^{1+n}}{|\mathbf{J}|}\begin{vmatrix} -(\mathbf{B}+\mathbf{C}\mathbf{D}_\Gamma)_{\bullet 1}^{[\cancel{n}]} & \mathbf{A}^{[\cancel{n},\cancel{1}]} & \mathbf{B}+\mathbf{C}\mathbf{D}_\Gamma^{[\cancel{n},\cancel{1}]} \\ -(\mathbf{E}_Y-\mathbf{D}_{(U+\delta)P/XY})_{\bullet 1} & (\mathbf{E}_N^{[-,\cancel{1}]})^{[-,\cancel{1}]} & (\mathbf{E}_Y-\mathbf{D}_{(U+\delta)P/XY})^{[-,\cancel{1}]} \end{vmatrix}\prod_{j\leq n}\beta_j X_j^* \tag{S243}$$

The sensitivity of host 1 infected density ($\partial I_1^*/\partial\alpha$) is computed using the previous two equations and equation (S232). Because $(-1)^{n^2} = (-1)^n$, the only difference between equations (S238) and (S243) is that the first columns of the matrices are different. Equations (S238) and (S243) can differ in sign when the differences between the first columns are large. Those conditions are discussed in Remark S2.3.

**Special Case 1: fixed host densities, equal uptake rates, and low interspecific transmission**: Assume the host densities ($N_i$) are fixed quantities, susceptible, infected, and recovered individuals have equal uptake rates ($u_{i,S_j} = u_{i,I_j} = u_{i,R_j}$ for all $i, j$), and interspecific transmission is very low ($\chi_{ii} \gg \chi_{ij}$ for $j \neq i$). The model dynamics are given by equation (S149) and the Jacobian, $\hat{\mathbf{J}}$, is given in equation (S150). Note that sign($|\hat{\mathbf{J}}|$) $= (-1)^{2n}$ because I assume the equilibrium is stable.

Because the host densities are fixed, the sensitivities of host 1 infection prevalence and host 1 infected density are proportional to each other and always have the same sign. The sensitivity of host 1 infection prevalence to the density of host $n$ is

$$\frac{\partial Y_1^*}{\partial N_n} = \sum_{k\leq n}\left(-\frac{\partial}{\partial N_n}\frac{dP_k}{dt}\right)\frac{(-1)^{n+k+1}}{|\hat{\mathbf{J}}|}\left|\hat{\mathbf{J}}^{[\not{n}+\not{k},\not{1}]}\right| \tag{S244}$$

$$= \sum_{k\leq n}\left(-\frac{\partial}{\partial N_n}\frac{dP_k}{dt}\right)\frac{(-1)^{n+k+1}}{|\hat{\mathbf{J}}|}\begin{vmatrix} \mathbf{D}_{\beta P/Y}^{[-,\not{1}]} & \mathbf{D}_{\beta X} \\ (\mathbf{E}_Y + \mathbf{E}_Z\mathbf{D}_\Gamma)^{[\not{k},\not{1}]} & -\mathbf{D}_{U+\delta}^{[\not{k},-]} \end{vmatrix} \tag{S245}$$

$$= \sum_{k\leq n}\left(-\frac{\partial}{\partial N_n}\frac{dP_k}{dt}\right)\frac{(-1)^{n+k+1}}{|\hat{\mathbf{J}}|}\begin{vmatrix} \mathbf{0}_{n\times n-1} & \mathbf{D}_{\beta X} \\ (\mathbf{E}_Y - \mathbf{D}_{(U+\delta)P/XY})^{[\not{k},\not{1}]} & -\mathbf{D}_{U+\delta}^{[\not{k},-]} \end{vmatrix} \tag{S246}$$

$$= \sum_{k\leq n} -\varphi_{kn}\frac{(-1)^{n+k+1+n(n-1)}}{|\hat{\mathbf{J}}|}\left|(\mathbf{E}_Y - \mathbf{D}_{(U+\delta)P/XY})^{[\not{k},\not{1}]}\right|\prod_{j\leq n}\beta_j X_j^* \tag{S247}$$

Proposition S2.5 shows that the sign of each term in equation (S247) is determined by $\varphi_{kj}$. Thus, the responses in focal host infection prevalence depend on whether host $n$ is a sink or source in each habitat.

If there is DDDT (meaning $u_{i,S_j} = 0$ for all $i, j$), then all hosts are sources for all habitats and increased density of host $n$ always results in increased prevalence of host 1 ($\partial Y_1^*/\partial N_n > 0$). If there is FDDT (meaning $\delta_i = 0$ for all $i$), then host $i$ must be a source for its own habitat (in order for the equilibrium to exist) and a sink for all other habitats (because interspecific transmission is sufficiently low). In addition, $\chi_{ii}$ being sufficiently larger than $\chi_{ij}$ for $i \neq j$ means that the leading order term of equation (S247) is $\varphi_{1n}Q_1$, where $Q_1 > 0$ is defined at the end of Proposition S2.5. Consequently, increased density of host $n$ decreases host 1 infection prevalence ($\partial Y_1^*/\partial N_n < 0$) when host $n$ sheds fewer infectious propagules than it takes up in habitat 1. Increased density of host $n$ increases host 1 infection prevalence ($\partial Y_1^*/\partial N_n > 0$) when host $n$ sheds more infectious propagules than it takes up in habitat 1; this requires very low rates of uptake by host $n$.

**Special Case 2: no interspecific competition, equal uptake rates, and low interspecific transmission**: Assume no interspecific competition, all individuals have equal uptake rates ($u_{i,V_j} = u$ for all $i, j$ and $V, W \in \{S, I, R\}$), susceptible and infected individuals have equal competitive abilities ($\partial f_{V_i}/\partial S_j = \partial f_{V_i}/\partial I_j$ for all $i, j$), and interspecific transmission is very low ($\chi_{ii} \gg \chi_{ij}$ for $j \neq i$). In particular for the latter, I assume $\chi_{ij} = \mathcal{O}(\epsilon)$ for $i \neq j$ where $\epsilon \ll \chi_{ii}$ for any $i$.

Under these assumptions, $\mathbf{A}$ and $\mathbf{B}$ are diagonal matrices with negative diagonal entries because $\mathbf{A}_{ij} = 0$ and $\mathbf{B}_{ij} = 0$ for $i \neq j$, $\mathbf{E}_Y$ is a positive matrix whose off-diagonal entries are $\mathcal{O}(\epsilon)$, and $\mathbf{E}_N = \mathbf{D}_{\chi Y} - \mathbf{1}u + \mathcal{O}(\epsilon)$ where $\mathbf{D}_{\chi Y}$ is a diagonal matrix with entries $\chi_{ii}Y_i$ and $\mathbf{1}$ is an $n \times n$ matrix of ones. To leading order, the following determinant is

$$\left|(\mathbf{E}_Y - \mathbf{D}_{(U+\delta)P/XY})^{[\not{1},\not{1}]}\right| = \prod_{i=2}^{n} \left[\chi_{ii}N_i^* - \frac{(U_i^* + \delta_i)P_i^*}{X_i^* Y_i^*}\right] + \mathcal{O}(\epsilon). \tag{S248}$$

In addition, the inverse of $(\mathbf{E}_Y - \mathbf{D}_{(U+\delta)P/XY})^{[\not{1},\not{1}]}$ has diagonal entries $[\chi_{ii}N_i^* - (U_i^* + \delta)/X_i^* Y_i^*]^{-1}$ and negative off-diagonal entries that are $\mathcal{O}(\epsilon)$.

After applying Schur's formula, to leading order equation (S238) simplifies to

$$\frac{\partial Y_1^*}{\partial \alpha} = \left(-\frac{\partial}{\partial \alpha}\frac{dN_n}{dt}\right)\frac{(-1)^{1+n^2}}{|\mathbf{J}|}\begin{vmatrix} \mathbf{A}^{[\not{n},-]} & (\mathbf{B}+\mathbf{C}\mathbf{D}_\Gamma)^{[\not{n},\not{1}]} \\ \mathbf{E}_N & (\mathbf{E}_Y + \mathbf{E}_Z\mathbf{D}_\Gamma - \mathbf{D}_{(U+\delta)P/XY})^{[-,\not{1}]} \end{vmatrix} \prod_{j\le n}(\beta_j X_j^*) \tag{S249}$$

$$= \left(-\frac{\partial}{\partial \alpha}\frac{dN_n}{dt}\right)\frac{(-1)^{1+n^2}}{|\mathbf{J}|}\prod_{j\le n}(\beta_j X_j^*)\left|(\mathbf{E}_Y - \mathbf{D}_{(U+\delta)P/XY})^{[\not{1},\not{1}]}\right| \tag{S250}$$

$$\times \left|\begin{bmatrix} \mathbf{A}^{[\not{n},-]} \\ \mathbf{E}_N[1,\bullet] \end{bmatrix} - \begin{bmatrix} (\mathbf{B}+\mathbf{C}\mathbf{D}_\Gamma)^{[\not{n},\not{1}]} \\ (\mathbf{E}_Y - \mathbf{D}_{(U+\delta)P/XY})^{[-,\not{1}]}_{[1\bullet]} \end{bmatrix}\left[(\mathbf{E}_Y - \mathbf{D}_{(U+\delta)P/XY})^{[\not{1},\not{1}]}\right]^{-1}\mathbf{E}_n^{[\not{1},-]}\right| \tag{S251}$$

$$= \left(-\frac{\partial}{\partial \alpha}\frac{dN_n}{dt}\right)\frac{(-1)^{n^2}}{|\mathbf{J}|}\prod_{j\le n}\beta_j X_j^* \prod_{j=2}^{n}\left(\chi_{ii}N_i^* - \frac{(U_i^*+\delta_i)P_i^*}{X_i^* Y_i^*}\right)\mathbf{A}_{11}uP_1^* \prod_{j=2}^{n-1}\left(\frac{N_j^*(\mu_j + \Gamma_j\xi_j)\chi_{jj}Y_j^*}{\chi_{jj}N_j^* - (U_j^*+\delta_j)P_j^*/X_j^*Y_j^*} + \mathbf{A}_{jj}\right) \tag{S252}$$

$$= \left(-\frac{\partial}{\partial \alpha}\frac{dN_n}{dt}\right)\frac{(-1)^{n^2}}{|\mathbf{J}|}\prod_{j\le n}\beta_j X_j^* \prod_{j=2}^{n}\left(\chi_{ii}N_i^* - \frac{(U_i^*+\delta_i)P_i^*}{X_i^* Y_i^*}\right)\mathbf{A}_{11}uP_1^* \prod_{j=2}^{n-1}\left(\frac{-(\mu_j + \Gamma_j\xi_j)X_j^*Y_j^*}{Y_j^* + Z_j^*} + \mathbf{A}_{jj}\right) \tag{S253}$$

where the last equality used $(U_i^* + \delta_i)P_i^* = \sum_j \chi_{ij}Y_j^*N_j^* = \chi_{ii}Y_i^*N_i^* + \mathcal{O}(\epsilon)$. Equation (S253) is positive because $(-1)^{n^2} = (-1)^n$, $\text{sign}(|\mathbf{J}|) = (-1)^{3n}$, $\chi_{ii}N_i^* - (U_i^* + \delta_i)P_i^*/X_i^*Y_i^* < 0$, and $\mathbf{A}_{ii} < 0$ for all $i$. As a result, $dY_1^*/dN_n^*$ is negative. Thus, increased density of host $n$ always lower focal host infection prevalence. Intuitively, this occurs because host $n$ is a sink in all habitats but its own, so it reduces transmission to the focal host, which lowers focal host infection prevalence.

<u>Frequency-Dependent Direct Transmission</u>: FDDT corresponds to $\delta_i = 0$ and $u = 1$. Because $dY_1^*/dN_n^*$ is negative, when interspecific transmission is sufficiently low relative to intraspecific transmission, increased density of any other host species reduces host 1 infection prevalence.

<u>Density-Dependent Direct Transmission</u>: DDDT corresponds to $u_{i,V_j} = 0$. Under this constraint, the leading order term of equation (S253) is

$$
\begin{aligned}
\frac{\partial Y_1^*}{\partial \alpha} = &\left(-\frac{\partial}{\partial \alpha}\frac{dN_n}{dt}\right)\frac{(-1)^{n^2}}{|\mathbf{J}|}\prod_{j\leq n}(\beta_j X_j^*)\prod_{j=2}^{n}\left(\chi_{ii}N_i^* - \frac{(U_i^*+\delta_i)P_i^*}{X_i^* Y_i^*}\right) \\
&\times \mathbf{A}_{11}\chi_{15}\left(-1 - \frac{X_n^* Y_n^*}{N_n^*(Y_n^*+Z_n^*)}\right)\prod_{j=2}^{n-1}\left(\frac{-(\mu_j+\Gamma_j\xi_j)X_j^* Y_j^*}{Y_j^*+Z_j^*} + \mathbf{A}_{jj}\right) + \mathcal{O}(\epsilon^2)
\end{aligned}
\tag{S254}
$$

Importantly, the extra factor $-1 - \frac{X_n^* Y_n^*}{N_n^*(Y_n^*+Z_n^*)}$ results in equation (S254 being negative, which means $dY_1^*/dN_n^*$ is positive. Consequently, if there is DDDT, no interspecific competition, and low interspecific transmission, then increases in host $n$ density cause an increase in focal host infection prevalence.

**Proposition S2.5.** *Assume the host densities are fixed, assume susceptible, infected, and recovered individuals have equal uptake rates (i.e., $u_{i,S_j} = u_{i,I_j} = u_{i,R_j}$ for all $i, j$), and the shedding rate of species $i$ into habitat $i$ is sufficiently greater than the shedding rate of species $j$ into species $i$ for $j \neq i$ (i.e., $\chi_{ij}$ sufficiently small relative to $\chi_{ij}$ for $i \neq j$). Then the sensitivity of host $i$ infection prevalence to host $j$ density is given by the formula*

$$
\frac{\partial Y_i^*}{\partial N_j} = \sum_k Q_k \varphi_{kj} \tag{S255}
$$

*where the $Q_k$ are positive constants.*

*Proof.* The off-diagonal entries of $\mathbf{E}_Y - \mathbf{D}_{(U+\delta)P/XY}$ are positive because the assumption $u_{i,S_j} = u_{i,I_j}$ implies $\mathbf{E}_Y[i,j] = \chi_{ij} > 0$ for $i \neq j$. Proposition S2.2 shows that $\mathbf{E}_Y - \mathbf{D}_{(U+\delta)P/XY}$ has negative diagonal entries. The row sums of matrix $\mathbf{E}_Y - \mathbf{D}_{(U+\delta)P/XY}$ satisfy

$$
\mathbf{E}_Y[i,i] + \frac{(U_i^*+\delta_i)P^*}{X_i^* Y_i^*} + \sum_{j\neq i}|\mathbf{E}_Y[i,j]| = \chi_{ii}N_i^* - \frac{(U_i^*+\delta_i)P_i^*}{X_i^* Y_i^*} + \sum_{j\neq i}\chi_{ij}N_j^* \tag{S256}
$$

$$
= \frac{1}{Y_i}\left[\chi_{ii}Y_i^* N_i^* - (U_i^*+\delta_i)P_i^*\right] + \frac{1}{X_i^*}\left[X_i^*\sum_{j\neq i}\chi_{ij}N_j^* - (U_i^*+\delta_i)P_i^*\right] - \frac{(U_i^*+\delta_i)P_i^* Z_i^*}{Y_i^*}. \tag{S257}
$$

The terms in the small brackets are negative because $dP_i/dt = \sum_j \chi_{ij}Y_j^* N_j^* - (U_i^* - \delta_i)P_i^* = 0$ at equilibrium and the terms in the large brackets are negative for sufficiently small $\chi_{ij}$ ($i \neq j$).

Combining the above, the matrix $\mathbf{E}_Y - \mathbf{D}_{(U+\delta)P/XY}$ is diagonally dominant with negative diagonal entries and positive off-diagonal entries. The Gershgorin Disk Theorem implies that that the sign of $|\mathbf{E}_Y - \mathbf{D}_{(U+\delta)P/XY}|$ is $(-1)^n$. Proposition S2.6 shows that those properties imply $(-1)^{k+1}\left|(\mathbf{E}_Y - \mathbf{D}_{(U+\delta)P/XY})^{[k,1]}\right| / |\mathbf{E}_Y - \mathbf{D}_{(U+\delta)P/XY}| < 0$. This means the sign of $\left|(\mathbf{E}_Y - \mathbf{D}_{(U+\delta)P/XY})^{[k,1]}\right|$ is $(-1)^{k+1+1+n}$. Recalling that $\text{sign}(|\hat{\mathbf{J}}|) = (-1)^{2n}$ and $n(n-1)$ is positive for any $n$ yields that the sign of each term in equation (S247) is given by

$$
-\varphi_{kn}\frac{(-1)^{n+k+1+n(n-1)}}{(-1)^{2n}}(-1)^{k+1+1+n} = \varphi_{kn} \tag{S258}
$$

This means the equation (S247) has the form in the statement of the proposition where

$$Q_k = \frac{(-1)^{n+k}}{|\hat{\mathbf{J}}|} \left|(\mathbf{E}_Y - \mathbf{D}_{(U+\delta)P/XY})^{[\cancel{k},\cancel{1}]}\right| \prod_{j \le n} \beta_j X_j^* > 0 \tag{S259}$$

□

**Proposition S2.6.** *Let* $\mathbf{M}$ *be an* $n \times n$ *matrix whose diagonal entries are negative (*$\mathbf{M}_{ii} < 0$*), whose off-diagonal entries are positive (*$\mathbf{M}_{ij} > 0$ *for* $i \neq j$*), and whose determinant satisfies* $sign(|\mathbf{M}|) = (-1)^n$*. Assume* $\mathbf{M}$ *is diagonally dominant, which means* $\mathbf{M}_{ii} + \sum_{j \neq i} |\mathbf{M}_{ij}| < 0$ *for all* $i$*. Then all minors of* $\mathbf{M}$ *satisfy*

$$(-1)^{i+j} \left|\mathbf{M}^{[\cancel{i},\cancel{j}]}\right| / |\mathbf{M}| < 0 \quad \textit{and} \quad sign(\left|\mathbf{M}^{[\cancel{i},\cancel{j}]}\right|) = (-1)^{n+i+j+1}. \tag{S260}$$

*Proof.* We prove the statement by induction. For the base case, consider a $2 \times 2$ matrix. Without loss of generality, set $i = 2$ and compute

$$(-1)^{2+1} \left|\mathbf{M}^{[\cancel{2},\cancel{1}]}\right| / |\mathbf{M}| = (-1)\mathbf{M}[1,2]/\mathbf{M} < 0 \tag{S261}$$

$$(-1)^{2+2} \left|\mathbf{M}^{[\cancel{2},\cancel{2}]}\right| / |\mathbf{M}| = (-1)\mathbf{M}[1,1]/\mathbf{M} < 0 \tag{S262}$$

For the inductive step, I assume the proposition holds for matrices of size $n \times n$ and prove that it holds for matrices of size $n+1 \times n+1$. Let $\mathbf{M}$ be an $n+1 \times n+1$ matrix satisfying $\text{sign}(|\mathbf{M}|) = (-1)^{n+1}$. Without loss of generality I set $i = n+1$. If $j = n+1$, then the result is immediate because $\mathbf{M}^{[\cancel{n+1},\cancel{n+1}]}$ is an $n \times n$ matrix whose determinant satisfies $\text{sign}(|\mathbf{M}^{[\cancel{n+1},\cancel{n+1}]}|) = (-1)^n$ and $\text{sign}(|\mathbf{M}|) = (-1)^{n+1}$. If $j \neq n+1$, then the minor is computed by expanding the determinant along column $n+1$ to get

$$\left|\mathbf{M}^{[\cancel{n+1},\cancel{j}]}\right| = \sum_{k \neq n+1} \left|\mathbf{M}^{[\cancel{n+1}\ \cancel{k},\cancel{j}\ \cancel{n+1}]}\right| \mathbf{M}[k,n+1](-1)^{k-1+n-1} \tag{S263}$$

$$= \sum_{k \neq n+1} \overbrace{\frac{(-1)^{j+k} \left|\mathbf{M}^{[\cancel{n+1}\ \cancel{k},\cancel{j}\ \cancel{n+1}]}\right|}{|\mathbf{M}^{[\cancel{n+1},\cancel{n+1}]}|}}^{\text{negative}} \times \underbrace{\mathbf{M}[k,n+1]}_{\text{positive}}(-1)^{k-1+n-1} \times (-1)^{j+k} \left|\mathbf{M}^{[\cancel{n+1},\cancel{n+1}]}\right| \tag{S264}$$

where $\left|\mathbf{M}^{[\cancel{n+1}\ \cancel{j},\cancel{k}\ \cancel{n+1}]}\right|$ is the determinant of $\mathbf{M}$ where rows $n+1$ and $j$ and columns $k$ and $n+1$ have been removed. Each factor on the first line of equation (S264) is negative because $\left|\mathbf{M}^{[\cancel{n+1}\ \cancel{k},\cancel{j}\ \cancel{n+1}]}\right|$ is the $k,j$ minor of the $n \times n$ matrix $\mathbf{M}^{[\cancel{n+1},\cancel{n+1}]}$, which satisfies the conditions of the proposition. Each factor $\mathbf{M}[k,n+1]$ is positive because the sum is restricted to $k \neq n+1$. The sign of $\left|\mathbf{M}^{[\cancel{n+1},\cancel{n+1}]}\right|$ is $(-1)^n$. Putting this all together, the sign of each term in the summation is $(-1)^{j+1}$. This means that

$$\text{sign}\left(\frac{(-1)^{j+n+1}\left|\mathbf{M}^{[\cancel{n+1},\cancel{j}]}\right|}{|\mathbf{M}|}\right) = \frac{(-1)^{j+n+1}(-1)^{j+1}}{(-1)^{n+1}} = -1. \tag{S265}$$

□

##### S2.5.3 Sensitivities of infection prevalence to competence parameters

**Summary of biological interpretation**: The general formulas for the sensitivities are too complex to determine general conditions under which they are positive versus negative. Special Cases 1 and 2 (defined in Section S2.1.1) show that when host densities are fixed or there is no interspecific competition (and the other conditions are satisfied), changes in parameter values that increase and decrease the competence of host $n$ ($\beta_n \chi_{nn}/u_{n,S_n}[\mu_n + m_n + \nu_n]$) cause host 1 infection prevalence to increase and decrease, respectively.

The signs of the sensitivities of host 1 infection prevalence and host 1 infected density often have the same sign. The signs of the sensitivities of host 1 infection prevalence and host 1 infected density can differ when (i) host 1 is more strongly regulated by disease than by intraspecific competition or (ii) host 1 is a sink or small source for habitat 1 and some subset of hosts $j > 1$ are large sources for habitat 1.

**Shedding rates ($\chi_{in}$)**: The sensitivity of host 1 infection prevalence to the shedding rate of host $j$ into habitat $i$ is

$$\frac{\partial Y_1^*}{\partial \chi_{ij}} = \frac{(-1)^{2n+i+n+1}}{|\mathbf{J}|}\left(-\frac{\partial}{\partial \chi_{ij}}\frac{dP_i}{dt}\right)|\mathbf{J}^{[\cancel{2n+i},\cancel{n+1}]}| \tag{S266}$$

$$= \frac{(-1)^{n+i}Y_j^* N_j^*}{|\mathbf{J}|}\begin{vmatrix} \mathbf{A} & (\mathbf{B}+\mathbf{C}\mathbf{D}_\Gamma)^{[-,\cancel{1}]} & \mathbf{0}_{n\times n} \\ -\mathbf{D}_{Y/N}\mathbf{A} & -\mathbf{D}_{\beta P/Y}^{[-,\cancel{1}]} - (\mathbf{D}_{Y/N}(\mathbf{B}+\mathbf{C}\mathbf{D}_\Gamma))^{[-,\cancel{1}]} & \mathbf{D}_{\beta X} \\ \mathbf{E}_N^{[\cancel{i},-]} & (\mathbf{E}_Y+\mathbf{E}_Z\mathbf{D}_\Gamma)^{[\cancel{i},\cancel{1}]} & -\mathbf{D}_{U+\delta}^{[\cancel{i},-]} \end{vmatrix} \tag{S267}$$

$$= \frac{(-1)^{n+i}Y_j^* N_j^*}{|\mathbf{J}|}\begin{vmatrix} \mathbf{A} & (\mathbf{B}+\mathbf{C}\mathbf{D}_\Gamma)^{[-,\cancel{1}]} & \mathbf{0}_{n\times n} \\ \mathbf{0}_{n\times n} & -\mathbf{D}_{\beta P/Y}^{[-,\cancel{1}]} & \mathbf{D}_{\beta X} \\ \mathbf{E}_N^{[\cancel{i},-]} & (\mathbf{E}_Y+\mathbf{E}_Z\mathbf{D}_\Gamma)^{[\cancel{i},\cancel{1}]} & -\mathbf{D}_{U+\delta}^{[\cancel{i},-]} \end{vmatrix} \tag{S268}$$

$$= \frac{(-1)^{n+i}Y_j^* N_j^*}{|\mathbf{J}|}\begin{vmatrix} \mathbf{A} & (\mathbf{B}+\mathbf{C}\mathbf{D}_\Gamma)^{[-,\cancel{1}]} & \mathbf{0}_{n\times n} \\ \mathbf{0}_{n\times n} & \mathbf{0}_{n\times n-1} & \mathbf{D}_{\beta X} \\ \mathbf{E}_N^{[\cancel{i},-]} & (\mathbf{E}_Y-\mathbf{D}_{(U+\delta)P/XY})^{[\cancel{i},\cancel{1}]} & -\mathbf{D}_{U+\delta}^{[\cancel{i},-]} \end{vmatrix} \tag{S269}$$

$$= \frac{(-1)^{n+i}Y_j^* N_j^*}{|\mathbf{J}|}\prod_{k\leq n}(\beta_k X_k^*)\begin{vmatrix} \mathbf{A} & (\mathbf{B}+\mathbf{C}\mathbf{D}_\Gamma)^{[-,\cancel{1}]} \\ \mathbf{E}_N^{[\cancel{i},-]} & (\mathbf{E}_Y-\mathbf{D}_{(U+\delta)P/XY})^{[\cancel{i},\cancel{1}]} \end{vmatrix} \tag{S270}$$

The sensitivity of host 1 total density to the shedding rate is

$$\frac{\partial N_1^*}{\partial \chi_{ij}} = \frac{(-1)^{2n+i+1}}{|\mathbf{J}|}\left(-\frac{\partial}{\partial \chi_{ij}}\frac{dP_i}{dt}\right)|\mathbf{J}^{[\cancel{2n+i},\cancel{1}]}| \tag{S271}$$

$$= \frac{(-1)^{i}Y_j^* N_j^*}{|\mathbf{J}|}\begin{vmatrix} \mathbf{A}^{[-,\cancel{1}]} & (\mathbf{B}+\mathbf{C}\mathbf{D}_\Gamma) & \mathbf{0}_{n\times n} \\ -(\mathbf{D}_{Y/N}\mathbf{A})^{[-,\cancel{1}]} & -\mathbf{D}_{\beta P/Y} - (\mathbf{D}_{Y/N}(\mathbf{B}+\mathbf{C}\mathbf{D}_\Gamma)) & \mathbf{D}_{\beta X} \\ \mathbf{E}_N^{[\cancel{i},\cancel{1}]} & (\mathbf{E}_Y+\mathbf{E}_Z\mathbf{D}_\Gamma)^{[\cancel{i},-]} & -\mathbf{D}_{U+\delta}^{[\cancel{i},-]} \end{vmatrix} \tag{S272}$$

$$= \frac{(-1)^{i}Y_j^* N_j^*}{|\mathbf{J}|}\begin{vmatrix} \mathbf{A}^{[-,\cancel{1}]} & (\mathbf{B}+\mathbf{C}\mathbf{D}_\Gamma) & \mathbf{0}_{n\times n} \\ \mathbf{0}_{n\times n-1} & -\mathbf{D}_{\beta P/Y} & \mathbf{D}_{\beta X} \\ \mathbf{E}_N^{[\cancel{i},\cancel{1}]} & (\mathbf{E}_Y+\mathbf{E}_Z\mathbf{D}_\Gamma)^{[\cancel{i},-]} & -\mathbf{D}_{U+\delta}^{[\cancel{i},-]} \end{vmatrix} \tag{S273}$$

$$= \frac{(-1)^{i} Y_j^* N_j^*}{|\mathbf{J}|} \begin{vmatrix} \mathbf{A}^{[-,\cancel{1}]} & \mathbf{B} + \mathbf{C}\mathbf{D}_\Gamma & \mathbf{0}_{n\times n} \\ \mathbf{0}_{n\times n-1} & \mathbf{0}_{n\times n} & \mathbf{D}_{\beta X} \\ \mathbf{E}_N^{[\cancel{i},\cancel{1}]} & (\mathbf{E}_Y - \mathbf{D}_{(U+\delta)P/XY})^{[\cancel{i},-]} & -\mathbf{D}_{U+\delta}^{[\cancel{i},-]} \end{vmatrix} \tag{S274}$$

$$= \frac{(-1)^{n+i} Y_j^* N_j^*}{|\mathbf{J}|} \prod_{k\leq n} (\beta_k X_k^*) \begin{vmatrix} -(\mathbf{B} + \mathbf{C}\mathbf{D}_\Gamma)_{\bullet 1} & \mathbf{A}^{[-,\cancel{1}]} & (\mathbf{B} + \mathbf{C}\mathbf{D}_\Gamma)^{[-,\cancel{1}]} \\ -(\mathbf{E}_Y - \mathbf{D}_{(U+\delta)P/XY})_{\bullet 1}^{[\cancel{i}]} & \mathbf{E}_N^{[\cancel{i},\cancel{1}]} & (\mathbf{E}_Y - \mathbf{D}_{(U+\delta)P/XY})^{[\cancel{i},\cancel{1}]} \end{vmatrix} \tag{S275}$$

The sensitivity of host 1 infected density ($\partial I_1^*/\partial \chi_{ij}$) is computing using the two previous equations and equation (S232). The only difference between equations (S270) and (S275) is that the first columns of the matrices are different. Equations (S270) and (S275) can differ in sign when the differences between the first columns are large. Those conditions are discussed in Remark S2.3.

**Uptake rates ($u_{i,V_n}$)**: Using nearly identical calculations to the above, the sensitivities to the uptake rate of class $V_j$ in habitat $i$ are

$$\frac{\partial Y_1^*}{\partial u_{i,V_j}} = \frac{(-1)^{n+i+1} V_j^* P_i^*}{|\mathbf{J}|} \prod_{k\leq n} (\beta_k X_k^*) \begin{vmatrix} \mathbf{A} & (\mathbf{B} + \mathbf{C}\mathbf{D}_\Gamma)^{[-,\cancel{1}]} \\ \mathbf{E}_N^{[\cancel{i},-]} & (\mathbf{E}_Y + \mathbf{D}_{(U+\delta)P/XY})^{[\cancel{i},\cancel{1}]} \end{vmatrix} \tag{S276}$$

$$\frac{\partial N_1^*}{\partial u_{i,V_j}} = \frac{(-1)^{n+i+1} V_j^* P_i^*}{|\mathbf{J}|} \prod_{k\leq n} (\beta_k X_k^*) \begin{vmatrix} -(\mathbf{B} + \mathbf{C}\mathbf{D}_\Gamma)_{\bullet 1} & \mathbf{A}^{[-,\cancel{1}]} & (\mathbf{B} + \mathbf{C}\mathbf{D}_\Gamma)^{[-,\cancel{1}]} \\ -(\mathbf{E}_Y - \mathbf{D}_{(U+\delta)P/XY})_{\bullet 1}^{[\cancel{i}]} & \mathbf{E}_N^{[\cancel{i},\cancel{1}]} & (\mathbf{E}_Y - \mathbf{D}_{(U+\delta)P/XY})^{[\cancel{i},\cancel{1}]} \end{vmatrix} \tag{S277}$$

The sensitivities of host 1 infection prevalence ($\partial Y_1^*/\partial u_{i,V_n}$) and host 1 infected density ($\partial I_1^*/\partial u_{i,V_n}$; computed using equation (S232)) can have different signs when the differences between the first columns of the matrices in those equations are large. Those conditions are discussed in Remark S2.3.

**Transmission parameter ($\beta_n$)**: The sensitivity of host 1 infection prevalence to the transmission parameter of host $n$ is

$$\frac{\partial Y_1^*}{\partial \beta_n} = \frac{(-1)^{2n+n+1}}{|\mathbf{J}|} \left( \frac{\partial}{\partial \beta_n} \frac{dY_n}{dt} \right) \left| \mathbf{J}^{[\cancel{2n},\cancel{n+1}]} \right| \tag{S278}$$

$$= \frac{(-1)^{n} X_n^* P_n^*}{|\mathbf{J}|} \begin{vmatrix} \mathbf{A} & (\mathbf{B} + \mathbf{C}\mathbf{D}_\Gamma)^{[-,\cancel{1}]} & \mathbf{0}_{n\times n} \\ -(\mathbf{D}_{Y/N}\mathbf{A})^{[\cancel{n},-]} & -\mathbf{D}_{\beta P/Y}^{[\cancel{n},\cancel{1}]} - (\mathbf{D}_{Y/N}(\mathbf{B} + \mathbf{C}\mathbf{D}_\Gamma))^{[\cancel{n},\cancel{1}]} & \mathbf{D}_{\beta X}^{[\cancel{n},-]} \\ \mathbf{E}_N & (\mathbf{E}_Y + \mathbf{E}_Z\mathbf{D}_\Gamma)^{[-,\cancel{1}]} & -\mathbf{D}_{U+\delta} \end{vmatrix} \tag{S279}$$

$$= \frac{(-1)^{n} X_n^* P_n^*}{|\mathbf{J}|} \begin{vmatrix} \mathbf{A} & (\mathbf{B} + \mathbf{C}\mathbf{D}_\Gamma)^{[-,\cancel{1}]} & \mathbf{0}_{n\times n} \\ \mathbf{0}_{n-1\times n} & -\mathbf{D}_{\beta P/Y}^{[\cancel{n},\cancel{1}]} & \mathbf{D}_{\beta X}^{[\cancel{n},-]} \\ \mathbf{E}_N & (\mathbf{E}_Y + \mathbf{E}_Z\mathbf{D}_\Gamma)^{[-,\cancel{1}]} & -\mathbf{D}_{U+\delta} \end{vmatrix} \tag{S280}$$

$$= \frac{(-1)^{n} X_n^* P_n^*}{|\mathbf{J}|} \begin{vmatrix} \mathbf{A} & (\mathbf{B} + \mathbf{C}\mathbf{D}_\Gamma)^{[-,\cancel{1}]} & \mathbf{0}_{n\times n} \\ \mathbf{0}_{n-1\times n} & \mathbf{0}_{n-1\times n} & \mathbf{D}_{\beta X}^{[\cancel{n},-]} \\ \mathbf{E}_N & (\mathbf{E}_Y - \mathbf{D}_{(U+\delta)P/XY})^{[-,\cancel{1}]} & -\mathbf{D}_{U+\delta} \end{vmatrix} \tag{S281}$$

$$
= \frac{X_n^* P_n^*}{|\mathbf{J}|}(U_n^* + \delta_n)\prod_{k<n}(\beta_k X_k^*) \begin{vmatrix} \mathbf{A} & (\mathbf{B}+\mathbf{C}\mathbf{D}_\Gamma)^{[-,\not{1}]} \\ \mathbf{E}_N^{[\not{n},-]} & (\mathbf{E}_Y - \mathbf{D}_{(U+\delta)P/XY})^{[\not{n},\not{1}]} \end{vmatrix} \tag{S282}
$$

Following similar steps, the sensitivity of host 1 total density to the shedding rate is

$$
\frac{\partial N_1^*}{\partial \beta_n} = \frac{(-1)^{2n+1}}{|\mathbf{J}|}\left(\frac{\partial}{\partial \beta_n}\frac{dY_n}{dt}\right)|\mathbf{J}^{[\not{2n},\not{1}]}| \tag{S283}
$$

$$
= \frac{X_n^* P_n^*}{|\mathbf{J}|}(U_n^* + \delta_n)\prod_{k<n}(\beta_k X_k^*) \begin{vmatrix} -(\mathbf{B}+\mathbf{C}\mathbf{D}_\Gamma)_{\bullet 1} & \mathbf{A}^{[-,\not{1}]} & (\mathbf{B}+\mathbf{C}\mathbf{D}_\Gamma)^{[-,\not{1}]} \\ -(\mathbf{E}_Y - \mathbf{D}_{(U+\delta)P/XY})_{\bullet 1}^{[\not{n}]} & \mathbf{E}_N^{[\not{n},\not{1}]} & (\mathbf{E}_Y - \mathbf{D}_{(U+\delta)P/XY})^{[\not{n},\not{1}]} \end{vmatrix} \tag{S284}
$$

The sensitivity of host 1 infected density ($\partial I_1^*/\partial \beta_n$) is computed using the two previous equations and equation (S232). Because the only difference between equations (S282) and (S284) is that the first columns of the matrices are different, equations (S282) and (S284) can differ in sign when the differences between the first columns are large. Those conditions are discussed in Remark S2.3.

**Disease-induced mortality rate ($\mu_n$)**: The sensitivity of host 1 infection prevalence to the disease-induced mortality rate of host $n$ is

$$
\frac{\partial Y_1^*}{\partial \mu_n} = \left(-\frac{\partial}{\partial \mu_n}\frac{dN_n}{dt}\right)\frac{(-1)^{n+n+1}}{|\mathbf{J}|}|\mathbf{J}^{[\not{n},\not{n+1}]}| + \left(-\frac{\partial}{\partial \mu_n}\frac{dY_n}{dt}\right)\frac{(-1)^{2n+n+1}}{|\mathbf{J}|}|\mathbf{J}^{[\not{2n},\not{n+1}]}| \tag{S285}
$$

$$
= \frac{-Y_n^* N_n^*}{|\mathbf{J}|}\begin{vmatrix} \mathbf{A}^{[\not{n},-]} & (\mathbf{B}+\mathbf{C}\mathbf{D}_\Gamma)^{[\not{n},\not{1}]} & \mathbf{0}_{n-1\times n} \\ \mathbf{0}_{n\times n} & -\mathbf{D}_{\beta P/Y}^{[-,\not{1}]} & \mathbf{D}_{\beta X} \\ \mathbf{E}_N & (\mathbf{E}_Y + \mathbf{E}_Z\mathbf{D}_\Gamma)^{[-,\not{1}]} & -\mathbf{D}_{U+\delta} \end{vmatrix} \tag{S286}
$$

$$
+ \frac{(-1)^{n+1}Y_n^*}{|\mathbf{J}|}\begin{vmatrix} \mathbf{A} & (\mathbf{B}+\mathbf{C}\mathbf{D}_\Gamma)^{[-,\not{1}]} & \mathbf{0}_{n\times n} \\ -(\mathbf{D}_{Y/N}\mathbf{A})^{[\not{n},-]} & -\mathbf{D}_{\beta P/Y}^{[\not{n},\not{1}]} - (\mathbf{D}_{Y/N}(\mathbf{B}+\mathbf{C}\mathbf{D}_\Gamma))^{[\not{n},\not{1}]} & \mathbf{D}_{\beta X}^{[\not{n},-]} \\ \mathbf{E}_N & (\mathbf{E}_Y + \mathbf{E}_Z\mathbf{D}_\Gamma)^{[-,\not{1}]} & -\mathbf{D}_{U+\delta} \end{vmatrix} \tag{S287}
$$

$$
= \frac{-Y_n^* N_n^*}{|\mathbf{J}|}\begin{vmatrix} \mathbf{A}^{[\not{n},-]} & (\mathbf{B}+\mathbf{C}\mathbf{D}_\Gamma)^{[\not{n},\not{1}]} & \mathbf{0}_{n-1\times n} \\ \mathbf{0}_{n\times n} & \mathbf{0}_{n\times n-1} & \mathbf{D}_{\beta X} \\ \mathbf{E}_N & (\mathbf{E}_Y - \mathbf{D}_{(U+\delta)P/XY})^{[-,\not{1}]} & -\mathbf{D}_{U+\delta} \end{vmatrix} \tag{S288}
$$

$$
+ \frac{(-1)^{n+1}Y_n^*}{|\mathbf{J}|}\begin{vmatrix} \mathbf{A} & (\mathbf{B}+\mathbf{C}\mathbf{D}_\Gamma)^{[-,\not{1}]} & \mathbf{0}_{n\times n} \\ \mathbf{0}_{n-1\times n} & \mathbf{0}_{n-1\times n} & \mathbf{D}_{\beta X}^{[\not{n},-]} \\ \mathbf{E}_N & (\mathbf{E}_Y - \mathbf{D}_{(U+\delta)P/XY})^{[-,\not{1}]} & -\mathbf{D}_{U+\delta} \end{vmatrix} \tag{S289}
$$

$$
\begin{aligned}
&= \frac{(-1)^{1+n^2}Y_n^* N_n^*}{|\mathbf{J}|}\begin{vmatrix} \mathbf{A}^{[\not{n},-]} & (\mathbf{B}+\mathbf{C}\mathbf{D}_\Gamma)^{[\not{n},\not{1}]} \\ \mathbf{E}_N & (\mathbf{E}_Y - \mathbf{D}_{(U+\delta)P/XY})^{[-,\not{1}]} \end{vmatrix}\prod_{j\le n}(\beta_j X_j^*) \\
&\quad + \frac{-Y_n^*}{|\mathbf{J}|}(U_n + \delta_n)\prod_{k<n}(\beta_k X_k^*)\begin{vmatrix} \mathbf{A} & (\mathbf{B}+\mathbf{C}\mathbf{D}_\Gamma)^{[-,\not{1}]} \\ \mathbf{E}_N^{[\not{n},-]} & (\mathbf{E}_Y - \mathbf{D}_{(U+\delta)P/XY})^{[\not{n},\not{1}]} \end{vmatrix}
\end{aligned} \tag{S290}
$$

where algebraic steps are the same as the calculations for $dY_1^*/dN_n^*$ and $\partial Y_1^*/\partial \beta_n$. Similar calculations also yield the sensitivity of host 1 total density,

$$\frac{\partial N_1^*}{\partial \mu_n} = \left(-\frac{\partial}{\partial \mu_n}\frac{dN_n}{dt}\right)\frac{(-1)^{n+1}}{|\mathbf{J}|}|\mathbf{J}^{[\cancel{n},\cancel{1}]}| + \left(-\frac{\partial}{\partial \mu_n}\frac{dY_n}{dt}\right)\frac{(-1)^{2n+1}}{|\mathbf{J}|}|\mathbf{J}^{[\cancel{2n},\cancel{1}]}| \tag{S291}$$

$$= \frac{(-1)^{1+n^2}Y_n^*N_n^*}{|\mathbf{J}|}\begin{vmatrix} -(\mathbf{B}+\mathbf{C}\mathbf{D}_\Gamma)^{[\cancel{n},\cancel{1}]} & \mathbf{A}_{\bullet 1}^{[\cancel{n}]} & (\mathbf{B}+\mathbf{C}\mathbf{D}_\Gamma)^{[\cancel{n},\cancel{1}]} \\ -(\mathbf{E}_Y+\mathbf{E}_Z\mathbf{D}_\Gamma-\mathbf{D}_{(U+\delta)P/XY})_{\bullet 1} & \mathbf{E}_N^{[-,\cancel{1}]} & (\mathbf{E}_Y-\mathbf{D}_{(U+\delta)P/XY})^{[-,\cancel{1}]} \end{vmatrix}\prod_{j\leq n}\beta_j X_j^*$$
$$+ \frac{-Y_n^*}{|\mathbf{J}|}(U_n^*+\delta_n)\prod_{k<n}(\beta_k X_k^*)\begin{vmatrix} -(\mathbf{B}+\mathbf{C}\mathbf{D}_\Gamma)_{\bullet 1} & \mathbf{A}^{[-,\cancel{1}]} & (\mathbf{B}+\mathbf{C}\mathbf{D}_\Gamma)^{[-,\cancel{1}]} \\ -(\mathbf{E}_Y-\mathbf{D}_{(U+\delta)P/XY})_{\bullet 1}^{[\cancel{n}]} & \mathbf{E}_N^{[\cancel{n},\cancel{1}]} & (\mathbf{E}_Y-\mathbf{D}_{(U+\delta)P/XY})^{[\cancel{n},\cancel{1}]} \end{vmatrix} \tag{S292}$$

The sensitivity of host 1 infected density ($\partial I_1^*/\partial \mu_n$) is computing using the two previous equations and equation (S232). Because the only differences between equations (S290) and (S292) are that the first columns of the matrices are different, equations (S290) and (S292) can differ in sign when the differences between the first columns are large. Those conditions are discussed in Remark S2.3.

**Recovery rate ($\nu_n$)**: Using the same algebraic steps as in the calculations for $\partial Y_1^*/\partial \mu_n$ and $\partial N_1^*/\partial \mu_n$, the sensitivities of host 1 infection prevalence and host 1 total density to the recovery rate of host $n$ are

$$\frac{\partial Y_1^*}{\partial \nu_n} = \left(-\frac{\partial}{\partial \nu_n}\frac{dN_n}{dt}\right)\frac{(-1)^{n+n+1}}{|\mathbf{J}|}|\mathbf{J}^{[\cancel{n},\cancel{n+1}]}| + \left(-\frac{\partial}{\partial \nu_n}\frac{dY_n}{dt}\right)\frac{(-1)^{2n+n+1}}{|\mathbf{J}|}|\mathbf{J}^{[\cancel{2n},\cancel{n+1}]}| \tag{S293}$$

$$= \frac{(-1)^{1+n^2}Y_n^*N_n^*\xi_n}{(m_n+\xi_n+\gamma_n)|\mathbf{J}|}\prod_{j\leq n}(\beta_j X_j^*)\begin{vmatrix} \mathbf{A}^{[\cancel{n},-]} & (\mathbf{B}+\mathbf{C}\mathbf{D}_\Gamma)^{[\cancel{n},\cancel{1}]} \\ \mathbf{E}_N & (\mathbf{E}_Y-\mathbf{D}_{(U+\delta)P/XY})^{[-,\cancel{1}]} \end{vmatrix}$$
$$+ \frac{-Y_n^*}{|\mathbf{J}|}(U_n^*+\delta_n)\prod_{k<n}(\beta_k X_k^*)\begin{vmatrix} \mathbf{A} & (\mathbf{B}+\mathbf{C}\mathbf{D}_\Gamma)^{[-,\cancel{1}]} \\ \mathbf{E}_N^{[\cancel{n},-]} & (\mathbf{E}_Y-\mathbf{D}_{(U+\delta)P/XY})^{[\cancel{n},\cancel{1}]} \end{vmatrix} \tag{S294}$$

$$\frac{\partial N_1^*}{\partial \nu_n} = \left(-\frac{\partial}{\partial \nu_n}\frac{dN_n}{dt}\right)\frac{(-1)^{n+1}}{|\mathbf{J}|}|\mathbf{J}^{[\cancel{n},\cancel{1}]}| + \left(-\frac{\partial}{\partial \nu_n}\frac{dY_n}{dt}\right)\frac{(-1)^{2n+1}}{|\mathbf{J}|}|\mathbf{J}^{[\cancel{2n},\cancel{1}]}| \tag{S295}$$

$$= \frac{(-1)^{1+n^2}Y_n^*N_n^*\xi_n}{(m_n+\xi_n+\gamma_n)|\mathbf{J}|}\prod_{j\leq n}(\beta_j X_j^*)\begin{vmatrix} -(\mathbf{B}+\mathbf{C}\mathbf{D}_\Gamma)_{\bullet 1}^{[\cancel{n}]} & \mathbf{A}^{[\cancel{n},-]} & (\mathbf{B}+\mathbf{C}\mathbf{D}_\Gamma)^{[\cancel{n},\cancel{1}]} \\ -(\mathbf{E}_Y-\mathbf{D}_{(U+\delta)P/XY})_{\bullet 1} & \mathbf{E}_N^{[-,\cancel{1}]} & (\mathbf{E}_Y-\mathbf{D}_{(U+\delta)P/XY})^{[-,\cancel{1}]} \end{vmatrix}$$
$$+ \frac{-Y_n^*}{|\mathbf{J}|}(U_n^*+\delta_n)\prod_{k<n}(\beta_k X_k^*)\begin{vmatrix} -(\mathbf{B}+\mathbf{C}\mathbf{D}_\Gamma)_{\bullet 1} & \mathbf{A}^{[-,\cancel{1}]} & (\mathbf{B}+\mathbf{C}\mathbf{D}_\Gamma)^{[-,\cancel{1}]} \\ -(\mathbf{E}_Y-\mathbf{D}_{(U+\delta)P/XY})_{\bullet 1}^{[\cancel{n}]} & \mathbf{E}_N^{[\cancel{n},\cancel{1}]} & (\mathbf{E}_Y-\mathbf{D}_{(U+\delta)P/XY})^{[\cancel{n},\cancel{1}]} \end{vmatrix} \tag{S296}$$

The sensitivity of host 1 infected density ($\partial I_1^*/\partial \nu_n$) is computing using the two previous equations and equation (S232). Because the only differences between equations (S294) and (S296) are that the first columns of the matrices are different, equations (S294) and (S296) can differ in sign when the differences between the first columns are large. Those conditions are discussed in Remark S2.3.

**Special Case 1: fixed host densities, equal uptake rates, and low interspecific transmission**: Assume the host densities ($N_i$) are fixed quantities, susceptible, infected, and recovered individuals have equal uptake rates ($u_{i,S_j} = u_{i,I_j} = u_{i,R_j}$ for all $i, j$), and interspecific transmission is very low ($\chi_{ii} \gg \chi_{ij}$ for $j \neq i$). The model dynamics are given by equation (S149) and the Jacobian, $\hat{\mathbf{J}}$, is given in equation (S150). Note that sign($|\mathbf{J}|$) = $(-1)^{2n}$ because I assume the equilibrium is stable.

Proposition S2.7 shows changes in parameter values that increase and decrease the competence of host $n$ ($\beta_n \chi_{nn}/u_{n,S_n}[\mu_n + \nu_n + m_n]$) cause increases and decreases, respectively, in host 1 infection prevalence. Because the host densities are fixed, the sensitivities of host 1 infection prevalence and host 1 infected density are proportional to each other and always have the same sign. This prediction automatically applies to models with frequency-dependent direct transmission and density-dependent direct transmission.

**Special Case 2: no interspecific competition, equal uptake rates, and low interspecific transmission**: Assume no interspecific competition, all individuals have equal uptake rates ($u_{i,V_j} = u$ for all $i, j$ and $V, W \in \{S, I, R\}$), susceptible and infected individuals have equal competitive abilities ($\partial f_{V_i}/\partial S_j = \partial f_{V_i}/\partial I_j$ for all $i, j$), and interspecific transmission is very low ($\chi_{ii} \gg \chi_{ij}$ for $j \neq i$). In particular for the latter, I assume $\chi_{ij} = \mathcal{O}(\epsilon)$ for $i \neq j$ where $\epsilon \ll \chi_{ii}$ for any $i$ and $\epsilon \ll u$.

Under these assumptions, $\mathbf{A}$ and $\mathbf{B}$ are diagonal matrices with negative diagonal entries because $\mathbf{A}_{ij} = 0$ and $\mathbf{B}_{ij} = 0$ for $i \neq j$, $\mathbf{E}_Y$ is a positive matrix whose off-diagonal entries are $\mathcal{O}(\epsilon)$, and $\mathbf{E}_N = \mathbf{D}_{\chi Y} - \mathbf{1}u + \mathcal{O}(\epsilon)$ where $\mathbf{D}_{\chi Y}$ is a diagonal matrix with entries $\chi_{ii} Y_i$ and $\mathbf{1}$ is an $n \times n$ matrix of ones. To leading order, the sensitivities of host 1 infection prevalence are

$$\begin{aligned}\frac{\partial Y_1^*}{\partial \chi_{ij}} =& \frac{(-1)^{n+1} Y_j^* N_j^*}{|\mathbf{J}|} \prod_{k \leq n} (\beta_k X_k^*)\, \mathbf{A}_{11} u P_1^* N_i^* (\mu_i + \Gamma_i \xi_i) \\ &\times \prod_{k \neq 1,i} \left( \chi_{kk} Y_k^* N_k^* (\mu_2 + \Gamma_2 \xi_2) + \mathbf{A}_{kk} \left[ \chi_{kk} N_k^* - \frac{(U_k^* + \delta_k) P_k^*}{X_k^* Y_k^*} \right] \right)\end{aligned} \tag{S297}$$

$$\begin{aligned}\frac{\partial Y_1^*}{\partial u_{i,V_j}} =& \frac{(-1)^{n} V_j^* P_i^*}{|\mathbf{J}|} \prod_{k \leq n} (\beta_k X_k^*)\, \mathbf{A}_{11} u P_1^* N_i^* (\mu_i + \Gamma_i \xi_i) \\ &\times \prod_{k \neq 1,i} \left( \chi_{kk} Y_k^* N_k^* (\mu_2 + \Gamma_2 \xi_2) + \mathbf{A}_{kk} \left[ \chi_{kk} N_k^* - \frac{(U_k^* + \delta_k) P_k^*}{X_k^* Y_k^*} \right] \right)\end{aligned} \tag{S298}$$

$$\begin{aligned}\frac{\partial Y_1^*}{\partial \beta_n} =& \frac{(-1)^{n+1} X_n^* P_n^*}{|\mathbf{J}|} (U_n^* + \delta_n) \prod_{k < n} (\beta_k X_k^*)\, \mathbf{A}_{11} u P_1^* N_n^* (\mu_n + \Gamma_n \xi_n) \\ &\times \prod_{k \neq 1,n} \left( \chi_{kk} Y_k^* N_k^* (\mu_2 + \Gamma_2 \xi_2) + \mathbf{A}_{kk} \left[ \chi_{kk} N_k^* - \frac{(U_k^* + \delta_k) P_k^*}{X_k^* Y_k^*} \right] \right)\end{aligned} \tag{S299}$$

$$
\begin{aligned}
\frac{\partial Y_1^*}{\partial \mu_n} =& \frac{(-1)^{n^2+1} Y_n^* N_n^*}{|\mathbf{J}|} \prod_{j\leq n} \left(\beta_j X_j^*\right) \prod_{j=2}^{n} \left( \chi_{ii} N_i^* - \frac{(U_i^* + \delta_i) P_i^*}{X_i^* Y_i^*} \right) \\
& \times \mathbf{A}_{11} u P_1^* \prod_{j=2}^{n-1} \left( \frac{-(\mu_j + \Gamma_j \xi_j) X_j^* Y_j^*}{Y_j^* + Z_j^*} + \mathbf{A}_{jj} \right) \\
& + \frac{(-1)^n Y_n^*}{|\mathbf{J}|} (U_n^* + \delta_n) \prod_{k<n} \left(\beta_k X_k^*\right) \mathbf{A}_{11} u P_1^* N_n^* (\mu_n + \Gamma_n \xi_n) \\
& \times \prod_{k\neq 1,n} \left( \chi_{kk} Y_k^* N_k^* (\mu_2 + \Gamma_2 \xi_2) + \mathbf{A}_{kk} \left[ \chi_{kk} N_k^* - \frac{(U_k^* + \delta_k) P_k^*}{X_k^* Y_k^*} \right] \right)
\end{aligned}
\tag{S300}
$$

$$
\begin{aligned}
\frac{\partial Y_1^*}{\partial \nu_n} =& \frac{(-1)^{n^2+1} Y_n^* N_n^* \xi_n}{(m_n + \xi_n + \gamma_n)|\mathbf{J}|} \prod_{j\leq n} \left(\beta_j X_j^*\right) \prod_{j=2}^{n} \left( \chi_{ii} N_i^* - \frac{(U_i^* + \delta_i) P_i^*}{X_i^* Y_i^*} \right) \\
& \times \mathbf{A}_{11} u P_1^* \prod_{j=2}^{n-1} \left( \frac{-(\mu_j + \Gamma_j \xi_j) X_j^* Y_j^*}{Y_j^* + Z_j^*} + \mathbf{A}_{jj} \right) \\
& + \frac{(-1)^n Y_n^*}{|\mathbf{J}|} (U_n^* + \delta_n) \prod_{k<n} \left(\beta_k X_k^*\right) \mathbf{A}_{11} u P_1^* N_n^* (\mu_n + \Gamma_n \xi_n) \\
& \times \prod_{k\neq 1,n} \left( \chi_{kk} Y_k^* N_k^* (\mu_2 + \Gamma_2 \xi_2) + \mathbf{A}_{kk} \left[ \chi_{kk} N_k^* - \frac{(U_k^* + \delta_k) P_k^*}{X_k^* Y_k^*} \right] \right)
\end{aligned}
\tag{S301}
$$

(S302)

Recall that $\mathrm{sign}(|\mathbf{J}|) = (-1)^{3n}$, $\chi_{kk} N_k^* - (U_k^* + \delta_k) P_k^* / X_k^* Y_k^* < 0$, and $\mathbf{A}_{kk} < 0$. Applying that to the above equations yields $\partial Y_1^* / \partial \chi_{ij} > 0$, $\partial Y_1^* / \partial u_{i,V_j} < 0$, $\partial Y_1^* / \partial \beta_i > 0$, $\partial Y_1^* / \partial \mu_n < 0$, and $\partial Y_1^* / \partial \mu_n < 0$. Thus, a change in a parameter that increases or decreases the competence of host $n$ causes an increase or decrease, respectively, in host 1 infection prevalence.

If the uptake rates are zero, then to leading order the sensitivities of host 1 infection prevalence are

$$
\begin{aligned}
\frac{\partial Y_1^*}{\partial \chi_{ij}} =& \frac{(-1)^n Y_j^* N_j^*}{|\mathbf{J}|} \prod_{k\leq n} \left(\beta_k X_k^*\right) \mathbf{A}_{11} \mathbf{A}_{ii} \chi_{ii} N_i^* \\
& \times \prod_{k\neq 1,i} \left( \chi_{kk} Y_k^* N_k^* (\mu_2 + \Gamma_2 \xi_2) + \mathbf{A}_{kk} \left[ \chi_{kk} N_k^* - \frac{(U_k^* + \delta_k) P_k^*}{X_k^* Y_k^*} \right] \right)
\end{aligned}
\tag{S303}
$$

$$
\begin{aligned}
\frac{\partial Y_1^*}{\partial u_{i,V_j}} =& \frac{(-1)^{n+1} V_j^* P_i^*}{|\mathbf{J}|} \prod_{k\leq n} \left(\beta_k X_k^*\right) \mathbf{A}_{11} \mathbf{A}_{ii} \chi_{ii} N_i^* \\
& \times \prod_{k\neq 1,i} \left( \chi_{kk} Y_k^* N_k^* (\mu_2 + \Gamma_2 \xi_2) + \mathbf{A}_{kk} \left[ \chi_{kk} N_k^* - \frac{(U_k^* + \delta_k) P_k^*}{X_k^* Y_k^*} \right] \right)
\end{aligned}
\tag{S304}
$$

$$
\begin{aligned}
\frac{\partial Y_1^*}{\partial \beta_n} =& \frac{(-1)^n X_n^* P_n^*}{|\mathbf{J}|} (U_n^* + \delta_n) \prod_{k<n} \left(\beta_k X_k^*\right) \mathbf{A}_{11} \mathbf{A}_{nn1} \chi_{1n} N_n^* \\
& \times \prod_{k\neq 1,n} \left( \chi_{kk} Y_k^* N_k^* (\mu_2 + \Gamma_2 \xi_2) + \mathbf{A}_{kk} \left[ \chi_{kk} N_k^* - \frac{(U_k^* + \delta_k) P_k^*}{X_k^* Y_k^*} \right] \right)
\end{aligned}
\tag{S305}
$$

$$
\begin{aligned}
\frac{\partial Y_1^*}{\partial \mu_n} =& \frac{(-1)^{n^2+1} Y_n^* N_n^*}{|\mathbf{J}|} \prod_{j \leq n} \left(\beta_j X_j^*\right) \prod_{j=2}^{n} \left( \chi_{ii} N_i^* - \frac{(U_i^* + \delta_i) P_i^*}{X_i^* Y_i^*} \right) \\
& \times \mathbf{A}_{11} \chi_{nn} \left( 1 + \frac{Y_n^* N_n^*}{N_n^*(Y) n^* + Z_n^*} \right) \prod_{j=2}^{n-1} \left( \frac{-(\mu_j + \Gamma_j \xi_j) X_j^* Y_j^*}{Y_j^* + Z_j^*} + \mathbf{A}_{jj} \right) \\
& + \frac{(-1)^{n+1} Y_n^*}{|\mathbf{J}|} (U_n^* + \delta_n) \prod_{k<n} \left(\beta_k X_k^*\right) \mathbf{A}_{11} \mathbf{A}_{nn} \chi_{1n} N_n^* \\
& \times \prod_{k \neq 1, n} \left( \chi_{kk} Y_k^* N_k^* (\mu_2 + \Gamma_2 \xi_2) + \mathbf{A}_{kk} \left[ \chi_{kk} N_k^* - \frac{(U_k^* + \delta_k) P_k^*}{X_k^* Y_k^*} \right] \right)
\end{aligned} \tag{S306}
$$

$$
\begin{aligned}
\frac{\partial Y_1^*}{\partial \nu_n} =& \frac{(-1)^{n^2+1} Y_n^* N_n^* \xi_n}{(m_n + \xi_n + \gamma_n)|\mathbf{J}|} \prod_{j \leq n} \left(\beta_j X_j^*\right) \prod_{j=2}^{n} \left( \chi_{ii} N_i^* - \frac{(U_i^* + \delta_i) P_i^*}{X_i^* Y_i^*} \right) \\
& \times \mathbf{A}_{11} \chi_{1n} \left( 1 + \frac{Y_n^* N_n^*}{N_n^*(Y) n^* + Z_n^*} \right) \prod_{j=2}^{n-1} \left( \frac{-(\mu_j + \Gamma_j \xi_j) X_j^* Y_j^*}{Y_j^* + Z_j^*} + \mathbf{A}_{jj} \right) \\
& + \frac{(-1)^{n+1} Y_n^*}{|\mathbf{J}|} (U_n^* + \delta_n) \prod_{k<n} \left(\beta_k X_k^*\right) \mathbf{A}_{11} \mathbf{A}_{nn} \chi_{1n} N_n^* \\
& \times \prod_{k \neq 1, n} \left( \chi_{kk} Y_k^* N_k^* (\mu_2 + \Gamma_2 \xi_2) + \mathbf{A}_{kk} \left[ \chi_{kk} N_k^* - \frac{(U_k^* + \delta_k) P_k^*}{X_k^* Y_k^*} \right] \right)
\end{aligned} \tag{S307}
$$

$$\tag{S308}$$

Note that the leading order terms of the sensitivities are $\mathcal{O}(\epsilon)$. The signs of the sensitivities are the same as the case when $\epsilon \ll u$.

**Proposition S2.7.** *Assume the host densities are fixed, susceptible, infected, and recovered individuals have equal uptake rates ($u_{i,S_j} = u_{i,I_j} = u_{i,R_j}$ for all $i, j$), and host $j$ has much smaller shedding rates than host $i$ into habitat $i$ (i.e., $\chi_{ij}$ for $i \neq j$ sufficiently small relative to $\chi_{ii}$). The sensitivities of host $i$ infection prevalence to the parameters of host $n$ satisfy*

$$
\frac{\partial Y_1^*}{\partial \chi_{in}} > 0, \quad \frac{\partial Y_1^*}{\partial u_{i,V_n}} < 0, \quad \frac{\partial Y_1^*}{\partial \beta_n} > 0, \quad \frac{\partial Y_1^*}{\partial \mu_n} < 0, \quad \frac{\partial Y_1^*}{\partial \nu_n} < 0
$$

*where $V \in \{S, I, R\}$.*

*Proof.* The Jacobian, $\hat{\mathbf{J}}$, for the $n$-habitat model with fixed host densities is given in equation (S150). The sensitivities of host 1 infection prevalence to the epidemiological parameters for host $n$ are

$$
\frac{\partial Y_1}{\partial \chi_{in}} = \frac{(-1)^{n+i+1} Y_n^* N_n^*}{|\hat{\mathbf{J}}|} \prod_{k \leq n} \beta_n X_n^* \left| (\mathbf{E}_Y - \mathbf{D}_{(U+\delta)P/XY})^{[\not{i}, \not{1}]} \right| \tag{S309}
$$

$$
\frac{\partial Y_1}{\partial u_{i,V_n}} = \frac{(-1)^{n+i+1} V_n^* P_i^*}{|\hat{\mathbf{J}}|} \prod_{k \leq n} \beta_n X_n^* \left| (\mathbf{E}_Y - \mathbf{D}_{(U+\delta)P/XY})^{[\not{i}, \not{1}]} \right| \tag{S310}
$$

$$\frac{\partial Y_1}{\partial \beta_n} = \frac{(-1)^{n+1+(n-1)^2} X_n^* P_n^*}{|\hat{\mathbf{J}}|}(U_n^* + \delta_n)\prod_{k<n} \beta_n X_n^* \left|(\mathbf{E}_Y - \mathbf{D}_{(U+\delta)P/XY})^{[\not{n},\not{1}]}\right| \tag{S311}$$

$$\frac{\partial Y_1}{\partial \mu_n} = \frac{(-1)^{n+(n-1)^2} Y_n^*}{|\hat{\mathbf{J}}|}(U_n^* + \delta_n)\prod_{k<n} \beta_n X_n^* \left|(\mathbf{E}_Y - \mathbf{D}_{(U+\delta)P/XY})^{[\not{n},\not{1}]}\right| \tag{S312}$$

$$\frac{\partial Y_1}{\partial \nu_n} = \frac{(-1)^{n+(n-1)^2} Y_n^*}{|\hat{\mathbf{J}}|}(U_n^* + \delta_n)\prod_{k<n} \beta_n X_n^* \left|(\mathbf{E}_Y - \mathbf{D}_{(U+\delta)P/XY})^{[\not{n},\not{1}]}\right| \tag{S313}$$

The proof of Proposition S2.5 shows that the matrix $\mathbf{E}_Y - \mathbf{D}_{(U+\delta)P/XY}$ has negative diagonal entries and positive off-diagonal entries and is diagonally dominant. The Gershgorin Disc Theorem implies $\text{sign}(|\mathbf{E}_Y - \mathbf{D}_{(U+\delta)P/XY}|) = (-1)^{-n}$. Thus, via Proposition S2.6, the signs of the minors of $\mathbf{E}_Y - \mathbf{D}_{(U+\delta)P/XY}$ are

$$\text{sign}([\mathbf{E}_Y - \mathbf{D}_{(U+\delta)P/XY}]^{[\not{i},\not{j}]}) = (-1)^{i+j+n+1}. \tag{S314}$$

Using the above to compute the signs of the partial derivatives yields the result. □

##### S2.5.4 Sensitivities of infection prevalence to transmission mode

Here, I use a continuous change of parameters to convert the $n$-habitat ET model (S147) from a frequency-dependent direct transmission (FDDT) form ($\delta_i = 0$) to a density-dependent direct transmission (DDDT) form ($u_{i,V_j} = 0$ for $V \in \{S, I, R\}$). The change of parameters holds all parameters constant except $\delta_i$ and $u_{i,V_j}$, which must necessarily change. The change of parameters holds constant the equilibrium host densities and frequencies for a particular assemblage of species. The biological interpretation of the constraint is that the densities and infection prevalences for a particular assemblage of host species are known, the pathogen's transmission mode is unknown, and the host densities and infection prevalences are unknown when a set of host species are added to or lost from the community. This method allows one to predict how changes in focal host infection prevalence and focal host infected density after the removal or addition of one or more host species depend on the pathogen transmission mode. Specifically, it identifies if greater increases (or smaller decreases) in focal host infection prevalence and infected density occur under DDDT or FDDT.

Note that in order to apply the following theory, one must know/assume the ratio of the uptake rates for all host species and classes, i.e., know the ratios of $u_{i,S_j}$, $u_{i,I_j}$, $u_{i,S_k}$ and $u_{i,I_k}$ for all $i, j, k$. That information is needed to define the change of parameters functions. While many FDDT studies use models corresponding to $u_{i,v_j} = 1$ and $\delta_i = 0$ (i.e., transmission rates of the form $\hat{\beta}_{ij} S_i I_j / \sum_k N_k$), my analysis allows for unequal uptake rates, which corresponds to FDDT model with weighted frequencies, e.g., $\hat{\beta}_{ij} S_i I_j / \sum_k w_k N_k$.

I focus on the case where increased shedding by any host species causes increased infection prevalence in all host species (i.e., $\partial Y_k^* / \partial \chi_{ij} > 0$ for all $i, j, k$). This condition is satisfied by Special Cases 1 and 2 in Section S2.1.1 and I expect it to be met in most systems. When it is not satisfied, the following predictions can be reversed.

**Summary of biological interpretation**: Additions of host species to a community are more likely to cause larger decreases or smaller increases in host 1 infection prevalence under

FDDT than DDDT when (i) the added host species are weak intraspecific and interspecific competitors, (ii) the added host species have lower competence than resident host species, and (iii) the interspecific shedding rates of added host species into the habitats of resident host species are lower than the interspecific shedding rates of resident host species into the habitats of resident host species. In contrast, additions of host species to a community are more likely to cause larger decreases or smaller increases in host 1 infection prevalence under DDDT than FDDT when (iv) the added host species are strong intraspecific and interspecific competitors, (v) the added host species have higher competence than resident host species, and (vi) the interspecific shedding rates of added host species into the habitats of resident host species are greater than the interspecific shedding rates of resident host species into the habitats of resident host species.

In many cases, the sensitivities of host 1 infection prevalence and host 1 infected density have the same sign. The signs of the sensitivities of host 1 infection prevalence and host 1 infected density can differ when (i) host 1 is more strongly regulated by disease than by intraspecific competition and (ii) host 1 is a sink or small source for habitat $j > 1$ and some subset of hosts $k > 1$ are large sources for habitat $j$. The justifications for all of the predictions are given after the calculations.

**Adding host species to a community**: Without loss of generality, assume host species $1, ..., m$ ($m < n$) are present in a community. Assume the species stably coexist at a disease-free equilibrium ($p'$; hereafter the 'subcommunity equilibrium). Let $N_i'$, $Y_i'$, and $P_i'$ denote the densities and prevalences at the equilibrium $p'$. Let $\mathbf{J}'$ denote the Jacobian evaluated at $p'$ and denote its submatrices using primes. I assume the competitive interactions between those species are consistent with stability. Mathematically, I assume $\text{sign}(|\mathbf{J}'|) = (-1)^{3m}$ and $\text{sign}(|\mathbf{A}'|) = (-1)^m$.

Each infectious propagule equation in the $n$-habitat NYZ model (S145) is rewritten as

$$\frac{dP_i}{dt} = \sum_j \chi_{ij} I_j - g_i(q) \sum_{j=1}^{n} \left( u_{i,S_j} S_i + u_{i,I_j} I_i + u_{i,R_j} R_i \right) P_i - h_i(q)\delta_i P_i \tag{S315}$$

where the change of parameters functions are defined by

$$g_i(q) = \begin{cases} \dfrac{-\delta_i}{U_i'} q + 1 + \dfrac{\delta_i}{U_i'}, & 0 \le q \le 1 \\ 2 - q, & 1 \le q \le 2 \end{cases} \tag{S316}$$

$$h_i(q) = \begin{cases} q, & 0 \le q \le 1 \\ \dfrac{-U_i'}{\delta_i}(2 - q) + 1 + \dfrac{U_i'}{\delta_i}, & 1 \le q \le 2 \end{cases} \tag{S317}$$

These functions transform the ET model from a FDDT form when $q = 0$ to a DDDT form when $q = 2$. The transformation holds constant the densities and frequencies at the $m$-host subcommunity equilibrium ($p'$).

Similar to the calculations for $\partial Y_1^*/\partial \chi_{ij}$ and $\partial I_1^*/\partial \chi_{ij}$ in equations (S270) and (S275), the sensitivities of host 1 infection prevalence and host 1 total density at the community equilibrium to the parameter $q$ are

$$0 \leq q \leq 1 : \frac{\partial Y_1^*}{\partial q} = \prod_{k \leq n} \beta_k X_k^* \sum_{i \leq n} \frac{\delta_i P_i^*}{U_i'} \left( U_i^* - U_i' \right) \frac{(-1)^{n+i} Y_i^* N_i^*}{|\mathbf{J}|} \begin{vmatrix} \mathbf{A} & (\mathbf{B} + \mathbf{C}\mathbf{D}_\Gamma)^{[-,\not{1}]} \\ \mathbf{E}_N^{[\not{i},-]} & (\mathbf{E}_Y - \mathbf{D}_{(U+\delta)P/XY})^{[\not{i},\not{1}]} \end{vmatrix} \tag{S318}$$

$$1 \leq q \leq 2 : \frac{\partial Y_1^*}{\partial q} = \prod_{k \leq n} \beta_k X_k^* \sum_{i \leq n} \delta_i \left( U_i^* - U_i' \right) \frac{(-1)^{n+i} Y_j^* N_j^*}{|\mathbf{J}|} \begin{vmatrix} \mathbf{A} & (\mathbf{B} + \mathbf{C}\mathbf{D}_\Gamma)^{[-,\not{1}]} \\ \mathbf{E}_N^{[\not{i},-]} & (\mathbf{E}_Y - \mathbf{D}_{(U+\delta)P/XY})^{[\not{i},\not{1}]} \end{vmatrix} \tag{S319}$$

$$0 \leq q \leq 1 : \frac{\partial N_1^*}{\partial q} = \prod_{k \leq n} \beta_k X_k^* \sum_{i \leq n} \frac{\delta_i P_i'}{U_i^*} \left( U_i^* - U_i' \right) \frac{(-1)^{n+i} Y_i^* N_i^*}{|\mathbf{J}|}$$
$$\times \begin{vmatrix} -(\mathbf{B} + \mathbf{C}\mathbf{D}_\Gamma)_{\bullet 1} & \mathbf{A}^{[-,\not{1}]} & (\mathbf{B} + \mathbf{C}\mathbf{D}_\Gamma)^{[-,\not{1}]} \\ (\mathbf{E}_Y - \mathbf{D}_{(U+\delta)P/XY})_{\bullet 1}^{[\not{i}]} & \mathbf{E}_N^{[\not{i},\not{1}]} & (\mathbf{E}_Y - \mathbf{D}_{(U+\delta)P/XY})^{[\not{i},\not{1}]} \end{vmatrix} \tag{S320}$$

$$1 \leq q \leq 2 : \frac{\partial N_1^*}{\partial q} = \prod_{k \leq n} \beta_k X_k^* \sum_{i \leq n} \delta_i \left( U_i^* - U_i' \right) \frac{(-1)^{n+i} Y_j^* N_j^*}{|\mathbf{J}|} \tag{S321}$$
$$\times \begin{vmatrix} -(\mathbf{B} + \mathbf{C}\mathbf{D}_\Gamma)_{\bullet 1} & \mathbf{A}^{[-,\not{1}]} & (\mathbf{B} + \mathbf{C}\mathbf{D}_\Gamma)^{[-,\not{1}]} \\ (\mathbf{E}_Y - \mathbf{D}_{(U+\delta)P/XY})_{\bullet 1}^{[\not{i}]} & \mathbf{E}_N^{[\not{i},\not{1}]} & (\mathbf{E}_Y - \mathbf{D}_{(U+\delta)P/XY})^{[\not{i},\not{1}]} \end{vmatrix} \tag{S322}$$

where the derivatives at $q = 1$ are interpreted as right and left derivatives. The sensitivity of host 1 infected density is computed using equation (S232).

**Removing species from a community with $n$ habitats**: The notation is the same as the above, where primes denote quantities evaluated at the subcommunity equilibrium. The infectious propagule equations are rewritten as in equation (S315) where the change of parameters functions are defined by

$$g_i(q) = \begin{cases} \dfrac{-\delta_i}{U_i^*} q + 1 + \dfrac{\delta_i}{U_i^*}, & 0 \leq q \leq 1 \\ 2 - q, & 1 \leq q \leq 2 \end{cases} \tag{S323}$$

$$h_i(q) = \begin{cases} q, & 0 \leq q \leq 1 \\ \dfrac{-U_i^*}{\delta_i}(2 - q) + 1 + \dfrac{U_i^*}{\delta_i}, & 1 \leq q \leq 2 \end{cases} \tag{S324}$$

These functions transform the ET model from a FDDT form when $q = 0$ to a DDDT form when $q = 2$. The transformation holds constant the host densities and frequencies at the $n$-host community equilibrium ($p^*$).

The sensitivity of host 1 infection prevalence at the subcommunity equilibrium to the parameter $q$ is

$$0 \leq q \leq 1 : \frac{\partial Y_1'}{\partial q} = \prod_{k \leq m} \beta_k X_k' \sum_{i \leq m} \frac{\delta_i P_i'}{U_i^*} \left( U_i^* - U_i' \right) \frac{(-1)^{m+i+1} Y_j' N_j'}{|\mathbf{J}'|} \begin{vmatrix} \mathbf{A}' & (\mathbf{B}' + \mathbf{C}'\mathbf{D}_\Gamma)^{[-,\not{1}]} \\ \mathbf{E}_N'^{[\not{i},-]} & (\mathbf{E}_Y' - \mathbf{D}_{(U+\delta)P/XY})^{[\not{i},\not{1}]} \end{vmatrix} \tag{S325}$$

$$1 \leq q \leq 2 : \frac{\partial Y_1'}{\partial q} = \prod_{k \leq m} \beta_k X_k' \sum_{i \leq m} \delta_i \left(U_i^* - U_i'\right) \frac{(-1)^{m+i+1} Y_j' N_j'}{|\mathbf{J}'|} \begin{vmatrix} \mathbf{A}' & (\mathbf{B}' + \mathbf{C}'\mathbf{D}_\Gamma)^{[-,\not{1}]} \\ \mathbf{E}_N'^{[\not{i},-]} & (\mathbf{E}_Y' - \mathbf{D}_{(U+\delta)P/XY})^{[\not{i},\not{1}]} \end{vmatrix} \tag{S326}$$

$$0 \leq q \leq 1 : \frac{\partial N_1'}{\partial q} = \prod_{k \leq m} \beta_k X_k' \sum_{i \leq m} \frac{\delta_i P_i'}{U_i^*} \left(U_i^* - U_i'\right) \frac{(-1)^{m+i+1} Y_j' N_j'}{|\mathbf{J}'|} \times \begin{vmatrix} (\mathbf{B}' + \mathbf{C}'\mathbf{D}_\Gamma)_{\bullet 1} & \mathbf{A}'^{[-,\not{1}]} & (\mathbf{B}' + \mathbf{C}'\mathbf{D}_\Gamma)^{[-,\not{1}]} \\ (\mathbf{E}_Y' - \mathbf{D}_{(U+\delta)P/XY})_{\bullet 1}^{[\not{i}]} & \mathbf{E}_N'^{[\not{i},\not{1}]} & (\mathbf{E}_Y' - \mathbf{D}_{(U+\delta)P/XY})^{[\not{i},\not{1}]} \end{vmatrix} \tag{S327}$$

$$1 \leq q \leq 2 : \frac{\partial N_1'}{\partial q} = \prod_{k \leq m} \beta_k X_k' \sum_{i \leq m} \delta_i \left(U_i^* - U_i'\right) \frac{(-1)^{m+i+1} Y_j' N_j'}{|\mathbf{J}'|} \times \begin{vmatrix} (\mathbf{B}' + \mathbf{C}'\mathbf{D}_\Gamma)_{\bullet 1} & \mathbf{A}'^{[-,\not{1}]} & (\mathbf{B}' + \mathbf{C}'\mathbf{D}_\Gamma)^{[-,\not{1}]} \\ (\mathbf{E}_Y' - \mathbf{D}_{(U+\delta)P/XY})_{\bullet 1}^{[\not{i}]} & \mathbf{E}_N'^{[\not{i},\not{1}]} & (\mathbf{E}_Y' - \mathbf{D}_{(U+\delta)P/XY})^{[\not{i},\not{1}]} \end{vmatrix} \tag{S328}$$

where $U_i^* - U_i'$ is defined in equation (S330). The sensitivity of host 1 infected density is computed using equation (S232).

**Justification for interpretation**: Here, I focus on equations (S318), (S319), (S320), and (S322), which define the responses to host species being additions. The interpretations are identical for equations (S325), (S326), (S327), and (S328), which define the responses to host species removals.

The only differences between equations (S318) and (S319) and equations (S320) and (S322) are the first columns of the matrices. The sensitivities $\partial Y_1^*/\partial q$ and $\partial I_1^*/\partial q$ are likely to have the same sign in many systems. However, the sensitivities can have opposite signs when the first columns of the matrices in equations (S320) and (S322) have entries with different signs or entries that differ in magnitude. Those conditions are discussed in Remark S2.3.

Assume $\partial Y_1^*/\partial \chi_{ij} > 0$ for all $i, j$. This means that increased shedding by any host species causes an increase in focal host infection prevalence. This condition is satisfied by Special Cases 1 and 2, and I expect it to be met in most systems. Under this assumption, the sign of each term in equations (S325) and (S326) is determined by the sign of $U_i^* - U_i'$. That quantity can be rewritten by solving $dI_i/dt|_{p^*} = 0$ and $dI_i/dt|_{p'} = 0$ for $I_i^*$ and $I_i'$, respectively, substituting into $dP_i/dt|_{p^*} = 0$ and $dP_i/dt|_{p'} = 0$, and solving for $U_i^*$ and $U_i'$. That yields

$$U_i^* - U_i' = \sum_{k > m} \frac{\chi_{ik}\beta_k}{m_k + \mu_k + \nu_k} S_k^* + \sum_{k \leq m} \frac{\chi_{ik}\beta_k}{m_k + \mu_k + \nu_k} (S_k^* - S_k') \tag{S329}$$

$$= \sum_{k > m} u_{i,S_k} \mathcal{R}_{0,k}(\infty) \frac{\chi_{ik}}{\chi_{kk}} S_k^* + \sum_{k \leq m} u_{i,S_k} \mathcal{R}_{0,k}(\infty) \frac{\chi_{ik}}{\chi_{kk}} (S_k^* - S_k'). \tag{S330}$$

The quantity $U_i^* - U_i'$ is more likely to be positive when resident host densities are higher in the presence of the added host species ($S_k^* - S_k' > 0$), resident host species have lower competence than added host species ($\mathcal{R}_{0,k}(\infty)$ is smaller for $k \leq m$ than $k > m$), and the ratios of interspecific shedding to intraspecific shedding are smaller for resident host species than added host species ($\chi_{ik}/\chi_{kk}$ is smaller for $k \leq m$ than $k > m$). The quantity $U_i^* - U_i'$ can be negative if resident host densities are higher in the absence of the added host species ($S_k^* - S_k' < 0$), resident host species have higher competence than added host species ($\mathcal{R}_{0,k}(\infty)$ is larger for $k \leq m$ than $k > m$), and the ratios of interspecific shedding to intraspecific shedding are larger for resident host species than added host species ($\chi_{ik}/\chi_{kk}$ is larger for $k \leq m$ than $k > m$).

The above yields the following predictions. Adding host species to a community is more likely to cause larger decreases (or smaller increases) in host 1 infection prevalence under FDDT than DDDT when (i) the added host species have weak intraspecific interactions and weak interspecific interactions on resident host species, (ii) the added host species have lower competence than the resident host species, and (iii) when comparing the ratios of interspecific to intraspecific shedding rates ($\chi_{ij}/\chi_{ii}$), added host species have smaller ratios than resident host species for the habitats of resident hosts. In comparison, adding host species to a community is more likely to cause larger increases (or smaller decreases) in host 1 infection prevalence under FDDT than DDDT when (iv) the added host species have strong intraspecific interactions and strong interspecific interactions on resident host species, (ii) the added host species have higher competence than the resident host species, and (iii) when comparing the ratios of interspecific to intraspecific shedding rates ($\chi_{ij}/\chi_{ii}$), added host species have larger ratios than resident host species for the habitats of resident hosts.

### S3 Application of theory to previous modeling studies

In this section, I apply my sensitivity-based theory to prior studies that have explored the relationships between host species richness and the three metrics of disease. I show how my theory explains the patterns and results in those prior studies. Models from a few studies (Strauss et al. 2015, Dallas et al. 2016) were not analyzed because the parameter values in those studies did not yield an $n$-host endemic equilibrium, which is necessary to compute the sensitivities of infection prevalence and infected density at equilibrium. I also do not address Cortez and Duffy (2021) and Cortez (2021) because the sensitivity results in those studies are mathematically equivalent to the sensitivities of host 1 infection prevalence in the 1-host NY model (S4); see Sections S1.5.2, S1.5.3, and S1.5.4. Studies of direct transmission models are addressed first followed by studies of environmental transmission models; each subsection header lists the study name and the disease metric(s) used in the study.

#### S3.1 Direct Transmission Models

##### S3.1.1 Begon et al. (1992) – $I_1^*/N_1^*$

Begon et al. (1992) analyzed a two-host SI-model with density-dependent direct transmission, where there is no interspecific competition and infected and susceptible individuals have equal reproductive output and equal competitive abilities. This corresponds to the $n$-habitat NY model (S147) where $n = 2$, $u_{i,V_j} = 0$ for all $i, j$ and $V \in \{S, I\}$ because there is DDDT, $\nu_i = \Gamma_i = 0$ because there is no recovered class, $\mathbf{A}_{ij} = 0$ and $\mathbf{B}_{ij} = 0$ for $i \neq j$ because there is no interspecific competition, $\mathbf{B}_{ii} = -\mu N_i^*$, $\mathbf{E}_N[i,j] = \chi_{ij} Y_j^*$, $\mathbf{E}_Y[i,j] = \chi_{ij} N_j^*$, and $|\mathbf{J}| > 0$.

The authors found that the addition of a second host species increased infection prevalence in the first species. This result is explained by equation (S238) in Section S2.5.2. Specifically, under the constraints listed in the previous paragraph, the sensitivity simplifies to

$$\frac{\partial Y_1^*}{\partial \alpha} = \left(-\frac{\partial}{\partial \alpha}\frac{dN_2}{dt}\right)\frac{1}{|\mathbf{J}|}\mathbf{A}_{11}\beta_1 X_1^*\left[\chi_{12}\beta_2 P_2^* \delta_2\right] < 0 \tag{S331}$$

Dividing by $\partial N_2^*/\partial \alpha < 0$ yields that host 1 infection prevalence increases with increased density of host 2 ($dY_1^*/dN_2^* > 0$).

##### S3.1.2 Bowers and Turner (1997) – $\mathcal{R}_0^C$, $I_1^*/N_1^*$

Bowers and Turner (1997) analyzed a two-host SI-model with density-dependent direct transmission, where interspecific competition was present and infected and susceptible individuals have equal reproductive output and equal competitive abilities. This corresponds to the $n$-habitat NY model (S147) where $n = 2$, $u_{i,V_j} = 0$ for all $i, j$ and $V \in \{S, i\}$, $\nu_i = \Gamma_i = 0$ because there is no recovered class, $\mathbf{A}_{ij} < 0$ for all $i, j$, $\mathbf{B}_{ij} = 0$ for $i \neq j$, $\mathbf{B}_{ii} = -\mu N_i^*$, $\mathbf{E}_N[i,j] = \chi_{ij} Y_j^*$, $\mathbf{E}_Y[i,j] = \chi_{ij} N_j^*$, and $|\mathbf{J}| > 0$.

Richness-$\mathcal{R}_0^C$ relationships: The authors did not directly address how host species richness affects $\mathcal{R}_0^C$. However, Figure 4a of Bowers and Turner (1997) shows that the pathogen can invade when only host 1 is present and it cannot invade when hosts 1 and 2 coexist and host

2 is a sufficiently strong interspecific competitor. This means that the addition of host 2 reduced $\mathcal{R}_0^C$ when host 2 is a sufficiently strong interspecific competitor. This agrees with Special Case A of section S2.4.2 where the sensitivity is given by equation (S184),

$$\frac{d\mathcal{R}_0^C}{dN_n^\ddagger} = \mathcal{R}_0^C \sum_k \frac{1}{N_k^\ddagger}\frac{dN_k^\ddagger}{dN_n^\ddagger} y_k x_k. \tag{S332}$$

When host 2 is a sufficiently weak interspecific competitor, the equation is positive, which means host 2 increases $\mathcal{R}_0^C$. However, if host is a sufficiently strong interspecific competitor, then the equation will be negative because $dN_1^\ddagger/dN_2^\ddagger$ is large and negative. This causes the addition of host 2 to decrease $\mathcal{R}_0^C$.

Richness-infection prevalence relationships: The authors did not directly address how host species richness affects focal host infection prevalence. However, they did observe cases where the addition of a second host species caused the pathogen to go extinct. The authors found that introducing host 2 caused the pathogen to go extinct when host 2 was a sufficiently strong interspecific competitor. This result is explained by equation (S238) in Section S2.5.2. Under the constraints listed in the previous paragraph, the sensitivity simplifies to

$$\frac{\partial Y_1^*}{\partial\alpha} = \overbrace{\left(-\frac{\partial}{\partial\alpha}\frac{dN_2}{dt}\right)\frac{(-1)^{1+2^2}}{|\mathbf{J}|}(\beta_1 X_1^*)}^{\text{negative}} \times \left\{ \overbrace{\mathbf{A}_{11}\mathbf{E}_N[1,2]\left(-\frac{\delta_2 P_2^*}{X_2^*Y_2^*}\right)}^{\text{positive}} \underbrace{-\mathbf{A}_{12}\left(-\mathbf{E}_N[1,2]\mathbf{E}_Y[2,1]+\mathbf{E}_N[1,1]\left[\mathbf{E}_Y[2,2]-\frac{\delta_2 P_2^*}{X_2^*Y_2^*}\right]\right)}_{\text{negative}}\right\} \tag{S333}$$

where $\mathbf{E}_Y[2,2] - \delta_2 P_2^*/X_2^*Y_2^* < 0$. Dividing by $\partial N_2^*/\partial\alpha < 0$ yields that host 1 infection prevalence increases when interspecific competition is sufficiently low ($dY_1^*/dN_2^* > 0$ when $\mathbf{A}_{12}$ small in magnitude) and host 1 infection prevalence decreases when interspecific competition is sufficiently high ($dY_1^*/dN_2^* < 0$ when $\mathbf{A}_{12}$ large in magnitude). Hence, if host 2 is a sufficiently strong interspecific competitor, then its introduction can cause host 1 infection prevalence to decrease (including decreasing to extinction).

##### S3.1.3 Dobson (2004) – $\mathcal{R}_0^C$

Dobson (2004) analyzed an $n$-host SIR model with either DDDT or FDDT where there was no interspecific competition. The study assumed the low interspecific transmission rates defined by $\hat{\beta}ij = c_{ij}(\hat{\beta}_{ii} + \hat{\beta}_{jj})/2$ where $c_{ij} \leq 0.05$ for $i \neq j$. The DDDT form of the Dobson (2004) model corresponds to the $n$-habitat model (S144) where $u_{j,V_i} = 0$ for all $i, j$ and $V \in \{S, I, R\}$. The FDDT form of the Dobson (2004) model corresponds to the $n$-habitat model (S144) where $\delta_i = 0$ and $u_{j,V_i} = 1$ for all $i, j$ and $V \in \{S, I, R\}$.

Variation in interspecific transmission rates: Figure 2 of Dobson (2004) shows that $\mathcal{R}_0^C$ increases with increased interspecific transmission (compare $\mathcal{R}_0^C$ values across top 3 and bottom 3 curves). Greater interspecific transmission in the Dobson (2004) models corresponds to larger $\chi_{ij}$ values ($i \neq j$) in the $n$-habitat model (S144). Increased $\mathcal{R}_0^C$ with increased interspecific transmission matches the positive sign of the sensitivity $\partial\mathcal{R}_0/\partial\chi_{ij}$ in Section S2.4.4.

Richness-$\mathcal{R}_0^C$ relationships for DDDT: Figure 2 of Dobson (2004) shows that $\mathcal{R}_0^C$ increases with increased host species richness under DDDT (top three curves increasing). Increased $\mathcal{R}_0^C$ with increased host species richness under DDDT matches the positive sign of $d\mathcal{R}_0/dN_i^\ddagger$ in Special Case A of Section S2.4.2, which assumes DDDT and no interspecific competition.

Richness-$\mathcal{R}_0^C$ relationships for FDDT: Figure 2 of Dobson (2004) shows that $\mathcal{R}_0^C$ decreases with increased host species richness under FDDT (bottom three curves decreasing). Decreased $\mathcal{R}_0^C$ with increased host species richness under FDDT matches the negative sign of $d\mathcal{R}_0/dN_n^\ddagger$ in Special Case C of Section S2.4.2. That special case assumes FDDT, no interspecific competition, equal weights, and very low interspecific transmission, which matches the assumptions in Dobson (2004).

###### S3.1.4 Faust et al. (2017) – $I_1^*/N_1^*$

Faust et al. (2017) studied an $n$-host model with either DDDT or FDDT where interspecific competition was absent. The interspecific direct transmission parameters were defined as the geometric mean of the intraspecific direct transmission parameters for the interacting species ($\hat{\beta}_{ij} = \sqrt{\hat{\beta}_{ii}\hat{\beta}_{jj}}$). This results in interspecific transmission rates that are not small relative to intraspecific transmission rates. The sensitivities for the $n$-habitat model are too complex to analyze in general when interspecific transmission is comparable to intraspecific transmission. Nonetheless, the sensitivity results in this study help explain the numerical results in Faust et al. (2017).

Richness-infection prevalence relationships for DDDT: The right column of Figure 1 and Figure 2a in Faust et al. (2017) show that increased host species richness results in higher infection prevalence for all host species when there is DDDT. Increased prevalence with increased host species richness matches the prediction from Special Case 2 of Section S2.5.2 when there is no interspecific competition, DDDT, and low interspecific transmission; see equation (S254). Increased prevalence with increased host species richness also matches the prediction from the 1-habitat model. Specifically, under the assumptions of no interspecific competition and DDDT, the formula for $dY_1^*/dN_n^*$ in equation (S74) of Section S1.5.2 reduces to $dY_1^*/dN_n^* = Q'\varphi_n > 0$ where $Q'$ is a positive constant and $\varphi_n > 0$.

Richness-infection prevalence relationships for FDDT: The middle column of Figure 1 and Figure 2b,c in Faust et al. (2017) show that increased host species richness results in lower infection prevalence for all host species under FDDT, unless the added host species have very high intraspecific transmission parameters (which also implies high interspecific transmission parameters). Increased prevalence with increased host species richness matches the prediction from Special Case 2 of Section S2.5.2 when there is no interspecific competition, FDDT, and low interspecific transmission; see equation (S253) and surrounding text. The patterns in Faust et al. (2017) also match the prediction from the 1-habitat model. In particular, under the assumption of no interspecific competition, the formula for $dY_1^*/dN_n^*$ in equation (S74) of Section S1.5.2 reduces to $dY_1^*/dN_n^* = Q'\varphi_n$ where $Q'$ is a positive constant. If host $n$ has a sufficiently low intraspecific transmission rate, then $\chi_n$ will be small and $\varphi_n$ will be negative. From this I predict decreases host 1 infection prevalence when host $n$ is added to the community. On the other hand, if host $n$ has a sufficiently high intraspecific transmission rate, then $\chi_n$ will be large and $\varphi_n$ will be positive. From this I predict increases host 1 infection prevalence when host $n$ is added to the community. These predictions align with what was observed in the FDDT simulations in Faust et al. (2017).

##### S3.1.5 Huang et al. (2019) – $\mathcal{R}_0^C$

Huang et al. (2019) analyzed a three-host SEIR model with DDDT where there is no interspecific host competition. Hosts 1 and 2 can transmit the disease whereas host 3 is a dead end host that cannot transmit the pathogen. The authors analytically computed $\mathcal{R}_0^C$ on page 947 of that study. To apply the sensitivity formulas from this study, I first reduce the dimension of the Huang et al. (2019) model by removing the exposed classes via a quasi-steady-state approximation. This does not affect the value of $\mathcal{R}_0^C$ and puts the model in a form that corresponds to the $n$-habitat model (S144) where $u_{j,S_i} = 0$ for all $i, j$. Also, because host 3 is a dead end host that cannot transmit the disease, $\mathcal{R}_0^C$ and its sensitivities are computed using $n = 2$.

Variation in recruitment rates: Figure 5a-c of Huang et al. (2019) shows that $\mathcal{R}_0^C$ increases with increased recruitment rates of either host. In the Huang et al. (2019) model, increased recruitment of a host is mathematically equivalent to decreased $\alpha$ in our sensitivities to host density. This means that the effects of increased recruitment are predicted by $-\partial\mathcal{R}_0^C/\partial\alpha$, whose formula is given in equation (S183) of Section S2.4.2. Because there is no interspecific competition, the disease-free equilibrium densities satisfy $\partial N_j^\ddagger/\partial\alpha < 0$ when $\alpha$ is the recruitment rate of host $j$ and $\partial N_j^\ddagger/\partial\alpha = 0$ otherwise. Thus, under the assumptions of DDDT and no interspecific competition, equation (S183) simplifies to

$$\frac{\partial\mathcal{R}_0^C}{\partial\alpha} = \mathcal{R}_0^C \frac{1}{N_j^\ddagger}\frac{\partial N_j^\ddagger}{\partial\alpha} y_j x_j < 0 \tag{S334}$$

where $\alpha$ is the competition coefficient for host $j$. Because increased reproduction implies reduced $\alpha$, the sensitivity is positive.

Variation in vaccination rates: Figure 5b of Huang et al. (2019) shows that $\mathcal{R}_0^C$ decreases with increased vaccination rates. In the Huang et al. (2019) model, an increased vaccination rate for host $j$ is mathematically equivalent to increased $\alpha$ in our sensitivities to host density. The formula for the sensitivity is the same as the above. Because increased vaccination implies increased $\alpha$, the sensitivity is negative.

Variation in transmission rates: Figure 4 shows that $\mathcal{R}_0^C$ increases with increased transmission. Increased transmission rates in the Huang et al. (2019) model corresponds to larger values of $\chi_{ij}$. As shown in Section S2.4.4, the sensitivity of $\mathcal{R}_0^C$ to $\chi_{ij}$ is always positive for all $i, j$.

##### S3.1.6 Joseph et al. (2013) – $\mathcal{R}_0^C$

Joseph et al. (2013) analyzed an $n$-host models with DDDT or FDDT. The DDDT model corresponds to the $n$-habitat model (S144) where $u_{i,V_j} = 0$ for all $i, j$ and $V \in \{S, I, R\}$ and the FDDT model corresponds to the $n$-habitat model (S144) where $\delta_i = 0$ and $u_{j,S_i} = 1$ for all $i, j$ and $V \in \{S, I, R\}$. In that study, "subtractive extirpation" implies no interspecific competition and "compensatory extirpation" implies interspecific competition being present. The simulations measured how $\mathcal{R}_0^C$ changed when a host species was removed from a five-host community. In the simulations, species varied in body size and transmission rates ($\hat{\beta}_{ii}$ in that study) and treatments differed in terms of which host was removed and whether the relationship between body size and host competence was negative, neutral, or positive. The study considered interspecific transmission rates that were equal to or less than the average of the intraspecific transmission rates ($\hat{\beta}_{ij} = c_{ij}(\hat{\beta}_{ii} + \hat{\beta}_{jj})/2$ for $0 \leq c_{ij} \leq 1$). Because my theory is for deterministic systems, I only focus on the simulations with "deterministic removal" wherein host species with the largest body size were removed.

Variation in interspecific transmission rates: Figure 3 shows increasing interspecific transmission rates ($c_{ij}$, $i \neq j$) resulted in bigger decreases in R0 as species are removed. This is consistent with my theory since R0C is an increasing function of $c_{ij}$, which corresponds to $\chi_{ij}$ in the $n$-habitat model.

Richness-$\mathcal{R}_0^C$ relationships for DDDT - no interspecific competition: Consider the results in Figure 2 of Joseph et al. (2013) for DDDT systems with subtractive extirpations (i.e., no interspecific competition). Subtractive extirpations always decrease in $\mathcal{R}_0^C$ (top right panel of Figure 2) and the decreases in $\mathcal{R}_0^C$ are larger with positive host competence-body size relationships (meaning high competence hosts are more likely to be removed; top right panel of Figure 3). This matches the predictions from the sensitivity equation (S184) in Special Case A of Section S2.4.2 for DDDT models where interspecific competition is absent. Specifically, when interspecific competition is absent, $d\mathcal{R}_0^C/dN_n^\ddagger$ is positive and proportional to $x_n y_n$, which is larger when host $n$ has higher competence.

Richness-$\mathcal{R}_0^C$ relationships for DDDT - yes interspecific competition: Consider the results in Figure 2 of Joseph et al. (2013) for DDDT systems with compensatory extirpations (which implies there is interspecific competition). Compensatory extirpations are more likely to increase $\mathcal{R}_0^C$ under negative host competence-body size relationships (left side of bottom right panel of Figure 2) and are more likely to decrease $\mathcal{R}_0^C$ under positive host competence-body size relationships (right side of bottom right panel of Figure 2). Equation (S184) in Special Case A of Section S2.4.2 can explain the shift form positive to negative changes in $\mathcal{R}_0^C$ (also shown in the bottom right panel of Figure 3). When the density of host $n$ is decreased, the value of $\mathcal{R}_0^C$ decreases at rate $\mathcal{R}_0 x_n y_n/N_n^\ddagger$ due to reduced density of host $n$ and increases at rate $\mathcal{R}_0 x_k y_k[d\ln(N_k^\ddagger)/dN_n^\ddagger]$ due to increased density of hosts $k \neq n$. For the simulations with negative host competence-body size relationships, low competence hosts are lost which means that $x_n y_n$ is small relative to $x_k y_k$ ($k \neq n$). Consequently, $\partial\mathcal{R}_0^C/\partial N_n^\ddagger$ is negative and $\mathcal{R}_0^C$ will increase as host $n$ is removed. For the simulations with positive host competence-body size relationships, high competence hosts are lost which means that $x_n y_n$ is larger than $x_k y_k$ ($k \neq n$). In this case, $\partial\mathcal{R}_0^C/\partial N_n^\ddagger$ is positive or negative depending on how much larger $x_n y_n$ is relative to $x_k y_k$ and the strength of interspecific competition. Hence, $\mathcal{R}_0^C$ is more likely to increase when high competence hosts are removed if interspecific competition is stronger and $\mathcal{R}_0^C$ is more likely to decrease when high competence hosts are removed if interspecific competition is weaker.

Richness-$\mathcal{R}_0^C$ relationships for FDDT - no interspecific competition: Consider the results in Figure 2 of Joseph et al. (2013) for FDDT systems with subtractive extirpations (i.e., no interspecific competition). Subtractive extirpations almost always increase $\mathcal{R}_0^C$ under negative host competence-body size relationships (left side of top left panel of Figure 2). In contrast, subtractive extirpations increase or decrease $\mathcal{R}_0^C$ under positive host competence-body size relationships (right side of top left panel of Figure 2). These patterns can be explained by the sensitivity equation (S186) in Special Case B of Section S2.4.2 for FDDT models with no interspecific competition and equal weights (i.e., $u_{i,V_j} = 1$). For negative host competence-body size relationships, the low competence hosts being removed have small $x_n y_n$ values. This means equation (S186) is negative, which results in $\mathcal{R}_0^C$ increasing as host $n$ is decreased toward zero and ultimately removed. For positive host competence-body size relationships, the high competence hosts being removed have larger $x_n y_n$ values. In this case, equation (S186) is positive when the removed host species are initially at low frequency in the community ($N_n^\ddagger / \sum_k N_k^\ddagger$ small) whereas equation (S186) is negative when the removed host species are initially at high frequency in the community ($N_i^\ddagger / \sum_k N_k^\ddagger$ large). Thus, under positive host competence-body size relationships, removal of a host species can increase or decrease $\mathcal{R}_0^C$ depending on its density and frequency in the community. This explains different patterns in the left and right sides of the top left panel of Figure 2 in Joseph et al. (2013).

Richness-$\mathcal{R}_0^C$ relationships for FDDT - yes interspecific competition: Consider the results in Figure 2 of Joseph et al. (2013) for FDDT systems with compensatory extirpations (which implies there is interspecific competition). Removal of a host species increases $\mathcal{R}_0^C$ more often than not (bottom left of Figure 2). This pattern cannot be explained fully because equation (S185) is too complex to analyze in full detail. However, insight about potential driving mechanisms can be gained by analyzing equation (S187) in Special Case C of Section S2.4.2, which assumes FDDT, equal weights, and very low interspecific transmission. Equation (S187) shows that the sensitivity is $d\mathcal{R}_0^C / dN_n^\ddagger$ is negative, which implies removal of host species will cause increases in $\mathcal{R}_0^C$. The sign of equation (S187) can change if interspecific transmission is sufficiently high. Thus, I predict that removals increased $\mathcal{R}_0^C$ when interspecific transmission was low and increased $\mathcal{R}_0^C$ when interspecific transmission was higher. In addition, removals were more likely to increase $\mathcal{R}_0^C$ under negative host competence-body size relationships because of the $n$th term in equation (S185) is more negative under negative host competence-body size relationships than positive host competence-body size relationships; see previous paragraph.

##### S3.1.7 McCormack and Allen (2007) – $\mathcal{R}_0^C$

McCormack and Allen (2007) analyzed an $n$-host SIS model with either DDDT or FDDT where there is no interspecific competition. The DDDT form of the model corresponds to the $n$-habitat model (S144) where $u_{j,V_i} = 0$ for all $i, j$ and $V \in \{S, I, R\}$. The FDDT form of the model corresponds to the $n$-habitat model (S144) where $\delta_i = 0$ and $u_{i,V_i} = 0$ for all $i$ and $V \in \{S, I, R\}$, and $u_{j,S_i} = 0$ for $i \neq j$.

Richness-$\mathcal{R}_0^C$ relationships for DDDT: Theorem 3.2 of McCormack and Allen (2007) shows that for the DDDT form of the model, $\mathcal{R}_0^C$ increases whenever a new host is added to the community. This is explained by Special Case A of Section S2.4.2. In particular, under DDDT and no interspecific competition, $d\mathcal{R}_0^C / dN_n^\ddagger > 0$. Hence, adding a species to a community will increase $\mathcal{R}_0^C$.

Richness-$\mathcal{R}_0^C$ relationships for FDDT: Theorem 3.2 of McCormack and Allen (2007) also shows that for the FDDT form of the model, $\mathcal{R}_0^C$ increases whenever a new host is added to the community. In comparison, the sensitivity of $\mathcal{R}_0^C$ in Special Case D of Section S2.4.2 shows that $d\mathcal{R}_0/dN_n^\ddagger = 0$. These two results do no contradict because $\mathcal{R}_0^C$ is a piecewise-constant function of host density. Said another way, $\mathcal{R}_0^C$ is discontinuous at $N_n^\ddagger = 0$ and constant for all positive values of $N_i^\ddagger$. This results in $\mathcal{R}_0^C$ increasing when a host is added to the community (the result in McCormack and Allen (2007)), but $\mathcal{R}_0^C$ being independent of host density, provided the host species is present (see the result in Special Case D of Section S2.4.2).

##### S3.1.8 Mihaljevic et al. (2014) – $\mathcal{R}_0^C$

Mihaljevic et al. (2014) analyzed an $n$-host models with DDDT or FDDT. The DDDT model corresponds to the $n$-habitat model (S144) where $u_{j,V_i} = 0$ for all $i, j$ and $V \in \{S, I, R\}$ and the FDDT model corresponds the same model where $\delta_i = 0$ and $u_{j,V_i} = 1$ for all $i, j$. In that study, the relationship between host species richness and total host abundance defines the effects of interspecific competition: "additive" corresponds to no interspecific competition and "compensatory" and "saturating" correspond to interspecific competition being present and possibly depending on the number of species present in the community. In the simulations, most species had $\mathcal{R}_{0,i} < 1$ and the interspecific direct transmission parameters were small compared to the intraspecific direct transmission parameters ($\hat{\beta}_{ij} = c_{ij}(\hat{\beta}_{ii} + \hat{\beta}_{jj})/2$ where $c_{ij} = 0.05$ for $i \neq j$).

The sensitivity results in this study can explain the patterns for DDDT systems in Figure 2A-D of Mihaljevic et al. (2014).

Richness-$\mathcal{R}_0^C$ relationships for DDDT - no interspecific competition: Consider the results in Figure 2A of Mihaljevic et al. (2014) for DDDT systems without interspecific competition (the "additive" case). The relationship between $\mathcal{R}_0^C$ and host species richness is always positive. This is explained by equation (S184) in Special Case A of Section S2.4.2. Specifically, in the absence of interspecific competition, $d\mathcal{R}_0^C/dN_n^\ddagger$ is always positive, which means host additions will always increase $\mathcal{R}_0^C$.

Richness-$\mathcal{R}_0^C$ relationships for DDDT - yes interspecific competition: Consider the results in Figure 2B-D of Mihaljevic et al. (2014) for DDDT systems with interspecific competition (the "compensatory" and "saturating" cases). The relationship between $\mathcal{R}_0^C$ and host species richness is positive when there is a steep relationship between host species richness and total host density (left sides of Figure 2CD). In comparison, the relationship between $\mathcal{R}_0^C$ and host species richness is negative when there is a shallow relationship between host species richness and total host density (Figure B and right sides of Figure 2CD). These patterns are explained by equation (S184) in Special Case A of Section S2.4.2. When there is a steep relationship between host species richness and total host abundance, increases in density of host $n$ will cause small decreases in the density of host $k \neq n$ ($dN_k^\ddagger/dN_n^\ddagger$ negative and small in magnitude). This means that equation (S184) has a positive term for $k = n$ and small negative terms for $k \neq n$. This results in equation (S184) being positive and $\mathcal{R}_0^C$ increasing when species are added to the community. In contrast, when there is a shallow relationship between host species richness and total host density, increases in density of host $n$ will cause large decreases in the density of host $k \neq n$ ($dN_k^\ddagger/dN_n^\ddagger$ negative and large in magnitude). This means that equation (S184) has a positive term for $k = n$ and large negative terms for

$k \neq n$. This results in equation (S184) being negative and $\mathcal{R}_0^C$ decreasing when species are added to the community.

Richness-$\mathcal{R}_0^C$ relationships for FDDT - no interspecific competition: Consider the results in Figure 2E of Mihaljevic et al. (2014) for FDDT systems without interspecific competition (the "additive" case). The relationship between $\mathcal{R}_0^C$ and host species richness is negative. This pattern is explained by equation (S187) in Special Case C of Section S2.4.2. Specifically, the authors assume most species have $\mathcal{R}_{0,i} < 1$ and very low interspecific transmission coefficients. This in combination with the assumption of no interspecific competition means that equation (S187) only has as single term (for the added host) and it is negative. Thus, increased density of a host species causes a decrease in $\mathcal{R}_0^C$.

Richness-$\mathcal{R}_0^C$ relationships for FDDT - yes interspecific competition: Consider the results in Figure 2F of Mihaljevic et al. (2014) for FDDT systems with interspecific competition (the "saturating" case). The relationship between $\mathcal{R}_0^C$ and host species richness is negative. Because the authors assume most species have $\mathcal{R}_{0,i} < 1$ and very low interspecific transmission coefficients, the patterns satisfy the conditions of Special Case C of Section S2.4.2. Thus, equation (S187) is negative, which means increased density of any species causes a decrease in $\mathcal{R}_0^C$.

##### S3.1.9 O'Regan et al. (2015) – $\mathcal{R}_0^C$

O'Regan et al. (2015) analyzed a two-host SIS model with either DDDT or FDDT. The DDDT version of the model corresponds to the $n$-habitat model (S144) where $u_{j,V_i} = 0$ for all $i, j$ and $V \in \{S, I, R\}$. The FDDT version of the model corresponds to the $n$-habitat model (S144) where $\delta_i = 0$ and $u_{j,V_i} = 1$ for all $i, j$. Note that "additive" density regulation corresponds to no interspecific competition and "substitutive" density regulation corresponds to strong, symmetric interspecific competition.

Variation in transmission and competence: Figure 3 of O'Regan et al. (2015) shows that $\mathcal{R}_0^C$ increases with increased $b$ and decreases with increased $\sigma$. When translated to the notation in this study, the variation in those parameters corresponds to increases in $\chi_{ij}$ $(i \neq j)$ and decreases in $\mathcal{R}_{0,2}$ (e.g., decreases in $\beta_2$). As shown in Section S2.4.4, increases in $\chi_{ij}$ increase $\mathcal{R}_0^C$ because $\partial\mathcal{R}_0^C/\partial\chi_{ij} > 0$ and decreases in $\beta_i$ decrease $\mathcal{R}_0^C$ because $\partial\mathcal{R}_0^C/\partial\beta_i > 0$.

Richness-$\mathcal{R}_0^C$ relationships for DDDT - no interspecific competition: Table 2 of O'Regan et al. (2015) predicts that if there is additive density regulation (i.e., no interspecific competition) and DDDT, then the addition of a second host always increases $\mathcal{R}_0^C$. This prediction can be explained by the sensitivity of $\mathcal{R}_0^C$ to the density of host 2 in Special Case A of Section S2.4.2. For the two-host model in O'Regan et al. (2015), the sensitivity $d\mathcal{R}_0^C/dN_2^\ddagger$ in equation (S184) reduces to

$$\frac{d\mathcal{R}_0^C}{dN_2^\ddagger} = \mathcal{R}_0^C \left( \frac{y_2 x_2}{N_2^\ddagger} + \frac{y_1 x_1}{N_1^\ddagger} \frac{dN_1^\ddagger}{dN_2^\ddagger} \right). \tag{S335}$$

When there is no interspecific competition, the derivative $dN_1^\ddagger/dN_2$ is zero. This results in equation (S335) being positive, which means the addition of host 2 increases $\mathcal{R}_0^C$.

Richness-$\mathcal{R}_0^C$ relationships for DDDT - yes interspecific competition: Table 2 of O'Regan et al. (2015) predicts that if there is substitutive density regulation (i.e., interspecific competition is present) and DDDT, then the addition of a second host increases $\mathcal{R}_0^C$ unless host 2 has a sufficiently low $\mathcal{R}_{0,2}$ (i.e., sufficiently low competence) and interspecific competition is sufficiently strong (i.e., $\alpha_{12}, \alpha_{21}$ are sufficiently large). This prediction also can be explained by the sensitivity of $\mathcal{R}_0^C$ to the density of host 2 in Special Case A of Section S2.4.2. The sensitivity $d\mathcal{R}_0^C/dN_2^\ddagger$ is defined as in the previous equation. When interspecific competition is present, the derivative $dN_1^\ddagger/dN_2$ is negative. In this case, the sensitivity is more likely to be negative when $y_2x_2$ is smaller, $y_1x_1$ is larger, and $dN_1^\ddagger/dN_2^\ddagger$ larger in magnitude. Biologically, these conditions mean host 2 has lower competence, host 1 has higher competence, and host 2 is a stronger interspecific competitor. This matches the result from O'Regan et al. (2015) where the addition of host 2 reduced $\mathcal{R}_0^C$ when host 2 had low competence and was a strong interspecific competitor.

Richness-$\mathcal{R}_0^C$ relationships for FDDT - no interspecific competition: Table 2 of O'Regan et al. (2015) predicts that if there is additive density regulation (i.e., no interspecific competition) and FDDT, then the addition of a second host decreases $\mathcal{R}_0^C$ unless host 2 has a high $\mathcal{R}_{0,i}$ or interspecific transmission is sufficiently large compared to intraspecific transmission ($b = \beta_{12}\beta_{21}/\beta_{11}\beta_{22}$ large in magnitude). This prediction can be explained by the sensitivity of $\mathcal{R}_0^C$ to the density of host 2 in Special Case B of Section S2.4.2. When there is no in specific competition, the sensitivity $d\mathcal{R}_0^C/dN_2^\ddagger$ in equation (S186) reduces to

$$\frac{d\mathcal{R}_0^C}{dN_2^\ddagger} = \frac{\mathcal{R}_0^C}{N_2^\ddagger}\left[x_2y_2 - \frac{N_1^\ddagger}{N_1^\ddagger + N_2^\ddagger}\right]. \tag{S336}$$

The equation is negative unless $x_2y_2$ is sufficiently large. This means equation is positive only if host 2 has higher competence (which also implies a higher $\mathcal{R}_{0,i}$) and higher interspecific transmission rate; see Section S2.4.3. Thus, the addition of host 2 will decrease $\mathcal{R}_0^C$ unless host 2 has high competence (high $\mathcal{R}_{0,i}$) or there is sufficiently high interspecific transmission.

Richness-$\mathcal{R}_0^C$ relationships for FDDT - yes interspecific competition: Table 2 of O'Regan et al. (2015) predicts that if there is substitutive density regulation (i.e., strong interspecific competition) and FDDT, then the addition of a second host decreases $\mathcal{R}_0^C$ when intraspecific transmission is lower than interspecific transmission and host 2 is has a lower $\mathcal{R}_{0,i}$. This prediction also can be explained by the sensitivity of $\mathcal{R}_0^C$ to the density of host 2 in Special Case B of Section S2.4.2. When there is interspecific competition, the sensitivity $d\mathcal{R}_0^C/dN_2^\ddagger$ in equation (S185) reduces to

$$\frac{d\mathcal{R}_0^C}{dN_2^\ddagger} = \mathcal{R}_0^C\left(\frac{1}{N_1^\ddagger}\frac{dN_1^\ddagger}{dN_2^\ddagger}\left[x_1y_1 - \frac{N_1^\ddagger}{N_1^\ddagger + N_2^\ddagger}\right] + \frac{1}{N_2^\ddagger}\left[x_2y_2 - \frac{N_2^\ddagger}{N_1^\ddagger + N_2^\ddagger}\right]\right). \tag{S337}$$

Because $dN_1^\ddagger/dN_2^\ddagger < 0$, the equation is negative when $x_1y_1$ is large and $x_2y_2$ is small. This occurs when host 1 has higher competence, host 2 has lower competence, and interspecific transmission is lower. Hence, the addition of host species 2 decreases $\mathcal{R}_0^C$ when host species 2 has lower competence and there is low interspecific transmission.

##### S3.1.10 Peixoto and Abramson (2006) – $I_1^*/N_1^*$

Peixoto and Abramson (2006) studied a two-host model with density-dependent direct transmission. Their model corresponds to the $n$-habitat model (S144) where $u_{j,V_i} = 0$ for all $i, j$

and $V \in \{S, I, R\}$. The authors assume there is interspecific host competition but host 2 cannot be infected by the pathogen. This means $\chi_{12} = \chi_{22} = 0$, $\beta_2 = 0$, and there is no $Y_2$ variable. Consequently, matrix $\mathbf{B}$ of the Jacobian does not have a second column and matrices $\mathbf{E}_N$ and $\mathbf{E}_Y$ do not have second rows; the other matrices are adjusted accordingly.

Variation in host 2 carrying capacity: Figure 3 shows that host 1 infection prevalence decreases increases in the carrying capacity of host 2. Increases in host 2 carrying capacity are mathematical equivalent to decreases in the parameter $\alpha$ in equation (S238) of Section (S2.4.2). Under the conditions listed in the previous paragraph, that equation simplifies to

$$\frac{dY_1^*}{d\alpha} = \left(-\frac{\partial}{\partial \alpha}\frac{dN_2}{dt}\right)\frac{(-1)^{1+2^2}}{|\mathbf{J}|}\beta_1 X_1 \left| \begin{bmatrix} \mathbf{A}_{11} & \mathbf{A}_{12} & \mathbf{B}_{11} \\ \mathbf{A}_{21} & \mathbf{A}_{22} & 0 \\ \chi_{11} Y_1^* & 0 & \chi_1 N_1^* - \delta_1 P_1^*/X_1^* Y_1^* \end{bmatrix}^{[2,3]} \right| \tag{S338}$$

$$= \left(-\frac{\partial}{\partial \alpha}\frac{dN_2}{dt}\right)\frac{-1}{|\mathbf{J}|}\beta_1 X_1 \mathbf{A}_{12} \chi_{11} Y_1^* \tag{S339}$$

Note that $|\mathbf{J}| > 0$, $\mathbf{A}_{12} < 0$, $\left(\frac{\partial}{\partial K}\frac{dN_2}{dt}\right) > 0$, and $\partial N_2^*/\partial K_2 > 0$. Consequently, the equation is negative and host 1 infection prevalence decreases with increases in the carrying capacity of host 2.

Richness-infection prevalence relationships: Figure 2 of Peixoto and Abramson (2006) shows that host 1 infection prevalence decreases with increases in the density of host 2. That pattern is explained by the equation from above. In particular, dividing by $\partial N_2^*/\partial \alpha < 0$ yields $dY_1^*/dN_1^* = [\partial Y_1^*/\partial K_2]/[\partial N_2^*/\partial K_2] > 0$.

##### S3.1.11 Roberts and Heesterbeek (2018) – $\mathcal{R}_0^C$, $I_1^*/N_1^*$, $I_1^*$

Roberts and Heesterbeek (2018) considered the same three metrics of disease ($\mathcal{R}_0^C$, $Y_1^*$, and $I_1^*$) and showed how elasticities of those metrics could be used to explore when the host species richness has similar or different effects on the metrics. The analysis in Roberts and Heesterbeek (2018) focused on a two-host model with DDDT or FDDT, a two-host ET model without uptake of infectious propagules, and an $n$-host model with DDDT or FDDT. The results in Roberts and Heesterbeek (2018) and the current study are related because if $dX^*/dY^*$ is the sensitivity of $X^*$ to $Y^*$, then $[Y^*/X^*][dX^*/dY^*]$ is the elasticity of $X^*$ to $Y^*$.

The sensitivity formulas for $d\mathcal{R}_0/dN_i^\ddagger$, $dY_1^*/dN_i^*$, and $dI_1^*/dN_i^*$ in this study generalize the elasticity formulas in Roberts and Heesterbeek (2018). The following explains the relations between the formulas in this study and those in Roberts and Heesterbeek (2018) and it shows how my results can explain the numerical results in Roberts and Heesterbeek (2018). Note that when discussing the numerical results I only focus on Figures 1-3 because Figures 4-6 explore the effects of variation in the mortality rate of the focal host 1 (this study only focuses on the effects of variation in characteristics of alternative host species). Also, note that the simulations in Figures 1-3 assume no disease-induced mortality, no recovery, and susceptible and infected individuals have equal competitive abilities, which implies $\mathbf{B}_{ij} = 0 = \mathbf{C}_{ij} = 0$ for all $i, j$.

Reproduction number results: Equation (2.4) of Roberts and Heesterbeek (2018) defines the elasticity of $\mathcal{R}_0^C$ for a two-host model with DDDT or FDDT. The term in parentheses is a special case of sensitivity $d\mathcal{R}_0^C/dN_2^\ddagger$ in equation (S174) of Section S2.4. Similarly, the last equation of section 2.3 of Roberts and Heesterbeek (2018) defines the elasticity of $\mathcal{R}_0^C$ for a two-host model with ET where this is no uptake. The term in parentheses is also a special case of sensitivity $d\mathcal{R}_0^C/dN_2^\ddagger$ in equation (S174) of Section S2.4. In particular, the ET model in Roberts and Heesterbeek (2018) behaves like a DDDT model at equilibrium, which means the sensitivity of $\mathcal{R}_0^C$ is given by equation (S184) in Special Case A of Section S2.4; notice the similarity of the forms of that equation and the equation in Roberts and Heesterbeek (2018). Finally, the last equation of Section 3.3 in Roberts and Heesterbeek (2018) presents a formal definition of the elasticity of $\mathcal{R}_0^C$ for an $n$-host model. Equation (S174) of Section S2.4 provides an analytical way to calculate and interpret the sensitivity, which applies to models with ET, DDDT, FDDT, and transmission intermediate to DDDT and FDDT.

The sensitivity $d\mathcal{R}_0^C/dN_2^\ddagger$ in equation (S174) explains the patterns for elasticity of $\mathcal{R}_0^C$ in Figures 1-3. Those numerical results are for a two-host DDDT model (corresponding to $u_{j,S_i} = 0$ for all $i, j$ in the $n$-habitat model (S144)) where there is asymmetric interspecific competition (the Lotka-Volterra competition coefficients used in the simulations are $\alpha_{11} = \alpha_{22} = 0.15, \alpha_{12} = 0.1, \alpha_{21} = 0.075$). The general pattern across Figures 1-3 is that increased density of host 2 increases $\mathcal{R}_0^C$ (i.e., $[N_2^\ddagger/\mathcal{R}_0][d\mathcal{R}_0/dN_2^\ddagger] > 0$) unless host 2 has a much lower value of $\mathcal{R}_{0,i}$ than host 1 (Figure 1). For the model used to generate the figures, the sensitivity $d\mathcal{R}_0^C/dN_2^\ddagger$ in equation (S174) reduces to

$$\frac{d\mathcal{R}_0^C}{dN_2^\ddagger} = \mathcal{R}_0^C \left( \frac{y_2 x_2}{N_k^\ddagger} + \frac{y_1 x_1}{N_k^\ddagger} \frac{dN_1^\ddagger}{dN_2^\ddagger} \right). \tag{S340}$$

The sensitivity is negative only if $dN_1^\ddagger/dN_2^\ddagger$ is sufficiently large and negative and $x_1 y_1$ is sufficiently large relative to $x_2 y_2$. Biologically, this means that the sensitivity is negative only if interspecific competition is sufficiently strong and host 1 has sufficiently greater competence than host 2 (i.e., $\mathcal{R}_{0,1}$ sufficiently larger than $\mathcal{R}_{0,2}$). This matches the patterns in Figures 1-3 of Roberts and Heesterbeek (2018).

Infection prevalence and infected density results: The sensitivity formulas in Section S2.5.2 generalize the elasticity formulas in Section 3.3 of Roberts and Heesterbeek (2018). In addition, those formulas can explain the elasticities for $Y_1^*$ and $I_1^*$ in Figures 1-3 of Roberts and Heesterbeek (2018). Figures 1-3 assume two host species, DDDT, and asymmetric interspecific competition (the Lotka-Volterra competition coefficients used in the simulations are $\alpha_{11} = \alpha_{22} = 0.15, \alpha_{12} = 0.1, \alpha_{21} = 0.075$). The interspecific transmission parameters are defined by $\beta_{11} = \beta_{22} = \epsilon\sqrt{\beta_{11}\beta_{22}}$. Translated to the $n$-habitat model (S144), increases in $\epsilon$ lead to increases in $\chi_{12}$ and $\chi_{21}$.

The general pattern across Figures 1-3 is that the elasticity of host 1 infection prevalence to host 2 density is negative when there is no interspecific transmission and can be positive when interspecific transmission is sufficiently large. To see how these patterns are explained by the sensitivities in Section S2.5.2, I simplify the formula for the sensitivity $\partial Y_1^*/\partial\alpha$ in equation (S238) and divide by $\partial N_2^*/\partial\alpha$ to get

$$
\begin{aligned}
\frac{dY_1^*}{dN_2^*} = & \overbrace{\left(-\frac{\partial}{\partial \alpha}\frac{dN_2}{dt}\right)\frac{(-1)^{1+2^2}}{|\mathbf{J}|}\prod_{j\neq 2}(\beta_j X_n^*)\Big/\left(\frac{\partial N_2^*}{\partial \alpha}\right)}^{\text{positive}} \\
& \times \Bigg\{ \overbrace{\mathbf{A}_{11}\left(\mathbf{E}_N[1,2]\left[\mathbf{E}_Y[2,2]-\frac{\delta_2}{X_2^* Y_2^*}\right]-\mathbf{E}_N[2,2]\mathbf{E}_Y[1,2]\right)}^{\text{positive}} \\
& \underbrace{-\mathbf{A}_{12}\left(-\mathbf{E}_N[1,2]\mathbf{E}_Y[2,1]+\mathbf{E}_N[1,1]\left[\mathbf{E}_Y[2,2]-\frac{\delta_2 P_2^*}{X_2^* Y_2^*}\right]\right)\Bigg\}}_{\text{negative}}.
\end{aligned}
\tag{S341}
$$

The signs of terms indicated in the equation follow from $|\mathbf{J}| > 0$, $\mathbf{A}_{ij} < 0$, DDDT implying $\mathbf{E}_N$ and $\mathbf{E}_Y$ have positive entries, and Proposition S2.2 implying $\mathbf{E}_Y[i,i]+(U_i^*+\delta_i)/X_i^*Y_i^* = \mathbf{E}_Y[i,i]+\delta_i/X_i^*Y_i^* < 0$ for DDDT models. Increases in $\epsilon$ in the Roberts and Heesterbeek (2018) model correspond to increases in $\chi_{12}$ and $\chi_{21}$ in the $n$-habitat NY model (S147), which in turn causes $\mathbf{E}_N[1,2]$ and $\mathbf{E}_Y[1,2]$ to increase. When there is no interspecific transmission ($\epsilon = 0$, which implies $\mathbf{E}_N[i,j] = \mathbf{E}_Y[i,j] = 0$ for $i \neq j$), equation (S341) is negative because the first term in braces is zero. Because intraspecific competition is assumed to be greater than interspecific competition, the positive term in braces in equation (S341) increases faster than the negative term in braces as $\epsilon$ increases. Consequently, the elasticity $(N_2^*/Y_1^*)(dY_1^*/dN_2^*)$ in Figures 1-3 of Roberts and Heesterbeek (2018) is negative when there is no interspecific transmission and becomes more positive as interspecific transmission increases.

The pattern for the elasticity of host 1 infected density to host 2 density, $[N_2^*/I_1^*][\partial I_1^*/\partial N_2^*]$, in Figures 1-3 of Roberts and Heesterbeek (2018) is similar to the above: the elasticity is negative when there is no interspecific transmission and becomes less negative as interspecific transmission increases. To see how this pattern pattern is explained by the sensitivities in Section S2.5.2, I simplify the formula for the sensitivity $\partial N_1^*/\partial \alpha$ in equation (S243) and divide by $\partial N_2^*/\partial \alpha$ to get

$$
\begin{aligned}
\frac{dN_1^*}{dN_2^*} = & \overbrace{\left(-\frac{\partial}{\partial \alpha}\frac{dN_2}{dt}\right)\frac{-1}{|\mathbf{J}|}\prod_{j\neq 2}(\beta_j X_n^*)\Big/\left(\frac{\partial N_2^*}{\partial \alpha}\right)}^{\text{positive}} \\
& \times \mathbf{A}_{11}\left(\overbrace{\left[\mathbf{E}_Y[1,1]-\frac{\delta_1 P_1^*}{X_1^* Y_1^*}\right]\left[\mathbf{E}_Y[2,2]-\frac{\delta_2 P_2^*}{X_2^* Y_2^*}\right]}^{\text{positive}}\overbrace{-\mathbf{E}_Y[1,2]\mathbf{E}_Y[2,1]}^{\text{negative}}\right)
\end{aligned}
\tag{S342}
$$

Recall that the sensitivity of host 1 infected density is computed as $dI_1^*/dN_2^* = N_1^*(dY_1^*/dN_2^*) + Y_1^*(dN_1^*/dN_2^*)$. When there is no interspecific transmission ($\epsilon = 0$, which implies $\mathbf{E}_N[i,j] = \mathbf{E}_Y[i,j] = 0$ for $i \neq j$), equations (S341) and (S342) are negative, which explains the negative values of the elasticity of host 1 infected density in Figures 1-3. Both equations become more positive with increased interspecific transmission (i.e., larger $\epsilon$), which explains the slopes of the elasticity curves in Figures 1-3.

##### S3.1.12 Rudolf and Antonovics (2005) – $I_1^*/N_1^*$

Rudolf and Antonovics (2005) studied a two-host SI model with either DDDT or FDDT. The DDDT version of the model corresponds to the $n$-habitat model (S144) where $u_{j,S_i} = 0$ for all $i, j$ and $V \in \{S, I, R\}$. The FDDT version of the model corresponds to the $n$-habitat model (S144) where $\delta_i = 0$ and $u_{j,S_i} = 1$ for all $i, j$ and $V \in \{S, I, R\}$. In that study, "additive" density regulation corresponds to no interspecific competition and "substitutive" density regulation corresponds to strong, symmetric interspecific competition.

Richness-infection prevalence relationships for DDDT - no interspecific competition: Figure 2 of Rudolf and Antonovics (2005) predicts that introduction of a second host increases host 1 infection prevalence for additive density regulation (i.e., no interspecific competition). This pattern can be explained by equation (S238) of Section (S2.4.2). After dividing by $\partial N_2^*/\partial\alpha$, the sensitivity to the density of host 2 simplifies to

$$\frac{\partial Y_1^*}{\partial N_2^*} = \overbrace{\left(-\frac{\partial}{\partial\alpha}\frac{dN_2}{dt}\right)\frac{(-1)^{1+2^2}}{|\mathbf{J}|}\prod_{j\neq 2}(\beta_j X_n^*)\left(\frac{\partial N_2^*}{\partial\alpha}\right)^{-1}}^{\text{positive}}$$
$$\times\left\{\overbrace{\mathbf{A}_{11}\mathbf{E}_N[1,2]\left(-\frac{\delta_2 P_2^*}{X_2^* Y_2^*}\right)}^{\text{positive}}\underbrace{-\mathbf{A}_{12}\left(-\mathbf{E}_N[1,2]\mathbf{E}_Y[2,1] + \mathbf{E}_N[1,1]\left[\mathbf{E}_Y[2,2] - \frac{\delta_2 P_2^*}{X_2^* Y_2^*}\right]\right)\right\}}_{\text{negative}} \tag{S343}$$

where $\mathbf{E}_N[i,j] > 0$, $\mathbf{E}_Y[i,j] > 0$, $\mathbf{E}_Y[2,2] - \delta_2 P_2^*/X_2^* Y_2^* < 0$, and $\partial N_2^*/\partial\alpha < 0$. When interspecific competition is absence ($\mathbf{A}_{12} = 0$), the sensitivity $dY_1^*/dN_2^*$ is positive.

Richness-infection prevalence relationships for DDDT - yes interspecific competition: Figure 2 of Rudolf and Antonovics (2005) predicts that introduction of a second host decreases host 1 infection prevalence for substitutive density regulation (i.e., sufficiently strong interspecific competition). This prediction is explained by the previous equation. Specifically, the sensitivity can become negative when interspecific is sufficiently strong ($\mathbf{A}_{12}$ large in magnitude).

Richness-infection prevalence relationships for FDDT: Figure 2 predicts that introduction of a second host decreases host 1 infection prevalence under additive density regulation (i.e., no interspecific competition). For a two-host systems with FDDT and equal uptake rates, the sensitivity $\partial Y_1^*/\partial N_2^*$ simplifies to

$$\frac{dY_1^*}{dN_2^*} = \overbrace{\left(-\frac{\partial}{\partial\alpha}\frac{dN_2}{dt}\right)\frac{(-1)^{1+2^2}}{|\mathbf{J}|}\beta_1 X_1^* \beta_2 X_2^* \left(\frac{\partial N_2^*}{\partial\alpha}\right)^{-1}}^{\text{positive}} \tag{S344}$$

$$\times\left\{-\mathbf{A}_{11}\mathbf{E}_N[1,2]\left[\frac{U_2^* P_2^*}{X_2^* Y_2^*}\right] - \mathbf{A}_{12}\left[\mathbf{E}_N[1,1]\left(\mathbf{E}_Y[2,2] - \frac{U_2^* P_2^*}{X_2^* Y_2^*}\right) - \mathbf{E}_Y[2,1]\mathbf{E}_N[1,2]\right]\right\}. \tag{S345}$$

When interspecific transmission is low, $\mathbf{E}_N[i,j]$ and $\mathbf{E}_Y[i,j]$ are negative for $i \neq j$ and $\mathbf{E}_N[i,i]$ and $\mathbf{E}_Y[i,i]$ are positive for all $i$. Because $\partial N_2^*/\partial\alpha < 0$, this results in $dY_1^*/dN_2^*$

being negative for all levels of interspecific competition. This matches the prediction in Rudolf and Antonovics (2005).

However, $dY_1^*/dN_2^*$ can be positive under FDDT if interspecific transmission is sufficiently high. In particular, high interspecific transmission in the Rudolf and Antonovics (2005) model means $\mathbf{E}_Y[i,j]$ and $\mathbf{E}_N[i,j]$ are positive for $i \neq j$. If those positive values are large enough, then the term multiplied by $\mathbf{A}_{11}$ in equation (S345) will change signs and $dY_1^*/dN_2^*$ will be positive. As a result, introduction of host 2 causes an increase in host 1 infection prevalence. This possibility was missed in Rudolf and Antonovics (2005) likely because of the parameter values they considered. For example, if the parameter values in Figure 1a of Rudolf and Antonovics (2005) are altered slightly to $r_1 = 0.017$, $\beta_{12} = 0.4$, and $\beta_{21} = 0.1$, then host 2 increases host 1 infection prevalence.

#### S3.2 Environmental Transmission Models

##### S3.2.1 Cáceres et al. (2014) – $I_1^*/N_1^*$, $I_1^*$

Cáceres et al. (2014) analyzed a resource explicit two-host model where the second host competed for resources, could take up infection propagules, but could not be infected by the pathogen. This means $\chi_{12} = \chi_{22} = 0$, $\beta_2 = 0$, and there is no $Y_2$ variable. Consequently, matrix $\mathbf{B}$ of the Jacobian does not have a second column and matrices $\mathbf{E}_N$ and $\mathbf{E}_Y$ do not have second rows; the other matrices are adjusted accordingly. The model in Cáceres et al. (2014) assumed resource dependent shedding rates and uptake rates. Here, I will assume those parameters are fixed and show that the patterns observed in Cáceres et al. (2014) are consistent the predictions from the sensitivities.

Richness-infection prevalence relationships: The bottom right panels of Figures 3 and 4 in Cáceres et al. (2014) show that the second host always reduced host 1 infection prevalence. This outcome is consistent with equation (S238) of Section (S2.4.2). That equation reduces to

$$\frac{dY_1^*}{d\alpha} = \left(-\frac{\partial}{\partial\alpha}\frac{dN_2}{dt}\right)\frac{(-1)^{1+2^2}}{|\mathbf{J}|}\beta_1 X_1 \left|\begin{bmatrix} \mathbf{A}_{11} & \mathbf{A}_{12} & \mathbf{B}_{11} \\ \mathbf{A}_{21} & \mathbf{A}_{22} & 0 \\ \mathbf{E}_N[1,1] & \mathbf{E}_N[1,2] & \chi_1 N_1^* - (U^*+\delta)/X_1^* Y_1^* \end{bmatrix}^{[\not{2},\not{3}]}\right| \tag{S346}$$

$$= \left(-\frac{\partial}{\partial\alpha}\frac{dN_2}{dt}\right)\frac{1}{|\mathbf{J}|}\beta_1 X_1 \left(\mathbf{A}_{11}\mathbf{E}_N[1,2] - \mathbf{A}_{12}\mathbf{E}_N[1,1]\right) > 0 \tag{S347}$$

where $|\mathbf{J}| > 0$; $\mathbf{E}_N[1,2] < 0$ because host 2 can take up infectious propagules but not shed them; and $\mathbf{E}_N[1,1] > 0$ because it is a necessary condition for the equilibrium to exist. Dividing by $\partial N_2^*/\partial K_2 < 0$ yields $dY_1^*/dN_2^* < 0$. Thus, increased density of host 2 always reduces host 1 infection prevalence. I note that the two terms in the parentheses show that the reduction in host 1 infection prevalence occurs via resource competition ($\mathbf{A}_{12}\mathbf{E}_N[1,1]$) and uptake of infectious propagules ($\mathbf{A}_{11}\mathbf{E}_N[1,2]$).

Richness-infected densities relationships: The top right panels of Figures 3 and 4 in Cáceres et al. (2014) show that the second host always reduced host 1 infected density. This outcome is also consistent with the predictions from the sensitivities. First, I compute the sensitivity of host 1 total density as

$$\frac{dN_1^*}{d\alpha} = \left(-\frac{\partial}{\partial\alpha}\frac{dN_2}{dt}\right)\frac{(-1)^{1+2^2}}{|\mathbf{J}|}\beta_1 X_1 \left| \begin{bmatrix} \mathbf{A}_{11} & \mathbf{A}_{12} & \mathbf{B}_{11} \\ \mathbf{A}_{21} & \mathbf{A}_{22} & 0 \\ \mathbf{E}_N[1,1] & \mathbf{E}_N[1,2] & \chi_1 N_1^* - (U^*+\delta)/X_1^* Y_1^* \end{bmatrix}^{[2,1]} \right| \tag{S348}$$

$$= \overbrace{\left(-\frac{\partial}{\partial\alpha}\frac{dN_2}{dt}\right)\frac{1}{|\mathbf{J}|}\beta_1 X_1}^{\text{positive}} \left( \overbrace{-\mathbf{B}_{11}\mathbf{E}_N[1,2]}^{\text{negative}} + \overbrace{\mathbf{A}_{12}\left[\chi_1 N_1^* - \frac{U^*+\delta}{X_1^* Y_1^*}\right]}^{\text{positive}} \right) \tag{S349}$$

In most cases, this equation will be positive because the negative term in parentheses is smaller in magnitude than positive term. Dividing by $\partial N_2^*/\partial K_2 < 0$ yields $dN_1^*/dN_2^* < 0$. Using equation (S232) to combine the two previous equations, yields $dI_1^*/dN_2^* < 0$, which is consistent with the observed decrease in host 1 infected density when host 2 is present. In principle, host 2 can increase host 1 infected density if the uptake rate of host 2 is very high ($\mathbf{E}_N[1,2]$ large in magnitude) or the mortality rate from the disease is very high ($\mathbf{B}_{11}$ large in magnitude); this causes the negative term in parentheses to be larger in magnitude than the positive term. This was not observed because those conditions are not satisfied by the parameter values used in Cáceres et al. (2014).

##### S3.2.2 Espira et al. (2022) – $\mathcal{R}_0^C$

Espira et al. (2022) analyzed a two-habitat environmental transmission model, which when written in my notation has the form

$$\begin{aligned} \frac{dS_i}{dt} &= \beta_i S_i (P_i + \theta_i P_j) \\ \frac{dI_i}{dt} &= \beta_i S_i (P_i + \theta_i P_j) - \nu_i I_i \\ \frac{dR_i}{dt} &= \nu_i I_i \\ \frac{dP_i}{dt} &= \chi_i I_i - \left(\frac{u_i}{\epsilon_i} N_i + \theta_j \frac{u_j}{\epsilon_i} N_j\right) P_i - \delta P_i \end{aligned} \tag{S350}$$

where $\epsilon_i$ is the size of the habitat for species $i$, and $\theta_i$ is the fraction of habitat of species $i$ that overlaps with the habitat of species $j$. The uptake rate for species $i$, $u_i$, is assumed to be the same in all habitats and independent of infection status (i.e., $u_{j,S_i} = u_{j,I_i} = u_{j,R_i} = u_i$). In the original study, the link between the transmission and uptake parameters was explicitly denoted by $\beta_i = \pi_i u_i = \pi_i \kappa_i \rho_i$. The analysis in Espira et al. (2022) focused on the pathogen pick-up efficiency for each host,

$$\eta_i = \frac{u_i N_i}{u_i N_i + \delta \epsilon_i} \tag{S351}$$

and the ratio of the basic reproduction numbers for the two host species, $\mathcal{R}_{0,2}/\mathcal{R}_{0,1}$, where the basic reproduction number for each host is

$$\mathcal{R}_{0,i} = \frac{\beta_i \chi_i}{\nu_i} \frac{\epsilon_i}{u_i + \delta\epsilon_i} = \frac{\pi_i \chi_i \epsilon_i}{\nu_i} \frac{u_i N_i}{u_i N_i + \delta\epsilon_i} = \frac{\pi_i \chi_i \epsilon_i}{\nu_i} \eta_i. \tag{S352}$$

The non-zero submatrix of the next generation matrix that is used to compute the community reproduction number is

$$\mathbf{M} = \begin{pmatrix} \frac{\beta_1\chi_1 N_1\epsilon_1}{\nu_1(u_1N_1+\theta_2u_2N_2+\delta\epsilon_1)} & \frac{\beta_1\chi_2N_1\epsilon_1\epsilon_2\theta_1}{\nu_2(u_2N_2+\theta_1u_1N_1+\delta\epsilon_2)} \\ \frac{\beta_2\chi_1N_2\epsilon_1\epsilon_2\theta_2}{\nu_1(u_1N_1+\theta_2u_2N_2+\delta\epsilon_1)} & \frac{\beta_2\chi_2N_2\epsilon_2}{\nu_2(u_2N_2+\theta_1u_1N_1+\delta\epsilon_2)} \end{pmatrix} \tag{S353}$$

$$= \begin{pmatrix} \mathcal{R}_{0,1}\frac{u_1N_1+\epsilon_1}{u_1}\frac{\epsilon_1}{u_1N_1+\theta_2u_2N_2+\delta\epsilon_1} & \mathcal{R}_{0,2}\frac{\beta_1}{\beta_2}\frac{u_2N_2+\epsilon_2}{u_2N_2}\frac{N_1\epsilon_1\epsilon_2\theta_1}{u_2N_2+\theta_1u_1N_1+\delta\epsilon_2} \\ \mathcal{R}_{0,1}\frac{\beta_2}{\beta_1}\frac{u_1N_1+\epsilon_1}{u_1N_1}\frac{N_2\epsilon_1\epsilon_2\theta_2}{u_1N_1+\theta_2u_2N_2+\delta\epsilon_1} & \mathcal{R}_{0,2}\frac{u_2N_2+\epsilon_2}{u_2}\frac{\epsilon_2}{u_2N_2+\theta_1u_1N_1+\delta\epsilon_2} \end{pmatrix} \tag{S354}$$

Variation of epidemiological parameters: The numerical results in Figures 3, 4, S1, and S2 of Espira et al. (2022) assumed fixed degradation rates ($\mu$) and habitat sizes ($\epsilon_1, \epsilon_2$). The figures show that when $\mathcal{R}_{0,2}$ and $\eta_2$ are varied independently, the community reproduction number, $\mathcal{R}_0^C$, increases with increased $\mathcal{R}_{0,2}$ and decreased $\eta_2$. Because $\mathcal{R}_{0,2}$ depends on $\eta_2$ (see equation (S352)), independent variation of $\mathcal{R}_{0,2}$ and $\eta_2$ effectively means that variation in $\mathcal{R}_{0,2}$ is due to variation in the ratio $\pi_2\chi_2/\nu_2$ and variation in $\eta_2$ is compensated by variation in $\pi_2\chi_2/\nu_2$ such that $\mathcal{R}_{0,2}$ remains constant.

The sensitivity-based theory in this study explain why $\mathcal{R}_0^C$ increases with increased $\mathcal{R}_{0,2}$ and decreased $\eta_2$. First, consider increases in $\mathcal{R}_{0,2}$ due to increases in the ratio $\pi_2\chi_2/\nu_2$. This corresponds to increases in $\beta_2$, increases in $\chi_2$, or decreases in $\nu_2$ in model (S350). Increases in $\mathcal{R}_0^C$ are guaranteed because the entries of $\mathbf{M}$ satisfy $\partial\mathbf{M}_{ij}/\partial\beta_2 \geq 0$, $\partial\mathbf{M}_{ij}/\partial\chi_2 \geq 0$, and $\partial\mathbf{M}_{ij}/\partial\nu_2 \leq 0$. Second, consider decreases in $\eta_2$ that are compensated by increases in $\pi_2\chi_2/\nu_2$ such that $\mathcal{R}_{0,2}$ remains constant. Using the form of $\mathbf{M}$ in equation (S354), this corresponds to decreasing $u_2$ or increasing $\delta$. Increases in $\mathcal{R}_0^C$ are guaranteed with those changes in parameters because the entries of $\mathbf{M}$ satisfy $\partial\mathbf{M}_{ij}/\partial u_2 \leq 0$ and $\partial\mathbf{M}_{ij}/\partial\delta \geq 0$.

##### S3.2.3 Fenton et al. (2015) – $\mathcal{R}_0^C$

Fenton et al. (2015) analyzed an environmental transmission model with an alternative form. The model describes the changes in the densities of susceptible and infected individuals in each host population ($S_i$, $I_i$) and the densities of infectious propagules released by host $i$ ($P_i$). Rewriting the model using the notation from this study yields,

$$
\begin{aligned}
\frac{dS_i}{dt} &= \underbrace{F_i(\cdot)}_{\text{reproduction}} - \underbrace{\sum_{j=1}^{n} \beta_{ij} S_i P_j}_{\text{infection}} - \overbrace{m_i S_i}^{\text{mortality}} + \overbrace{\gamma_i R_i}^{\text{loss of immunity}} \\
\frac{dI_i}{dt} &= \underbrace{\sum_{j=1}^{n} \beta_{ij} S_i P_j}_{\text{infection}} - \underbrace{(\mu_i + m_i) I_i}_{\text{mortality}} - \underbrace{\nu_i I_i}_{\text{recovery}} \\
\frac{dR_i}{dt} &= \underbrace{\nu_i I_i}_{\text{recovery}} - \underbrace{(\xi_i + m_i) I_i}_{\text{mortality}} - \underbrace{\gamma_i I_i}_{\text{loss of immunity}} \\
\frac{dP_i}{dt} &= \underbrace{\chi_i I_i}_{\text{propagule shedding}} - \underbrace{\delta_i P_i}_{\text{degradation}}
\end{aligned}
\tag{S355}
$$

All parameters are interpreted in a similar way except that $\beta_{ij}$ is the transmission coefficient for host $i$ encountering infectious propagules shed by host $j$ and $\chi_i$ is the shedding rate for host $i$. Note that the model assumes no interspecific host competition.

Model (S355) reduces to a density-dependent direct transmission model when there is a separation of time scales between the dynamics of the host classes and the infectious propagules. This is because the transmission rates become $\hat{\beta}_{ij} S_i I_j = \chi_i \beta_{ij} S_i I_j / \delta_i$, which is identical to the transmission form in model (S162). Consequently, all of the results in this study for DDDT models apply to model (S355) and can explain the results in Fenton et al. (2015).

Variation in habitat overlap: Fenton et al. (2015) found that $\mathcal{R}_0^C$ always increased with increased habitat overlap (parameter $\omega$ in their model); see the analysis of equation (4) of that study. Increased $\mathcal{R}_0^C$ with increased habitat overlap is explained in the following way. Increased habitat overlap corresponds to increased $\beta_{ij}$ ($i \neq j$) in model (S355), which in turn corresponds to increased $\chi_{ij}$ in the $n$-habitat SIR model (S144). The sensitivity results in Section S2.4.4 show that $\partial \mathcal{R}_0 / \partial \chi_{ij} > 0$.

Richness-$\mathcal{R}_0^C$ relationships: Fenton et al. (2015) found that $\mathcal{R}_0^C$ always increased with increased host species richness. Increased $\mathcal{R}_0^C$ with increased host species richness is explained by the positive sign of $d\mathcal{R}_0 / dN_i^{\ddagger}$ in equation (S184) of Special Case 1 in Section S2.4.2 when interspecific competition is absent. I note that the positive sign of equation (S184) also explains why it is possible for the pathogen to invade the two-host community in Figures 1 and 2 of Fenton et al. (2015) when $\mathcal{R}_{0,i} < 1$ for both host species.

##### S3.2.4 Rudge et al. (2013) – $\mathcal{R}_0^C$, $I_1^*/N_1^*$

Rudge et al. (2013) also analyzed the environmental transmission model (S355). As noted above, model (S355) assumes no interspecific competition and it reduces to a density-dependent direct transmission model when there is a separation of time scales between the dynamics of the host classes and the infectious propagules. Consequently, all of the results in this study for DDDT models without interspecific competition apply to model (S355) and can explain the results in Rudge et al. (2013).

Richness-$\mathcal{R}_0^C$ relationships: Tables 2 of Rudge et al. (2013) shows that removal of a host species never increases $\mathcal{R}_0^C$ (each entry in the bottom row of Table 1 is greater than or equal to all entries in the corresponding column in Table 2). Table 1 of of Rudge et al. (2013) shows that the estimated value of $\mathcal{R}_0^C$ is always larger than each individual $\mathcal{R}_{0,i}$ (bottom row of Table 1 larger than all other rows). Figure 3 of Rudge et al. (2013) shows that $\mathcal{R}_0^C$ decreases with the reduction of rodent density. These three patterns are explained by equation (S184) in Special Case A of section S2.4.2. In particular, that equation is always positive when there is DDDT and no interspecific competition. Hence, host additions always increase $\mathcal{R}_0^C$ and removals or reduced density always decrease $\mathcal{R}_0^C$.

Infection prevalence results: Figure 2 of Rudge et al. (2013) shows that vaccination of bovines and removal of bovines cause reductions in prevalence of all host species. The sensitivities of host infection prevalence for the $n$-habitat model are too complex to analyze when interspecific transmission is not small relatively to intraspecific transmission. However, the patterns in Figure 2 of Rudge et al. (2013) match the predictions from (i) the sensitivities $\partial Y_1^*/\partial\alpha$ and $dY_1^*/dN_n^*$ computed for the $n$-habitat model when intraspecific transmission is much less than interspecific transmission and (ii) the sensitivities $\partial Y_1^*/\partial\alpha$ and $dY_1^*/dN_n^*$ computed for the 1-habitat model. Thus, the sensitivity results in this study help explain (but may not completely explain) the numerical results in Rudge et al. (2013).

First, Figure 2 of Rudge et al. (2013) shows that infection prevalence decreases with increased vaccination. Vaccination in the Rudge et al. (2013) model is mathematically equivalent to reducing the reproduction rate of bovines. This means that the effects of increased vaccination are predicted by the sign of $\partial Y_1^*/\partial\alpha$. The negative slope in Figure 2 matches the prediction from Special Case 2 of Section S2.5.2 when there is no interspecific competition, DDDT, and low interspecific transmission; see equation (S254) and note that it has a negative sign. The negative slope also matches the prediction from the 1-habitat model. Specifically, under the assumptions of no interspecific competition and DDDT, the formula for $\partial Y_1^*/\partial\alpha$ in equation (S73) of Section S1.5.2 reduces to $\partial Y_1^*/\partial\alpha = -Q'\varphi_n > 0$ where $Q'$ is a positive constant and $\varphi_n > 0$. Thus, increased vaccination is predicted to decrease infection prevalence.

Second, Figure 2 of Rudge et al. (2013) shows that infection prevalence decreases with decreased bovine density. The effect of reduced density is predicted by the sign of $\partial Y_1^*/\partial N_n^* = (\partial Y_1^*/\partial\alpha)/(\partial N_n^*/\partial\alpha)$, where $\partial Y_1^*/\partial N_n^*$ is negative. Using arguments from the previous paragraph, the negative slope in Figure 2 matches the predictions from Special Case 2 of Section S2.5.2 and the 1-habitat model.

##### S3.2.5 Searle et al. (2016) – $\mathcal{R}_0^C$, $I_1^*/N_1^*$, $I_1^*$

Searle et al. (2016) analyzed a 1-habitat environmental transmission model with two-host species, *Daphnia dentifera* and *D. lumholtzi*. After converting their model into the form of model (2) and equation (3) from the main text, the model parameters are defined as in Table S4. Note that (a) the non-disease mortality rate is zero because it has been absorbed into the reproduction rates and disease induced mortality rates and (b) there are no recovered classes because recovery from infection is not possible. Comparing the quantities in the bottom half of the Table S4 reveals that *D. lumholtzi* has higher competence than *D. dentifera*

($\mathcal{R}_{0,L}(\infty) > \mathcal{R}_{0,D}(\infty)$), both hosts are sources but *D. lumholtzi* is a larger source than *D. dentifera* ($\varphi_L$ and $\varphi_D$ are positive for biologically reasonable spore densities and $\varphi_L > \varphi_D$), and the pathogen basic reproduction number is larger for *D. lumholtzi* than *D. dentifera* (larger $\mathcal{R}_{0,L} > \mathcal{R}_{0,D}$).

**Table S4**: Parameters from (Searle et al. 2016) model

| Host | $r_i$ | $b_i$ | $\alpha_{ii}$ | $\alpha_{ij}$ | $\beta_i$ | $m_i$ | $\mu_n$ | $\chi_i$ | $u_{S_i}$ | $u_{I_i}$ |
|---|---|---|---|---|---|---|---|---|---|---|
| *D. dentifera* | 0.206 | 0.75 | 1/97.5 | 2.63/97.5 | $1.45{\cdot}10^{-5}$ | 0 | 0.05 | 6000 | 0.0348 | 0.0186 |
| *D. lumholzti* | 0.246 | 0.75 | 1/12.8 | -0.286/12.8 | $4.87{\cdot}10^{-5}$ | 0 | 0.05 | 6200 | 0.0361 | 0.0171 |

| | $\mathcal{R}_{0,i}(\infty)$ | $\mathcal{R}_{0,i}$ | $\varphi_i$ | $N_i^\ddagger$ |
|---|---|---|---|---|
| *D. dentifera* | 1.74 | 1.52 | $2552 - 0.0186P^*$ | 36.4 |
| *D. lumholtzi* | 6.04 | 2.90 | $5173 - 0.0171P^*$ | 23.2 |

The degradation rate is $\delta = 0.5$. See Searle et al. (2016) for units.

Richness-$\mathcal{R}_0^C$ relationships: $\mathcal{R}_0^C$ for the two-host disease-free equilibrium is 2.79. This means that adding *D. dentifera* reduces $\mathcal{R}_0^C$ whereas adding *D. lumholtzi* increases $\mathcal{R}_0^C$. This pattern agrees with sensitivities of $\mathcal{R}_0^C$, which are

$$\frac{\partial \mathcal{R}_0^C}{N_L^\ddagger} = \frac{u_{S_L}}{U^\ddagger + \delta}(\mathcal{R}_{0,L}(\infty) - \mathcal{R}_0^C) + \frac{u_{S_D}}{U^\ddagger + \delta}(\mathcal{R}_{0,D}(\infty) - \mathcal{R}_0^C)(-\alpha_{DL}) \tag{S356}$$

$$\frac{\partial \mathcal{R}_0^C}{N_D^\ddagger} = \frac{u_{S_D}}{U^\ddagger + \delta}(\mathcal{R}_{0,D}(\infty) - \mathcal{R}_0^C) + \frac{u_{S_L}}{U^\ddagger + \delta}(\mathcal{R}_{0,L}(\infty) - \mathcal{R}_0^C)(-\alpha_{LD}) \tag{S357}$$

In the first equation, both terms are positive because (first term) increased *D. lumholtzi* density increases the density of a high competence host and (second term) decreased density of *D. dentifera* reduces the density of a low competence host. Thus, increased density of *D. lumholtzi* increases $\mathcal{R}_0^C$. In the second equation,the first term is negative because increased *D. dentifera* density increases the density of a low competence host whereas the second term is positive because increased density of *D. lumholtzi* increases the density of a high competence host. The first term is larger in magnitude than the second because the coefficient $\alpha_{LD}$ is small in magnitude. Thus, increased density of *D. dentifera* decreases $\mathcal{R}_0^C$.

Richness-infection prevalence relationships: The infection prevalences of both species at the two-host endemic equilibrium are higher than the infection prevalences at the single-host endemic equilibria. Thus each host increases infection prevalence in the other. The following explains how these responses can be computed from the Jacobian.

When evaluated at the coexistence equilibrium, the Jacobian for the system is

$$\mathbf{J} = \begin{pmatrix} -0.0585 & -0.154 & -1.83 & 0 & 0 \\ 0.0839 & -0.293 & 0 & -1.20 & 0 \\ 0.000783 & 0.00206 & -0.08281 & 0 & 2.96 \cdot 10^{-7} \\ -0.00321 & 0.0112 & 0 & -0.172 & 5.09 \cdot 10^{-7} \\ 7.87 & 2420 & 229000 & 146000 & -1.80 \end{pmatrix} \tag{S358}$$

where the order of the columns is $N_D, N_L, Y_D, Y_L, P$. Note that the values of the matrix are consistent with our calculated values from above. In particular, both hosts are sources ($J_{51}$ and $J_{52}$ positive); *D. lumholtzi* is the larger source ($J_{52} > J_{51}$); *D. dentifera* is more strongly regulated by interspecific competition than intraspecific competition ($J_{11}$ smaller in magnitude than $J_{12}$); and *D. lumholtzi* is more strongly regulated by intraspecific competition than interspecific competition ($J_{22}$ larger in magnitude than $J_{21}$).

The effect of increased density of *D. lumholtzi* on infection prevalence in *D. dentifera* is explained by equation (S74) in Section S1.5.2. In particular, the term $Q(-1)^2|\mathbf{A}^{[\not{2},\not{2}]}|\varphi_2 Y_2^*/|\mathbf{J}|$ in equation (S74) is positive because host 2 is a source; the term $Q(-1)^1|\mathbf{A}^{[\not{2},\not{1}]}|\varphi_1 Y_1^*/|\mathbf{J}|$ is negative because host 1 is a source and there is interspecific competition; and the sum of the two terms is positive because host 2 is a much larger source than host 1 ($\varphi_2$ much larger in magnitude than $\varphi_1$).

The effect of increased density of *D. dentifera* on infection prevalence in *D. lumholtzi* is explained by the same equations. Specifically, the term $Q(-1)^2|\mathbf{A}^{[\not{1},\not{1}]}|\varphi_1 Y_2^*/|\mathbf{J}|$ in equation (S74) is positive because host 1 is a source; the term $Q(-1)^1|\mathbf{A}^{[\not{1},\not{2}]}|\varphi_1 Y_1^*/|\mathbf{J}|$ is positive because host 2 is a source and host 1 has positive effects on host 2 ($\mathbf{A}_{21} > 0$); and the sum of the two terms is positive. Note that the response would be the same if host 1 was a weak interspecific competitor ($\mathbf{A}_{21}$ negative and sufficiently small in magnitude), because the term $Q(-1)^1|\mathbf{A}^{[\not{1},\not{2}]}|\varphi_1 Y_1^*/|\mathbf{J}|$ would be negative, but smaller in magnitude than the term $Q(-1)^2|\mathbf{A}^{[\not{1},\not{1}]}|\varphi_1 Y_2^*/|\mathbf{J}|$ in equation (S74).

Richness-infected density relationships: Infected density of *D. dentifera* decreases when *D. lumholtzi* is added to the system; this means the responses in infection prevalence and infected density are in the opposite direction. In comparison, infected density of *D. lumholtzi* increases when *D. dentifera* is added to the system; this means the responses in infection prevalence and infected density are in the same direction. The following explains how these responses can be computed from the Jacobian.

The effect of increased density of *D. lumholtzi* on infected density in *D. dentifera* is explained by equations (S74), (S81), and (S82) in Section S1.5.2. In particular, compare the terms in the first columns of the matrices in equation (S82): $S_1^* \vec{E}_N[1] \approx 143$ is positive because host 1 a source and $\theta_n \approx 72700$ is very large because host 2 is a much larger source than host 1. Larger differences in the magnitudes between $S_1^* \vec{E}_N[1]$ and $\theta_n$ result in the second determinant in equation (S82) being the opposite sign and larger in magnitude than the first determinant. As a result, while the sensitivity $dY_1^*/dN_2^*$ is positive, the sensitivity $dI_1^*/dN_2^*$ is negative because introduction of host 2 causes very large reductions in the density of host 1 ($dN_1^*/dN_2^*$ large and negative).

The effect of increased density of *D. dentifera* on infected density in *D. lumholtzi* is explained by the same equations. In particular, compare the terms in the first columns of the matrices in equation (S82): $S_2^* \mathbf{A}_{22} = 1.58$ is much larger in magnitude than $\vec{\omega}_n[1] = 0.06$ and the magnitude of $S_2^* \vec{E}_N[2] = 13000$ is similar to that of $\theta_n = 72700$. This results in the second determinant in equation (S82) being the smaller in magnitude than the first determinant. Said another way, the magnitude of $dN_1^*/dN_2^*$ is smaller than $dY_1^*/dN_2^*$. As a result, the sensitivities $dY_1^*/dN_2^*$ and $dI_1^*/dN_2^*$ have the same signs and host 1 increases both the infection prevalence and infected density in host 2.

### S4 Parameter values for figures

The figures were generated using a 4-host version of model (2a-d) from the main text, with host reproduction rates defined by the Lokta-Volterra equations (3). The values of $\mathcal{R}_0^C$ were computed by solving for the unique disease-free equilibrium and evaluating the equation for $\mathcal{R}_0^C$ in Box 2. The values of focal host infection prevalence and infected density were computed by solving for the unique endemic equilibrium. All calculations were done in Maple v2019.2 (Maple Version 2019.2).

For each row of Figures 1, 2, 3, and 5, curves of the same color were generated using the same parameter values. Black circles correspond to cases where the three metrics have sensitivities of the same sign, blue squares correspond to cases where the sign of the sensitivity for $\mathcal{R}_0^C$ is the opposite of the sign of the sensitivities for focal host infection prevalence and infected density (via mechanisms 1 or 2), and red triangles correspond to cases where focal host infection prevalence and infected density have sensitivities of opposite signs (because of mechanism 3). The only exception is the red triangles in the bottom row of Figure 4, where the sign of the sensitivity for $\mathcal{R}_0^C$ is the opposite of the signs for the other two metrics.

Bold entries in each table denote the parameter values that differ between curves in the same panels; this highlights the specific parameter values causing the differences between simulations.

**Table S4**: Parameter values for Figure 1

| **Parameter** | Panels A-C (top row) | | | Panels D-F (bottom row) | | |
|---|---|---|---|---|---|---|
| | Black | Blue | Red | Black | Blue | Red |
| $r_i$ | 1 | 1 | 1 | 1 | 1 | 1 |
| $b_i$ | 1 | 1 | 1 | 1 | 1 | 1 |
| $\alpha_{ii}$ | 1 | 1 | 1 | 1 | 1 | 1 |
| $\alpha_{ij}$ | **0.1** | **0.75** | **0.1** | **0.1** | **0.1** | **0.6** |
| $e_{ij}$ | 1 | 1 | 1 | 1 | 1 | 1 |
| $m_i$ | 0 | 0 | 0 | 0 | 0 | 0 |
| $(\beta_1, \beta_2, \beta_3)$ | (0.5,0.5,0.5) | (0.5,0.5,0.5) | (0.5, 0.5, 0.5) | (1,1,1) | (1,1,1) | (1,1,1) |
| $\beta_4$ | 0 - 2 | 0 - 2 | 0 - 2 | 1 | 1 | 1 |
| $(\mu_1, \mu_2, \mu_3)$ | (**0.5**,1,1) | (**0.5**,1,1) | (**3**, 1,1) | (0.1,0.1,0.1) | (0.1,0.1,0.1) | (0.1,0.1,0.1) |
| $\mu_4$ | 0.5 | 0.5 | 0.5 | 0 - 2 | 0 - 6 | 0 - 2 |
| $\nu_i$ | 1 | 1 | 1 | 1 | 1 | 1 |
| $(\xi_1, \xi_2, \xi_3, \xi_4)$ | 0.1 | 0.1 | 0.1 | 0 | 0 | 0 |
| $(\gamma_1, \gamma_2, \gamma_3, \gamma_4)$ | 1 | 1 | 1 | 1 | 1 | 1 |
| $(\chi_1, \chi_2, \chi_3, \chi_4)$ | (5,5,5,25) | (5,5,5,25) | (5,5,5,25) | (1,10,10,**9**) | (1,10,10,**1**) | (1,10,10,**9**) |
| $u_{S_i} = u_{I_i} = u_{R_i}$ | 1 | 1 | 1 | 2 | 2 | 2 |
| $\delta$ | 1 | 1 | 1 | 1 | 1 | 1 |

**Table S5**: Parameter values for Figure 2

| **Parameter** | Panels A-C (top row) | | Panels D-F (bottom row) | |
|---|---|---|---|---|
| | Black | Blue | Black | Blue |
| $r_i$ | 1.1 | 1.1 | 1.1 | 1.1 |
| $b_i$ | 0.2 | 0.2 | 0.2 | 0.2 |
| $(\alpha_{11}, \alpha_{22}, \alpha_{33})$ | (1,1,1) | (1,1,1) | (1,1,1) | (1,1,1) |
| $\alpha_{44}$ | 1-9 | 1-9 | 0.1-10.1 | 0.1-10.1 |
| $\alpha_{ij}$ | **0** | * | **0** | ‡ |
| $e_{ij}$ | 1 | 1 | 1 | 1 |
| $m_i$ | 0.1 | 0.1 | 0.1 | 0.1 |
| $(\beta_1, \beta_2, \beta_3, \beta_4)$ | (2,2,3,1) | (2,2,3,1) | (1,2,3,2) | (1,2,3,2) |
| $(\mu_1, \mu_2, \mu_3, \mu_4)$ | 0.25 | 0.25 | 0.25 | 0.25 |
| $\nu_i$ | 0.1 | 0.1 | 0.1 | 0.1 |
| $xi_i$ | 0.1 | 0.1 | 0.1 | 0.1 |
| $\gamma_i$ | 1 | 1 | 1 | 1 |
| $(\chi_1, \chi_2, \chi_3, \chi_4)$ | (5,10,10,20) | (5,10,10,20) | (20,1,20,1) | (20,1,20,1) |
| $u_{S_i} = u_{I_i} = u_{R_i}$ | 20 | 20 | 20 | 20 |
| $\delta$ | 30 | 30 | 20 | 20 |

$*$ Each competition coefficient, $\alpha_{ij}$, is determined by the $i, j$ entry of the matrix $\begin{pmatrix} 1 & 0.1 & 0.8 & 0.9 \\ 0.9 & 1 & 0.6 & 0.9 \\ 0.1 & 0.5 & 1 & 0.9 \\ 0.4 & 0.7 & 0.7 & - \end{pmatrix}$.

‡ Each competition coefficient, $\alpha_{ij}$, is determined by the $i, j$ entry of the matrix $\begin{pmatrix} 1 & 0.1 & 0.7 & 0.8 \\ 0.7 & 1 & 0.6 & 0.9 \\ 0.9 & 0.9 & 1 & 0.1 \\ 0.1 & 0.5 & 0.9 & - \end{pmatrix}$.

**Table S6**: Parameter values for Figure 3

| **Parameter** | Panels A-C (top row) | | | Panels D-F (bottom row) | | |
|---|---|---|---|---|---|---|
| | Black | Blue | Red | Black | Blue | Red |
| $r_i$ | 0.5 | 0.5 | 0.5 | 1 | 1 | 1 |
| $b_i$ | 1 | 1 | 1 | 1 | 1 | 1 |
| $(\alpha_{11}, \alpha_{22}, \alpha_{33})$ | (1,1,1) | (1,1,1) | (1,1,1) | (1,1,1) | (1,1,1) | (1,1,1) |
| $\alpha_{44}$ | 0.1-10 | 0.1-10 | 0.1-10 | 0.1-10 | 0.1-10 | 0.1-10 |
| $\alpha_{ij}$ | 0 | 0 | 0 | **0** | **0** | **0.2** |
| $e_{ij}$ | 1 | 1 | 1 | 1 | 1 | 1 |
| $m_i$ | 0 | 0 | 0 | 0 | 0 | 0 |
| $(\beta_1, \beta_2, \beta_3, \beta_4)$ | (1,1,1,**1**) | (1,1,1,**6**) | (1,1,1,**1**) | (1,1,1,0.3) | (1,1,1,0.3) | (1,1,1,0.3) |
| $(\mu_1, \mu_2, \mu_3, \mu_4)$ | (**0.5**,0.5,0.5,0.5) | (**0.5**,0.5,0.5,0.5) | (**2**,0.5,0.5,0.5) | (0.5,0.5,0.5,0.5) | (0.5,0.5,0.5,0.5) | (0.5,0.5,0.5,0.5) |
| $\nu_i$ | 1 | 1 | 1 | 1 | 1 | 1 |
| $xi_i$ | 0 | 0 | 0 | 0.1 | 0.1 | 0.1 |
| $\gamma_i$ | 1 | 1 | 1 | 1 | 1 | 1 |
| $(\chi_1, \chi_2, \chi_3, \chi_4)$ | (10,10,10,**30**) | (10,10,10,**3**) | (10,10,10,**30**) | (20,20,20,**10**) | (20,20,20,**50**) | (20,20,20,**50**) |
| $u_{S_i} = u_{I_i} = u_{R_i}$ | 4 | 4 | 4 | 5 | 5 | 5 |
| $\delta$ | 1 | 1 | 1 | 1 | 1 | 1 |

**Table S7**: Parameter values for Figure 4

| **Parameter** | Panel A | Panel D | Panel B,C | Panel E,F |
|---|---|---|---|---|
| $r_i$ | 1 | 1 | 1 | 10 |
| $b_i$ | 1 | 1 | 1 | 1 |
| $\alpha_{ij}$ | Matrix $\mathbf{M}_A$ | Matrix $\mathbf{M}_D$ | Matrix $\mathbf{M}_{BC}$ | Matrix $\mathbf{M}_{EF}$ |
| $e_{ij}$ | 1 | 1 | 1 | 1 |
| $m_i$ | 0 | 0 | 0.1 | 0 |
| $(\beta_1, \beta_2, \beta_3, \beta_4)$ | (3,4,5,6) | (9,4,3,1) | (16,2.3,1.6,1.3) | (4.5,2.5,2,1.5) |
| $\mu_i$ | 1 | 1 | (2,0.3,0.3,0.3) | 0.25 |
| $\nu_i$ | 0 | 0 | 0.1 | 0.1 |
| $xi_i$ | 0 | 0 | 0.1 | 0.1 |
| $\gamma_i$ | 0 | 0 | 1 | 1 |
| $(\chi_1, \chi_2, \chi_3, \chi_4)$ | (1,1,1,1) | (1,1,1,1) | (10, 12, 14, 16) | (15,2,1.5,1) |
| $u_{S_i} = u_{I_i} = u_{R_i}$ | (1,1,1,1) | (1,1,1,1) | 20 | 20 |
| $\delta$ | 1 | 1 | 40 | 40 |

For all black curves, there is no interspecific competition, which means $\alpha_{ij} = 0$ for $i \neq j$. The intraspecific competition coefficients ($\alpha_{ii}$) for the black curves and all competition coefficients for the blue curves are given by the entries of the matrices

$$\mathbf{M}_A = \begin{pmatrix} 0.5 & 1.5 & 3 & 5 \\ 0.2 & 1.8 & 3 & 5 \\ 1.4 & 0.5 & 3.1 & 5 \\ 1 & 1 & 2 & 6 \end{pmatrix}, \mathbf{M}_D = \begin{pmatrix} 1 & 1.6 & 0.029 & 0.34 \\ 0.25 & 1.9 & 0.44 & 5.2 \\ 0.5 & 0.17 & 1 & 3.8 \\ 0.9 & 0.4 & 0.2 & 2 \end{pmatrix}, \mathbf{M}_{BC} = \begin{pmatrix} 1 & 0.72 & 0.6 & 0.6 \\ 0.72 & 1 & 0.6 & 0.6 \\ 0.6 & 0.6 & 1 & 0.6 \\ 0.6 & 0.6 & 0.6 & 1 \end{pmatrix},$$

$$\mathbf{M}_{EF} = \begin{pmatrix} 1 & 1.6 & 0.029 & 0.34 \\ 0.25 & 2 & 0.44 & 5.2 \\ 0.5 & 0.17 & 1 & 3.8 \\ 0.9 & 0.4 & 0.2 & 2 \end{pmatrix}.$$

**Table S8**: Parameter values for Figure 5

| **Parameter** | Panels A-C (top row) | | | Panels D-F (bottom row) | | |
|---|---|---|---|---|---|---|
| | Black | Blue | Red | Black | Blue | Red |
| $r_i$ | 0.25 | 0.25 | 0.25 | 1.1 | 1.1 | 1.1 |
| $b_i$ | 0.2 | 0.2 | 0.2 | 0.2 | 0.2 | 0.2 |
| $\alpha_{ii}$ | 1 | 1 | 1 | 1 | 1 | 1 |
| $\alpha_{ij}$ for $j \neq 4$ | 0 | 0 | 0 | 0.3 | 0.3 | 0.3 |
| $\alpha_{i4}$ for $i \neq 4$ | 0 | 0 | 0 | **0.3** | **0.6** | **0.3** |
| $e_{ij}$ | 1 | 1 | 1 | 1 | 1 | 1 |
| $m_i$ | 0 | 0 | 0 | 0.1 | 0.1 | 0.1 |
| $(\beta_1, \beta_2, \beta_3, \beta_4)$ | (1,1,1,1) | (1,1,1,1) | (1,1,1,1) | (1,1,1,1) | (1,1,1,1) | (1,1,1,1) |
| $\mu_i$ | 0.1 | 0.1 | 0.1 | 0.25 | 0.25 | 0.25 |
| $\nu_i$ | 0.1 | 0.1 | 0.1 | 0.1 | 0.1 | 0.1 |
| $xi_i$ | 0.1 | 0.1 | 0.1 | 0.1 | 0.1 | 0.1 |
| $\gamma_i$ | 1 | 1 | 1 | 1 | 1 | 1 |
| $(\chi_1, \chi_2, \chi_3, \chi_4)$ | (10,10,10,**5**) | (10,10,10,**20**) | (10,10,10,**90**) | (100,100, 100, **100**) | (100,100, 100, **100**) | (100,100, 100, **400**) |
| $u_{S_i} = u_{I_i} = u_{R_i}$ | 20 | 20 | 20 | 100 | 100 | 100 |
| $\delta$ | 10 | 10 | 10 | 170 | 170 | 170 |
| $g_1(q)$ for $\mathcal{R}_0^C$ | $7/6 - q/6$ | $7/6 - q/6$ | $7/6 - q/6$ | 1.9970.997$q$ | 1.997−0.997$q$ | 1.997−0.997$q$ |
| $h_1(q)$ for $\mathcal{R}_0^C$ | $q$ | $q$ | $q$ | $q$ | $q$ | $q$ |
| $g_2(q)$ for $\mathcal{R}_0^C$ | $2 - q$ | $2 - q$ | $2 - q$ | $2 - q$ | $2 - q$ | $2 - q$ |
| $h_2(q)$ for $\mathcal{R}_0^C$ | -5+6$q$ | -5+6$q$ | -5+6$q$ | -0.0026+1.0026$q$ | -0.0026+1.0026$q$ | -0.0026+1.0026$q$ |
| $g_1(q)$ for $I_1^*/N_1^*$ and $I_1^*$ | 1.244−0.2441$q$ | 1.244−0.2441$q$ | 1.244−0.2441$q$ | 2.027−1.027$q$ | 2.027−1.027$q$ | 2.027−1.027$q$ |
| $h_1(q)$ for $I_1^*/N_1^*$ and $I_1^*$ | $q$ | $q$ | $q$ | $q$ | $q$ | $q$ |
| $g_2(q)$ for $I_1^*/N_1^*$ and $I_1^*$ | $2 - q$ | $2 - q$ | $2 - q$ | $2 - q$ | $2 - q$ | $2 - q$ |
| $h_2(q)$ for $I_1^*/N_1^*$ and $I_1^*$ | -3.09+4.09$q$ | -3.09+4.09$q$ | -3.09+4.09$q$ | 0.0267+0.973$q$ | 0.0267+0.973$q$ | 0.0267+0.973$q$ |

The functions $g_i(q)$ and $h_i(q)$ define the change of parameters used to convert the ET model from a FDDT form ($q = 0$) to a DDDT ($q = 2$). The change of parameters for $\mathcal{R}_0^C$ hold constant the value of $\mathcal{R}_0^C$ for the subcommunity equilibrium; see equations (S59) and (S60) of Section S1.4.3. The change of parameters for $I_1^*/N_1^*$ and $I_1^*$ hold constant the densities and frequencies at the endemic subcommunity equilibrium ($p'$); see equations (S123) and (S124) of Section S1.5.4. The same change of parameters is used in each row of Figure 4 because only parameters of the added host species (i.e., host 4) differ between the simulations.
